# Type I interferon–driven lung pathology restricts T cell accumulation and motility during tuberculosis

**DOI:** 10.64898/2025.12.05.692491

**Authors:** Maxime Caouaille, Léa Fromont, Tomoyo Shinkawa, Marion Faucher, Aizat Iman Abdul Hamid, Annie Behar, Serge Mazères, Yaël Duvergé, Emeline Decarpentrie, Yohan Lorreyte, Emmanuelle Näser, Samuel M Behar, Emma Lefrançais, Olivier Neyrolles, Denis Hudrisier

## Abstract

CD8^+^ T cells are elicited during tuberculosis yet how susceptible lung environments shape their maintenance and behavior is unclear. We examined pulmonary T cell responses to *Mycobacterium tuberculosis* in resistant control and *Sp140^-/-^* mice, a model of type I interferon (IFN-I)-driven susceptibility. Resistant mice generated diverse CD8^+^ effector- and memory-like subsets producing TNF and IFNγ, whereas *Sp140^-/-^* mice showed a broad loss of pulmonary CD8^+^ T cells. Single-cell RNA sequencing revealed altered states in CD8^+^ T cells from susceptible mice with exhaustion and IFN-I transcriptional programs. IFNAR blockade rescued CD8^+^ T cell numbers, diversity, cytokine production, reduced bacterial burden and lung pathology, with comparable benefits also valid for CD4^+^ T cells. Intravital microscopy of infected lungs further showed that T cell dynamics and motility within lesions were restricted under exuberant IFN-I signaling but fully restored by IFNAR blockade. Thus, IFN-I-driven tissue pathology restricts T cell accumulation and motility during tuberculosis.

**Teaser:** In tuberculosis, excessive type I interferon signaling reshapes the lung environment, limiting T cell accumulation and motility within infected lesions.

## Introduction

Tuberculosis (TB), caused by *Mycobacterium tuberculosis*, remains a major global health threat, with over 10.5 million new cases and over a million deaths in 2024 (*1*). It is estimated that *M. tuberculosis* has infected roughly a quarter of the world’s population (*1*). Remarkably, only 5-10% of infected individuals develop active TB disease, highlighting the overall effectiveness of human immune defenses against *M. tuberculosis*. Although considerable progress has been made in understanding TB immunity, critical questions remain regarding the mechanisms that mediate protective immune responses. Evidence from human studies and animal models, in particular mice and non-human primates (NHPs), has shown that CD4^+^ T cells are essential for TB control (*2–4*), and excessive neutrophil recruitment into the lungs is detrimental (*5–8*). However, variations in CD4^+^ T cell and neutrophil responses do not fully account for differences in disease outcomes, nor has analysis of CD4^+^ T cells provided sufficient predictive or therapeutic value for vaccine development (*9*, *10*). Thus, understanding the contributions of additional immune cell subsets to protection against TB remains a key goal.

The role of CD8^+^ T cells in immunity to *M. tuberculosis* infection in mice is more complicated than that of CD4^+^ T cells, and this has been the subject of some controversy over the years. Early studies in mice showed that deficiency of total CD8^+^ T cells (e.g. CD8α^-/-^), or of CD8^+^ T cells restricted by classical MHC class I molecules (β2m^-/-^), increases susceptibility to *M. tuberculosis* (*11–14*); however, antibody-mediated depletion experiments suggested a relatively modest role for conventional CD8^+^ T cells during acute infection (*11*, *15*). By contrast, depletion of CD8^+^ T cells during latent infection in mice or in *M. tuberculosis*-infected old mice increased bacterial burden in the lungs, whereas depletion of CD4^+^ T cells had little effect (*15*, *16*). Studies in NHPs demonstrated the importance of CD8α^+^ cells both in natural immunity and in the protection afforded by intravenous BCG vaccination, an effect that is likely mediated by multiple CD8α^+^ populations, predominantly innate lymphoid cells (*17–20*). In humans and experimentally infected animals, *M. tuberculosis* elicits robust CD8^+^ T cell responses that correlate with bacterial control (*21–27*). In addition, adoptive transfer experiments showed that CD8^+^ T cells can restrict *M. tuberculosis in vivo* (*28*). These CD8^+^ T cells contribute to protection through multiple mechanisms, including direct cytotoxicity, cytokine production, and regulation of inflammatory responses. In mice, polyclonal, infection-elicited, CD8^+^ T cells can restrict *M. tuberculosis in vitro* (*29*) and *in vivo* (*11*, *30–32*), and vaccine-induced CD8^+^ T cell responses enhance pulmonary TB control (*33*, *34*). Collectively, these studies highlight the contribution of CD8^+^ T cells to optimal control of *M. tuberculosis* infection (*4*, *31*, *35–37*).

CD8^+^ T cells consist of diverse subsets defined by their tissue localization, migratory property, effector functions, and lifespan, including tissue-resident memory, effector, and central memory populations (*38–46*). The contribution of dedicated memory CD8^+^ T cell subsets in various acute and chronic pathologies has been well established (*47*, *48*). In TB, although both human and animal studies have demonstrated a protective role for CD8^+^ T cells, the differences in the responses of these cells in susceptible and resistant hosts, particularly within the lungs, remain poorly understood. Notably, the ability of CD8^+^ T cells to traffic to infected lesions and exert effector functions may be shaped by host genetics and the local inflammatory environment, and elucidating how these factors regulate CD8^+^ T cell responses is important to advance our understanding of TB immunity and potentially to inform novel immunomodulatory strategies.

A growing body of evidence shows that the inflammatory environment, particularly cytokine signaling and neutrophil activity, plays a critical role in shaping T cell dynamics in TB (*5*, *8*, *49–51*). Type I interferons (IFN-I) in particular have been implicated in TB pathogenesis, with excessive IFN-I signaling associated with worsened disease outcomes in humans (*52*, *53*) and experimental models (*54–56*). Excessive IFN-I signaling has been shown to modulate the relative abundance of immune cells populations in the lungs of TB-infected mice (*5*, *8*, *57–59*). Two recent studies showed that IFN-I signaling drives a pathological accumulation of neutrophils in lungs of TB-susceptible C3HeB/FeJ mice, precluding the infiltration of protective CD4^+^ T cells into lesions. Depletion of neutrophils or blockade of IFN-I signaling in these mice reversed the relative abundance of neutrophils and CD4^+^ T cells in lung lesions and decreased bacterial burden (*6*, *7*). Notably, CD8^+^ T cell numbers were also reduced in C3HeB/FeJ mice (*6*), potentially contributing to their susceptibility to *M. tuberculosis*. The mechanisms underlying the effects of IFN-I on T cell responses in TB remains incompletely understood.

Sp140 is a key regulator of IFN-I responses that is associated with immune-mediated diseases and bacterial infection, such as *Legionella pneumophila* and *M. tuberculosis* in mice (*8*, *60–62*). Located within the super-susceptibility to tuberculosis-1 (*sst1*) locus, Sp140 has recently been shown to restrain IFN-I responses during infection (*8*, *60–62*), with mice lacking Sp140 exhibiting exacerbated IFN-I production and heightened susceptibility to *M. tuberculosis*, phenotypes that was completely reversed upon genetic ablation of IFN-I signaling (*60*).

Here, we investigate how the inflammatory microenvironment shapes pulmonary CD8^+^ T cell responses during aerosol *M. tuberculosis* infection in mice in the presence and absence of *Sp140*. Using high-dimensional cytometry, single-cell transcriptomics, and intravital imaging, we show that unrestrained IFN-I signaling results in a collapse of CD8^+^ T cell numbers and function and impairs T cell motility within TB lesions in susceptible Sp140-deficient mice. Because CD4^+^ T cell numbers were likewise reduced in *Sp140^-/-^* mice and rescued by IFNAR blockade, our findings uncover a previously unrecognized role for Sp140 in safeguarding adaptive immunity by limiting IFN-I-mediated pathology and identify additional IFN-I-dependent barriers that impede T cell dynamics in TB lesions.

## Results

### Profound dysregulation of pulmonary CD8^+^ T cells in *Sp140*^-/-^ mice

To investigate the contribution of CD8^+^ T cells during *M. tuberculosis* infection, we compared pulmonary T cell responses in resistant (*Sp140*^+/+^) and susceptible (*Sp140*^-/-^) mice following aerosol infection with the Erdman strain of *M. tuberculosis* (*60*). *Sp140*^+/+^ mice survived beyond day 56 while *Sp140*^-/-^ mice succumbed to infection before day 28, with uncontrolled bacterial replication and severe lung pathology with necrotic lesions, necessitating euthanasia for ethical reasons (Supplemental Figure 1). Using a 27-antibody spectral flow cytometry panel that included a fluorescently labeled anti-CD45 antibody injected intravenously 5 minutes before sacrifice to distinguish cells residing within the lung parenchyma (CD45iv^-^) from those within the vasculature (CD45iv^+^) (*63*), we observed a trend toward reduced CD8^+^ T cells numbers in *Sp140*^-/-^ mice between day 21 post-infection and later time points (Figure 1A). Unsupervised analysis revealed a robust diversity of CD8^+^ T cells subsets within the CD8^+^ T cell compartment in control mice, which included naïve and activated cells as well as cell displaying classical phenotypic markers of memory cells (referred to as “memory-like” throughout the manuscript since the pathogen is never eliminated in our context), most of which expressed TCRβ and CD8αβ (Supplemental Figure 2). Unsupervised clustering analysis was not possible in *Sp140*^-/-^ mice due to the low numbers of CD8^+^ T cells in the lungs of these mice (Figure 1A).

**Figure 1.**
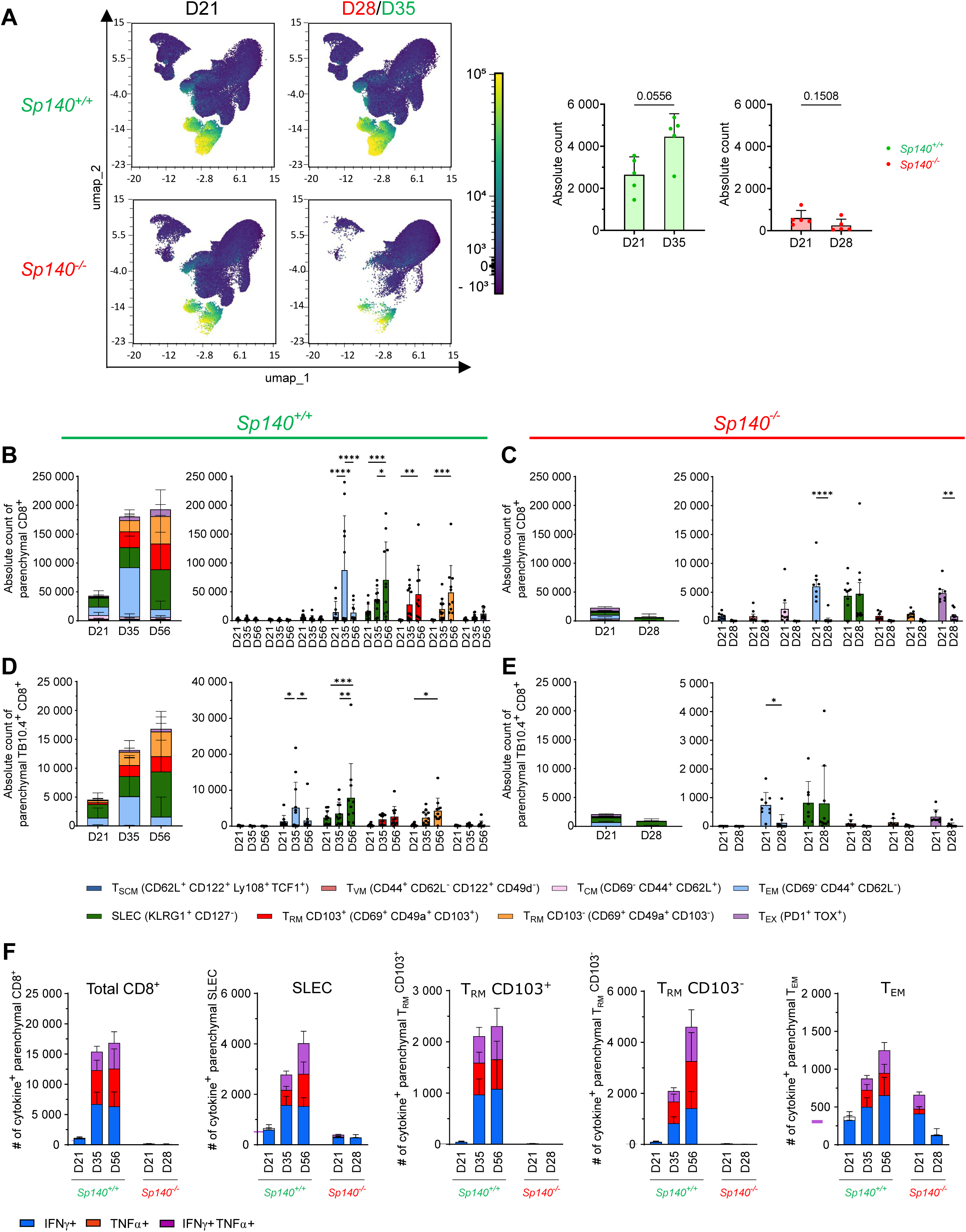
Profound dysregulation of pulmonary CD8^+^ T cells in *Sp140*^-/-^ mice. *Sp140^+/+^* and *Sp140^-/-^* mice were infected by aerosol route with 100 CFUs of the Erdman strain of *M. tuberculosis* and, at the indicated time points, were sacrificed and their lungs analyzed by flow cytometry. A. UMAP representation of live lung cells based on spectral cytometry data, with CD8 expression displayed as a continuous color scale. Samples include equal numbers of cells from *Sp140^+/+^* (days 21 (n=5) and 35 (n=5)) and *Sp140^-/-^* (days 21 (n=5) and 28 (n=5)) mice. Quantification of the CD8-expressing cell cluster is shown (right). Each data point corresponds to one individual mouse. Bars and error bars indicate the mean ± standard deviation. Non parametric Mann-Whitney test was used. B. Absolute numbers of the main parenchymal lung CD8^+^ T cell subsets in *Sp140^+/+^* mice at days 21 (n=11, mean number of total CD8^+^ T cell: 100,184), 35 (n=12, mean number of total CD8^+^ T cell: 368,332), and 56 (n=11, mean number of total CD8^+^ T cell: 296,671) post-infection. Stacked (left) and individual subpopulations (right) representations are shown. Each data point corresponds to one individual mouse. Bars and error bars indicate the mean ± standard deviation. Data are pooled from two independent experiments, and unpaired two-way ANOVA tests with Tukey correction were used. C. As in B) for *Sp140^-/-^* mice at days 21 (n=8, mean number of total CD8^+^ T cell: 52,775) and 28 (n=10, mean number of total CD8^+^ T cell: 13,906) post-infection. D. As in B) but for parenchymal TB10.4-specific CD8^+^ T cell subsets in *Sp140^+/+^* mice at days 21 (mean number of total TB10.4-specific CD8^+^ T cell: 6,257), 35 (mean number of total TB10.4-specific CD8^+^ T cell: 21,047), and 56 (mean number of total TB10.4-specific CD8^+^ T cell: 22,494) post-infection. Stacked (left) and individual subpopulations (right) representations are shown. E. As in D) but for *Sp140^-/-^* mice at days 21 (n=8, mean number of total CD8^+^ T cell: 2,388) and 28 (n=10, mean number of total CD8^+^ T cell: 1,587) post-infection. The color code for individual CD8^+^ T cell subsets, presented below, is common to all panels B-E. F. Absolute numbers of total lung parenchymal CD8^+^ T cells and CD8^+^ T cell subpopulations producing IFNγ, TNFα, or both after stimulation with *M. tuberculosis* peptides. Data for *Sp140^+/+^* mice are shown at days 21 (n=11), 35 (n=12), and 56 (n=11) (left) and for *Sp140^-/-^* mice at days 21 (n=8) and 28 (n=10) (right). Bars and error bars indicate the mean ± standard deviation.

To assess CD8^+^ T cell subsets in *Sp140*^-/-^ mice, we established a gating strategy using both surface and intracellular markers (Supplemental Figure 3) that allowed us to distinguish effector and memory-like CD8^+^ T cell subpopulations and characterize their kinetics in the lungs over the course of infection. We focused on CD8^+^ T cell subsets with well-characterized roles in various pathological contexts (*38*, *39*, *41*, *64*), including short-lived effector (SLEC), exhausted (T_EX_), stem cell memory-like (T_SCM_), central memory-like (T_CM_), effector memory-like (T_EM_), virtual memory-like (T_VM_), and resident memory-like (T_RM_) cells, with T_EM_ cells comprising two subpopulations: “true” T_EM_ and peripheral memory-like T cells (T_PM_) (*65*). We noted much fewer CD8^+^ T cells in *Sp140*^-/-^ mice compared with controls. At day 21 post-infection, *Sp140*^+/+^ mice harbored a diverse array of activated and memory-like CD8^+^ T-cell populations in the lung parenchyma, predominated by SLEC, T_CM_, and T_EM_ subsets, with minimal representation by T_EX_, T_RM_, T_SCM_, and T_VM_ subsets (Figure 1B). CD8^+^ T cells were present in limited numbers in the lungs of *Sp140*^-/-^ mice (Figure 1C) but, when distributed across subsets, the overall diversity was broad and resembled *Sp140*^+/+^ mice at day 21 (Supplemental Figure 4A). Notably, T_EX_ cells were significantly enriched in *Sp140*^-/-^ mice but virtually absent in *Sp140*^+/+^ mice at day 21 (Figure 1B, C).

The differences in CD8^+^ T cell composition in the lung parenchyma were more pronounced at later time points, with cell numbers increasing substantially in wild-type mice by day 35 and both CD103^+^ and CD103^-^ T_RM_ cells becoming more prominent (Figure 1B, C). Within the T_EM_ subset in wild-type mice, T_PM_ cells predominated within the lung parenchyma, whereas “true” T_EM_ cells were more abundant in the pulmonary microvasculature (Supplemental Figure 4B). Consistent with the spectral flow cytometry data, more than 90% of CD8^+^ T cells in the lungs expressed TCRβ and CD8αβ (Supplemental Figure 4C), corresponding to conventional CD8^+^ T cells. T cell numbers in *Sp140*^-/-^ mice, by contrast, declined between day 21 and day 28, accompanied by a collapse in subset diversity, with nearly all populations disappearing and only a small number of SLEC and T_EX_ cells persisting (Figure 1C). Results were similar when analyzing TB10.4-specific CD8^+^ T cells identified using an H-2K^b^/TB10.4 tetramer (Figure 1D, E and Supplemental Figure 4D) (*66*). The CD8^+^ T cell compartment in the lung vasculature of *Sp140*^-/-^ mice showed a similar decrease in numbers and subset diversity as seen in the parenchyma, which was distinct from wild-type mice (Supplemental Figure 4E-F). These results show that although CD8^+^ T cell responses are effectively initiated and traffic to the lungs in both mouse strains, their numbers are dramatically lower and progressively wane over the course of infection in *Sp140*^-/-^ mice. Confirming these results, co-transfer of Sp140-competent and -deficient T cells in *M. tuberculosis*-infected *Rag2^-/-^*mice revealed the lack of direct impact of Sp140 deficiency in T-cell activation and accumulation in lung parenchyma (Supplemental Figure 4G, H). In *Sp140^-/-^* mice, parenchymal CD4^+^ T cells and B cells were also reduced, whereas neutrophils were increased and other myeloid populations were not significantly altered, indicating a broader shift in pulmonary immune composition rather than a selective CD8^+^ T cell defect (Supplemental Figure 4I-K).

To measure CD8^+^ T cell function, we assessed cytokine production after *ex vivo* stimulation with a cocktail of CD4^+^ and CD8^+^ T cell antigens on the H-2^b^ background (Table S1 and Supplemental Figure 5A). In *Sp140*^+/+^ mice, multiple parenchymal CD8^+^ T cell subpopulations produced TNFα and/or IFNγ, with T_RM_ (CD103^+^ plus CD103^-^) being proportionally overrepresented among cytokine producers (Figure 1F and Supplemental Figure 5A, B). As expected, T_EX_, T_CM_, T_VM_, and T_SCM_ subsets did not produce cytokines (Supplemental Figure 5A, B). In the lung vasculature, TNFα and IFNγ were mostly produced by SLEC and T_EM_ populations (Supplemental Figure 5C). In *Sp140*^-/-^mice, T_EM_ and SLEC were the only CD8^+^ T cell subsets that produced TNFα and/or IFNγ at day 21, and cytokine production decreased over time, becoming nearly undetectable by day 28 (Figure 1F, Supplemental Figure 5C, D). Similarly, only modest cytokine production was detected in the vasculature from SLEC and T_EM_ cells (Supplemental Figure 5D). Together, these findings reveal a profound dysregulation of the CD8^+^ T cell compartment in *Sp140*^-/-^ mice, with severely decreased cell numbers, collapse of phenotypic diversity, and impaired effector cytokine production. These results indicate that SP140 preserves CD8^+^ T cell responses during TB, and its absence results in a limited T cell accumulation and function within TB lesions.

### CD8^+^ T cells from *Sp140*^-/-^ mice display signatures of type I IFN and exhaustion pathways

To explore the unexpected differences in CD8^+^ T cell subpopulation diversity between *Sp140*^+/+^ and *Sp140*^-/-^ mice in an unbiased manner, we performed a global scRNA-seq analysis of CD8^+^ T cells to gain deeper insight into their phenotypic and functional heterogeneity and to identify potential mechanisms underlying these disparities. To recover sufficient cells, mice were infected with a lower inoculum (30 CFUs); this diminished the strain-differences in CD8^+^ T cell numbers and diversity observed at 100 CFUs (Supplemental Figure 6A), but *Sp140*^-/-^ mice still developed a severe disease trajectory across multiple independent experiments (Supplemental Figure 6B). scRNA-seq performed on parenchymal CD8^+^ T cells (Supplemental Figure 6C) confirmed substantial heterogeneity among lung-infiltrating CD8^+^ T cells in both genotypes (Figure 2A), with eleven clusters shared across both genotypes but with distinct relative proportions (Figure 2B-D). A small fraction of myeloid cells (cluster 9) was detected but were considered to be contaminants because they lacked *Cd8* expression (Table S2). Functional gene-expression patterns (Figure 2E) revealed an activation continuum across clusters 0-2, with expression of genes associated with activation, polyfunctionality, and exhaustion progressively increasing across clusters. Cluster 0 occurred at similar frequencies in both genotypes, whereas clusters 1 and 2 were enriched in CD8^+^ T cells from *Sp140*^-/-^ mice (Figure 2B, D). These clusters also expressed high levels of cytokines and cytotoxic molecules, a signature associated both with protective capacity and functional impairment due proteotoxic stress (*67*), the latter seeming more consistent with our data. As expected, the IFN-I response, concentrated in cluster 5, was enriched in CD8^+^ T cells from *Sp140*^-/-^ mice, whereas naive-like clusters 4 and 7 were enriched in CD8^+^ T cells from *Sp140*^+/+^ mice (Figure 2D, E). Cluster 3 remained after QC filtering and expressed tissue-residency-associated markers such as *Itgae* and *Itga1*, but also displayed relatively elevated mitochondrial transcript abundance, consistent with stressed or lower-quality cells. Accordingly, this cluster was retained for transparency but was not used as a primary basis for biological interpretation. The presence of *P2rx7* transcripts is compatible with the preferential loss of P2X7R^+^ T_RM_ cells under exposure to tissue damage (Table S2) (*68*). Cluster 6 comprised proliferating CD8^+^ T cells, while clusters 8 and 10 contained rare γδT cells and Tc17 cells (Figure 2A).

**Figure 2.**
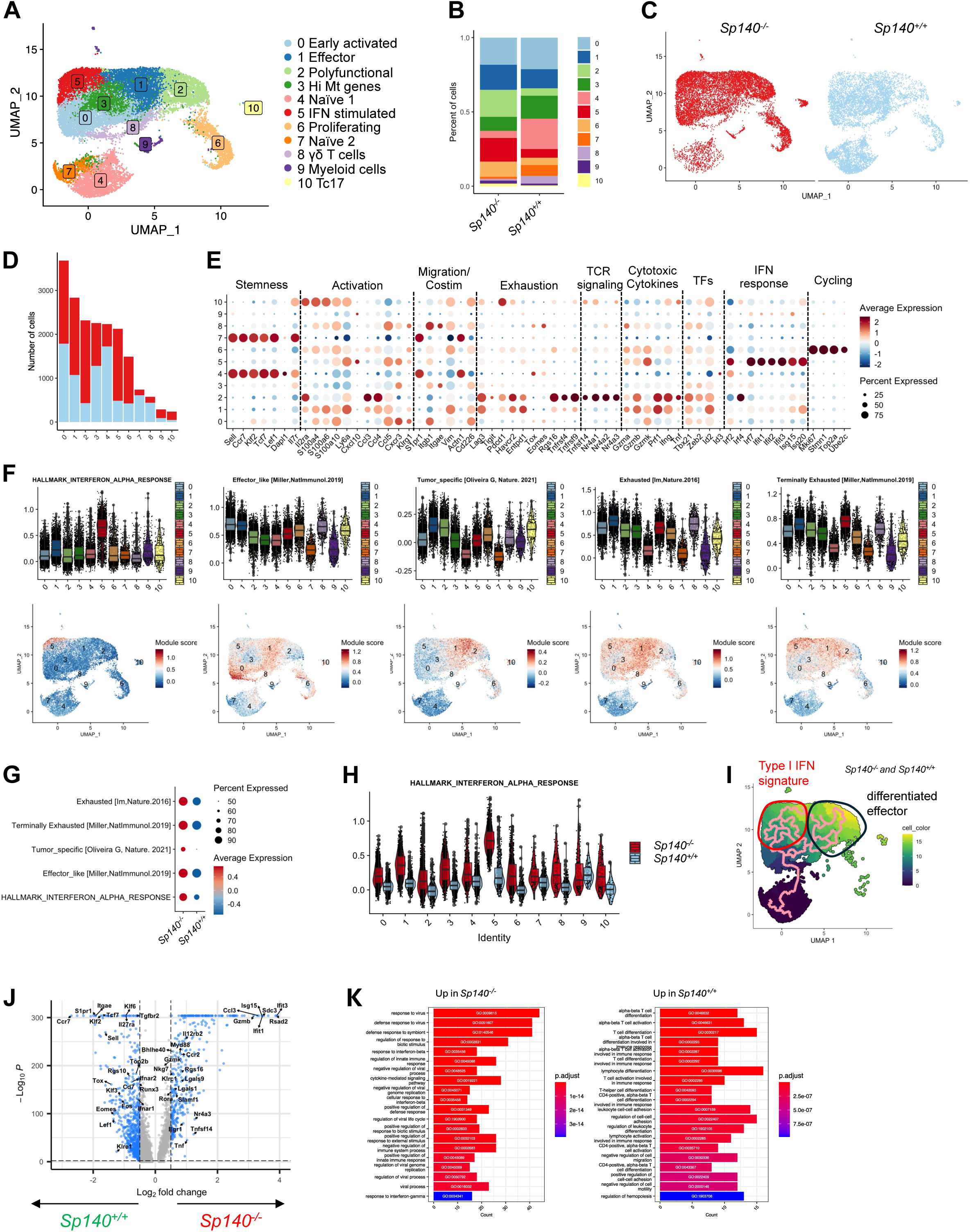
CD8^+^ T cells from *Sp140*^-/-^ mice display signatures of type I IFN and exhaustion pathways. A. UMAP visualization of scRNAseq data of lung CD8^+^ T cells from *Sp140^+/+^* and *Sp140^-/-^* mice infected with *M. tuberculosis* for 35 and 28 days, respectively. B. A stacked bar plot depicts the cluster distribution in CD8^+^ T cells from *Sp140^+/+^* and *Sp140^-/-^* mice. C. Distribution of CD8^+^ T cells from *Sp140^+/+^* (blue) and *Sp140^-/-^* (red) mice on the UMAP. D. A stacked bar plot depicts the sample distribution in CD8^+^ T cells from *Sp140^+/+^* (blue) and *Sp140^-/-^* (red) mice. E. Dot plot of selected function-associated genes among the CD8^+^ T cell clusters. F. Violin plots depict the enrichment score of indicated CD8^+^ T cell signatures from published studies, and UMAP is colored based on the enrichment scores. G. Dot plot showing the enrichment scores of indicated CD8^+^ T cell signatures from published studies in CD8^+^ T cells from *Sp140^+/+^* and *Sp140^-/-^* mice. H. Violin plot showing the enrichment score of IFN-I response in individual cluster of CD8^+^ T cells from *Sp140^+/+^* and *Sp140^-/-^* mice. I. Single-cell trajectories constructed using Monocle3. UMAP shows cells colored by pseudotime values along the trajectory (pink line). Clusters 6, 9, and 10 are excluded from the analysis. J. Volcano plot of DEGs between CD8^+^ T cells from *Sp140^+/+^* and CD8^+^ T cells from *Sp140^-/-^* mice. Genes with log2FC >0.5 and an adjusted p value < 0.01 are designated by a blue-filled circle. K. GSEA of differentially expressed genes in CD8^+^ T cells from *Sp140^-/-^* (left) versus CD8^+^ T cells from *Sp140^+/+^* (right) mice cells with log2FC>0.5 and adjusted P value < 0.01 using MSigDB immunologic signature gene sets (C7). Adjusted P values are denoted in the bar.

To refine cluster identities, we projected published gene-set signatures (*69–71*) onto our data. Interferon-α, tumor-response, and exhaustion signatures scored highest in clusters 1, 2, and 5, which were overrepresented in CD8^+^ T cells from *Sp140*^-/-^ mice, suggesting a link between Sp140 deficiency, heightened exhaustion and IFN-I-driven transcriptional programs in CD8^+^ T cells (Figure 2F). Enrichment analyses using external CD8^+^ T cell datasets (*70*) corroborated this conclusion, with *Sp140^-/-^* profiles strongly enriched for effector-like, exhaustion, and IFN-I-responsive genes (Figure 2G and Supplemental Figure 6D). Reference mapping against CD8^+^ T cell states from acute (Armstrong) versus chronic (clone 13) LCMV infection (*72*) again recapitulated the pattern: *Sp140^-/-^* CD8^+^ T cells aligned with chronic effector-like, exhausted, and ISG states, whereas control cells aligned with acute naïve and effector states (Figure 2H).

Pseudotime analysis using Monocle3 further supported this model, positioning clusters 2 and 5 at terminal endpoints associated with exhaustion- and IFN-I-enriched programs, respectively (Figure 2I). Differential gene-expression and Gene Ontology analyses across the full dataset showed that *Sp140^-/-^* CD8^+^ T cells were skewed toward distinct IFN-I/antiviral response and exhaustion pathways (Supplemental Figure 6E, Table S3), whereas *Sp140^+/+^* cells were enriched for immune-activation and differentiation programs (Figure 2J-K). Collectively, these data reveal profound transcriptional reprogramming of CD8^+^ T cells in the Sp140-deficient environment, driving them toward IFN-I-skewed, exhausted states and providing a mechanistic explanation for the impaired CD8^+^ T cell response observed in *Sp140*^-/-^ mice upon *M. tuberculosis* infection.

### IFN-I drives deficient responses in both CD8^+^ and CD4^+^ T cells and high bacterial loads in *Sp140*^-/-^ mice

IFN-I expression is a hallmark of *Sp140*^-/-^ mice infected with *M. tuberculosis* and has been reported to profoundly reshape myeloid cell responses (*8*, *60*). Consistent with this, several T cell clusters that are unique to, or markedly overrepresented in, *Sp140*^-/-^ mice exhibit a strong transcriptional signature typical of a type I IFN response (Figure 2). To evaluate the impact of IFN-I on T cell responses, we administered a blocking anti-IFNAR monoclonal antibody to *Sp140*^-/-^ mice. Consistent with previous reports (*50*, *60*, *73*, *74*), anti-IFNAR treatment markedly reduced bacterial loads in *Sp140*^-/-^ mice at day 28, restoring them to levels comparable to those in *Sp140*^+/+^ mice (Figure 3A). This was accompanied by substantial improvement in lung pathology and stabilization of body weight in anti-IFNAR-treated *Sp140*^-/-^ mice (Figures 3B, C). Neutrophil extracellular trap formation (NETosis) is amplified by IFN-I and has been shown to be detrimental in TB infection (*5*, *8*, *59*). Histone citrullination (cit-H3), a marker of NETosis, was markedly enhanced in *Sp140*^-/-^ mice but reverted to levels comparable with *Sp140*^+/+^ control mice following anti-IFNAR treatment (Figure 3D). IFNAR blockade also restored CD8^+^ T cell numbers in *Sp140*^-/-^ mice and promoted their infiltration into lung near bacteria-containing lesions (Figure 3D). In marked contrast, CD8^+^ T cells were sparse and remained distant from both bacteria-rich and cit-H3-rich lesions in untreated *Sp140*^-/-^ mice, suggesting that IFN-I-associated lesion remodeling is linked to altered distribution and motility of CD8^+^ T cells within infected lung tissue (Figure 3D). Importantly, IFN-I blockade restored CD8^+^ T cell responses, rescuing both cell numbers and phenotypic diversity (Figure 3E), with similar results observed when gating on *M. tuberculosis*-specific CD8^+^ T cells (Supplemental Figure 7A). This restoration was evident across both supervised and unsupervised analyses, and in both the lung parenchyma and vasculature (Supplemental Figure 7B-D). Importantly, CD4^+^ T cell underwent the same collapse in *Sp140^-/-^* mice and were fully restored upon anti-IFNAR treatment (Supplemental Figure 7E). Cytokine production by CD8^+^ T cells was also fully restored following anti-IFNAR treatment (Figure 3F). In contrast, administration of anti-IFNAR to *Sp140*^+/+^ mice had no effect on bacterial loads, body weight loss, or T cell responses (Supplemental Figure 8A-D). Finally, this approach confirmed our scRNAseq study showing a link between IFN-I signature and increased exhaustion on both CD8^+^ and CD4^+^ T cells (Supplemental Figure 8E). Together, these results indicate that *Sp140* deficiency limits CD8^+^ T cell accumulation, positioning, and functional potential within TB lesions, in association with IFN-I-dependent remodeling of the infected lung environment.

**Figure 3.**
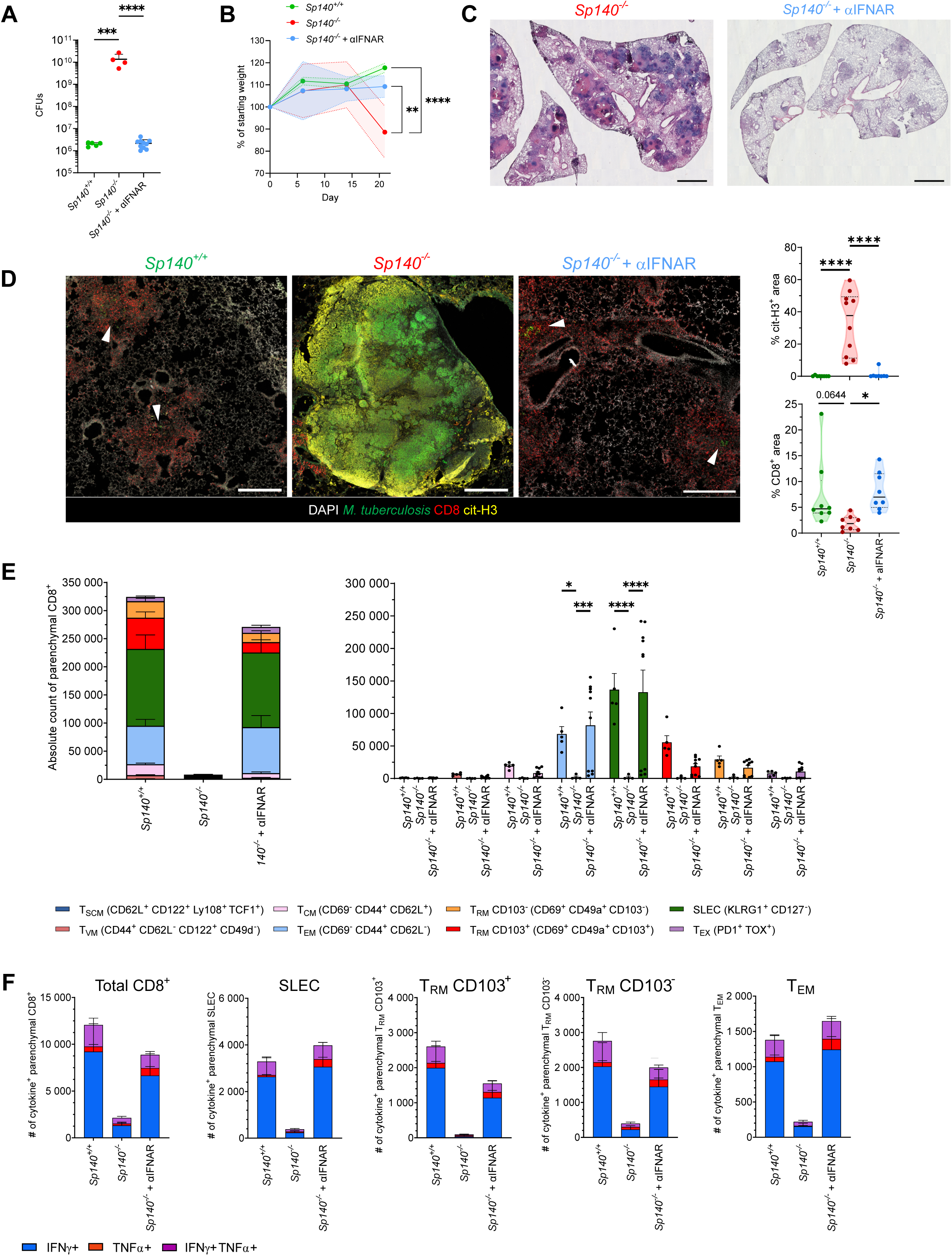
IFN-I drives deficient responses in both CD8^+^ and CD4^+^ T cells and high bacterial loads in *Sp140*^-/-^ mice. *Sp140^-/-^* mice infected with 100 CFUs of the Erdman strain of *M. tuberculosis* were treated with or without a blocking mAb against IFNAR between days 7 and 27 post-infection, and then their lungs were collected and analyzed. Untreated *Sp140^+/+^* mice were used for comparison. A. Pulmonary bacterial loads at 28 days post-infection in *Sp140^+/+^* (n=5), untreated *Sp140^-/-^* (n=4), and anti-IFNAR-treated *Sp140^-/-^* mice (n=10). A representative experiment is shown. Each data point corresponds to one individual mouse. Bars and error bars indicate the mean ± standard deviation. One-way ANOVA tests with Tukey correction was used. B. Weight monitoring from day 0 to day 21 in the same groups of mice. Body weight was monitored over time and expressed as a percentage of the initial body weight (baseline set to 100%). The shaded area corresponds to the standard deviation. One-way ANOVA tests with Tukey correction was used at day 21. C. Representative hematoxylin and eosin staining of lung sections (20 µm) from untreated and anti-IFNAR-treated *Sp140^-/-^* mice at 28 days post-infection. Scale bar, 1000 µm. D. Representative confocal images of lung sections (20 µm) from *Sp140^+/+^* mice (day 35 post infection, left), untreated *Sp140^-/-^* mice (day 28 post infection, middle), and anti-IFNAR-treated *Sp140^-/-^* mice (day 28 post infection, right) infected with *M. tuberculosis*-GFP (bacterial foci indicated by white arrows). CD8^+^ T cells are shown in red, NETs (citrullinated histone H3) in yellow, and nuclei (DAPI) in white. Scale bar, 500 µm. Quantification of NETs and CD8^+^ T cell areas is shown (right). Each data point corresponds to measurements taken from distinct microscopic fields of view from at least 3 different mice per group. One-way ANOVA tests with Tukey correction was used. E. Representation in absolute numbers of the main parenchymal lung CD8^+^ T cells subpopulations induced during *M. tuberculosis* infection of *Sp140^+/+^* (n=5, mean number of total CD8^+^ T cell: 498,063) and *Sp140^-/-^* treated (n=10, mean number of total CD8^+^ T cell: 338,597) or not (n=4, mean number of total CD8^+^ T cell: 15,119) with anti-IFNAR on day 28. Stacked representation (left) and individual subpopulation (right) are shown. Bars and error bars indicate the mean ± standard deviation. Two-way ANOVA tests with Tukey correction was used. F. Absolute numbers of IFNγ, TNFα, or both producing lung parenchymal CD8^+^ T cells and main producing CD8^+^ subpopulations after stimulation with *M. tuberculosis* peptides of *Sp140^+/+^* (n=5) and *Sp140^-/-^* treated (n=10) or not (n=4) with anti-IFNAR on day 28. A representative experiment is shown alongside two independent ones. Bars and error bars indicate the mean ± standard deviation.

### The type I interferon-dominant environment impairs both CD8^+^ and CD4^+^ T cell mobility within TB lesions

The loss of the CD8^+^ T cell response in *Sp140*^-/-^ mice and its restoration following IFNAR blockade (Figures 1 and 3) mirrors previous observations in TB, in which susceptible C3HeB/FeJ mice displayed reduced pulmonary CD4^+^ T cell accumulation after infection with the virulent *M. tuberculosis* HN878 strain (*6*). As shown above, the T cell response is only minimally affected at early time points in *Sp140^-/-^* mice (Supplementary Figure 4A), whereas the detrimental effects of Sp140 deficiency (*60*) and excessive IFN-I signaling (*8*) during *M. tuberculosis* infection have been shown to act prominently through the myeloid compartment. These observations led us to test whether the impaired T cell response in *Sp140^-/-^* mice could result, at least in part, from an IFN-I-dependent alteration of the infected lung environment rather than from an obligate T cell-intrinsic defect. To address this question, we used the LiveLungTB platform (*75*), which enables intravital two-photon imaging of lungs from live, mechanically ventilated mice infected with virulent *M. tuberculosis* under BSL-3 conditions.

For this, we adoptively transferred labeled CD8^+^ T cells from *M. tuberculosis*-infected *Sp140*^+/+^ donors into *Sp140*^-/-^ recipients (Figure 4A). To optimize transfer timing, we monitored IFN-I-driven tissue damage by tracking the kinetics of cit-H3 accumulation (Supplemental Figure 9A, B), which confirmed T cell exclusion from TB lesions (*6*). Transfers were performed between days 21 and 28, when IFN-I-dependent pathology was established (Figure 3D). Total CD4^+^ T cells from the same donors were co-transferred at a 1:1 ratio with CD8^+^ T cells to provide helper signals (*29*). Parallel transfers were performed in anti-IFNAR-treated *Sp140*^-/-^ mice, which display reduced inflammation and tissue pathology (Figure 3). Twenty-four hours after transfer, intravital lung imaging was performed to evaluate T cell localization and motility within infected lungs during 60-minute time-lapse acquisitions.

**Figure 4.**
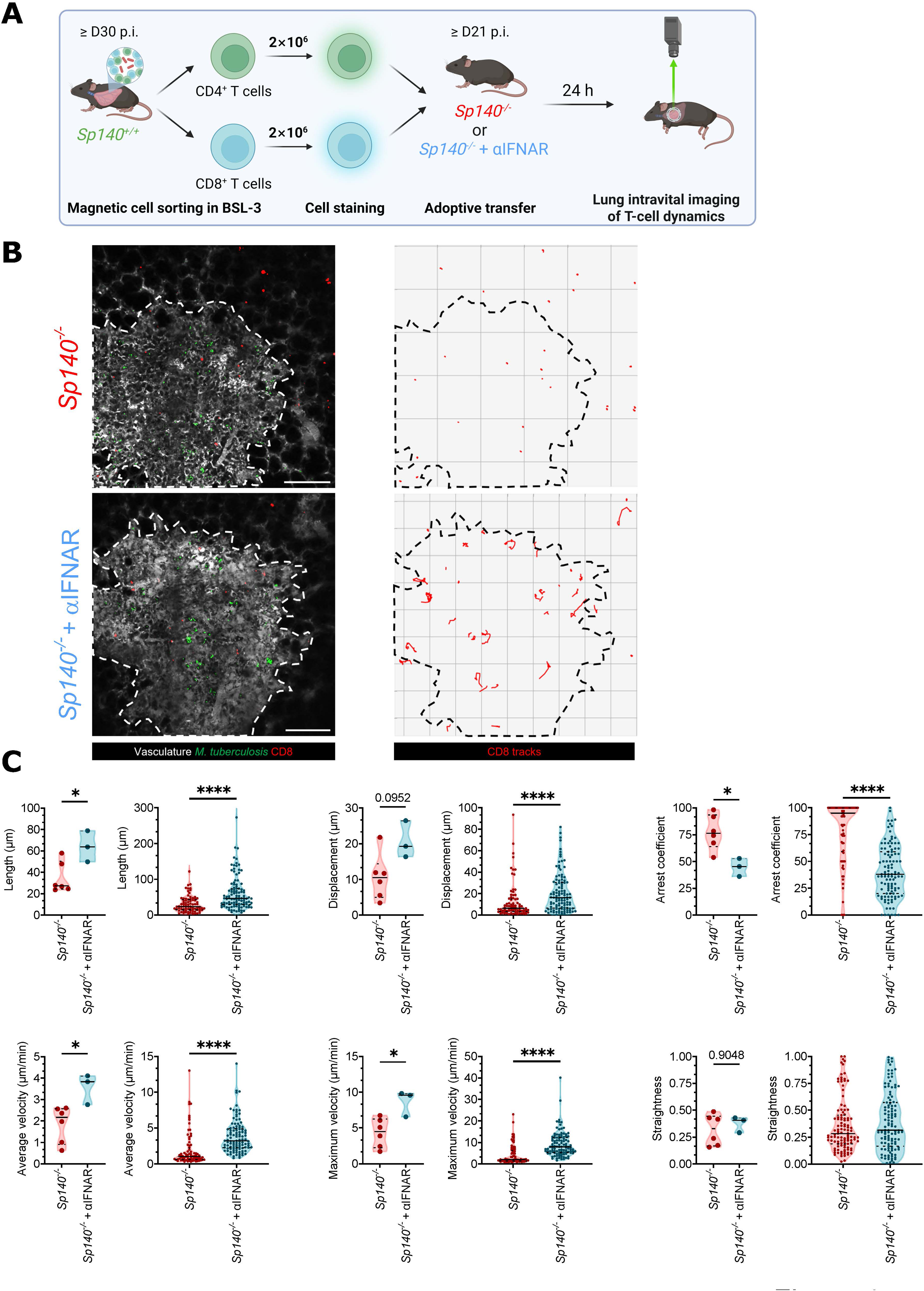
The type I interferon-dominant environment impairs both CD8^+^ and CD4+ T cell mobility and access to TB lesions. A. Schematic representation of the adoptive transfer and lung live imaging experiment. Briefly, both lung CD8^+^ and CD4^+^ T cells were isolated from infected *Sp140^+/+^* mice, labelled and adoptively transferred into *Sp140^-/-^* recipient mice treated or not with anti-IFNAR. Lung intravital imaging was performed 24 hours post-transfer. B. Representative intravital microscopy images of lungs from *M. tuberculosis*-mTurquoise-infected mice. Bacteria appear in green, transferred CD8^+^ T cells are in red, and lesion boundaries, identified by dextran staining and *M. tuberculosis* fluorescence, are outlined by a dotted line. Images show *Sp140^-/-^* mice (day 21 post infection, top) and anti-IFNAR-treated *Sp140^-/-^* mice (day 28 post infection, bottom). The first frame of the video is shown (left), with corresponding CD8^+^ T cell tracks over the full acquisition period (right). Scale bar, 200 µm. C. Quantification of CD8^+^ T cell motility parameters during intravital imaging. Per-animal motility (left) and pooled track (right) measurements for transferred CD8^+^ T cells in infected lungs are shown for each measured parameter. For per-animal motility analysis, each dot represents the mean value of the measured parameters of CD8^+^ T cell motility in a single mouse. For pooled tracks analysis, each dot represents a single tracked transferred CD8^+^ T cells. Parameters include track length, displacement, arrest coefficient (% of of time T cell instantaneous speed < 2 µm/min), average and maximum velocity and straightness. Non parametric Mann-Whitney test was used. *Sp140*^-/-^ group: n = 6 mice, 111 tracked cells in total, with 16, 6, 25, 33, 23, and 8 tracked cells per mouse. *Sp140*^-/-^ + anti-IFNAR group: n = 3 mice, 115 tracked cells in total, with 35, 23, and 57 tracked cells per mouse. Data were acquired from two independent experiments.

Albeit transferred CD8^+^ T cells successfully reached the lung parenchyma and infiltrated granulomatous lesions in *M. tuberculosis*-infected *Sp140*^-/-^ mice (Figure 4B and Supplemental Figure 9C, D), they exhibited low velocities and presented arrest coefficient close to 100%, indicating that they were essentially immobile within lesions (Figure 4C and Supplemental Movie 1). In sharp contrast, CD8^+^ T cells transferred into anti-IFNAR-treated *Sp140*^-/-^ mice efficiently accessed lesions (Figure 4B) and showed higher motility, characterized by longer track lengths and displacement, higher mean and maximum velocities, a reduced arrest coefficient and higher cumulative displacement (Figure 4C, Supplemental Figures 9E, F and Supplemental Movie 2). However, straightness was unaffected by the treatment and the value centered around 0.5 suggests that, when moving, T cells keep using a similar pattern of displacement (Figure 4C). Because CD4^+^ T cells were co-transferred in these experiments (Figure 4A) and have been reported to accumulate poorly within TB lesions in susceptible mice (*6*, *7*) and to be similarly affected in *Sp140^-/-^* mice in a IFN-I-dependent manner (Supplemental Figure 7E), we also analyzed their dynamics. Consistent with the CD8^+^ phenotype, CD4^+^ T cells in untreated *Sp140*^-/-^ mice exhibited minimal motility, whereas anti-IFNAR treatment markedly enhanced all motility parameters (Supplemental Figure 10A-C, Supplemental Movies 3 & 4). Control datasets obtained with fluorescent anti-Ly6G labeling revealed that neutrophils remain motile both inside and outside lesions of *Sp140^-/-^* mice (Supplemental Movie 5). In addition, dextran administration revealed continuous perfusion and vascular remodeling in *Sp140^-/-^* mice clearly demonstrating that the behavior of T cells is not due to tissue nonviability (Supplemental Movie 6).

Together, these findings demonstrate that type I IFN profoundly impairs T cell immunity in *Sp140*^-/-^ mice and that IFN-I-driven environmental changes are associated with altered numbers, diversity and function of endogenous T cells and altered motility of transferred T cells, with IFNAR blockade reversing both phenotypes. The data show that an IFN-I-dominated inflammatory milieu coincides with limited CD8^+^ and CD4^+^ T cell access to pulmonary lesions *in vivo* but, as shown here for the first time, also restricts their movement within these structures, revealing a potentially critical mechanism of immune evasion in TB.

## Discussion

T cells play a central and well-established role in immunity to TB in both humans and animal models. Classically, differences in T cell responses between susceptible and resistant hosts have been evaluated based on functional parameters, such as cytokine production and cytolytic capacity. Because elimination of *M. tuberculosis*-infected cells requires direct contact with T cells (*76*), understanding the molecular and spatial determinants of successful T cell access to lesions is critical. This is particularly important considering that T cells exist in various lineages and subsets, displaying not only diverse biological functions but also variable lifetimes, migration and residency patterns, and tissue-positioning properties (*38–40*, *43*, *77–79*).

Prior work identified *Sp140* as a gene within the murine *sst1* locus that confers resistance to TB by restraining deleterious type I IFN pathways (*8*, *60*). However, to date, the consequences of *Sp140* deficiency have been mostly studied in innate cells (*60*, *80*).

Our findings establish that *Sp140* deficiency profoundly impairs adaptive immunity during *M. tuberculosis* infection, with both CD8^+^ and CD4^+^ T cells excluded from lung lesions in *Sp140*-deficient mice. In *Sp140*^+/+^ mice, pulmonary CD8^+^ T cells expand robustly and differentiate into multiple effector and memory-like subsets, including T_RM_ and T_EM_ populations capable of producing cytokines (IFNγ and TNF) that are associated with protection against TB (*81–84*). In contrast, mice lacking *Sp140* showed a collapse in the numbers, subset diversity, and cytokine production of CD8^+^ T cells in the lungs by 3-4 weeks post-infection, with a substantial fraction of CD8^+^ T cells displaying an exhausted phenotype. Previous studies have shown that exhaustion of CD4^+^ T cells leads to loss of protection in models of chronic TB infection (*29*, *85*, *86*). The collapse of CD8^+^ and CD4^+^ T cells coincided with extreme susceptibility to TB in *Sp140*^-/-^ animals, which develop severe symptoms, including uncontrolled bacterial growth, tissue damage, weight loss and immunopathology. These results confirm that Sp140 is essential for innate defenses against TB and show that its expression is also essential for preserving T cell immunity, indirectly enabling long-term, T cell–mediated protection against *M. tuberculosis* infection.

Dysregulated IFN-I signaling drives the susceptibility of *Sp140*^-/-^ mice (*8*, *60*, *74*) and underlies the T cell attrition we report here. Single-cell RNA sequencing and phenotypic analysis revealed that CD8^+^ T cells from *Sp140*-deficient mice bear strong IFN-I-induced transcriptional signature and exhaustion programs, which correlate with T cell dysfunction. Several features of these signatures (e.g. upregulation of stemness genes) overlap with those seen in subsets of exhausted CD8^+^ T cells in models of melanoma and chronic viral infection (*69–71*). The cells expressing high levels of exhaustion markers, TCR signaling genes, and cytokines are distinct from the population exhibiting the strongest type I IFN signature (Figure 2E, F and Supplemental Figure 6E). Therefore, there may be two independents dysfunctional CD8^+^ T cell compartments in *Sp140^-/-^* mice: (1) CD8^+^ T cells encountering APCs that present high levels of *M. tuberculosis* antigens, and (2) CD8^+^ T cells exposed to a type I IFN–rich environment. Although it is unclear how these two pathways are interconnected, scRNA-seq analysis enabled us to distinguish these two potential CD8^+^ T cell impairments. Importantly, this phenotype appears distinct from previously described models of IFN-I induced CD8^+^ T cell impairment, including mouse tumor models in which IFN-I signaling directly induces exhaustion programs (*87*, *88*).

To avoid the marked reduction in CD8^+^ T cells observed in *Sp140*^-/-^ mice, we used a lower infectious dose to mitigate pathology and recover sufficient numbers of the cells of interest. Although this lower dose consistently produced a similar disease trajectory, including reduced CD8^+^ T cell numbers, subtle differences in immune cell recruitment may influence the interpretation of the observed phenotypes. This is particularly relevant given the magnitude of the early neutrophil response, which was recently reported to impair T cell access to infected macrophages (*7*). Because the scRNA-seq experiment was performed on pooled cells from 8-12 mice per condition and was not independently replicated, it should be interpreted as a discovery dataset rather than a replicated sample-level comparison. Accordingly, we use these data to define candidate CD8^+^ T cell states and transcriptional programs that are supported by orthogonal flow-cytometric, functional, IFNAR-blockade, histological, and imaging experiments. The rescue of immunity and reduction in bacterial loads after IFNAR blockade in Sp140-deficient mice mirrors prior findings showing that *Sp140*- or *sst1^S^*-mediated susceptibility to TB is largely driven by IFN-I (*74*). CD4^+^ T cells have a well-established role in immunity to *M. tuberculosis*; however, their abundance does not consistently correlate with protection in TB, indicating that additional immune components contribute to host defense. CD8^+^ T cells are strong candidates, supported by studies in both mice and non-human primates demonstrating context-dependent increases in susceptibility when CD8^+^ T cells are absent or depleted [(*89*) and references therein]. Although our current study did not directly asses a requirement for CD8^+^ T cells in the setting of an anti-IFNAR-mediated restoration of bacterial control, future work will incorporate CD8^+^ T cell depletion and targeted reconstitution to determine their necessity and sufficiency in this model.

Excess IFN-I has been shown to undermine host defense in TB by inducing immunosuppressive factors, including IL-10 (*90*) and IL-1Ra (*74*), and by skewing innate responses toward pathological inflammation (*5*, *8*, *54–56*, *60*, *73*, *74*, *91*). *Sp140*^-/-^ mice exhibit hallmarks of IFN-I-driven immunopathology, including a key role for increased neutrophil influx and extensive NET formation, as shown in previous studies (*7*, *8*, *52*, *53*). Blocking IFN-I signaling has been shown to reverse essentially all immune defects described in TB-susceptible mouse strains, including reversing the loss of T cells [our study in *Sp140*^-/-^ mice and the study by Naik *et al.* in *Irgm1^-/-^* mice (*73*)], and reducing neutrophil accumulation and NETosis in C3HeB/FeJ mice (*6*, *8*, *59*), indicating that IFN-I drives both T cell attrition and tissue-damaging innate responses. However, whether elevated bacterial burden and associated pathology, aside from IFN-I signaling, can also constrain T cell numbers and diversity remains untested.

IFN-I are strong and key regulators of CD8^+^ T cell responses exerting complex positive and negative effects on CD8^+^ T cells: they can deliver growth-inhibitory signals or conversely promote survival and clonal expansion of the CD8^+^ T cell pool, depending on the pathogen and/or the effector or memory status of the cells and/or the acute or chronic inflammation. At the functional level, IFN-I enhance cytotoxicity but can either up- or down-modulate IFNγ production. Finally by shaping the magnitude of the primary response, IFN-I determine the size, trafficking and effector function of the downstream memory CD8^+^ T cell compartment (*90*, *92*, *93*). It therefore came as a surprise that excessive IFN-I production in *Sp140*^-/-^ mice has a profoundly deleterious impact on all CD8^+^ T cells, regardless of their effector or memory-like status and function. Although our study mainly focused on CD8^+^ T cells, we made similar observations for CD4^+^ T cells and for B cells, indicating that both arms of the adaptive response were curtailed. These mice, like *sst1^S^* mice, have a normal T cell compartment and do not display obvious signs of altered T cell activation at steady-state (*8*, *94*). Consistent with this, IFN-I-dependent collapse of both CD4^+^ and CD8^+^ T cell responses has also been reported in *Irgm1^-/-^* mice (*73*). Previous studies failed to show a fundamental defect in T cell activation in the lymph nodes of *M. tuberculosis*-infected mice exposed to excess IFN-I either via direct cytokine administration or co-infection with a virus (*54*, *56*). Many of the TB lesions present in the lungs have been linked to IFN-I (*59*, *95*, *96*) and, in some reports, IFN-I correlated with a reduced presence of T cells in the lungs of *M. tuberculosis*-infected mice (*5*, *6*, *56*, *59*). Together, these findings support a model in which excessive IFN-I signaling in *Sp140^-/-^*mice creates a hostile infected lung environment that secondarily remodels the T cell compartment. In this model, IFN-I-driven pathology is initiated predominantly through non-T cell targets, particularly the myeloid compartment, leading to altered tissue organization, impaired T cell dynamics, and progressive acquisition of dysfunctional transcriptional states by CD8^+^ T cells. Consistent with this interpretation, macrophage-specific conditional deletion of IFNAR reduces the susceptibility of *Sp140^-/-^* mice to TB to levels comparable to those observed in fully IFNAR-deficient mice on the same genetic background (*8*). These data therefore favor a predominantly environmental model, in which the altered transcriptional and functional state of CD8^+^ T cells reflect sustained exposure to an IFN-I-dominated, inflammatory, and structurally perturbed lung environment. However, we cannot exclude that IFN-I also exerts more subtle, temporally restricted, or subset-specific intrinsic effects on CD8^+^ or CD4^+^ T cells during infection. IFN-I signaling can have divergent effects on T cells depending on the timing of exposure, differentiation state, antigenic context, and inflammatory milieu. Definitive dissection of these possibilities will require T cell lineage- or subset-specific perturbation of IFNAR signaling during *M. tuberculosis* infection.

We hypothesized that, in the absence of Sp140, unrestrained IFN-I production results in excessive inflammation and the formation of physical and molecular barriers that prevent T cells from effectively populating and protecting the lung.

Flow cytometry or still imaging can only partially capture key parameters of T cell accessibility and dynamics in the lungs. Intravital imaging, the most appropriate approach to overcome the limitations of still imaging (*97*), is particularly challenging to implement in the context of the lungs, especially in a BSL3 environment (*98*, *99*). Leveraging adoptive T cell transfer and intravital lung imaging allowed us to test our hypothesis and directly visualize how an IFN-I-rich environment can physically constrain T cells, with dramatically reduced mobility of the transferred cells that were able to penetrate bacterium-rich lesions in *Sp140*^-/-^ hosts. In contrast, in anti-IFNAR-treated *Sp140*^-/-^ mice, T cells migrated freely into the lung parenchyma and accumulated near *M. tuberculosis* foci, closely resembling the behavior of CD4^+^ T cells observed in infected lungs of WT C57BL/6 mice (*75*). These results are consistent with the recent study of Branchett *et al.*, which demonstrates that IFN-I signaling promotes exclusion of CD4^+^ T cells from lesions (*6*). They are also compatible with live imaging of granuloma explants showing that intralesional CD4^+^ T cell motility was reduced after exacerbation of the pathology through SIV co-infection (*100*). Our results extend this concept by revealing that excessive IFN-I signaling not only limits T cell entry into infected areas but also constrains their motility within lesions, which could contribute to the inhibition of effective local anti-TB immunity.

These observations align with recent reports that elevated IFN-I signaling can hinder T cell migration in a pleiotropic manner by modulating multiple parameters, such as chemokine production and the tissue microenvironment. For instance, virus-induced IFN-I has been shown to suppress CXCL9 and CXCL10 production by myeloid cells, impairing the pulmonary influx of *M. tuberculosis*-specific Th1 cells (*56*). Additionally, IFN-I-driven inflammation may create structural barriers to T cell entry, such as excessive extracellular matrices or NETs (*8*, *59*). High levels of citrullinated histones (cit-H3) in the pulmonary lesions in *Sp140*^-/-^ mice indicate abundant NET formation, which can trap bacteria but also exacerbate tissue damage (*5*, *8*, *59*) and have been shown to correlate with exclusion of CD4^+^ T cells from lesions (*6*, *7*). Our data confirm that T cells are excluded from infected sites when IFN-I is rampant, a condition reversed by neutralizing this cytokine signaling. In non-human primate, IFN-I-expressing cells surround but do not infiltrate cit-H3^+^ areas (*59*). Since T cells are excluded from cit-H3 areas, whether T cells are in close proximity of IFN-I-producing cells remain to be investigated. Also, imaging T cell positioning respective to local IFN-I concentration in TB lesions was not determined in our study and may help addressing this question. Thus, one key mechanism by which Sp140 enables effective TB immunity is by keeping IFN-I responses in check, ensuring that T cells can navigate to and engage pathogens in the lung tissue. This supports the idea that physical or functional barriers, whose identity yet remains to be fully characterized, undermine the effectiveness of T cell-mediated immunity in TB (*97*).

A limitation of our study relies on the fact that magnitude of the T cell phenotype in *Sp140^-/-^* mice is influenced by infectious dose. Lowering the inoculum reduces the magnitude of the phenotype but does not eliminate the underlying Sp140-dependent inflammatory program. Most importantly, our central mechanistic conclusion is that IFNAR blockade in *Sp140^-/-^*, but not *Sp140^+/+^,* mice restored CD8^+^ T cell numbers and reduced bacterial burden and lung pathology. Thus, although elevated bacterial burden and tissue damage may amplify T cell loss, the impaired T cell response is not simply a nonspecific consequence of higher inoculum. Rather, it is driven by the exaggerated IFN-I-dependent pathological milieu that develops in the absence of Sp140. Testing lower inocula would be informative, but would primarily address the threshold and kinetics of pathology development rather than the mechanism identified here. Yet, interpreting the restoration of endogenous T cell responses in susceptible *Sp140^-/-^* or C3HeB/FeJ mice, treated with anti-IFNAR, is challenging because of the potential effects of altered IFN-I milieus on initial T cell priming and differentiation, and subsequent effects on activation, migration, and survival. While providing the concept that IFN-I-dependent damages physically alter T cell dynamics, our results do not examine if IFN-I could affect intrinsically multiple CD8^+^ and CD4^+^ T cell subsets elicited during infection. T cell-specific perturbation of IFNAR signaling in individual lineages or subsets will be required to fully address this question but remains beyond the scope of the current study.

In summary, our study uncovers a critical intersection between innate immune regulation and adaptive immunity in TB. SP140 emerges as a key host factor that restrains IFN-I-driven pathology, thereby preserving the CD8^+^ and CD4^+^ T cell response for optimal control of *M. tuberculosis*. These findings place Sp140 within a growing framework whereby excessive IFN-I signaling is a dominant antagonist of protective TB immunity (*52*, *90–92*). Importantly, the parallels between Sp140-deficiency in mice and the elevated neutrophil driven IFN-I gene signature observed in TB patients, which correlates with severe disease and destructive lung pathology (*52*, *53*), suggest that our results have direct clinical relevance. This notion is further substantiated by genetic evidence showing that single-nucleotide polymorphisms in SP140 confer an increased risk of developing multiple chronic inflammatory diseases in humans [see (*61*) and references therein]. Excessive IFN-I signaling has been proposed to render TB an “atypical interferonopathy” (*101*). In this context, our findings highlight the therapeutic potential of targeting the IFN-I axis to restore host resistance. Transient IFNAR blockade or inhibition of downstream pathways could re-establish effective T cell functions and bacterial control in individuals with hyperinflammatory TB. More broadly, this work opens new directions for modulating the tissue microenvironment of TB lesions to enhance T cell activity. Future studies aim at defining the molecular “brakes” imposed by IFN-I, whether through altered chemokine gradients, metabolic suppression, or physical barriers, and determining how relieving these constraints can improve vaccine efficacy and treatment outcomes.

## Materials and Methods

### Mice

Male and female mice aged 6-10 weeks were used throughout the study. C57BL/6 (*Sp140^+/+^*) mice were purchased from Charles River France. *Sp140^-/-^* mice were kindly provided by Dr. Russel Vance (UC Berkeley, USA) and bred at the Institute of Pharmacology and Structural Biology (IPBS) UMR 5089 (agreement F31555005). *Rag2^-/-^* and Ly5.1 mice were purchase from Jackson laboratory and bred at the IPBS. All procedures complied with French regulations and were approved by the Ministry of Higher Education and Research (APAFIS agreements 38001, 34716, 47848 and 50470).

### Bacteria

Fluorescent derivatives of the wild-type *M. tuberculosis* strain Erdman were constructed by transformation of this strain with pGMCS-P1-GFP and pGMCS-P1-mTurquoise. Plasmids pGMCS-P1-GFP and pGMCS-P1-mTurquoise are integrative vectors in mycobacteria that confer resistance to streptomycin and constitutively express GFP or mTurquoise fluorescent reporters, respectively. They were constructed by Gateway cloning, as described previously (*102*).

*M. tuberculosis* Erdman and its fluorescent GFP or mTurquoise transformants were cultured either in suspension in Middlebrook 7H9 medium (BD) supplemented with 10% albumin-dextrose-catalase (ADC, BD) and 0.05% Tyloxapol (Sigma), or on 7H11 medium supplemented with 10% oleic acid-albumin-dextrose-catalase (OADC, BD). For infection, exponentially growing cultures were centrifuged at 2,301 × *g*, resuspended in phosphate-buffered saline (PBS; Gibco), and de-aggregated by vortexing with glass beads. Remaining clumps were removed by a low-speed spin (120 × *g*). Bacterial concentration was estimated by measuring the optical density at 600 nm (OD_600_), and the suspensions were adjusted in PBS for *in vivo* infection.

### Mouse infection

Mice were exposed to 0.4 × 10^4^ or 2 × 10^6^ bacteria in an inhalation tower (Buxco Inhalation Exposure System, DSI), which is calibrated to deliver approximately 30 or 100 CFUs, respectively, to the lungs per mouse. At 24 h post-infection, lung homogenates from ≥2 mice were plated to confirm the delivered dose. Bacterial burden was quantified by plating serial dilutions on 7H11 + 10% OADC and incubating for 3 weeks at 37 °C.

### Adoptive co-transfer of *Sp140^+/+^* and *Sp140^-/-^* T cells

Donor T cells were isolated from the spleens of naïve CD45.1^+^ *Sp140^+/+^* and CD45.2^+^ *Sp140^-/-^* mice by magnetic enrichment using anti-CD3 microbeads (Mojosort kit, BioLegend). Purified *Sp140^+/+^* and *Sp140^-/-^* T cells were mixed at a 1:1 ratio, and a total of 2 × 10^6^ T cells per mouse was administered intravenously to *M. tuberculosis*-infected *Rag2^-/-^* recipient mice at day 7 post-infection.

### *In vivo* treatment

Anti-IFNAR monoclonal antibody (500 µg; Bio X Cell, Cat# BE0241, RRID:AB_2687723) or isotype control was administered intraperitoneally every 2 days from day 7 to day 26 post-infection. Lungs from untreated, isotype-treated, or anti-IFNAR-treated mice were collected at day 28.

### Lung harvest and cell isolation

Five minutes before euthanasia, mice received an intravenous injection of anti-CD45 monoclonal antibody (5 µg) to distinguish parenchymal (CD45iv**^-^**) from intravascular (CD45iv^+^) leukocytes. Mice were anesthetized with isoflurane and euthanized by cervical dislocation. Lungs were aseptically harvested and homogenized in PBS (Difco) using a gentleMACS dissociator (C Tubes, Miltenyi). Homogenates were incubated with DNase I (0.1 mg/mL; Roche) and collagenase D (2 mg/mL; Roche) for 30 min at 37 °C, 5% CO_2_, filtered through 70 µm strainers, and centrifuged at 423 × *g* for 5 min. Red blood cells were lysed (150 mM NH_4_Cl, 10 mM KHCO_3_, 0.1 mM EDTA, pH 7.2; 5 min), washed, resuspended in Cell Staining Buffer (BioLegend), and filtered through 40 µm strainers.

### Spectral cytometry staining

Single-cell suspensions were stained in 96-well plates with an MHC-I tetramer loaded with the immunodominant TB10.4 antigen for 1 h at room temperature (RT), washed, and incubated with extracellular antibodies and viability dye for 20 min at RT. Cells were then fixed and permeabilized using the eBioscience^TM^ Foxp3/Transcription Factor Staining Buffer Set (30 min, per manufacturer’s instructions), followed by intracellular staining for 30 min at RT. For *M. tuberculosis* inactivation, cells were fixed in 4% paraformaldehyde (Fisher Scientific) for 2 h at RT. Samples were acquired on a Cytek Northern Lights 3L (16V-14B-8R) spectral cytometer (Cytek Biosciences). Spectral unmixing was performed with SpectroFlo (Cytek Biosciences). Conventional analyses were performed in FlowJo v10 (BD), and unsupervised analyses/visualizations were performed in OMIQ (Dotmatics). After data cleaning (PeacoQC), datasets were scaled and subsampled prior to UMAP generation. Clustering was subsequently carried out using FlowSOM.

### Intracellular cytokine staining

For cytokine detection, lung single-cell suspensions were incubated at 37 °C, 5% CO_2_ for 4 h with a pool of *M. tuberculosis*-derived peptides (Table S1) in the presence of anti-CD28 (2µg/mL; BioLegend, Cat# 102102, RRID:AB_312867) and monensin (2µM; eBioscience). Cells were then stained, fixed, and permeabilized as described above before intracellular staining.

### Cytometry antibodies and reagents

Cytometry staining was performed using the following antibodies and reagents: anti-CD103 BV605, clone 2E7 (BioLegend, Cat# 121433, RRID:AB_2629724); anti-CD115 PE, clone AFS98 (BioLegend, Cat# 135505, RRID:AB_1937254); anti-CD11b BV650, clone M1/70 (BioLegend, Cat# 101239, RRID:AB_11125575); anti-CD11c FITC, clone N418 (BioLegend, Cat# 117306, RRID:AB_313775); anti-CD122 PE-Dazzle, clone TM-b1 (BioLegend, Cat# 123217, RRID:AB_2572179); anti-CD127 APC-Cy7, clone A7R34 (BioLegend, Cat# 135040, RRID:AB_2566161); anti-CD170 BV785, clone E50-2440 (BD Biosciences, Cat# 740956, RRID:AB_2740581); anti-CD19 BV570, clone RA3-6B2 (BioLegend, Cat# 103237, RRID:AB_10900264), or Pacific Blue, clone 6D5 (BioLegend, Cat# 115523, RRID:AB_439718); anti-CD3 APC or BV570, clone 17A2 (BioLegend, Cat# 100236, RRID:AB_2561456; Cat# 100225, RRID:AB_10900444); anti-CD4 PE, clone GK1.5 (BioLegend, Cat# 100407, RRID:AB_312692), or PerCP-eFluor 710, clone RM4-5 (Thermo Fisher Scientific, Cat# 46-0042-82, RRID:AB_1834431); anti-CD44 PerCP, clone IM7 (BioLegend, Cat# 103035, RRID:AB_10639933); anti-CD45.1 FITC, clone A20 (BD Biosciences, Cat# 553775, RRID:AB_395043), or BV421, clone A20 (BioLegend, Cat# 110731, RRID:AB_10896425); anti-CD45.2 BV421, clone 104 (BioLegend, Cat# 109831, RRID:AB_10900256), BV711, clone 104 (BD Biosciences, Cat# 563685, RRID:AB_2738374), or Alexa Fluor 700, clone 104 (BD Biosciences, Cat# 560693, RRID:AB_1727491); anti-CD49a PE, clone HMa1 (BioLegend, Cat# 142604, RRID:AB_10945158), or BV711, clone Ha31/8 (BD Biosciences, Cat# 564863, RRID:AB_2738987); anti-CD49d BV480 or BV510, clone 9C10/MFR4.B (BD Biosciences, Cat# 746320, RRID:AB_2743644; Cat# 745007, RRID:AB_2742641); anti-CD62L PE-Fire 810, clone W18021D (BioLegend, Cat# 161205, RRID:AB_2910338); anti-CD64 PE-Dazzle 594, clone X54-5/7.1 (BioLegend, Cat# 139320, RRID:AB_2566559); anti-CD69 BV510, clone H1.2F3 (BioLegend, Cat# 104531, RRID:AB_2562326), or BV480, clone H1.2F3 (BD Biosciences, Cat# 746813, RRID:AB_2744067); anti-CD8α BV650 or Spark Blue 550, clone 53-6.7 (BioLegend, Cat# 100742, RRID:AB_2563056; Cat# 100779, RRID:AB_2832268); anti-CD8β Alexa Fluor 700, clone YTS156.7.7 (BioLegend, Cat# 126618, RRID:AB_2563949); anti-CX3CR1 Alexa Fluor 488, clone SA011F11 (BioLegend, Cat# 149022, RRID:AB_2565705); anti-Eomes eFluor 450, clone Dan11mag (Thermo Fisher Scientific, Cat# 48-4875-82, RRID:AB_2574062); anti-I-A/I-E PerCP-Cy5.5, clone M5/114.15.2 (BioLegend, Cat# 107626, RRID:AB_2191071); anti-IFNγ BV650, clone XMG1.2 (BioLegend, Cat# 505831, RRID:AB_11142685); anti-KLRG1 eFluor 506, clone 2F1 (Thermo Fisher Scientific, Cat# 69-5893-82, RRID:AB_2637143); anti-Ly108 BV750, clone 13G3 (BD Biosciences, Cat# 747169, RRID:AB_2871904); anti-Ly6A/E Alexa Fluor 700, clone D7 (BioLegend, Cat# 108142, RRID:AB_2565959); anti-Ly6C eFluor 450, clone HK1.4 (Thermo Fisher Scientific, Cat# 48-5932-82, RRID:AB_10805519); anti-Ly6G Alexa Fluor 700, clone 1A8 (BD Biosciences, Cat# 561236, RRID:AB_10611860; anti-PD-1 APC-Fire 810, clone 29F.1A12 (BioLegend, Cat# 135251, RRID:AB_2910292); anti-TCF1/TCF7 PE-Cy7, clone C63D9 (Cell Signaling Technology, Cat# 90511, RRID:AB_3086656); anti-TCRβ BV786, clone H57-597 (BioLegend, Cat# 109249, RRID:AB_2810347); anti-TIM-3 PE-Fire 640, clone RMT3-23 (BioLegend, Cat# 119749, RRID:AB_2927886); anti-TNFα Alexa Fluor 660, clone MP6-XT22 (Thermo Fisher Scientific, Cat# 606-7321-80, RRID:AB_2896303); and anti-TOX eFluor 660, clone TXRX10 (Thermo Fisher Scientific, Cat# 50-6502-82, RRID:AB_2574265). Fc receptor binding was blocked using anti-mouse CD16/32, clone 93 (BioLegend, Cat# 101320, RRID:AB_1574975). MHC-I/TB10.4 APC tetramers were obtained from the NIH Tetramer Core Facility as custom reagents.

### Preparation of OCT-embedded tissues

At designated time points, mice were euthanized by intraperitoneal injection of a lethal dose of sodium pentobarbital. The lungs were inflated with 1 mL of 2% ultrapure agarose and then harvested. Tissues were fixed for 24 h at 4 °C in periodate-lysine-paraformaldehyde (PLP) buffer [0.05 M phosphate buffer, 0.1 M L-lysine (Sigma-Aldrich), 2 mg/mL NaIO_4_ (ThermoFisher Scientific), 4% paraformaldehyde; pH 7.4]. Samples were cryoprotected in a sucrose gradient at 4 °C and embedded in optimal cutting temperature (OCT) compound. Frozen tissues were sectioned at 20 µm on a Leica CM1950 cryostat and mounted on SuperFrost Ultra Plus™ GOLD slides (Epredia).

### Tissue preparation for immunofluorescence

OCT sections were thawed in Milli-Q water (2 × 10 min, RT). Non-specific binding was blocked for 1 h at RT in PBS containing 5% goat serum (GeneTex), 5% bovine serum albumin (BSA; Euromedex), and 0.2% Triton X-100 (Sigma-Aldrich). Sections were incubated overnight at 4°C with primary antibodies diluted in PBS + 5% BSA + 0.2% Triton X-100. After washing, secondary antibodies were applied for 4 h at RT, followed by DAPI (5 min; Sigma-Aldrich). Sections were mounted in DAKO Fluorescent Mounting Medium (Agilent) and imaged on a Zeiss LSM 710 confocal microscope. Image processing was performed in ImageJ or QuPath.

### Image quantification

For immunofluorescence staining, the following antibodies were used: APC-conjugated anti-CD3, clone 17A2 (BioLegend, Cat# 100236, RRID:AB_2561456); Alexa Fluor 647-conjugated or purified anti-CD8α, clone 53-6.7 (BioLegend, Cat# 100724, RRID:AB_389326; Cat# 100702, RRID:AB_312741); anti-citrullinated histone H3, clone EPR20358-120 (Abcam, Cat# ab219407, RRID:AB_3665807); Alexa Fluor 555-conjugated goat anti-rabbit IgG (H+L) secondary antibody (Thermo Fisher Scientific, Cat# A-21428, RRID:AB_2535849); and Alexa Fluor 647-conjugated goat anti-rat IgG (H+L) secondary antibody (Thermo Fisher Scientific, Cat# A-21247, RRID:AB_141778). Quantification of citrullinated histone H3 and CD8α-positive signals in lung sections from *M. tuberculosis*-infected mice was performed using QuPath. Whole-section images were analyzed using a standardized pixel-classification thresholding approach to distinguish positive from negative pixels. The same thresholding parameters were applied uniformly to all samples.

### Single-cell RNA-seq sample preparation

Parenchymal (CD45iv^-^) CD8α^+^ T cells were sorted individually from each of 8-12 infected mice per genotype using a FACSAria Fusion (BD) in a BSL-3 facility. Initial sorting from each mouse was performed to ensure sufficient parenchymal CD8^+^ T cell numbers were present. Subsequently, cells were pooled by genotype to reach approximately 1 × 10^6^ cells. Cells from *Sp140^+/+^* mice were collected at day 35 post-infection and from *Sp140^-/-^* mice at day 28 post-infection (infection dose: 30 CFUs). Samples were then processed with the Chromium Fixed RNA Profiling workflow (10x Genomics) using the Single Cell Fixed RNA Sample Preparation Kit, following the manufacturer’s protocol. Briefly, cells were incubated in fixation buffer (1× Fix & Perm Buffer; 4% formaldehyde in nuclease-free water) for 24 h at 4 °C, centrifuged (850 × g, 5 min, 4 °C), resuspended in 1× quenching buffer, and supplemented with 0.1 volume of preheated Enhancer (10x Genomics PN-20000482). Nuclease-free glycerol was added to 10% (final), and samples were stored at -80 °C prior to sequencing (Single Cell Discoveries, Utrecht, Netherlands).

### scRNA-seq data processing and analysis

All downstream analyses were performed using Seurat (v.4.1.1) for R (v4.2.0). Genes detected in fewer than 3 cells, or cells with fewer than 200 or more than 7500 genes detected were excluded from analysis of the *Sp140^+/+^* and *Sp140^-/-^* CD8^+^ T cell datasets. Cells with >10% of transcripts originated from the mitochondria were deemed low-quality and excluded. The TCR genes (Tra[vjc], Trb[vdjc]) were removed to avoid any TCR gene-driven bias during clustering. Log-normalization by the NormalizeData function was performed and the top 2000 highly variable genes were determined using the FindVariableFeatures function. Samples from the *Sp140^+/+^* and *Sp140^-/-^* CD8^+^ T cell datasets were then merged using the SelectIntegrationFeatures, FindIntegrationAnchors, and IntegrateData function. After data scaling by ScaleData function, Principal Component Analysis (PCA) was performed using the RunPCA function. The optimal number of principal components (PCs) for downstream analysis was determined using an elbow plot and the first 15 PCs were selected. RunUMAP was performed and cells were clustered using FindNeighbours and FindClusters functions (res = 0.4). The cluster markers were found using the *FindAllMarkers* function (min.pct = 0.25, assay = “RNA”), and avg_log2 fold change (FC) and adjusted *p* value for top30 markers for each cluster are shown in Table S2. Barplot was visualized by dittoseq (v1.10.0).

### Gene signature scoring

The AddModuleScore function in Seurat (v.4.1.1) was used to compute average gene expression scores for published gene sets (Table S4).

### Differentially expressed gene analysis

Differentially expressed genes (DEGs) between *Sp140^+/+^* versus *Sp140^-/-^* CD8^+^ T cells were identified using the FindMarkers function (Table S5). Volcano plots were created with the R package EnhancedVolcano (v1.16.0). We used the *enrichGO* function in the clusterProfiler package (v4.6.2) to measure the enrichment of modules in GO terms across all three ontologies (“BP,” “MF,” and “CC”).

### Projection of our scRNA-seq data to published scRNA-seq datasets

The Run.ProjecTILs function in the ProjecTILs (v.3.3.0) was used to project our scRNA-Seq dataset onto processed scRNA-Seq data from the following sources: ref_TILAtlas_mouse_v1 (https://figshare.com/articles/dataset/ProjecTILs_murine_reference_atlas_of_tumor-infiltrating_T_cells_version_1/12478571?file=41398167) and GSE199565.

### Pseudotime trajectory analysis

Pseudotime was calculated using the Monocle3 (v.1.3.1) package. Using the SeuratWrappers (v.0.3.0) package, the integrated Seurat object was converted into a cell dataset object. Only activated classical CD8^+^ T cells (Seurat cluster 1-3,5) were retained for the pseudotime trajectory analysis. A cell with the highest expression level of the *Lef1* and *Sell* genes was set as the root node. Cells were clustered with *cluster_cells* using “Leiden” as a cluster_method. Genes of interest were ordered along pseudotime using the plot_genes_in_pseudotime function.

### Lung multiphoton intravital imaging

Lung multiphoton intravital microscopy (MP-IVM) under BSL3 conditions was used to monitor T cell dynamics, employing a dedicated surgical platform that enables mouse surgery and transfer within the BSL-3 environment, as well as a lung window specifically adapted for imaging lesions, as described (*75*). Briefly, *Sp140^-/-^* mice infected 21-28 days earlier with mTurquoise-expressing *M. tuberculosis* and treated or not with anti-IFNAR received an adoptive transfer of fluorescently labeled CD8^+^ and CD4^+^ T cells. Donor T cells were magnetically isolated from infected *Sp140*^+/+^ mice using anti-CD8 or anti-CD4 microbeads (MojoSort kit, Biolegend) and labeled with either CellTrace^TM^ Deep Red or CellTrace^TM^ Orange. Equal number of CD8^+^ and CD4^+^ T cells were mixed 1:1 (2 × 10^6^ each) prior to transfer. Due to laser excitation wavelength and filters emission configuration, CellTrace^TM^ Deep Red-labeled cells were robustly detected and tracked, whereas CellTrace^TM^ Orange-labeled cells were only faintly detectable and used to confirm engraftment. After 24 hours, mice were anesthetized with ketamine/xylazine (105 and 3 mg/kg, respectively, intraperitoneal), followed by subcutaneous injections of buprenorphine (0,1 mg/kg) and Ringer Lactate (10 mL/kg). Dextran, either Texas Red 70,000 MW (1250 μg, D1864 Invitrogen) or Cascade Blue 10,000 MW (1250 μg, D1864 Invitrogen), (50 µL of 25 mg/mL; Life Technologies) was administered intravenously before imaging to visualize vasculature. Together with the presence of fluorescent *M. tuberculosis*, dextran staining also allowed to identify lesions based on vasculature remodeling (Supplemental Movie 6-7). Mice were intubated for mechanical ventilation, and isoflurane (1%) was delivered during imaging. Two ribs were transected to allow the installation of a lung window over the lesion. Once transferred in the heated microscope isolator, suction (20-25 mmHg) was applied to gently immobilize the lung against the coverslip (Dexter Medical 0-250 mbar) of the window. MP-IVM was performed using a Trimscope II Miltenyi Biotec (LaVision Biotec) upright multi-photon microscope equipped with a 20×/1.0 water immersion objective and a Ti-Sapphire femtosecond laser, Chameleon-Ultra II (Coherent Inc.) tuned to 820 nm. Emission signals corresponding to mTurquoise/Cascade Blue, AlexaFluor488, CTO/Texas Red and Deep Red signals were collected in separate channels using 450/50, 525/50, 595/40 and LP650 filters, respectively. Four channel images were acquired at a rate of ∼1 frame per second for 405 × 405 µm field of view at a resolution of 512 × 512 pixel. Multi-tile three-dimensional time-lapse sequences of lesions were recorded for 30 minutes to 2 hours, with Z-stacks collected approximately every minute.

### Time-lapse video processing and T cell tracking analysis

Three-dimensional multi-tile videos were stitched and converted into 2D maximum-intensity projections, after removal of z-slices lacking relevant information. Fluorescence from the CTO/Texas Red channel was then subtracted from the Deep Red channel using channel arithmetic. Each channel was denoised independently prior to running the Bleach Correction and BaSiC (a tool for background and shading correction) plugins in ImageJ (National Institutes of Health). Channels were then merged and despeckled before export as MP4 videos at 10 frames pers second, including scale bars and time stamps. Representative images shown in the figures correspond to the first frames of the processed videos. For T cell tracking, upper and lower z-planes lacking relevant information were removed from stitched 3D multi-tile videos, prior to conversion into 2D maximum-intensity projections. Drift occurring over the course of the acquisition was corrected using the mTurquoise/Cascade Blue channel as a reference with the Correct 3D drift plugin in ImageJ. Quantification of T cell dynamics was performed as previously described using Trackmate plugin in ImageJ (*75*). To identify CTO signal within the Deep Red channel, green and red LUTs were applied to the respective channels, and yellow cells were excluded from tracking in the Deep Red channel. For each tracked cell, track length, displacement, straightness, instantaneous speed, average speed and maximum speed were quantified. The arrest coefficient, defined as the percentage of time during which a cell’s instantaneous speed remained below 2 µm/min, was also calculated. Lesions were delineated based on the remodeled and tortuous dextran-labeled vasculature in infected regions and the presence of mTurquoise-expressing *M. tuberculosis*.

### Statistical analysis

Sample sizes (*n*), statistical tests, and significance thresholds are indicated in the figure legends. Analyses were performed in Prism 10.0 (GraphPad). Depending on the experiment, either an unpaired ANOVA or an unpaired Student’s t-test was used, as specified in the figure legends. Significance thresholds were *p* < 0.05 (*)*, p* < 0.005 (**), *p* < 0.0005 (***), and *p* < 0.0001 (****).

## Acknowledgments

We thank Russel Vance for the kind gift of the *Sp140^-/-^* mice as well as Geanncarlo Lugo-Villarino and Yoann Rombouts for critical reading of our manuscript. We thank Jérémy Sintes for calibrating the aerosol delivery apparatus and Claude Gutierrez for the conception of plasmids to transform *M. tuberculosis.* We acknowledge the cytometry (Genotoul TRI-IPBS) and animal (Genotoul Anexplo-IPBS) facilities, particularly Flavie Moreau, Malory Blasco and Céline Berrone for technical support. TRI-IPBS is member of the national infrastructure France-BioImaging (https://ror.org/01y7vt929) supported by the French National Research Agency (ANR-24-INBS-0005 FBI BIOGEN); Anexplo-IPBS is member of the national infrastructure Celphedia (https://ror.org/00v2cdz24). This work was supported by Fondation Bettencourt Schueller (Grant Explore-TB to O.N.), ANRS MIE (Contract KILL-TB #206385 to D.H.), the European Commission Horizon Programme (Grant TBVAC-HORIZON #101080309 to O.N. and E.L. and Grant ITHEMYC to O.N.), the Fondation pour la Recherche Médicale (Grant Equipe FRM #EQU202103012733 to O.N.), National Institutes of Health (grants 1R01AI192333 and R01AI172905 to S.M.B., and grant R21 AI156407 to E.L and O.N.), ANRS MIE (PhD fellowship to L.F. and to M.C.), University of Toulouse (Tremplin 2022 #0003970 to E.L.). Figures were created with Biorender.com.

## Authors’ contributions

M.C. conducted experiments, analyzed data and wrote the manuscript. L.F. and A.-I.A.-H. performed intravital microscopy experiments. T.S. analyzed the scRNA-seq data of lung CD8^+^ T cells. M.F. generated fluorescent strains of *M. tuberculosis* and contributed to *in vivo* experiments. A.B. contributed to *in vivo* experiments. S.M. designed and built the surgical platform and lung window, coded all in-house Python scripts for cell tracking and behavior analysis, and performed image and movie analyses. Y.D. performed immunofluorescence staining and their quantification. E.D. contributed to the analysis of intravital microscopy experiments. Y.L. characterized the pool of peptides for T cell stimulation. E.N. helped design the antibody panels for spectral cytometry. S.M.B. provided financial support and directed the work of T.S. E.L. supervised, designed, performed and analyzed the intravital microscopy experiments. O.N. secured major funding and provided regular, in-depth scientific guidance, contributing to experimental design and key expertise. D.H. supervised the project, designed and analyzed experiments, and wrote the manuscript. All authors critically revised the manuscript.

## Competing Interest Statement

The authors declare that they have no competing interests

## Data availability statement

All data needed to evaluate the conclusions in the paper are present in the paper and/or the Supplementary Materials.

## Legend to Supplemental Figures

**Supplemental Figure 1 (related to Figure 1).**
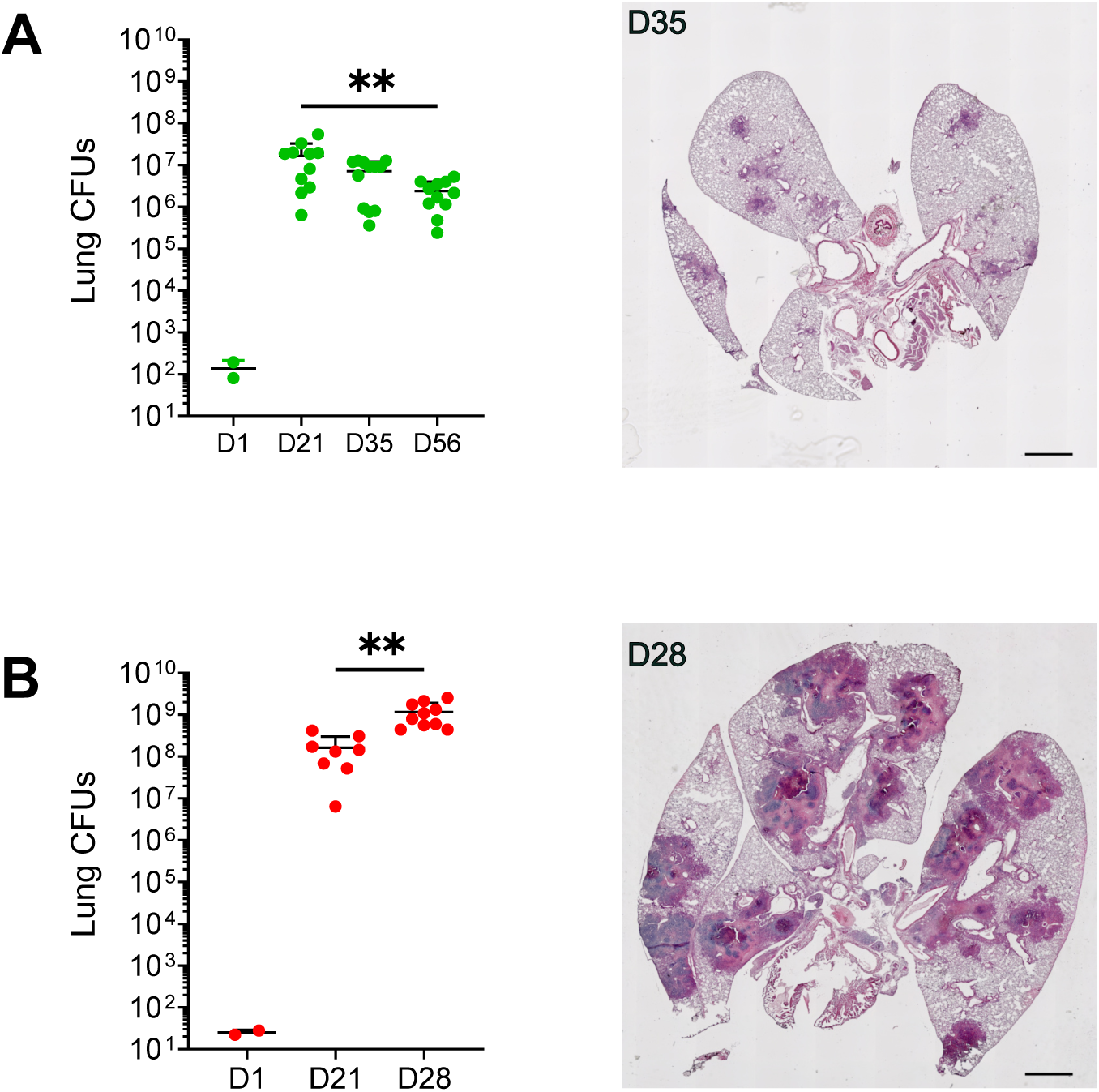
Bacterial loads and pathology in *Sp140^+/+^* and *Sp140^-/-^* mice infected with *M. tuberculosis. Sp140^+/+^* and *Sp140^-/-^* mice were exposed to the aerosol route with 100 CFUs of the Erdman strain of *M. tuberculosis,* and at the indicated time points, they were sacrificed, and their lungs were collected. A. Representative hematoxylin and eosin staining of lung sections (20 µm) from *Sp140^+/+^* mice at day 35 post-infection (bacterial loads shown for days 21 (n=11), 35 (n=12), and 56 (n=11)). Each data point corresponds to one individual mouse. Bars and error bars indicate the mean ± standard deviation. CFUs data are pooled from two independent experiments, and unpaired two-way ANOVA tests with Tukey correction were used. Histological images are representative of at least two independent experiments. Scale bar, 1000 µm. B. Same for *Sp140^-/-^* mice at day 28 post-infection (bacterial loads shown for days 21 (n=8) and 28 (n=10)).

**Supplemental Figure 2 (related to Figure 1).**
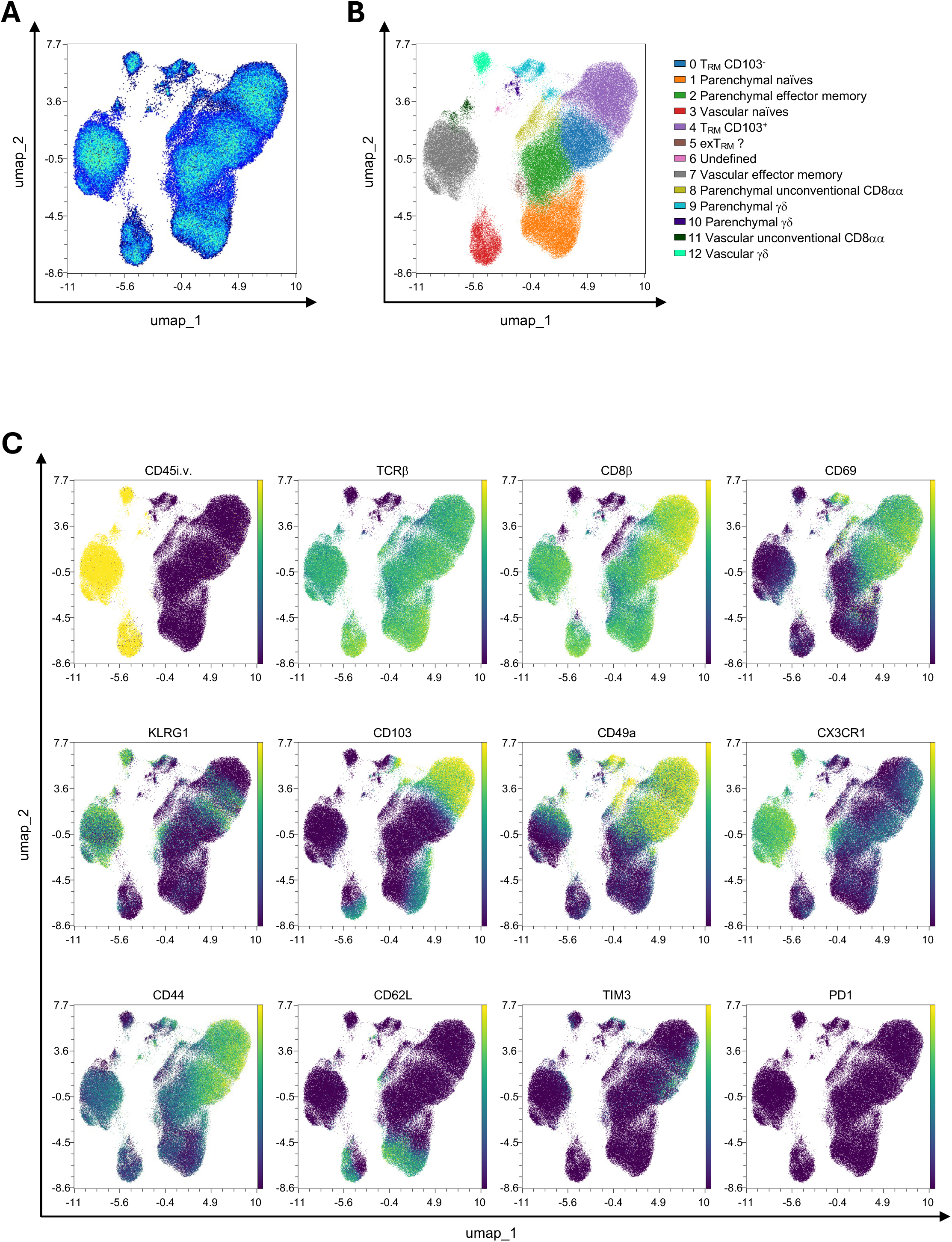
Unbiased analysis of spectral cytometry data describing CD8^+^ T cells from *Sp140^+/+^* mice infected with *M. tuberculosis. Sp140^+/+^* were exposed to the aerosol route with 100 CFUs of the Erdman strain of *M. tuberculosis* and were sacrificed at day 35 post-infection, and their lungs were collected. A. UMAP, performed using OMIQ, of the CD3^+^CD8α^+^ cells and represented using a density plot. B. Clustering performed using FlowSOM algorithm of the populations presented in A). C. UMAP representation with the expression of the indicated markers displayed as a continuous color scale.

**Supplemental Figure 3 (related to Figure 1).**
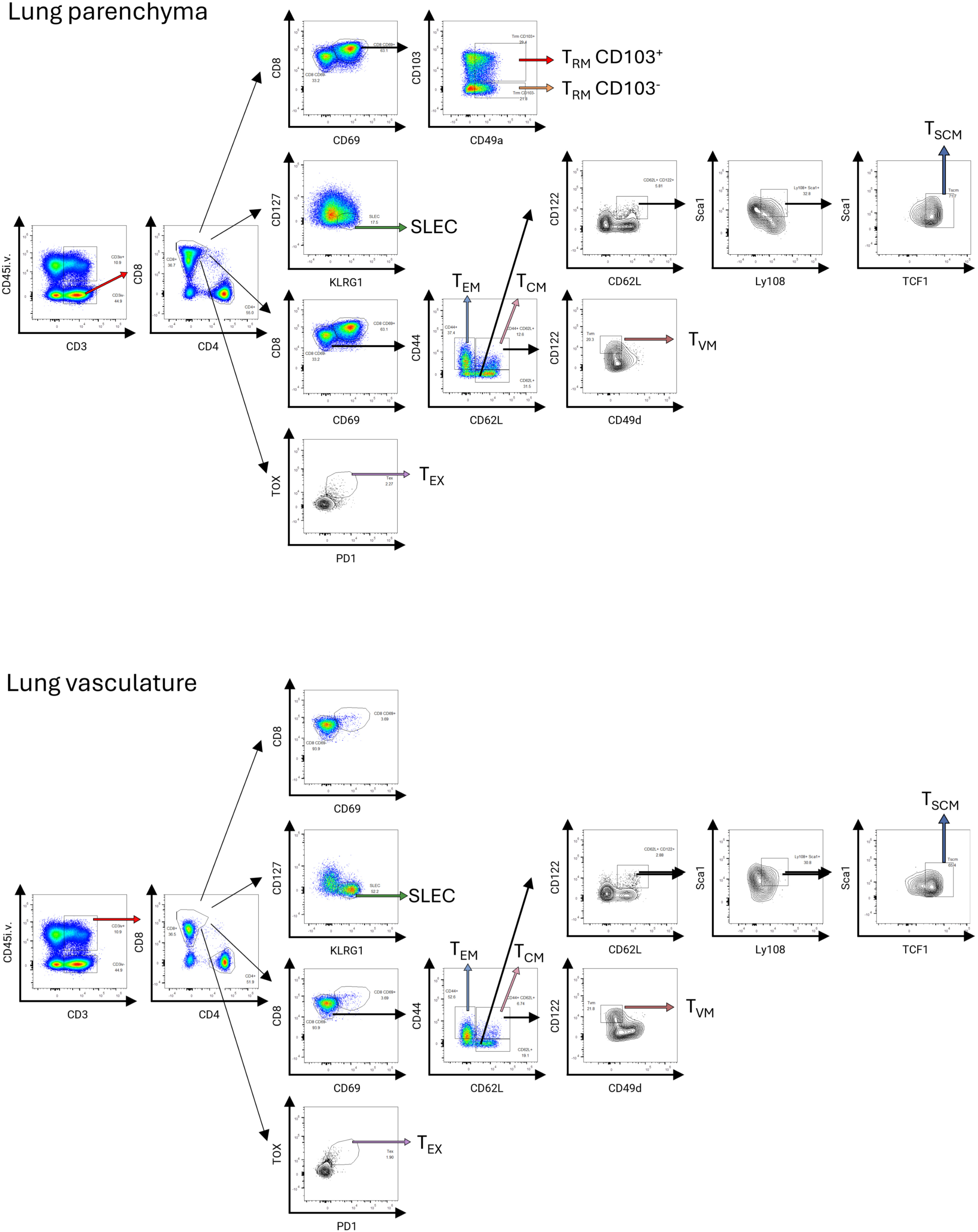
Gating strategy to analyze CD8^+^ T cells in *Sp140^+/+^* and *Sp140^-/-^* mice infected with *M. tuberculosis.* After excluding doublets, the live cells isolated from the lungs of infected mice are first separated based on the expression of CD3 and CD45iv, which allows us to analyze parenchymal T lymphocytes (CD45iv**^-^**, top panel) compared to vascular T lymphocytes (CD45iv^+^, bottom panel). The CD4 vs CD8 staining enables us to focus on CD8^+^ T cells. The various effector cells (short-lived effector cells, SLEC), memory-like cells (central memory-like, T_CM_; effector memory-like, T_EM_; resident memory-like CD103^+^ and CD103^-^, T_RM_; virtual memory-like, T_VM_; stem cell memory-like, T_SCM_), and exhausted cells (T_EX_) are identified using the indicated surface markers.

**Supplemental Figure 4. (related to Figure 1).**
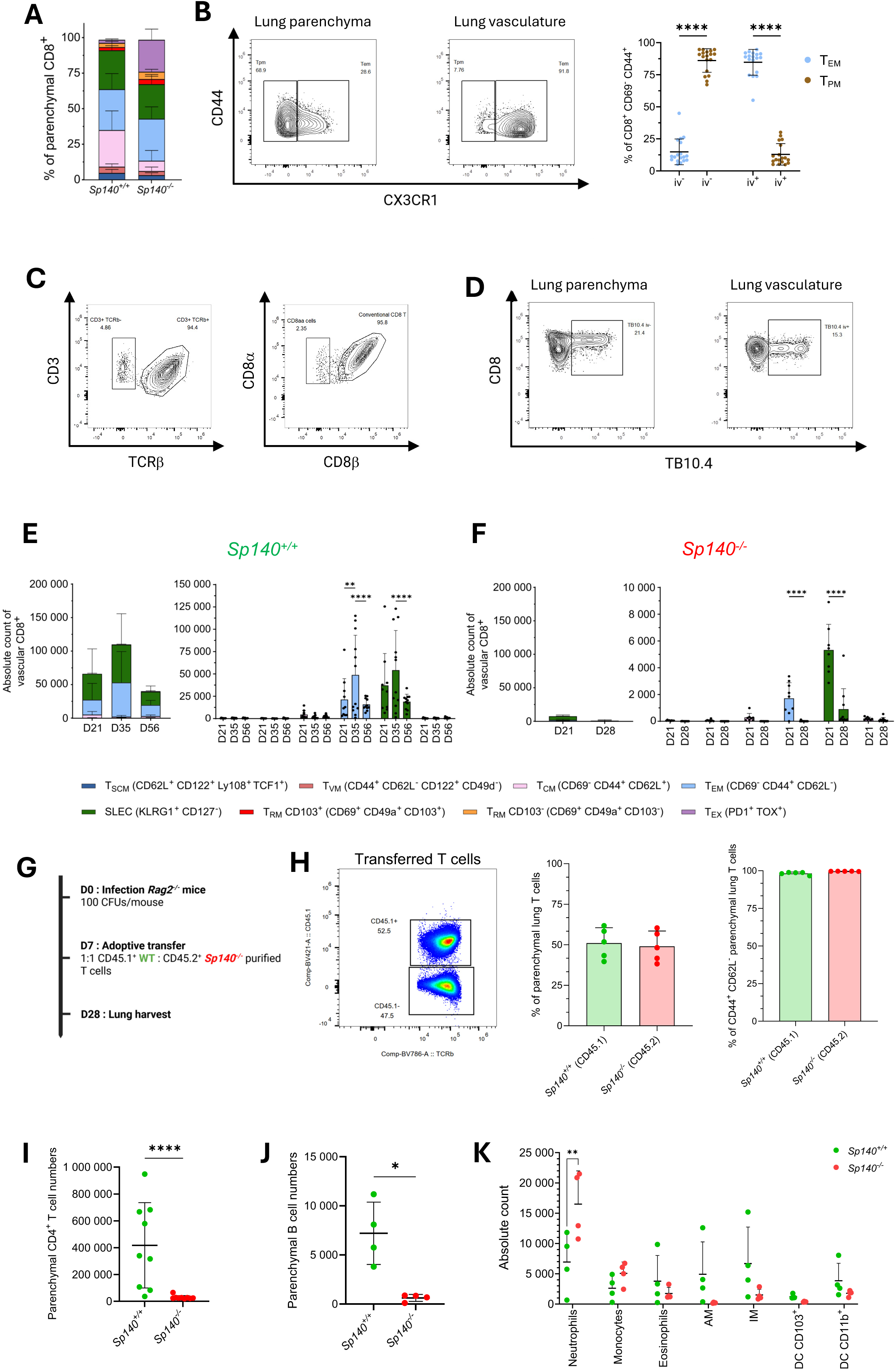
Further leukocyte subset definition in *Sp140^+/+^* and *Sp140^-/-^* mice infected with *M. tuberculosis*. A. Normalized CD8^+^ T cell diversity at day 21 in *Sp140^+/+^* and *Sp140^-/-^* mice. Bars and error bars indicate the mean ± standard deviation. B. Representation of T_EM_ (CX3CR1^+^) and T_PM_ (CX3CR1^int^) proportion in CD69^-^ CD44^+^ CD62L^-^ CD8^+^T cells in parenchyma (left) and vasculature (right). Quantification of each subpopulation is shown on the right. Each data point corresponds to one individual *Sp140^+/+^* mouse (n=18). Bars and error bars indicate the mean ± standard deviation. One-way ANOVA test with Tukey correction was used. C. Representation of CD3^+^TCRβ^+^ and CD3^+^TCRβ^-^ (on gated CD3^+^ cells) and CD8αα/CD8αβ (on gated CD8α^+^ cells). D. Representation of TB10.4-specific CD8^+^ T cells in parenchyma (left) and in vasculature (right). E. Absolute numbers of the main vascular lung CD8^+^ T cell subsets in *Sp140^+/+^* mice at days 21 (n=11, mean number of total CD8^+^ T cell: 67,566), 35 (n=12, mean number of total CD8^+^ T cell: 103,377), and 56 (n=11, mean number of total CD8^+^ T cell: 46,972) post-infection. Stacked (left) and individual subpopulations (right) representations are shown. Each data point corresponds to one individual mouse. Bars and error bars indicate the mean ± standard deviation. Data are pooled from two independent experiments, and unpaired two-way ANOVA tests with Tukey correction were used. F. As in E) but for *Sp140^-/-^* mice at days 21 (n=8, mean number of total CD8^+^ T cell: 11,600) and 28 (n=10, mean number of total CD8^+^ T cell: 2,084) post-infection. The color code for individual CD8^+^ T cell subsets in panels A, E, and F is identical to that used in Figure 1. G. Experimental design of the competitive adoptive T-cell co-transfer experiment. *Rag2^-/-^* mice were infected with *M. tuberculosis* and, 7 days later, received purified T cells from Sp140-competent CD45.1^+^ and Sp140-deficient CD45.2^+^ donor mice mixed at a 1:1 ratio. H. Ratio of parenchymal CD45.1^+^ and CD45.2^+^ T cells in the lungs (left panel) and the activated CD44^+^CD62L^low^ phenotype of the lung parenchymal T cells (right panel). An experiment with 5 mice per group representative of two independent ones is shown. Non parametric Mann-Whitney test was used. I. Absolute numbers of the total, lung parenchymal, CD4^+^ T cell (defined as CD45iv^-^ CD4^+^CD3^+^ live cells) in *Sp140^+/+^* and *Sp140^-/-^* mice at day 28. Each data point corresponds to one individual mouse. Bars and error bars indicate the mean ± standard deviation. Data are pooled from two independent experiments. Non parametric Mann-Whitney test was used J. As in I. but for B cells (defined as CD45iv^-^ CD19^+^ live cells). K. As in I. but for myeloid cells. The populations were defined as neutrophils (CD11b^hi^ Ly6C^+^ Ly6G^+^), monocytes (CD11b^hi^ Ly6C^+^ Ly6G^-^), eosinophils (non-neutrophils, non-monocytes, SiglecF^+^, CD11c^-^), alveolar macrophages (AM; non-eosinophils, CD64^+^, CD115^-^, SiglecF^+^, CD11c^+^), interstitial macrophages (IM, non-eosinophils, CD64^+^, CD115^-^, SiglecF^+^, CD11c^+^), CD103^+^ dendritic cells (DC CD103^+^; non-macrophages, I-A/I-E^+^ CD11b^-^ CD103^+^), CD11b^+^ dendritic cells (DC CD11b^+^; non-macrophages, I-A/I-E^+^ CD11b^+^ CD103^-^).

**Supplemental Figure 5 (related to Figure 1).**
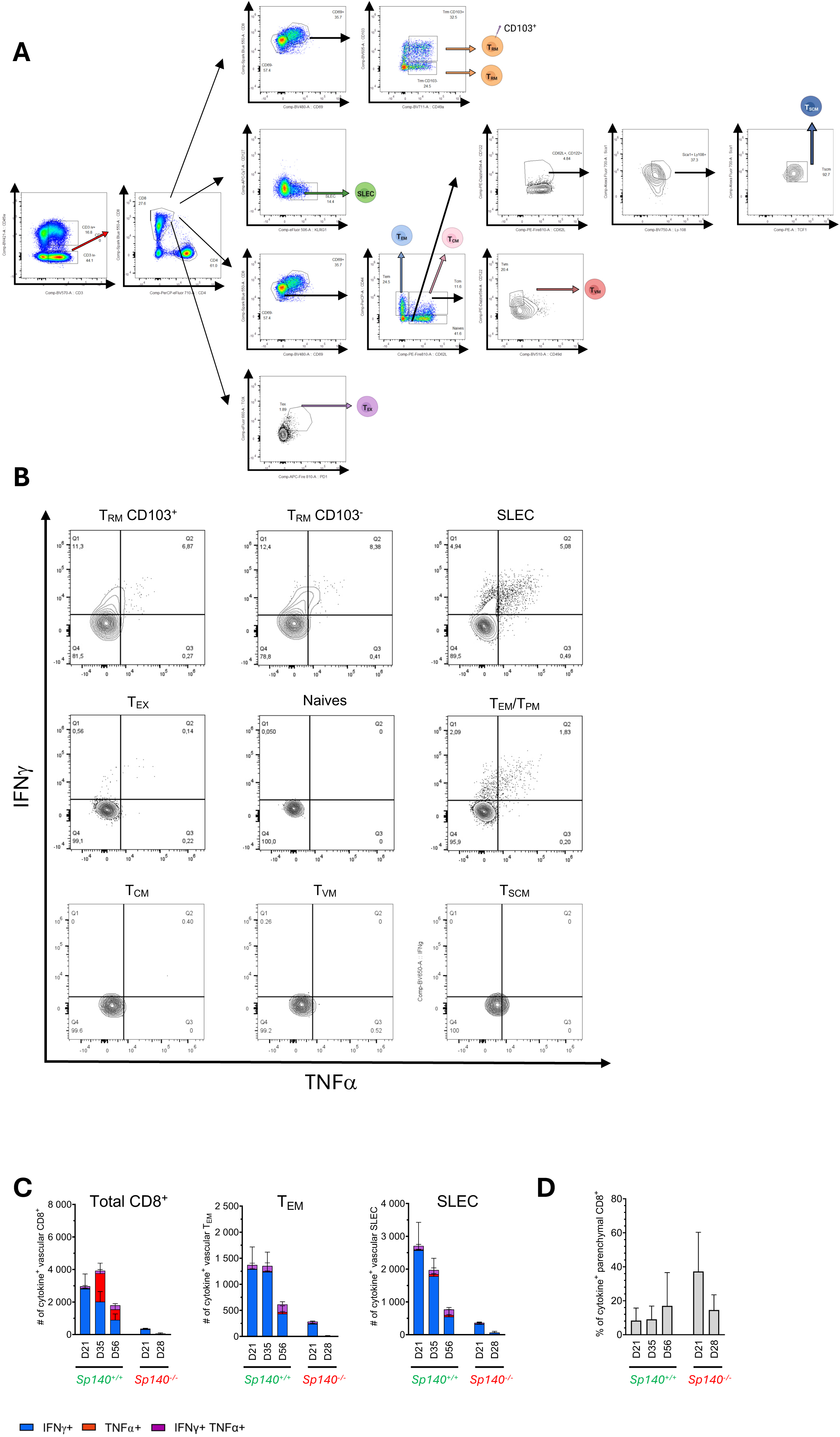
Cytokine production by CD8^+^ T cells in *Sp140^+/+^* and *Sp140^-/-^* mice infected with *M. tuberculosis*. A. Gating strategy to analyze CD8^+^ T cells in *Sp140^+/+^* and *Sp140^-/-^* mice infected with *M. tuberculosis* and after *in vitro* stimulation as described in the Materials and Methods section for cytokine production. After excluding doublets, the live cells isolated from the lungs of infected mice are first separated based on the expression of CD3 and CD45iv, which allows us to analyze parenchymal T lymphocytes (CD45iv**^-^**) compared to vascular T lymphocytes (CD45iv^+^). The CD4 vs CD8 staining enables us to focus on CD8^+^ T cells. The various effector cells (short-lived effector cells, SLEC), memory-like cells (central memory-like, T_CM_; effector memory-like, T_EM_; resident memory-like CD103^+^ and CD103^-^, T_RM_; virtual memory-like, T_VM_; stem cell memory-like, T_SCM_), and exhausted cells (T_EX_) are identified using the indicated surface markers. B. Expression of IFNγ and TNFα after *M. tuberculosis* peptide stimulation on CD8^+^ T cell subsets on day 56 infected *Sp140^+/+^* mice. C. Absolute numbers of IFNγ, TNFα, or both producing lung vascular CD8^+^ T cells and main producing CD8^+^ subpopulations after stimulation with *M. tuberculosis* peptides. Data for *Sp140^+/+^* mice are shown at days 21 (n=11), 35 (n=12), and 56 (n=11) (left) and for *Sp140^-/-^* mice at days 21 (n=8) and 28 (n=10) (right). Bars and error bars indicate the mean ± standard deviation. D. Proportions of lung parenchymal CD8^+^ T cells producing either or both IFNγ and TNFα, after stimulation with *M. tuberculosis* peptides. Data for *Sp140^+/+^* mice are shown at days 21 (n=11), 35 (n=12), and 56 (n=11) (left) and for *Sp140^-/-^* mice at days 21 (n=8) and 28 (n=10) (right). Bars and error bars indicate the mean ± standard deviation.

**Supplemental Figure 6 (related to Figure 2).**
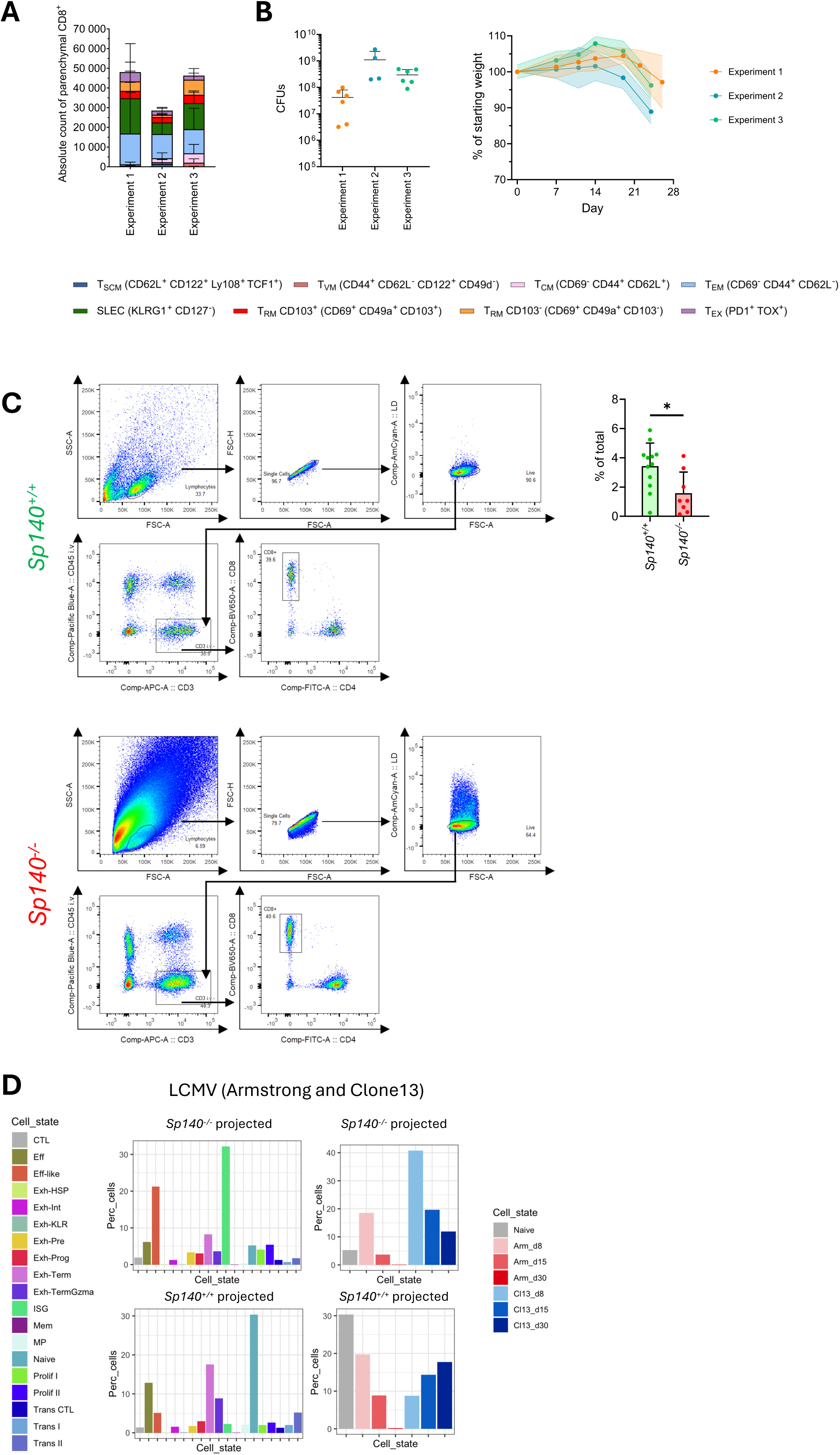
scRNAseq analysis of CD8^+^ T cells from *Sp140^+/+^* and *Sp140^-/-^* mice infected with *M. tuberculosis*. A. CD8^+^ T cell diversity and numbers in *Sp140^-/-^* mice infected with 30 CFUs of *M. tuberculosis*, analyzed as in Figure 1A-D. The color code for individual CD8^+^ T cell subsets is identical to that used in Figure 1. Three independent experiments are shown. B. Bacterial loads and weight loss in *Sp140^-/-^* mice infected with 30 CFUs of *M. tuberculosis* (to be compared with Supplemental Figure 1 and Figure 3A-B). For CFUs data, each data point corresponds to one individual mouse. Bars and error bars indicate the mean ± standard deviation. For weight monitoring, each dot represents the mean weight at the indicated time point, and the shaded area corresponds to the standard deviation. Three independent experiments are shown. C. Gating strategy used to sort CD45iv**^-^** CD8^+^ T cells from infected *Sp140^+/+^* (top 5 panels) and *Sp140^-/-^* (lower 5 panels) mice for scRNA-seq and percentages of CD8^+^ T cells from both genotypes (right). Each dot represents a mouse used as a source of CD8^+^ T cells for sorting, and an unpaired Mann-Whitney t-test was used for statistical analysis. D. CD8^+^ T cells from *Sp140^+/+^* or *Sp140^-/-^* mice projected onto CD8^+^ T cells from an LCMV infection dataset (GSE199565). E. Volcano plot of DEGs between IFN stimulated (Cluster 5) and other activated clusters (Cluster 0,1,2) in *Sp140^-/-^* dataset. Genes with log2FC > 0.5 and an adjusted p value < 0.01 are designated by a blue-filled circle.

**Supplemental Figure 7 (related to Figure 3).**
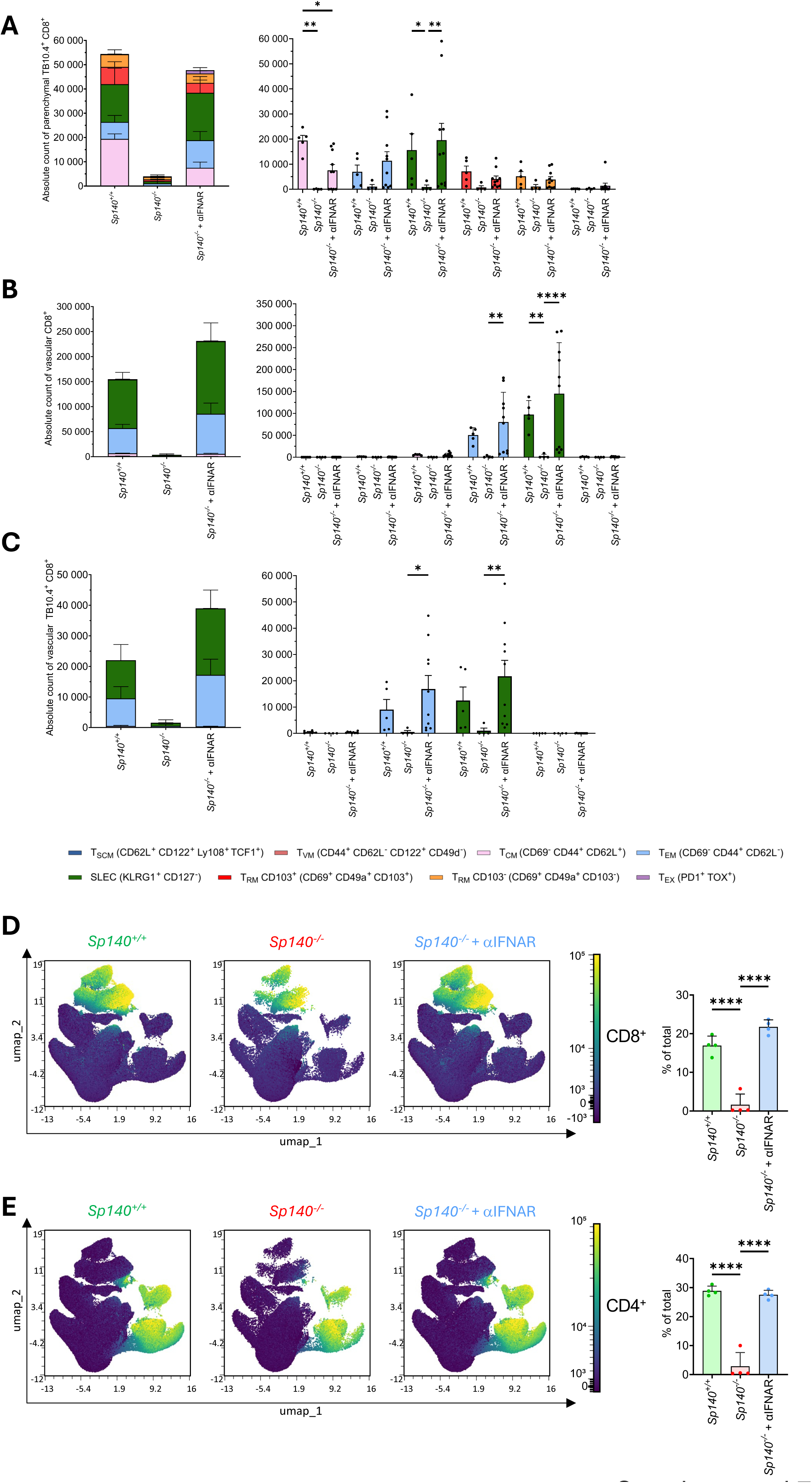
Impact of IFNAR blockade on pathology and T cell responses in *Sp140^+/+^* and *Sp140^-/-^* mice infected with *M. tuberculosis*. A. Representation in absolute numbers of the main parenchymal TB10.4-specific CD8^+^ T cell subsets induced during *M. tuberculosis* infection of *Sp140^+/+^* (n=5, mean number of total TB10.4-specific CD8^+^ T cell: 37,991) and *Sp140^-/-^* treated (n=10, mean number of total TB10.4-specific CD8^+^ T cell: 40,962) or not (n=4, mean number of total TB10.4-specific CD8^+^ T cell: 4,759) with anti-IFNAR on day 28. Stacked representation (left) and individual subpopulation (right) are shown. Each data point corresponds to one individual mouse. Bars and error bars indicate the mean ± standard deviation. Unpaired two-way ANOVA tests with Tukey correction were used. A representative experiment is shown alongside two independent ones. B. As in A, but for vascular CD8^+^ T cell subsets. *Sp140^+/+^* (n=5, mean number of total CD8^+^ T cell: 152,646) and *Sp140^-/-^* treated (n=10, mean number of total CD8^+^ T cell: 176,559) or not (n=4, mean number of total CD8^+^ T cell: 5,283) with anti-IFNAR on day 28 C. As in A, but for vascular TB10.4-specific CD8^+^ T cell subsets. *Sp140^+/+^* (n=5, mean number of total TB10.4-specific CD8^+^ T cell: 14,205) and *Sp140^-/-^* treated (n=10, mean number of total TB10.4-specific CD8^+^ T cell: 25,612) or not (n=4, mean number of total TB10.4-specific CD8^+^ T cell: 1,424) with anti-IFNAR on day 28. The color code for individual CD8^+^ T cell subsets in panels A-C is identical to that used in Figure 1. D. UMAP representation of live lung cells based on spectral cytometry data, with CD8 expression displayed as a continuous color scale. Samples include equal numbers of cells from *Sp140^+/+^* and anti-IFNAR-treated or not *Sp140^-/-^* mice. Quantification of the CD8-expressing cell cluster is shown (right). Bars and error bars indicate the mean ± standard deviation. Unpaired one-way ANOVA tests with Tukey correction were used. E. As in D but for CD4 expression.

**Supplemental Figure 8 (related to Figure 3).**
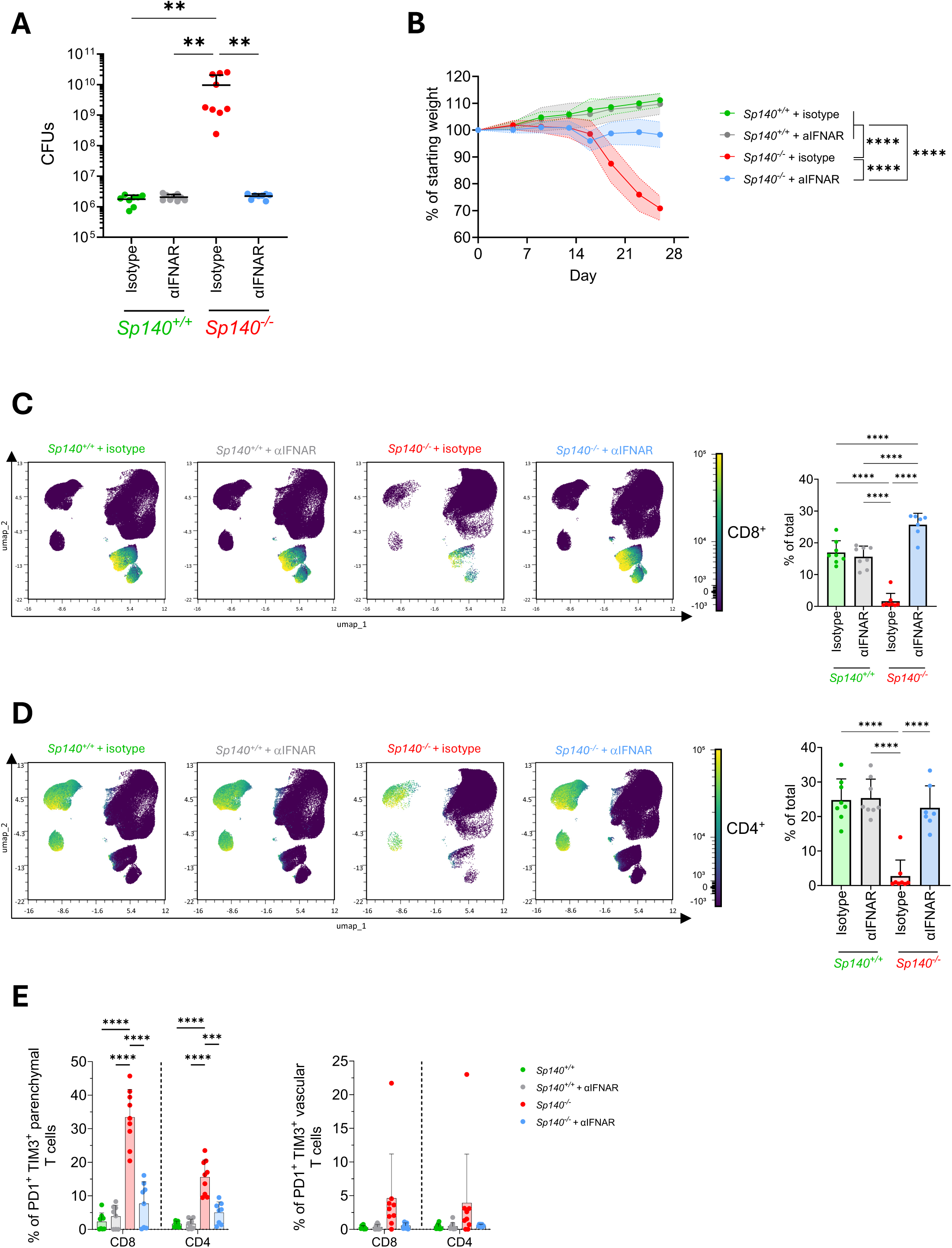
Impact of IFN-I on T cells from mice infected with *M. tuberculosis. Sp140^+/+^* and *Sp140^-/-^* mice were infected with 100 CFUs of the Erdman strain of *M. tuberculosis* and treated between days 7 and 27 post-infection with either an isotype control or a blocking monoclonal antibody (mAb) against IFNAR. Lungs were collected and analyzed at day 28 post-infection. A representative experiment among two is shown. A. Pulmonary bacterial loads at 28 days post-infection in isotype-treated-*Sp140^+/+^* (n=8), anti-IFNAR-treated *Sp140^+/+^* (n=9), isotype-treated *Sp140^-/-^* (n=9) and anti-IFNAR-treated *Sp140^-/-^* mice (n=8). Each data point corresponds to one individual mouse. Bars and error bars indicate the mean± standard deviation. One-way ANOVA tests with Tukey correction were used. B. Weight monitoring from day 0 to day 26 in the same groups of mice. Body weight was monitored over time and expressed as a percentage of the initial body weight (baseline set to 100%). The shaded area corresponds to the standard deviation. One-way ANOVA tests with Tukey correction were used at day 26. C. UMAP representation of live lung cells based on spectral cytometry data, with CD8 expression displayed as a continuous color scale. Samples include equal numbers of cells from *Sp140^+/+^* and *Sp140^-/-^* (isotype or anti-IFNAR treated) mice. Quantification of the CD8-expressing cell cluster is shown (bottom). Unpaired ANOVA tests with Tukey correction were used for statistical analysis. D. As in C but for CD4 expression. E. Representation in percentages of the parenchymal (left) and vascular (right) lung CD8^+^ and CD4^+^ T cells displaying an exhausted phenotype PD1^+^TIM3^+^. Bars and error bars indicate the mean ± standard deviation. Two-way ANOVA tests with Tukey correction was used.

**Supplemental Figure 9 (related to Figure 4).**
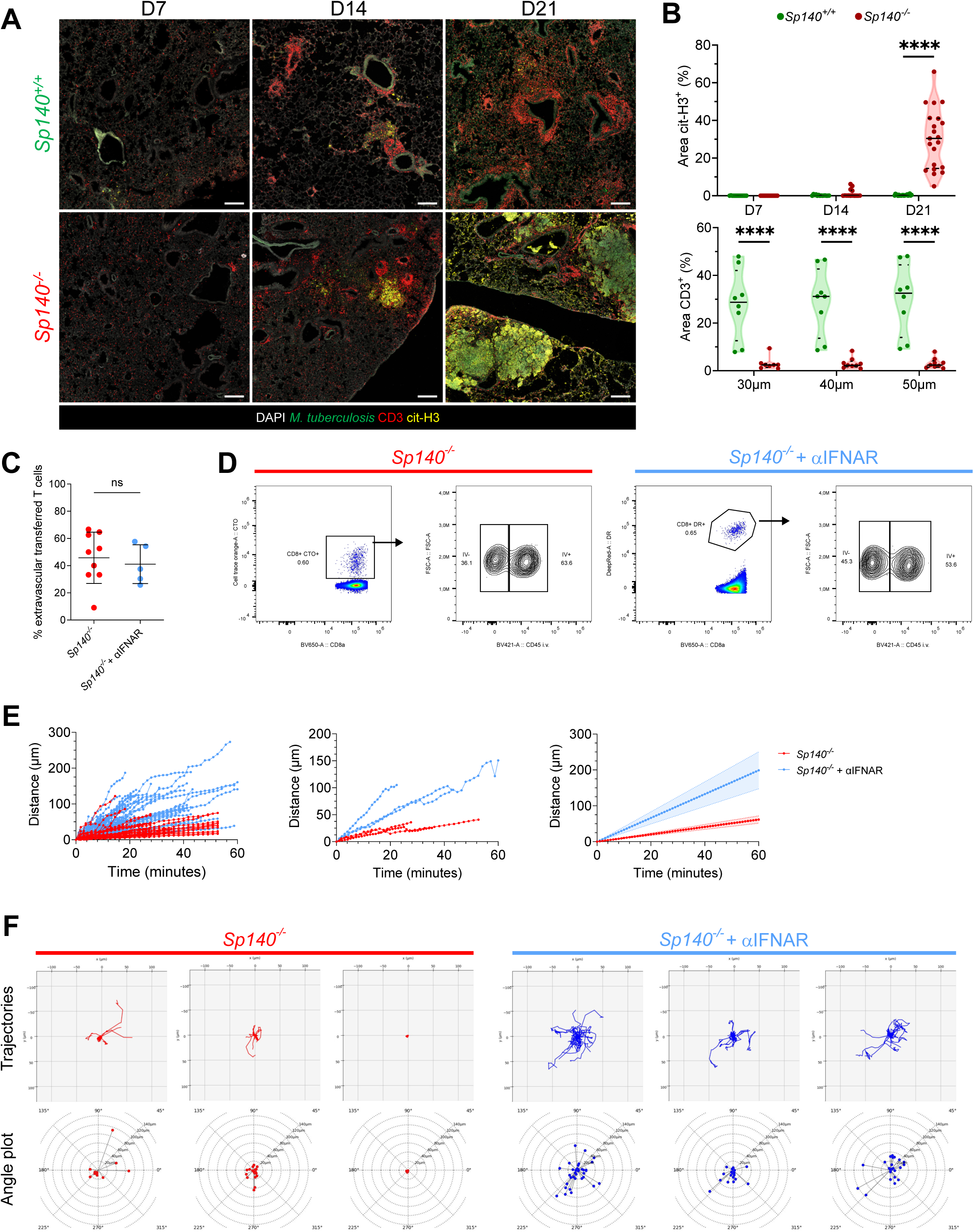
Imaging T cell positioning and mobility in *M.tuberculosis*-infected lungs. A. Representative confocal images of lung sections (20 µm) from *Sp140^+/+^* and *Sp140^-/-^* mice, infected with *M. tuberculosis*-GFP. NETs (citrullinated histone H3) in yellow, T cells (CD3^+^) in red and nuclei (DAPI) in white are shown at day 7 (left), day 14 (middle), and day 21 (right) post infection. Scale bar, 500 µm. B. Associated quantification of the NETs area is shown (top) and T cell exclusion from bacteria lesions (right). Briefly, polygons were drawn around the bacterial foci and, using the “expand annotation” tool in QuPath, they were sequentially extended by a radius of 30µm, 40µm, and 50µm. Quantification of NETs and CD3^+^ T cell areas is shown (right). Each data point corresponds to measurements taken from distinct microscopic fields of view from at least 2 different mice per group. C. Percentage of transferred, extravascular T cells analyzed based on the positioning of T cells respective to vessels identified by fluorescent dextran and total tracked T cells. Each dot represents one individual mouse. D. Proportions of transferred CD8^+^ T cells present in intra- *vs.* extra- vascular compartments recovered from MP-IVM imaging and after administration of a fluorescent anti-CD45 antibody prior to lung collection. A representative experiment is shown. E. Cumulative displacement analysis of transferred CD8^+^ T cells in infected lungs. Left: individual tracks are shown. Middle: mean value of per animal tracks. Right: Extrapolation of the linear regression for each mouse to standardize the time points kinetics from 1 to 60 min, with 1 min between each acquisition. Non parametric Mann-Whitney test was used. *Sp140*^-/-^ group: n = 6 mice, 111 tracked cells in total, with 16, 6, 25, 33, 23, and 8 tracked cells per mouse. *Sp140*^-/-^ + anti-IFNAR group: n = 3 mice, 115 tracked cells in total, with 35, 23, and 57 tracked cells per mouse. Data were acquired from two independent experiments. F. CD8^+^ T cell tracks, plotted from a common origin, from *Sp140^-/-^* (left, red) and *Sp140^-/-^* treated with anti-IFNAR (right, blue) from 3 representative mice across 2 independent experiments. Graphs show either the tracks of individual T cells tracked over time by intravital microscopy (solid colored line, top), illustrating cell displacement, confinement, and directional migration within the tissue, or the final position of individual T cells tracked over time by intravital microscopy (colored circle, bottom), illustrating a measure of directional persistence (angle plots).

**Supplemental Figure 10 (related to Figure 4).**
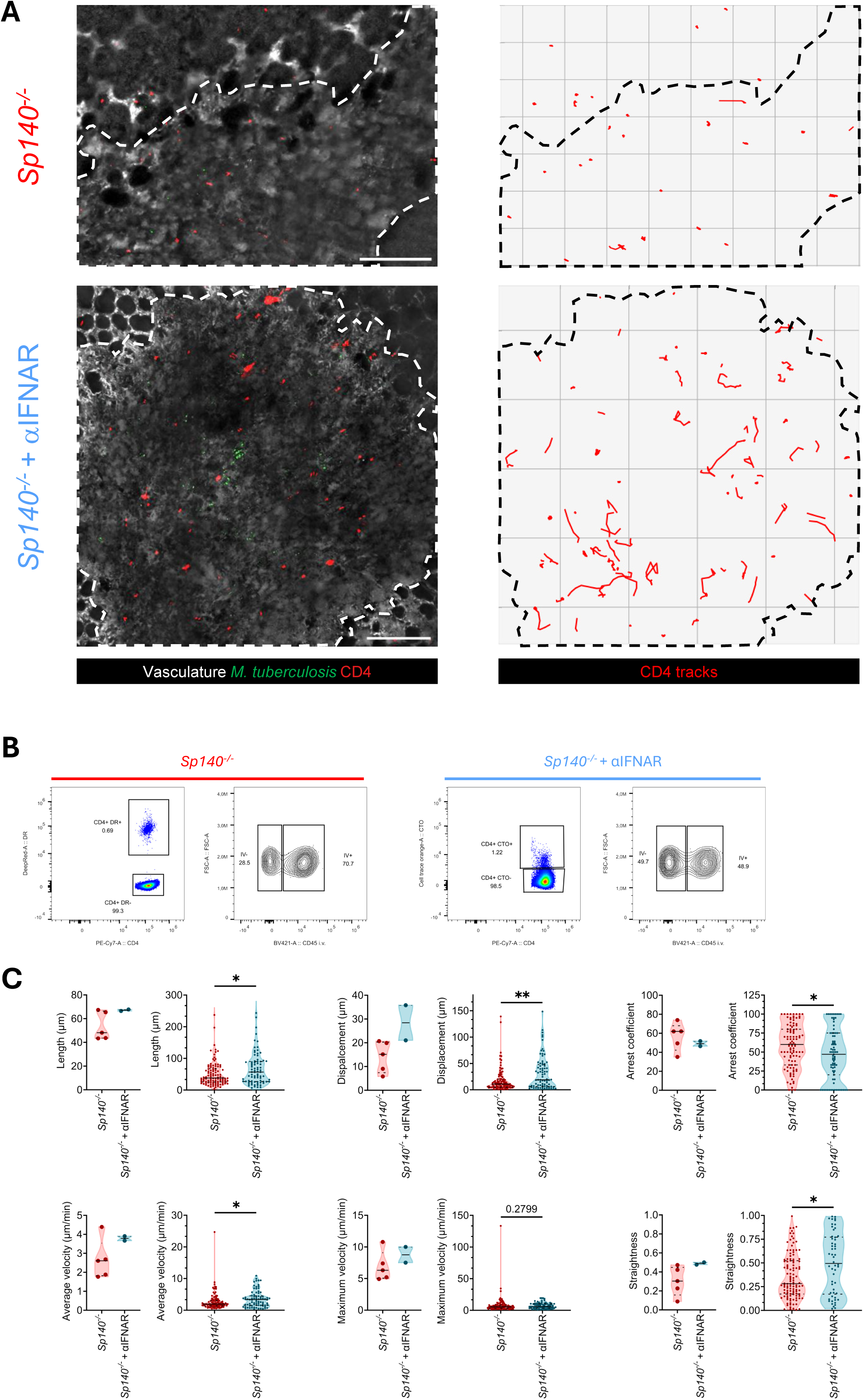
Imaging CD4^+^ T cell positioning and mobility in *M. tuberculosis*-infected lungs. A. Representative intravital microscopy images of lungs from *M. tuberculosis*-mTurquoise-infected mice. Bacteria appear in green, transferred CD4^+^ T cells are in red, and lesion boundaries, identified by dextran staining and *M. tuberculosis* fluorescence, are outlined by a dotted line. Images show *Sp140^-/-^* mice (day 28 post infection, top) and anti-IFNAR-treated *Sp140^-/-^* mice (day 27 post infection, bottom). The first frame of the video is shown (left), with corresponding CD4^+^ T cell tracks over the full acquisition period (right). Scale bar, 200 µm. B. Proportions of transferred CD4^+^ T cells present in intra- *vs.* extra- vascular compartments recovered from MP-IVM imaging and after administration of a fluorescent anti-CD45 antibody prior to lung collection. A representative experiment is shown. C. Quantification of CD4^+^ T cell motility parameters during intravital imaging. Per-animal motility (left) and pooled track (right) measurements for transferred CD4^+^ T cells in infected lungs are shown for each measured parameter. For per-animal motility analysis, each dot represents the mean value of the measured parameters of CD4^+^ T cell motility in a single mouse. For pooled tracks analysis, each dot represents a single tracked transferred CD4^+^ T cells. Parameters include track length, displacement, arrest coefficient (% of of time T cell instantaneous speed < 2 µm/min), average and maximum velocity and straightness. Non parametric Mann-Whitney test was used. *Sp140^-/-^* group: n = 5 mice, with 19, 27, 37, 27, and 6 tracked cells per mouse, respectively, for a total of 116 tracked cells. *Sp140^-/-^* + anti-IFNAR group: n = 2 mice, with 8 and 46 tracked cells per mouse, respectively, for a total of 54 tracked cells. Data were acquired from two independent experiments. Because only two anti-IFNAR-treated mice were imaged, statistical analysis was not performed at the animal level.

## Legend to Supplemental Movies

**Supplemental Movie 1.** Arrested CD8^+^ T cell migration in *Sp140*^-/-^ mice.

Lung intravital microscopy reveals severely restricted migratory behavior of CD8^+^ T cells in *Sp140*^-/-^ mice. Multiphoton intravital imaging was performed in *M. tuberculosis*-infected *Sp140*^-/-^ mice (day 21 post-infection) following adoptive transfer of DeepRed^+^ CD8^+^ T cells (red). The lung vasculature is visualized using Cascade Blue or Texas Red–dextran (grey), and bacteria are identified by mTurquoise expression (green). CD8^+^ T cell motility was tracked over a multi-tile, 2D maximum intensity projection time-lapse sequence (tracks in cyan). Time is displayed as h:min:s. Scale bar, 200 µm.

**Supplemental Movie 2.** Restored CD8^+^ T cell migratory dynamics in anti-IFNAR–treated *Sp140*^-/-^ mice.

Lung intravital microscopy shows that anti-IFNAR treatment restores dynamic CD8^+^ T cell migration within lesions in *Sp140*^-/-^ mice. Multiphoton intravital imaging was performed in *M. tuberculosis*-infected *Sp140*^-/-^ mice (day 28 post-infection), treated with anti-IFNAR, following adoptive transfer of DeepRed^+^ CD8^+^ T cells (red). The lung vasculature is visualized using Cascade Blue or Texas Red–dextran (grey), and bacteria are identified by mTurquoise expression (green). CD8^+^ T cell motility was tracked over a multi-tile, 2D maximum intensity projection time-lapse sequence (tracks in cyan). Time is displayed as h:min:s. Scale bar, 200 µm.

**Supplemental Movie 3.** Arrested CD4^+^ T cell migration in *Sp140*^-/-^ mice.

Lung intravital microscopy reveals severely restricted migratory behavior of CD4^+^ T cells in *Sp140*^-/-^ mice. Multiphoton intravital imaging was performed in *M. tuberculosis*-infected *Sp140*^-/-^ mice (day 28 post-infection) following adoptive transfer of DeepRed^+^ CD4^+^ T cells (red). The lung vasculature is visualized using Cascade Blue or Texas Red–dextran (grey), and bacteria are identified by mTurquoise expression (green). A multi-tile, 2D maximum intensity projection time-lapse sequence (tracks in cyan). Time is displayed as h:min:s. Scale bar, 200 µm.

**Supplemental Movie 4.** Restored CD4^+^ T cell migratory dynamics in anti-IFNAR–treated *Sp140*^-/-^ mice.

Lung intravital microscopy shows that anti-IFNAR treatment restores dynamic CD4^+^ T cell migration within lesions in *Sp140*^-/-^ mice. Multiphoton intravital imaging was performed in *M. tuberculosis*-infected *Sp140*^-/-^ mice (day 27 post-infection) treated with anti-IFNAR following adoptive transfer of DeepRed^+^ CD4^+^ T cells (red). The lung vasculature is visualized using Cascade Blue or Texas Red–dextran (grey), and bacteria are identified by mTurquoise expression (green). CD4^+^ T cell motility was tracked over a multi-tile, 2D maximum intensity projection time-lapse sequence (tracks in cyan). Time is displayed as h:min:s. Scale bar, 200 µm.

**Supplemental Movie 5.** Neutrophil migration in TB lesion of *Sp140^-/-^* mice.

Representative example of intravital multiphoton microscopy of neutrophils labeled with anti-Ly6G (green) injected intravenously in *Sp140^-/-^* mice imaged in healthy, bacteria-rich, or mixed lung regions. The lesion area is outlined by a white dotted line. Time is displayed as min:s. Scale bar, 50 µm.

**Supplemental Movie 6.** Vascular remodeling and perfusion in lung TB lesions.

Vascular architecture within lung TB lesions following intravenous injection of Cascade Blue or Texas Red-dextran (grey). The lesion area is outlined by a white dotted line. White asterisks show normal alveoli with no dextran leakage, and green asterisks show dextran-filled alveoli due to leakage and vascular permeability. The red arrows show perfused blood vessels. Time is displayed as min:s. Scale bar, 50 µm.

**Supplemental Movie 7.** Spatial distribution of bacteria in lung TB lesions.

Vascular architecture within lung TB lesions following intravenous injection of Cascade Blue or Texas Red-dextran (grey). *M. tuberculosis* is visualized by mTurquoise expression (green). The white dashed line indicates vascular remodeling within the lesion area. White asterisks indicate normal alveoli with no leakage, and red asterisks indicate dextran-filled alveoli with leakage and vascular permeability. Time is displayed as min:s. Scale bar, 50 µm.

## Notes

### Competing Interest Statement

The authors have declared no competing interest.

### Summary of Updates

Summary of major revisions to the manuscript We strengthened the notion that the observed T cell phenotype reflects a secondary effects imposed by the inflammatory tissue environment. Sp140-deficient T cells transferred into T cell-deficient hosts activated and accumulated in the lung after M. tuberculosis infection in a manner comparable to Sp140-competent T cells. These data further argue against an overt T cell-intrinsic defect caused by loss of Sp140, at least for the activation and accumulation parameters assessed. Our findings support a model in which Sp140 deficiency primarily promotes an IFN-I-driven pathological lung environment that secondarily constrains T cell accumulation, positioning, motility, and function. This model reconciles the apparent discrepancy between endogenous T cell phenotypes and adoptive-transfer imaging experiments: the dominant defect is not an obligate T cell-intrinsic failure, but rather an IFN-I-dependent tissue environment that imposes secondary constraints on otherwise competent T cells. We also substantially revised the intravital imaging component of the study. Imaging datasets were reprocessed to improve visualization, annotation, and interpretability. Additional validation and control videos, including dextran perfusion and Ly6G labeling, were added to better document vascular perfusion, tissue viability, and inflammatory-cell localization. We also reanalyzed T cell motility using animal-level statistics rather than pooled tracks alone. Individual tracks are now displayed by mouse and, where possible, by lesion, thereby highlighting biological heterogeneity while ensuring that statistical inference is based on independent biological replicates. Overall, the new experiments, revised analyses, added controls, and textual clarifications strengthen the mechanistic framework of the study. The revised manuscript now presents a clearer model in which Sp140 deficiency promotes an IFN-I-driven pathological lung environment that secondarily constrains T cell positioning, motility, accumulation, and function.

