## Supplementary material for "Type I interferon–driven lung pathology restricts T cell accumulation and motility during tuberculosis": Supp Tables

| Peptide sequence | Protein of origin | MHC | Reference |
|---|---|---|---|
| IMYNYPAM | TB10.4 | MHC-I | Woodworth J.S., Journal of Immunology, 2008 |
| GAPINSATAM | Mtb32A | MHC-I | Woodworth J.S., Journal of Immunology, 2008 |
| FAVTNDGVI | MPT64 | MHC-I | Doi T., Journal of Immunology, 2007 |
| SGVGNDLVL | Psts-3 | MHC-I | Romano M., Journal of Immunology, 2004 |
| AAIGNMTLL | PPE44 | MHC-I | Romano M., Vaccine, 2008 |
| INYEYAIV | Psts-1 | MHC-I | Zhu X., International Immunology, 1997 |
| INYLVPFL | CFP11 | MHC-I | Eweda G., Vaccine, 2010 |
| RMVINYLVPFL | CFP11 | MHC-I | Eweda G., Vaccine, 2010 |
| VGSLNGTYV | CFP17 | MHC-I | Eweda G., Vaccine, 2010 |
| GTYVNREPV | CFP17 | MHC-I | Eweda G., Vaccine, 2010 |
| FQDAYNAAGGHNAVF | Ag85 | MHC-II | S. Yanagisawa, International Immunology, 1997 |
| EQQWNFAGIEAAASA | ESAT6 | MHC-II | Brandt L., Journal of Immunology, 1996 |

Table S1

### Table S2

| cluster | gene | p_val | avg_log2FC | pct.1 | pct.2 | p_val_adj |
|---|---|---|---|---|---|---|
| 0 | Ccl5 | 0 | 1,068 | 0,995 | 0,926 | 0 |
| 0 | Itgb1 | 2,29E-306 | 0,956 | 0,829 | 0,637 | 4,25E-302 |
| 0 | S100a10 | 0 | 0,903 | 0,993 | 0,890 | 0 |
| 0 | S100a4 | 3,75E-160 | 0,855 | 0,742 | 0,564 | 6,98E-156 |
| 0 | Klrg1 | 4,32E-135 | 0,849 | 0,476 | 0,300 | 8,04E-131 |
| 0 | Cxcr3 | 8,71E-240 | 0,743 | 0,777 | 0,580 | 1,62E-235 |
| 0 | P2rx7 | 5,88E-192 | 0,710 | 0,580 | 0,357 | 1,09E-187 |
| 0 | Emp3 | 7,34E-267 | 0,681 | 0,880 | 0,703 | 1,36E-262 |
| 0 | Ifitm10 | 2,13E-166 | 0,678 | 0,622 | 0,432 | 3,97E-162 |
| 0 | Pdcd4 | 0 | 0,676 | 0,939 | 0,821 | 0 |
| 0 | Ahnak | 4,69E-289 | 0,661 | 0,964 | 0,841 | 8,72E-285 |
| 0 | Crip1 | 5,99E-245 | 0,645 | 0,935 | 0,801 | 1,11E-240 |
| 0 | Arhgdib | 0 | 0,644 | 0,998 | 0,961 | 0 |
| 0 | Adgre5 | 3,76E-255 | 0,641 | 0,921 | 0,803 | 6,99E-251 |
| 0 | Cd52 | 0 | 0,634 | 0,999 | 0,970 | 0 |
| 0 | Zyx | 4,33E-285 | 0,618 | 0,934 | 0,834 | 8,05E-281 |
| 0 | Mxd4 | 2,09E-165 | 0,589 | 0,662 | 0,488 | 3,89E-161 |
| 0 | Apbb1ip | 2,95E-263 | 0,589 | 0,919 | 0,833 | 5,48E-259 |
| 0 | Cd226 | 6,69E-215 | 0,584 | 0,877 | 0,764 | 1,24E-210 |
| 0 | Cd8b1 | 0 | 0,583 | 0,987 | 0,922 | 0 |
| 0 | Lsp1 | 5,70E-305 | 0,572 | 0,982 | 0,916 | 1,06E-300 |
| 0 | Tmsb4x | 0 | 0,560 | 0,999 | 0,996 | 0 |
| 0 | Ucp2 | 9,33E-297 | 0,556 | 0,989 | 0,947 | 1,73E-292 |
| 0 | Hcst | 4,64E-210 | 0,553 | 0,845 | 0,723 | 8,63E-206 |
| 0 | Ptpn18 | 2,46E-239 | 0,551 | 0,942 | 0,842 | 4,57E-235 |
| 0 | Sh3bgrl3 | 1,95E-275 | 0,550 | 0,984 | 0,907 | 3,61E-271 |
| 0 | Rac2 | 0 | 0,538 | 0,998 | 0,976 | 0 |
| 0 | Rnf166 | 1,96E-216 | 0,538 | 0,901 | 0,795 | 3,64E-212 |
| 0 | Sh2d1a | 2,54E-145 | 0,534 | 0,791 | 0,679 | 4,71E-141 |
| 0 | Gzmk | 1,89E-162 | 0,527 | 0,825 | 0,619 | 3,51E-158 |
| 1 | AA467197 | 0 | 1,757 | 0,875 | 0,405 | 0 |
| 1 | Gzmb | 8,38E-292 | 1,725 | 0,855 | 0,668 | 1,56E-287 |

| | | | | | | |
|---|---|---|---|---|---|---|
| 1 | Glrx | 0 | 1,529 | 0,955 | 0,584 | 0 |
| 1 | Ifitm2 | 3,63E-180 | 1,428 | 0,277 | 0,093 | 6,75E-176 |
| 1 | Gzma | 6,46E-25 | 1,368 | 0,583 | 0,500 | 1,20E-20 |
| 1 | Prf1 | 0 | 1,356 | 0,957 | 0,721 | 0 |
| 1 | Lag3 | 0 | 1,232 | 0,903 | 0,579 | 0 |
| 1 | S100a6 | 0 | 1,202 | 0,896 | 0,569 | 0 |
| 1 | Ctla2a | 0 | 1,131 | 0,986 | 0,772 | 0 |
| 1 | Id2 | 0 | 1,116 | 0,997 | 0,877 | 0 |
| 1 | Fgl2 | 0 | 1,104 | 0,953 | 0,656 | 0 |
| 1 | Vim | 0 | 1,069 | 0,995 | 0,917 | 0 |
| 1 | Acadl | 0 | 1,059 | 0,900 | 0,636 | 0 |
| 1 | S100a4 | 4,86E-203 | 1,004 | 0,803 | 0,562 | 9,03E-199 |
| 1 | Ly6a | 0 | 0,990 | 0,998 | 0,917 | 0 |
| 1 | Ifitm3 | 1,47E-167 | 0,982 | 0,366 | 0,155 | 2,73E-163 |
| 1 | Anxa2 | 0 | 0,978 | 0,953 | 0,656 | 0 |
| 1 | Cxcr6 | 0 | 0,972 | 0,939 | 0,756 | 0 |
| 1 | Hlx | 0 | 0,945 | 0,514 | 0,152 | 0 |
| 1 | Havcr2 | 0 | 0,909 | 0,724 | 0,300 | 0 |
| 1 | Entpd1 | 0 | 0,903 | 0,793 | 0,395 | 0 |
| 1 | Ctsd | 0 | 0,896 | 0,979 | 0,816 | 0 |
| 1 | Ccr5 | 0 | 0,879 | 0,990 | 0,807 | 0 |
| 1 | Icos | 1,62E-252 | 0,847 | 0,951 | 0,766 | 3,01E-248 |
| 1 | Gem | 2,95E-254 | 0,832 | 0,749 | 0,433 | 5,48E-250 |
| 1 | Cst7 | 0 | 0,832 | 0,962 | 0,723 | 0 |
| 1 | Mxd1 | 1,25E-196 | 0,807 | 0,870 | 0,644 | 2,32E-192 |
| 1 | Capg | 1,67E-269 | 0,805 | 0,681 | 0,353 | 3,10E-265 |
| 1 | Gstt1 | 7,67E-161 | 0,797 | 0,366 | 0,156 | 1,43E-156 |
| 1 | Sub1 | 0 | 0,796 | 0,981 | 0,810 | 0 |
| 2 | Ccl3 | 0 | 3,713 | 0,822 | 0,344 | 0 |
| 2 | Xcl1 | 1,35E-131 | 3,264 | 0,308 | 0,129 | 2,51E-127 |
| 2 | Ccl4 | 0 | 3,072 | 0,927 | 0,701 | 0 |
| 2 | Tnfrsf9 | 0 | 2,673 | 0,887 | 0,273 | 0 |
| 2 | Nr4a3 | 0 | 2,525 | 0,665 | 0,160 | 0 |

| | | | | | | |
|---|---|---|---|---|---|---|
| 2 | Rgs16 | 0 | 2,495 | 0,903 | 0,310 | 0 |
| 2 | Ifng | 0 | 2,455 | 0,931 | 0,670 | 0 |
| 2 | Il2ra | 0 | 2,076 | 0,826 | 0,343 | 0 |
| 2 | Irf8 | 0 | 2,067 | 0,879 | 0,334 | 0 |
| 2 | Nr4a1 | 0 | 1,982 | 0,764 | 0,390 | 0 |
| 2 | Egr3 | 0 | 1,902 | 0,413 | 0,077 | 0 |
| 2 | Nr4a2 | 0 | 1,793 | 0,728 | 0,284 | 0 |
| 2 | Tnfrsf4 | 0 | 1,781 | 0,605 | 0,139 | 0 |
| 2 | Sema7a | 0 | 1,630 | 0,778 | 0,217 | 0 |
| 2 | Havcr2 | 0 | 1,623 | 0,794 | 0,304 | 0 |
| 2 | Lag3 | 0 | 1,603 | 0,953 | 0,582 | 0 |
| 2 | Cdkn1a | 0 | 1,589 | 0,635 | 0,172 | 0 |
| 2 | Gzmb | 1,42E-268 | 1,571 | 0,891 | 0,669 | 2,63E-264 |
| 2 | Tnfsf14 | 0 | 1,562 | 0,501 | 0,173 | 0 |
| 2 | Slc7a5 | 0 | 1,549 | 0,754 | 0,314 | 0 |
| 2 | Rilpl2 | 0 | 1,513 | 0,881 | 0,453 | 0 |
| 2 | Prf1 | 0 | 1,473 | 0,926 | 0,733 | 0 |
| 2 | Myc | 2,08E-192 | 1,430 | 0,627 | 0,349 | 3,87E-188 |
| 2 | Tnf | 1,25E-195 | 1,382 | 0,489 | 0,212 | 2,33E-191 |
| 2 | Irf4 | 0 | 1,379 | 0,781 | 0,303 | 0 |
| 2 | Egr2 | 0 | 1,362 | 0,443 | 0,075 | 0 |
| 2 | Nfkbid | 2,65E-169 | 1,354 | 0,618 | 0,349 | 4,93E-165 |
| 2 | Egr1 | 1,68E-267 | 1,308 | 0,654 | 0,307 | 3,12E-263 |
| 2 | Gadd45b | 5,87E-278 | 1,307 | 0,770 | 0,436 | 1,09E-273 |
| 2 | Crtam | 1,16E-143 | 1,283 | 0,503 | 0,280 | 2,15E-139 |
| 3 | Rsrp1 | 0 | 1,229 | 0,937 | 0,873 | 0 |
| 3 | Itgae | 2,83E-100 | 1,204 | 0,454 | 0,272 | 5,26E-96 |
| 3 | Fosb | 9,40E-50 | 1,131 | 0,469 | 0,365 | 1,75E-45 |
| 3 | mt-Co2 | 1,72E-292 | 0,937 | 0,985 | 0,973 | 3,20E-288 |
| 3 | mt-Co3 | 2,43E-263 | 0,936 | 0,951 | 0,918 | 4,52E-259 |
| 3 | Neurl3 | 1,88E-58 | 0,927 | 0,618 | 0,550 | 3,50E-54 |
| 3 | Gabbr1 | 2,79E-193 | 0,912 | 0,745 | 0,593 | 5,19E-189 |
| 3 | mt-Cytb | 1,20E-216 | 0,904 | 0,915 | 0,838 | 2,23E-212 |

| | | | | | | |
|---|---|---|---|---|---|---|
| 3 | Ogt | 8,41E-294 | 0,902 | 0,898 | 0,864 | 1,56E-289 |
| 3 | mt-Atp6 | 3,88E-267 | 0,897 | 0,973 | 0,958 | 7,22E-263 |
| 3 | Itga1 | 1,23E-102 | 0,896 | 0,574 | 0,410 | 2,28E-98 |
| 3 | Vmp1 | 1,15E-220 | 0,892 | 0,910 | 0,894 | 2,14E-216 |
| 3 | Slc20a1 | 2,70E-159 | 0,885 | 0,792 | 0,706 | 5,01E-155 |
| 3 | Dennd4a | 9,56E-175 | 0,884 | 0,835 | 0,763 | 1,78E-170 |
| 3 | Adcy7 | 2,12E-247 | 0,876 | 0,852 | 0,781 | 3,93E-243 |
| 3 | Tnfaip3 | 2,48E-239 | 0,873 | 0,954 | 0,956 | 4,60E-235 |
| 3 | H2-Q6 | 0 | 0,855 | 0,980 | 0,972 | 0 |
| 3 | mt-Co1 | 1,69E-253 | 0,848 | 0,975 | 0,956 | 3,14E-249 |
| 3 | Baiap3 | 4,79E-109 | 0,837 | 0,649 | 0,515 | 8,91E-105 |
| 3 | P2rx7 | 2,44E-105 | 0,826 | 0,561 | 0,378 | 4,53E-101 |
| 3 | mt-Nd4l | 4,16E-180 | 0,817 | 0,841 | 0,723 | 7,74E-176 |
| 3 | Cdh1 | 9,70E-91 | 0,802 | 0,398 | 0,228 | 1,80E-86 |
| 3 | Arglu1 | 3,66E-207 | 0,799 | 0,844 | 0,759 | 6,79E-203 |
| 3 | Lilrb4a | 1,67E-91 | 0,792 | 0,735 | 0,631 | 3,10E-87 |
| 3 | Sorl1 | 1,32E-275 | 0,780 | 0,920 | 0,905 | 2,46E-271 |
| 3 | Ccnl2 | 1,37E-213 | 0,773 | 0,833 | 0,762 | 2,54E-209 |
| 3 | mt-Nd1 | 2,84E-194 | 0,768 | 0,943 | 0,909 | 5,28E-190 |
| 3 | Adgrg5 | 2,72E-185 | 0,765 | 0,873 | 0,824 | 5,06E-181 |
| 3 | mt-Atp8 | 4,04E-229 | 0,763 | 0,978 | 0,974 | 7,51E-225 |
| 3 | Clk1 | 1,45E-271 | 0,759 | 0,928 | 0,919 | 2,70E-267 |
| 4 | Sell | 0 | 3,144 | 0,982 | 0,217 | 0 |
| 4 | Ccr7 | 0 | 2,910 | 0,907 | 0,127 | 0 |
| 4 | Igfbp4 | 0 | 2,734 | 0,602 | 0,058 | 0 |
| 4 | Ighm | 0 | 2,421 | 0,899 | 0,118 | 0 |
| 4 | Klf2 | 0 | 2,174 | 0,975 | 0,465 | 0 |
| 4 | Tcf7 | 0 | 2,167 | 0,987 | 0,368 | 0 |
| 4 | Lef1 | 0 | 2,032 | 0,966 | 0,447 | 0 |
| 4 | S1pr1 | 0 | 1,985 | 0,910 | 0,309 | 0 |
| 4 | Dapl1 | 0 | 1,828 | 0,402 | 0,028 | 0 |
| 4 | Pde2a | 0 | 1,756 | 0,770 | 0,209 | 0 |
| 4 | Nsg2 | 0 | 1,732 | 0,774 | 0,081 | 0 |

| | | | | | | |
|---|---|---|---|---|---|---|
| 4 | Actn1 | 0 | 1,729 | 0,861 | 0,308 | 0 |
| 4 | Treml2 | 0 | 1,577 | 0,788 | 0,171 | 0 |
| 4 | Il6st | 0 | 1,552 | 0,892 | 0,454 | 0 |
| 4 | Eef1b2 | 0 | 1,513 | 0,997 | 0,878 | 0 |
| 4 | Cmah | 0 | 1,494 | 0,826 | 0,165 | 0 |
| 4 | Il6ra | 0 | 1,398 | 0,599 | 0,058 | 0 |
| 4 | Dgka | 0 | 1,375 | 0,985 | 0,836 | 0 |
| 4 | Fam241a | 0 | 1,359 | 0,860 | 0,521 | 0 |
| 4 | Tpt1 | 0 | 1,344 | 0,999 | 0,974 | 0 |
| 4 | Sh3bp5 | 0 | 1,323 | 0,677 | 0,119 | 0 |
| 4 | Gramd4 | 0 | 1,321 | 0,852 | 0,339 | 0 |
| 4 | Socs3 | 0 | 1,260 | 0,831 | 0,420 | 0 |
| 4 | Tmsb10 | 0 | 1,239 | 0,997 | 0,942 | 0 |
| 4 | Eef1g | 0 | 1,234 | 0,962 | 0,748 | 0 |
| 4 | Evl | 0 | 1,134 | 0,978 | 0,696 | 0 |
| 4 | Tdrp | 0 | 1,117 | 0,530 | 0,062 | 0 |
| 4 | Dph5 | 0 | 1,095 | 0,688 | 0,247 | 0 |
| 4 | St6gal1 | 0 | 1,095 | 0,546 | 0,048 | 0 |
| 4 | Satb1 | 0 | 1,080 | 0,983 | 0,792 | 0 |
| 5 | Ifit3 | 0 | 2,627 | 0,885 | 0,458 | 0 |
| 5 | Ifit1 | 0 | 2,310 | 0,838 | 0,333 | 0 |
| 5 | Cxcl10 | 0 | 2,290 | 0,412 | 0,113 | 0 |
| 5 | Rsad2 | 0 | 2,148 | 0,734 | 0,210 | 0 |
| 5 | Isg15 | 0 | 2,122 | 0,874 | 0,506 | 0 |
| 5 | Ifit1bl1 | 0 | 2,070 | 0,944 | 0,645 | 0 |
| 5 | Slfn5 | 0 | 2,028 | 0,607 | 0,164 | 0 |
| 5 | Ifit2 | 0 | 1,890 | 0,761 | 0,393 | 0 |
| 5 | Daxx | 0 | 1,831 | 0,891 | 0,633 | 0 |
| 5 | Cmpk2 | 0 | 1,749 | 0,704 | 0,217 | 0 |
| 5 | Isg20 | 0 | 1,711 | 0,916 | 0,637 | 0 |
| 5 | Rnf213 | 0 | 1,673 | 0,963 | 0,858 | 0 |
| 5 | Gbp2 | 0 | 1,646 | 0,963 | 0,779 | 0 |
| 5 | Phf11b | 0 | 1,639 | 0,913 | 0,688 | 0 |

| | | | | | | |
|---|---|---|---|---|---|---|
| 5 | Usp18 | 0 | 1,606 | 0,832 | 0,493 | 0 |
| 5 | Bst2 | 0 | 1,596 | 0,944 | 0,782 | 0 |
| 5 | Oas3 | 0 | 1,581 | 0,866 | 0,515 | 0 |
| 5 | Samhd1 | 0 | 1,561 | 0,985 | 0,929 | 0 |
| 5 | Ifih1 | 0 | 1,518 | 0,787 | 0,370 | 0 |
| 5 | Plac8 | 0 | 1,516 | 0,906 | 0,823 | 0 |
| 5 | Ifi209 | 0 | 1,510 | 0,965 | 0,819 | 0 |
| 5 | Ifi208 | 0 | 1,470 | 0,789 | 0,417 | 0 |
| 5 | Irf7 | 0 | 1,453 | 0,915 | 0,664 | 0 |
| 5 | Dhx58 | 0 | 1,443 | 0,814 | 0,438 | 0 |
| 5 | Ifit3b | 0 | 1,425 | 0,665 | 0,160 | 0 |
| 5 | Nt5c3 | 5,05E-268 | 1,423 | 0,733 | 0,527 | 9,38E-264 |
| 5 | Slfn1 | 0 | 1,419 | 0,925 | 0,721 | 0 |
| 5 | Trafd1 | 0 | 1,402 | 0,895 | 0,670 | 0 |
| 5 | Zbp1 | 0 | 1,389 | 0,979 | 0,855 | 0 |
| 5 | Rtp4 | 0 | 1,386 | 0,899 | 0,569 | 0 |
| 6 | Hist1h2ap | 0 | 3,898 | 0,776 | 0,193 | 0 |
| 6 | Hist1h2ae | 0 | 3,169 | 0,688 | 0,106 | 0 |
| 6 | Mki67 | 0 | 3,158 | 0,942 | 0,225 | 0 |
| 6 | Stmn1 | 0 | 3,044 | 0,953 | 0,104 | 0 |
| 6 | Hist1h1b | 0 | 3,009 | 0,732 | 0,095 | 0 |
| 6 | Top2a | 0 | 2,869 | 0,906 | 0,239 | 0 |
| 6 | Rrm2 | 0 | 2,734 | 0,800 | 0,080 | 0 |
| 6 | Hist1h1e | 0 | 2,424 | 0,955 | 0,663 | 0 |
| 6 | Hist1h1a | 0 | 2,409 | 0,640 | 0,097 | 0 |
| 6 | Ccna2 | 0 | 2,297 | 0,726 | 0,019 | 0 |
| 6 | Ube2c | 0 | 2,153 | 0,636 | 0,051 | 0 |
| 6 | Tubb5 | 0 | 2,152 | 0,983 | 0,717 | 0 |
| 6 | Ccnb2 | 0 | 2,147 | 0,697 | 0,031 | 0 |
| 6 | Tuba1b | 0 | 2,038 | 0,928 | 0,432 | 0 |
| 6 | Hist1h1d | 0 | 2,032 | 0,669 | 0,127 | 0 |
| 6 | Pclaf | 0 | 2,019 | 0,832 | 0,079 | 0 |
| 6 | Ncapd2 | 0 | 2,003 | 0,925 | 0,243 | 0 |

| | | | | | | |
|---|---|---|---|---|---|---|
| 6 | Birc5 | 0 | 1,994 | 0,730 | 0,026 | 0 |
| 6 | Cdk1 | 0 | 1,930 | 0,786 | 0,061 | 0 |
| 6 | Kif11 | 0 | 1,880 | 0,735 | 0,075 | 0 |
| 6 | Uhrf1 | 0 | 1,854 | 0,879 | 0,159 | 0 |
| 6 | Rrm1 | 0 | 1,819 | 0,910 | 0,285 | 0 |
| 6 | Cdca3 | 0 | 1,800 | 0,709 | 0,033 | 0 |
| 6 | Nusap1 | 0 | 1,773 | 0,667 | 0,032 | 0 |
| 6 | Tacc3 | 0 | 1,770 | 0,826 | 0,092 | 0 |
| 6 | Mcm3 | 0 | 1,761 | 0,889 | 0,300 | 0 |
| 6 | Tpx2 | 0 | 1,745 | 0,744 | 0,037 | 0 |
| 6 | Mcm5 | 0 | 1,725 | 0,895 | 0,196 | 0 |
| 6 | Hist1h3c | 0 | 1,711 | 0,558 | 0,013 | 0 |
| 6 | Mcm7 | 0 | 1,685 | 0,885 | 0,259 | 0 |
| 7 | Ccr7 | 0 | 2,451 | 0,909 | 0,191 | 0 |
| 7 | Sell | 0 | 2,323 | 0,963 | 0,281 | 0 |
| 7 | Klf2 | 2,31E-274 | 1,989 | 0,962 | 0,508 | 4,30E-270 |
| 7 | S1pr1 | 0 | 1,986 | 0,897 | 0,359 | 0 |
| 7 | Il6st | 3,34E-267 | 1,977 | 0,882 | 0,491 | 6,20E-263 |
| 7 | Cmah | 0 | 1,903 | 0,850 | 0,219 | 0 |
| 7 | Vps37b | 1,44E-194 | 1,887 | 0,938 | 0,840 | 2,67E-190 |
| 7 | Lef1 | 1,87E-288 | 1,848 | 0,932 | 0,491 | 3,47E-284 |
| 7 | Txk | 2,15E-241 | 1,814 | 0,893 | 0,594 | 3,99E-237 |
| 7 | Ifngr2 | 0 | 1,794 | 0,473 | 0,069 | 0 |
| 7 | Il4ra | 3,93E-211 | 1,782 | 0,818 | 0,495 | 7,31E-207 |
| 7 | Dgka | 1,51E-303 | 1,772 | 0,976 | 0,849 | 2,81E-299 |
| 7 | Gramd3 | 1,24E-251 | 1,759 | 0,972 | 0,853 | 2,31E-247 |
| 7 | Socs3 | 1,24E-206 | 1,739 | 0,832 | 0,454 | 2,30E-202 |
| 7 | Dusp10 | 3,02E-103 | 1,727 | 0,687 | 0,476 | 5,62E-99 |
| 7 | Actn1 | 1,08E-231 | 1,684 | 0,801 | 0,356 | 2,01E-227 |
| 7 | Slc12a7 | 4,70E-207 | 1,651 | 0,862 | 0,557 | 8,74E-203 |
| 7 | Il7r | 5,92E-176 | 1,646 | 0,907 | 0,654 | 1,10E-171 |
| 7 | Bcl2 | 2,31E-160 | 1,624 | 0,833 | 0,552 | 4,28E-156 |
| 7 | Pik3ip1 | 4,10E-235 | 1,595 | 0,687 | 0,240 | 7,63E-231 |

| | | | | | | |
|---|---|---|---|---|---|---|
| 7 | Il6ra | 0 | 1,590 | 0,593 | 0,103 | 0 |
| 7 | Igfbp4 | 8,44E-256 | 1,563 | 0,523 | 0,106 | 1,57E-251 |
| 7 | Fam241a | 1,06E-163 | 1,508 | 0,822 | 0,551 | 1,97E-159 |
| 7 | Arhgef18 | 1,33E-216 | 1,506 | 0,942 | 0,757 | 2,47E-212 |
| 7 | Gramd4 | 2,01E-217 | 1,502 | 0,797 | 0,383 | 3,74E-213 |
| 7 | Rab3ip | 1,34E-160 | 1,491 | 0,705 | 0,375 | 2,50E-156 |
| 7 | Pde2a | 2,55E-166 | 1,487 | 0,652 | 0,260 | 4,74E-162 |
| 7 | Dph5 | 4,92E-201 | 1,466 | 0,688 | 0,284 | 9,14E-197 |
| 7 | Fam169b | 4,80E-166 | 1,429 | 0,806 | 0,525 | 8,92E-162 |
| 7 | mt-Cytb | 6,00E-169 | 1,412 | 0,939 | 0,844 | 1,11E-164 |
| 8 | Trdc | 0 | 3,753 | 0,591 | 0,062 | 0 |
| 8 | Klre1 | 0 | 2,211 | 0,834 | 0,221 | 0 |
| 8 | Cd7 | 5,67E-201 | 2,127 | 0,873 | 0,412 | 1,05E-196 |
| 8 | Tcrg-V7 | 3,03E-150 | 2,073 | 0,286 | 0,045 | 5,64E-146 |
| 8 | Spry2 | 5,54E-248 | 2,009 | 0,693 | 0,180 | 1,03E-243 |
| 8 | Tcrg-C4 | 0 | 2,008 | 0,725 | 0,088 | 0 |
| 8 | Fcer1g | 8,09E-200 | 1,932 | 0,334 | 0,047 | 1,50E-195 |
| 8 | Tcrg-C2 | 0 | 1,912 | 0,855 | 0,205 | 0 |
| 8 | Ikzf2 | 5,96E-286 | 1,679 | 0,815 | 0,230 | 1,11E-281 |
| 8 | Klrk1 | 1,81E-214 | 1,677 | 0,970 | 0,539 | 3,36E-210 |
| 8 | Tcrg-C1 | 0 | 1,675 | 0,585 | 0,057 | 0 |
| 8 | Cd244a | 0 | 1,630 | 0,662 | 0,122 | 0 |
| 8 | Lat2 | 3,37E-298 | 1,426 | 0,726 | 0,165 | 6,26E-294 |
| 8 | Gzma | 2,11E-10 | 1,369 | 0,540 | 0,512 | 3,92E-06 |
| 8 | Itgae | 2,32E-43 | 1,233 | 0,505 | 0,288 | 4,31E-39 |
| 8 | Itgax | 3,27E-81 | 1,136 | 0,730 | 0,414 | 6,07E-77 |
| 8 | Itga1 | 8,94E-108 | 1,065 | 0,801 | 0,418 | 1,66E-103 |
| 8 | Ccl5 | 3,44E-102 | 1,061 | 0,995 | 0,938 | 6,39E-98 |
| 8 | Itgb1 | 1,89E-58 | 1,043 | 0,859 | 0,669 | 3,52E-54 |
| 8 | Fgl2 | 4,17E-97 | 1,041 | 0,955 | 0,693 | 7,76E-93 |
| 8 | Chn2 | 5,53E-151 | 1,040 | 0,549 | 0,158 | 1,03E-146 |
| 8 | Klrd1 | 1,02E-122 | 1,020 | 0,969 | 0,707 | 1,90E-118 |
| 8 | Ier5l | 3,62E-42 | 1,003 | 0,462 | 0,241 | 6,73E-38 |

| | | | | | | |
|---|---|---|---|---|---|---|
| 8 | Rgs2 | 1,96E-38 | 0,951 | 0,587 | 0,371 | 3,64E-34 |
| 8 | Plcg2 | 2,83E-131 | 0,897 | 0,613 | 0,225 | 5,25E-127 |
| 8 | Tcrg-V1 | 2,92E-241 | 0,821 | 0,329 | 0,037 | 5,43E-237 |
| 8 | Lgals3 | 1,80E-45 | 0,811 | 0,669 | 0,410 | 3,35E-41 |
| 8 | Trat1 | 1,14E-96 | 0,805 | 0,510 | 0,185 | 2,12E-92 |
| 8 | Zfp36l2 | 1,78E-56 | 0,805 | 0,986 | 0,933 | 3,30E-52 |
| 8 | Tcrg-V4 | 3,87E-281 | 0,800 | 0,312 | 0,028 | 7,19E-277 |
| 9 | S100a8 | 4,04E-208 | 6,534 | 0,291 | 0,019 | 7,51E-204 |
| 9 | Cd74 | 2,28E-60 | 5,305 | 0,474 | 0,176 | 4,23E-56 |
| 9 | H2-Aa | 6,12E-237 | 4,781 | 0,443 | 0,042 | 1,14E-232 |
| 9 | Lyz2 | 1,07E-138 | 4,760 | 0,325 | 0,038 | 1,98E-134 |
| 9 | H2-Eb1 | 1,19E-183 | 4,619 | 0,429 | 0,051 | 2,20E-179 |
| 9 | H2-Ab1 | 7,98E-132 | 4,320 | 0,433 | 0,073 | 1,48E-127 |
| 9 | Tyrobp | 0 | 3,492 | 0,391 | 0,015 | 0 |
| 9 | Fth1 | 7,30E-10 | 3,318 | 0,637 | 0,922 | 1,36E-05 |
| 9 | Fcer1g | 1,98E-134 | 3,004 | 0,374 | 0,051 | 3,68E-130 |
| 9 | Fabp5 | 7,00E-12 | 2,710 | 0,284 | 0,168 | 1,30E-07 |
| 9 | Mpeg1 | 5,63E-250 | 2,710 | 0,315 | 0,019 | 1,05E-245 |
| 9 | Cebpb | 7,19E-06 | 2,653 | 0,401 | 0,397 | 0,133676174 |
| 9 | Ctsh | 0 | 2,528 | 0,332 | 0,005 | 0 |
| 9 | Tgfbi | 0 | 2,476 | 0,294 | 0,003 | 0 |
| 9 | Ctsz | 1,07E-07 | 2,472 | 0,367 | 0,321 | 0,001980474 |
| 9 | Msrb1 | 5,85E-06 | 2,454 | 0,388 | 0,383 | 0,108745103 |
| 9 | Alox5ap | 0 | 2,391 | 0,298 | 0,004 | 0 |
| 9 | Tnfaip2 | 0 | 2,346 | 0,284 | 0,005 | 0 |
| 9 | Cybb | 0 | 2,333 | 0,284 | 0,007 | 0 |
| 9 | Ifi30 | 7,20E-19 | 2,251 | 0,291 | 0,142 | 1,34E-14 |
| 9 | Lgals3 | 2,04E-08 | 2,246 | 0,450 | 0,417 | 0,000378371 |
| 9 | Spi1 | 0 | 2,236 | 0,322 | 0,003 | 0 |
| 9 | Ifitm3 | 2,51E-17 | 2,063 | 0,346 | 0,185 | 4,67E-13 |
| 9 | Tgm2 | 1,54E-191 | 2,030 | 0,270 | 0,018 | 2,86E-187 |
| 9 | Ifitm2 | 2,19E-26 | 1,956 | 0,304 | 0,118 | 4,07E-22 |
| 9 | Ncf2 | 4,64E-195 | 1,906 | 0,280 | 0,019 | 8,61E-191 |

| | | | | | | |
|---|---|---|---|---|---|---|
| 9 | Fcgr3 | 6,15E-291 | 1,898 | 0,273 | 0,011 | 1,14E-286 |
| 9 | Marcks | 0 | 1,782 | 0,260 | 0,003 | 0 |
| 9 | Plin2 | 1,72E-22 | 1,760 | 0,287 | 0,125 | 3,20E-18 |
| 9 | Psap | 0,009294362 | 1,699 | 0,433 | 0,792 | 1 |
| 10 | Il17a | 0 | 3,455 | 0,339 | 0,003 | 0 |
| 10 | Tmem176b | 0 | 3,420 | 0,983 | 0,048 | 0 |
| 10 | Tmem176a | 0 | 3,411 | 0,962 | 0,047 | 0 |
| 10 | Ramp1 | 0 | 3,160 | 0,962 | 0,106 | 0 |
| 10 | Rorc | 0 | 2,797 | 0,860 | 0,014 | 0 |
| 10 | Selenop | 3,51E-119 | 2,612 | 0,750 | 0,214 | 6,53E-115 |
| 10 | Zbtb16 | 0 | 2,429 | 0,682 | 0,007 | 0 |
| 10 | Il1r1 | 0 | 2,246 | 0,818 | 0,004 | 0 |
| 10 | Klrb1b | 0 | 2,246 | 0,407 | 0,017 | 0 |
| 10 | Maf | 6,94E-271 | 2,169 | 0,780 | 0,097 | 1,29E-266 |
| 10 | Cpd | 2,44E-232 | 2,101 | 0,894 | 0,170 | 4,53E-228 |
| 10 | Capg | 1,97E-124 | 2,065 | 0,936 | 0,396 | 3,66E-120 |
| 10 | Ly6g5b | 9,47E-194 | 2,036 | 0,818 | 0,158 | 1,76E-189 |
| 10 | Smox | 2,43E-145 | 1,870 | 0,775 | 0,183 | 4,52E-141 |
| 10 | Ltb4r1 | 0 | 1,851 | 0,771 | 0,037 | 0 |
| 10 | Il1r2 | 2,44E-60 | 1,754 | 0,318 | 0,061 | 4,53E-56 |
| 10 | Mmp25 | 0 | 1,732 | 0,610 | 0,006 | 0 |
| 10 | Tff1 | 8,08E-40 | 1,690 | 0,335 | 0,094 | 1,50E-35 |
| 10 | Furin | 1,38E-73 | 1,688 | 0,975 | 0,707 | 2,57E-69 |
| 10 | S100a4 | 4,59E-76 | 1,670 | 0,979 | 0,594 | 8,52E-72 |
| 10 | Irs2 | 1,47E-81 | 1,611 | 0,581 | 0,154 | 2,73E-77 |
| 10 | Pdcd1 | 7,08E-59 | 1,523 | 0,839 | 0,412 | 1,31E-54 |
| 10 | Serpinb1a | 3,41E-104 | 1,514 | 0,614 | 0,139 | 6,33E-100 |
| 10 | Tcrg-C1 | 1,93E-290 | 1,432 | 0,682 | 0,065 | 3,58E-286 |
| 10 | Plxnd1 | 0 | 1,414 | 0,559 | 0,033 | 0 |
| 10 | Blk | 0 | 1,404 | 0,352 | 0,001 | 0 |
| 10 | Sdc4 | 0 | 1,370 | 0,492 | 0,021 | 0 |
| 10 | Podnl1 | 5,53E-155 | 1,370 | 0,653 | 0,119 | 1,03E-150 |
| 10 | Slc27a6 | 0 | 1,370 | 0,669 | 0,005 | 0 |

| | | | | | | |
|---|---|---|---|---|---|---|
| 10 | Olfr60 | 4,30E-169 | 1,369 | 0,775 | 0,144 | 7,99E-165 |

### Table S3

| HALLMARK_INTERFERON_ALPHA_RESPONSE | Tumor-specific [Oliveira G et al,Nature. 2021] | Exhausted [Im SJ et al, Nature. 2016] | Effector-like [Miller BC et al, Nat Immunol. 2019] | Terminally Exh [Miller BC et al, Nat Immunol. 2019] |
|---|---|---|---|---|
| Trim25 | Krt86 | Pdcd1 | 1810058I24Rik | 2900026A02Rik |
| Il15 | Rdh10 | Tox | Abracl | Abi3 |
| Stat2 | Tyms | Il21r | Acp5 | Adam19 |
| Procr | Hmox1 | Sh2d2a | Actb | Akna |
| Wars1 | Gng4 | Cd8a | Adgre5 | Apobec3 |
| Psma3 | Cxcl13 | Nkg7 | Ahnak | Arhgap9 |
| Il4ra | Afap1l2 | Fgl2 | AI467606 | Arl6ip1 |
| Ly6e | Acp5 | Entpd1 | Anxa2 | Armc7 |
| Trim21 | Myo1e | Fasl | Anxa6 | AW112010 |
| Elf1 | Layn | Ccl5 | Apbb1ip | Bcl2a1b |
| Irf9 | Tns3 | Apol7e | Arhgdib | Cbx4 |
| Psme2 | Tnfsf4 | Arhgap9 | Arl4c | Ccl3 |
| Psme1 | Akap5 | Nedd9 | Arl6ip5 | Ccl4 |
| Ifit3 | Havcr2 | Cd38 | Arpc2 | Ccl5 |
| Casp8 | Entpd1 | Pvrig | Arpc5 | Cd160 |
| Ifi27 | Slc2a8 | Trbc2 | Atp1b3 | Cd164 |
| Trim26 | Mcm5 | Chn2 | Atp2b1 | Cd27 |
| Csf1 | Cav1 | 2900026A02Rik | Atp5h | Cd3e |
| Samd9l | Golim4 | Pstpip1 | Bhlhe40 | Cd3g |
| Usp18 | Vcam1 | Chst12 | Bin2 | Cd7 |
| Psmb9 | Pon2 | Cd3e | Calm1 | Cd82 |
| Psmb8 | Mtss1 | Cxcr6 | Capn2 | Cd8a |
| Uba7 | Cd38 | Cd3g | Ccl5 | Cst7 |
| Cxcl10 | Tox2 | Adgrg1 | Ccnd3 | Cxcr6 |
| Eif2ak2 | Csf1 | Prkch | Cd247 | Dapk2 |
| C1s1 | Galnt2 | Tap2 | Cd48 | Dtx1 |
| Isg15 | Fxyd2 | Irf1 | Cd8b1 | Dusp2 |
| Irf7 | Plpp1 | Tapbpl | Cdc42 | Efhd2 |
| Cxcl11 | Lmcd1 | AW112010 | Cdkn2d | Eif4a2 |
| Nub1 | Myl6b | Arl6ip1 | Cmpk1 | Fam189b |
| Txnip | Lag3 | Gng2 | Crip1 | Fasl |
| Adar | Igflr1 | Adam19 | Ctsd | Foxn3 |
| Ripk2 | Ccdc50 | Gimap7 | Cx3cr1 | Fyn |
| Ifitm3 | Cd27 | Gzmk | Cyba | Gimap1 |
| Gmpr | Cdkn2a | | Cyth4 | Gimap6 |
| Lap3 | Cd70 | | Dnajc15 | Gimap7 |
| Herc6 | Abhd6 | | Dusp5 | Glrx |
| Lpar6 | Ctla4 | | Emp3 | Gm12216 |
| Ube2l6 | Pdcd1 | | Fam107b | Gng2 |
| Rtp4 | Gem | | Fam65b | Gramd1a |
| Epsti1 | Nusap1 | | Flna | Gzma |
| Tmem140 | Tox | | Fxyd5 | Gzmb |
| Ifitm1 | Cxcr6 | | Glipr2 | Gzmk |
| Bst2 | Nmb | | Glud1 | H2-Q4 |
| Ifi35 | Hopx | | Gmfg | Hcst |
| Ifih1 | Clic3 | | Gna15 | Hspa5 |
| Pnpt1 | Inpp5f | | H2-D1 | Id2 |

| | | | | |
|---|---|---|---|---|
| Ogfr | Snap47 | | H2-Q7 | Ifi47 |
| Parp14 | Tshz2 | | H2afz | Ifit3 |
| Ccrl2 | Sit1 | | H3f3a | Il21r |
| Mvb12a | Pycard | | Hsd11b1 | Isg15 |
| Cmtr1 | Adgrg1 | | Ier2 | Itk |
| Batf2 | Prf1 | | Ier5 | Itpkb |
| Trim14 | Ptms | | Ifngr1 | Lag3 |
| Sp110 | Cks1b | | Il17ra | Lax1 |
| Trafd1 | Hipk2 | | Il18r1 | Lrrk1 |
| Gbp3 | Chst12 | | Itgb2 | Mbnl1 |
| Nmi | Lsp1 | | Itgb7 | Mxd4 |
| Isg20 | Fam3c | | Kcnab2 | Nr4a2 |
| Rsad2 | Slc1a4 | | Kcnn4 | Pdcd1 |
| Dhx58 | Nudt1 | | Klf2 | Pfdn5 |
| Parp9 | Dnph1 | | Klf3 | Pla2g16 |
| Ifitm2 | | | Klrc1 | Plac8 |
| Rnf31 | | | Klrd1 | Prdx5 |
| Ifi30 | | | Klrk1 | Prkch |
| Tdrd7 | | | Krtcap2 | Psmb10 |
| Parp12 | | | Lamb3 | Psmb8 |
| Slc25a28 | | | Laptm5 | Psme1 |
| Oasl1 | | | Lfng | Ptger4 |
| Oas1a | | | Lgals1 | Ptpn18 |
| Ddx60 | | | Lgals3 | Ptpn22 |
| Helz2 | | | Lrrc58 | Rgs1 |
| Lamp3 | | | Lrrfip1 | Rgs2 |
| Ifi44 | | | Lsp1 | Rgs3 |
| Ncoa7 | | | Ly6c2 | Rtp4 |
| Tent5a | | | Ms4a4b | Runx3 |
| Trim12c | | | Myl6 | Serpina3g |
| B2m | | | Myo1f | Serpinb6b |
| Cnp | | | Myo1g | Sh2d2a |
| Plscr1 | | | Neat1 | Shisa5 |
| Ifi44l | | | Nfkbia | Sipa1 |
| Cd74 | | | Nkg7 | Slc3a2 |
| Casp1 | | | Nptn | Stat1 |
| Il7 | | | Nt5c | Stk17b |
| Irf1 | | | Ostf1 | Tap2 |
| Irf2 | | | Pim1 | Tapbp |
| Cd47 | | | Ppp1r12a | Tapbpl |
| Mov10 | | | Prex1 | Tnfrsf1b |
| Mx2 | | | Prkcq | Tox |
| Sell | | | Prr13 | Ucp2 |
| Tap1 | | | Psap | Vmp1 |
| Ifit2 | | | Pycard | Zbp1 |
| Lgals3bp | | | Racgap1 | |
| Cmpk2 | | | Rap1b | |
| | | | Rasa3 | |

| | | | |
|---|---|---|---|
| | | | Rasgrp2<br>Reep5<br>Rhoa<br>Rora<br>S100a10<br>S100a11<br>S100a13<br>S100a4<br>S100a6<br>S1pr4<br>Selplg<br>Sept11<br>Sgk1<br>Sh3bgrl3<br>Snx5<br>Spn<br>Stk24<br>Tagln2<br>Thy1<br>Tm6sf1<br>Tmem50a<br>Tnfaip8l2<br>Tprgl<br>Tsc22d3<br>Tspo<br>Tuba1a<br>Txk<br>Txndc5<br>Vim<br>Xrn2<br>Ywhaq<br>Zeb2<br>Zyx |

### Table S4

| gene | p_val | avg_log2FC | pct.1 | pct.2 | p_val_adj |
|---|---|---|---|---|---|
| Gzmb | 0 | 3,03 | 0,938 | 0,398 | 0 |
| Ccl3 | 0 | 2,79 | 0,573 | 0,193 | 0 |
| Ifit3 | 0 | 2,67 | 0,751 | 0,203 | 0 |
| Isg15 | 0 | 2,41 | 0,776 | 0,265 | 0 |
| Ifi27l2a | 0 | 2,08 | 0,855 | 0,534 | 0 |
| Gzmf | 3,15E-206 | 1,91 | 0,109 | 0,002 | 5,85E-202 |
| Ifit1 | 0 | 1,84 | 0,606 | 0,123 | 0 |
| AA467197 | 0 | 1,81 | 0,648 | 0,263 | 0 |
| Usp18 | 0 | 1,79 | 0,747 | 0,264 | 0 |
| Ifit1bl1 | 0 | 1,67 | 0,801 | 0,527 | 0 |
| Ccl4 | 0 | 1,66 | 0,847 | 0,582 | 0 |
| Gzma | 0 | 1,59 | 0,714 | 0,263 | 0 |
| Isg20 | 0 | 1,54 | 0,807 | 0,496 | 0 |
| Tnfrsf9 | 0 | 1,54 | 0,489 | 0,175 | 0 |
| Bst2 | 0 | 1,53 | 0,896 | 0,682 | 0 |
| Rsad2 | 0 | 1,52 | 0,452 | 0,044 | 0 |
| Ifng | 0 | 1,49 | 0,816 | 0,563 | 0 |
| Ifitm3 | 0 | 1,47 | 0,319 | 0,024 | 0 |
| Lag3 | 0 | 1,45 | 0,765 | 0,458 | 0 |
| Il2ra | 0 | 1,44 | 0,532 | 0,242 | 0 |
| Plac8 | 0 | 1,42 | 0,916 | 0,728 | 0 |
| Ifitm1 | 1,82E-241 | 1,41 | 0,17 | 0,021 | 3,39E-237 |
| Rnf213 | 0 | 1,41 | 0,918 | 0,81 | 0 |
| Prf1 | 1,43E-292 | 1,37 | 0,82 | 0,679 | 2,66E-288 |
| Cxcl10 | 2,67E-196 | 1,35 | 0,215 | 0,062 | 4,96E-192 |
| Ifit2 | 0 | 1,33 | 0,587 | 0,246 | 0 |
| Oas3 | 0 | 1,32 | 0,711 | 0,362 | 0 |
| Pik3ap1 | 0 | 1,32 | 0,814 | 0,467 | 0 |
| Rtp4 | 0 | 1,32 | 0,784 | 0,386 | 0 |
| Phf11b | 0 | 1,31 | 0,841 | 0,555 | 0 |
| Ifi211 | 0 | 1,30 | 0,481 | 0,074 | 0 |
| Irf7 | 0 | 1,26 | 0,827 | 0,527 | 0 |
| Dhx58 | 0 | 1,24 | 0,67 | 0,245 | 0 |
| Daxx | 0 | 1,22 | 0,774 | 0,524 | 0 |
| Ifih1 | 0 | 1,22 | 0,593 | 0,2 | 0 |
| Ifi208 | 0 | 1,21 | 0,645 | 0,23 | 0 |
| Havcr2 | 0 | 1,20 | 0,501 | 0,195 | 0 |
| Slfn5 | 0 | 1,19 | 0,332 | 0,068 | 0 |
| Zbp1 | 0 | 1,19 | 0,932 | 0,791 | 0 |
| Ifitm2 | 8,02E-259 | 1,17 | 0,194 | 0,031 | 1,49E-254 |
| Irf8 | 0 | 1,15 | 0,537 | 0,235 | 0 |
| Oas1a | 0 | 1,12 | 0,559 | 0,176 | 0 |
| Xaf1 | 0 | 1,10 | 0,821 | 0,503 | 0 |
| Tnfrsf4 | 0 | 1,08 | 0,31 | 0,055 | 0 |
| Icos | 0 | 1,08 | 0,864 | 0,708 | 0 |

Up in SP140KO
Up in WT

| | | | | | |
|---|---|---|---|---|---|
| Gzmc | 1,30E-187 | 1,06 | 0,132 | 0,016 | 2,41E-183 |
| Ly6a | 0 | 1,06 | 0,98 | 0,867 | 0 |
| Id2 | 0 | 1,06 | 0,954 | 0,822 | 0 |
| Trafd1 | 0 | 1,05 | 0,804 | 0,562 | 0 |
| Slfn1 | 0 | 1,05 | 0,852 | 0,611 | 0 |
| Lgals3bp | 0 | 1,03 | 0,942 | 0,848 | 0 |
| Sema7a | 0 | 1,02 | 0,441 | 0,096 | 0 |
| Cmpk2 | 0 | 1,01 | 0,419 | 0,09 | 0 |
| Ifi209 | 0 | 1,01 | 0,893 | 0,765 | 0 |
| Hif1a | 0 | 1,01 | 0,785 | 0,585 | 0 |
| Tnfrsf18 | 0 | 0,99 | 0,709 | 0,504 | 0 |
| Mxd1 | 1,70E-294 | 0,98 | 0,761 | 0,575 | 3,16E-290 |
| Cdkn1a | 0 | 0,97 | 0,343 | 0,089 | 0 |
| Sytl3 | 0 | 0,94 | 0,721 | 0,397 | 0 |
| Phf11c | 0 | 0,93 | 0,697 | 0,385 | 0 |
| Pkm | 0 | 0,93 | 0,874 | 0,738 | 0 |
| Sdc3 | 0 | 0,90 | 0,35 | 0,042 | 0 |
| Nt5c3 | 5,47E-299 | 0,90 | 0,652 | 0,425 | 1,02E-294 |
| Rilpl2 | 0 | 0,89 | 0,626 | 0,357 | 0 |
| Rbm3 | 0 | 0,88 | 0,87 | 0,677 | 0 |
| Ms4a4c | 2,16E-295 | 0,88 | 0,553 | 0,301 | 4,01E-291 |
| Ifit3b | 0 | 0,88 | 0,367 | 0,032 | 0 |
| Herc6 | 0 | 0,88 | 0,757 | 0,53 | 0 |
| Il18rap | 0 | 0,87 | 0,883 | 0,664 | 0 |
| Pim1 | 0 | 0,87 | 0,866 | 0,652 | 0 |
| Casp3 | 0 | 0,85 | 0,731 | 0,456 | 0 |
| Entpd1 | 0 | 0,85 | 0,602 | 0,273 | 0 |
| Slfn8 | 0 | 0,84 | 0,794 | 0,517 | 0 |
| Helz2 | 0 | 0,84 | 0,878 | 0,753 | 0 |
| Acadl | 7,08E-265 | 0,84 | 0,746 | 0,589 | 1,32E-260 |
| Pml | 0 | 0,84 | 0,768 | 0,512 | 0 |
| Slc7a5 | 1,31E-214 | 0,83 | 0,462 | 0,253 | 2,43E-210 |
| Phf11a | 0 | 0,82 | 0,508 | 0,141 | 0 |
| Casp4 | 0 | 0,82 | 0,678 | 0,36 | 0 |
| Srgn | 0 | 0,80 | 0,98 | 0,955 | 0 |
| Sv2c | 0 | 0,80 | 0,297 | 0,039 | 0 |
| Pdcd1 | 0 | 0,80 | 0,544 | 0,26 | 0 |
| Cd86 | 0 | 0,79 | 0,543 | 0,267 | 0 |
| Parp9 | 0 | 0,77 | 0,813 | 0,571 | 0 |
| Mapkapk2 | 0 | 0,77 | 0,838 | 0,674 | 0 |
| Glrx | 1,59E-216 | 0,76 | 0,724 | 0,536 | 2,95E-212 |
| Tpi1 | 3,81E-292 | 0,76 | 0,445 | 0,209 | 7,07E-288 |
| Ccrl2 | 0 | 0,76 | 0,322 | 0,048 | 0 |
| Gstt1 | 2,31E-209 | 0,75 | 0,265 | 0,092 | 4,29E-205 |
| Oasl2 | 0 | 0,74 | 0,288 | 0,067 | 0 |
| Gadd45b | 6,41E-156 | 0,74 | 0,558 | 0,377 | 1,19E-151 |

| | | | | | |
|---|---|---|---|---|---|
| Il10ra | 0 | 0,72 | 0,707 | 0,421 | 0 |
| Hk2 | 0 | 0,72 | 0,338 | 0,09 | 0 |
| Eif2ak2 | 0 | 0,72 | 0,596 | 0,282 | 0 |
| Serpinb9 | 4,54E-238 | 0,71 | 0,638 | 0,419 | 8,43E-234 |
| Stx11 | 0 | 0,71 | 0,705 | 0,421 | 0 |
| Plek | 0 | 0,71 | 0,777 | 0,518 | 0 |
| Nr4a3 | 2,60E-93 | 0,70 | 0,278 | 0,154 | 4,82E-89 |
| Cish | 0 | 0,69 | 0,744 | 0,453 | 0 |
| Trim30d | 0 | 0,68 | 0,544 | 0,229 | 0 |
| Trim30a | 0 | 0,68 | 0,736 | 0,511 | 0 |
| Clic4 | 0 | 0,68 | 0,629 | 0,35 | 0 |
| Dtx3l | 0 | 0,68 | 0,89 | 0,784 | 0 |
| Ube2l6 | 0 | 0,67 | 0,344 | 0,083 | 0 |
| Batf | 1,47E-265 | 0,67 | 0,712 | 0,506 | 2,73E-261 |
| Hlx | 6,63E-289 | 0,66 | 0,302 | 0,088 | 1,23E-284 |
| Naa20 | 0 | 0,66 | 0,743 | 0,52 | 0 |
| Il12rb1 | 5,58E-305 | 0,65 | 0,746 | 0,517 | 1,04E-300 |
| Aldoa | 4,86E-248 | 0,65 | 0,841 | 0,729 | 9,03E-244 |
| Wdfy1 | 0 | 0,65 | 0,605 | 0,281 | 0 |
| Zbtb32 | 0 | 0,64 | 0,278 | 0,026 | 0 |
| Rgs16 | 1,31E-191 | 0,64 | 0,475 | 0,27 | 2,44E-187 |
| Gbp7 | 0 | 0,64 | 0,855 | 0,689 | 0 |
| Parp12 | 8,02E-296 | 0,64 | 0,462 | 0,213 | 1,49E-291 |
| Lamc1 | 0 | 0,64 | 0,62 | 0,307 | 0 |
| Arsb | 0 | 0,64 | 0,754 | 0,522 | 0 |
| Mif4gd | 1,09E-303 | 0,64 | 0,775 | 0,58 | 2,03E-299 |
| Setbp1 | 0 | 0,63 | 0,536 | 0,262 | 0 |
| Parp14 | 1,10E-216 | 0,63 | 0,854 | 0,728 | 2,05E-212 |
| Chmp4b | 0 | 0,63 | 0,84 | 0,641 | 0 |
| Prdm1 | 2,60E-293 | 0,62 | 0,546 | 0,285 | 4,83E-289 |
| Ifi206 | 5,05E-239 | 0,62 | 0,786 | 0,632 | 9,39E-235 |
| Marcksl1 | 1,22E-195 | 0,62 | 0,262 | 0,093 | 2,27E-191 |
| Rab27a | 0 | 0,62 | 0,828 | 0,628 | 0 |
| Il12rb2 | 2,46E-260 | 0,61 | 0,665 | 0,415 | 4,58E-256 |
| Serpina3g | 2,09E-208 | 0,61 | 0,775 | 0,582 | 3,88E-204 |
| Il1r2 | 1,64E-192 | 0,61 | 0,112 | 0,005 | 3,05E-188 |
| Serpina3f | 5,35E-299 | 0,61 | 0,335 | 0,106 | 9,94E-295 |
| Ddx58 | 8,21E-236 | 0,60 | 0,731 | 0,548 | 1,53E-231 |
| Ifi35 | 0 | 0,60 | 0,779 | 0,571 | 0 |
| Gbp2 | 8,61E-133 | 0,60 | 0,85 | 0,738 | 1,60E-128 |
| Hist1h2ap | 2,26E-16 | 0,60 | 0,264 | 0,209 | 4,20E-12 |
| Gem | 2,34E-137 | 0,60 | 0,556 | 0,388 | 4,34E-133 |
| Gbp9 | 3,68E-202 | 0,60 | 0,802 | 0,638 | 6,84E-198 |
| Nfkbia | 1,42E-171 | 0,60 | 0,855 | 0,731 | 2,63E-167 |
| Bcl2a1b | 0 | 0,59 | 0,772 | 0,523 | 0 |
| Sub1 | 9,79E-269 | 0,59 | 0,877 | 0,785 | 1,82E-264 |

| | | | | | |
|---|---|---|---|---|---|
| Lgals9 | 3,78E-172 | 0,59 | 0,766 | 0,642 | 7,02E-168 |
| Ldha | 2,22E-260 | 0,59 | 0,916 | 0,84 | 4,13E-256 |
| Ddx60 | 2,83E-274 | 0,59 | 0,45 | 0,209 | 5,26E-270 |
| Ifi214 | 1,02E-290 | 0,58 | 0,628 | 0,374 | 1,90E-286 |
| Txn1 | 3,05E-300 | 0,58 | 0,826 | 0,663 | 5,67E-296 |
| Dgat1 | 7,99E-225 | 0,58 | 0,66 | 0,453 | 1,49E-220 |
| Bcl2l1 | 1,25E-278 | 0,58 | 0,517 | 0,264 | 2,32E-274 |
| Fkbp5 | 1,31E-286 | 0,58 | 0,553 | 0,296 | 2,43E-282 |
| Ddt | 6,81E-171 | 0,57 | 0,556 | 0,364 | 1,26E-166 |
| Ptms | 4,40E-267 | 0,57 | 0,799 | 0,581 | 8,17E-263 |
| Coro2a | 7,69E-250 | 0,57 | 0,77 | 0,555 | 1,43E-245 |
| Map2k3 | 1,70E-206 | 0,57 | 0,658 | 0,463 | 3,16E-202 |
| Stat2 | 8,90E-253 | 0,56 | 0,597 | 0,36 | 1,65E-248 |
| Trim25 | 9,79E-296 | 0,56 | 0,731 | 0,488 | 1,82E-291 |
| Sdcbp2 | 4,16E-288 | 0,56 | 0,407 | 0,164 | 7,73E-284 |
| Ccr2 | 1,39E-228 | 0,56 | 0,534 | 0,286 | 2,57E-224 |
| Chchd10 | 3,50E-148 | 0,56 | 0,356 | 0,191 | 6,51E-144 |
| Prelid1 | 0 | 0,56 | 0,84 | 0,696 | 0 |
| Ifi204 | 0 | 0,56 | 0,17 | 0,005 | 0 |
| Gzmk | 1,57E-223 | 0,55 | 0,766 | 0,527 | 2,91E-219 |
| Tmem184b | 7,62E-224 | 0,55 | 0,606 | 0,385 | 1,42E-219 |
| Gpr65 | 3,81E-212 | 0,55 | 0,692 | 0,509 | 7,08E-208 |
| Gbp3 | 5,06E-256 | 0,55 | 0,732 | 0,509 | 9,41E-252 |
| Vps54 | 4,63E-293 | 0,55 | 0,927 | 0,854 | 8,60E-289 |
| Anxa2 | 1,25E-146 | 0,54 | 0,767 | 0,619 | 2,33E-142 |
| Nfkbid | 1,03E-103 | 0,54 | 0,451 | 0,296 | 1,91E-99 |
| Oas2 | 4,80E-263 | 0,54 | 0,247 | 0,059 | 8,92E-259 |
| H2-T24 | 2,29E-237 | 0,53 | 0,575 | 0,327 | 4,25E-233 |
| Atf6 | 2,80E-255 | 0,53 | 0,68 | 0,449 | 5,21E-251 |
| Tor3a | 4,32E-254 | 0,53 | 0,403 | 0,178 | 8,02E-250 |
| Maf | 6,57E-146 | 0,53 | 0,157 | 0,041 | 1,22E-141 |
| Egr1 | 6,03E-85 | 0,52 | 0,412 | 0,273 | 1,12E-80 |
| Tor1aip1 | 5,63E-262 | 0,52 | 0,907 | 0,822 | 1,05E-257 |
| Usp25 | 1,05E-247 | 0,52 | 0,91 | 0,831 | 1,95E-243 |
| Bhlhe40 | 1,67E-234 | 0,51 | 0,888 | 0,704 | 3,11E-230 |
| Slc16a3 | 8,37E-237 | 0,51 | 0,287 | 0,096 | 1,56E-232 |
| Sh3bp2 | 0 | 0,51 | 0,515 | 0,239 | 0 |
| Alcam | 1,51E-289 | 0,50 | 0,272 | 0,067 | 2,80E-285 |
| Rhbdf2 | 1,02E-261 | 0,50 | 0,527 | 0,288 | 1,90E-257 |
| Smpdl3b | 0 | 0,50 | 0,452 | 0,181 | 0 |
| Ifi203 | 6,28E-92 | 0,50 | 0,821 | 0,774 | 1,17E-87 |
| Ms4a4b | 0 | 0,50 | 0,98 | 0,977 | 0 |
| Tg | 0 | 0,50 | 0,212 | 0,026 | 0 |
| Samsn1 | 4,72E-156 | 0,50 | 0,623 | 0,445 | 8,77E-152 |
| Slfn2 | 2,22E-223 | 0,50 | 0,958 | 0,899 | 4,12E-219 |
| Ide | 8,47E-272 | 0,50 | 0,764 | 0,578 | 1,57E-267 |

| | | | | | |
|---|---|---|---|---|---|
| Mdfic | 3,95E-198 | 0,50 | 0,573 | 0,366 | 7,34E-194 |
| Slc25a19 | 1,81E-179 | 0,49 | 0,584 | 0,395 | 3,36E-175 |
| B4galt5 | 2,11E-176 | 0,49 | 0,782 | 0,632 | 3,93E-172 |
| Ctla2a | 1,21E-106 | 0,49 | 0,856 | 0,74 | 2,25E-102 |
| Gnptab | 3,39E-219 | 0,49 | 0,702 | 0,511 | 6,30E-215 |
| Tnfsf14 | 2,55E-62 | 0,48 | 0,256 | 0,16 | 4,73E-58 |
| N4bp1 | 1,57E-200 | 0,48 | 0,763 | 0,585 | 2,92E-196 |
| Grina | 1,88E-198 | 0,48 | 0,88 | 0,769 | 3,49E-194 |
| Sco1 | 8,05E-301 | 0,48 | 0,365 | 0,128 | 1,50E-296 |
| Klrc1 | 8,35E-190 | 0,48 | 0,799 | 0,568 | 1,55E-185 |
| Mt1 | 3,67E-211 | 0,48 | 0,22 | 0,058 | 6,83E-207 |
| Samhd1 | 4,86E-44 | 0,48 | 0,94 | 0,93 | 9,03E-40 |
| Tspo | 1,36E-213 | 0,48 | 0,869 | 0,746 | 2,53E-209 |
| Ccnyl1 | 3,93E-232 | 0,47 | 0,446 | 0,215 | 7,31E-228 |
| Hsh2d | 2,46E-291 | 0,47 | 0,456 | 0,199 | 4,58E-287 |
| Cysltr2 | 1,96E-184 | 0,47 | 0,692 | 0,488 | 3,64E-180 |
| Oasl1 | 1,62E-298 | 0,46 | 0,209 | 0,028 | 3,02E-294 |
| Dusp4 | 1,39E-185 | 0,46 | 0,305 | 0,126 | 2,58E-181 |
| Egr3 | 3,20E-82 | 0,45 | 0,16 | 0,068 | 5,95E-78 |
| B2m | 0 | 0,45 | 0,984 | 0,973 | 0 |
| Chd7 | 3,66E-151 | 0,45 | 0,786 | 0,658 | 6,81E-147 |
| Ddit4 | 8,86E-162 | 0,45 | 0,623 | 0,428 | 1,65E-157 |
| Ogfr | 1,41E-226 | 0,44 | 0,853 | 0,708 | 2,62E-222 |
| Serpinb6b | 8,09E-229 | 0,44 | 0,377 | 0,162 | 1,50E-224 |
| Tnf | 3,87E-50 | 0,44 | 0,29 | 0,192 | 7,18E-46 |
| Atp10a | 5,59E-186 | 0,44 | 0,652 | 0,445 | 1,04E-181 |
| Slc39a10 | 3,39E-127 | 0,44 | 0,462 | 0,299 | 6,29E-123 |
| Arpc4 | 6,20E-255 | 0,44 | 0,741 | 0,542 | 1,15E-250 |
| Ms4a6d | 3,39E-260 | 0,44 | 0,379 | 0,15 | 6,30E-256 |
| Bak1 | 9,51E-232 | 0,43 | 0,754 | 0,555 | 1,77E-227 |
| Mapkapk3 | 6,62E-196 | 0,43 | 0,744 | 0,545 | 1,23E-191 |
| Myd88 | 9,66E-232 | 0,43 | 0,679 | 0,457 | 1,80E-227 |
| Gbp6 | 1,18E-164 | 0,43 | 0,753 | 0,575 | 2,20E-160 |
| Hist1h2ae | 7,32E-31 | 0,43 | 0,18 | 0,118 | 1,36E-26 |
| Nkg7 | 2,67E-213 | 0,43 | 0,982 | 0,959 | 4,97E-209 |
| Ifi47 | 1,84E-130 | 0,43 | 0,917 | 0,855 | 3,42E-126 |
| Glipr2 | 7,22E-187 | 0,43 | 0,846 | 0,714 | 1,34E-182 |
| Gpr171 | 8,75E-159 | 0,43 | 0,897 | 0,811 | 1,63E-154 |
| Esm1 | 1,20E-183 | 0,43 | 0,23 | 0,073 | 2,23E-179 |
| Nampt | 3,16E-192 | 0,42 | 0,506 | 0,29 | 5,88E-188 |
| Dgkh | 3,74E-284 | 0,42 | 0,336 | 0,109 | 6,96E-280 |
| Ly6e | 3,16E-225 | 0,42 | 0,988 | 0,971 | 5,88E-221 |
| Frmd4b | 1,15E-204 | 0,42 | 0,435 | 0,223 | 2,13E-200 |
| Evi2a | 8,98E-176 | 0,42 | 0,501 | 0,294 | 1,67E-171 |
| S100a6 | 1,22E-101 | 0,41 | 0,695 | 0,524 | 2,26E-97 |
| Tmem163 | 1,05E-224 | 0,41 | 0,485 | 0,255 | 1,95E-220 |

| | | | | | |
|---|---|---|---|---|---|
| Irgm1 | 1,91E-140 | 0,41 | 0,813 | 0,671 | 3,55E-136 |
| Adar | 4,53E-187 | 0,41 | 0,838 | 0,699 | 8,42E-183 |
| Capza2 | 7,08E-223 | 0,41 | 0,716 | 0,534 | 1,32E-218 |
| Jaml | 5,56E-120 | 0,41 | 0,843 | 0,708 | 1,03E-115 |
| Farp1 | 2,61E-191 | 0,41 | 0,425 | 0,219 | 4,85E-187 |
| Smchd1 | 5,45E-84 | 0,41 | 0,834 | 0,749 | 1,01E-79 |
| Epsti1 | 1,50E-134 | 0,41 | 0,949 | 0,905 | 2,79E-130 |
| Lilrb4a | 6,44E-198 | 0,41 | 0,747 | 0,516 | 1,20E-193 |
| Rnf114 | 3,13E-171 | 0,41 | 0,873 | 0,769 | 5,83E-167 |
| Pfkp | 4,79E-134 | 0,40 | 0,848 | 0,768 | 8,89E-130 |
| Ctla4 | 5,63E-96 | 0,40 | 0,828 | 0,725 | 1,05E-91 |
| Lgals1 | 2,40E-155 | 0,40 | 0,69 | 0,503 | 4,45E-151 |
| Nek6 | 2,78E-129 | 0,40 | 0,419 | 0,249 | 5,17E-125 |
| Plp2 | 2,92E-166 | 0,40 | 0,637 | 0,441 | 5,42E-162 |
| Lgals3 | 9,37E-63 | 0,40 | 0,473 | 0,349 | 1,74E-58 |
| Nmi | 2,36E-152 | 0,40 | 0,781 | 0,623 | 4,39E-148 |
| Nr4a1 | 1,34E-19 | 0,40 | 0,466 | 0,398 | 2,49E-15 |
| Furin | 9,18E-62 | 0,39 | 0,759 | 0,651 | 1,71E-57 |
| Cd47 | 4,16E-258 | 0,39 | 0,971 | 0,957 | 7,73E-254 |
| Psma5 | 4,09E-178 | 0,39 | 0,741 | 0,567 | 7,60E-174 |
| Spats2 | 3,62E-303 | 0,39 | 0,233 | 0,039 | 6,74E-299 |
| Arf5 | 2,50E-215 | 0,39 | 0,902 | 0,815 | 4,64E-211 |
| Irf4 | 1,52E-95 | 0,39 | 0,427 | 0,28 | 2,82E-91 |
| Hilpda | 7,33E-132 | 0,39 | 0,292 | 0,142 | 1,36E-127 |
| Arl14ep | 4,95E-123 | 0,39 | 0,361 | 0,207 | 9,20E-119 |
| Ms4a6b | 1,56E-187 | 0,39 | 0,974 | 0,959 | 2,90E-183 |
| Lmnb1 | 1,10E-127 | 0,39 | 0,733 | 0,573 | 2,05E-123 |
| Gadd45g | 2,63E-123 | 0,39 | 0,325 | 0,168 | 4,89E-119 |
| Ppa1 | 2,81E-176 | 0,39 | 0,481 | 0,275 | 5,23E-172 |
| Hcls1 | 6,60E-238 | 0,39 | 0,937 | 0,88 | 1,23E-233 |
| Smim3 | 2,73E-238 | 0,38 | 0,38 | 0,161 | 5,07E-234 |
| Nr4a2 | 1,53E-56 | 0,38 | 0,39 | 0,275 | 2,85E-52 |
| Cd160 | 2,54E-108 | 0,38 | 0,475 | 0,307 | 4,72E-104 |
| Bcl2l11 | 2,27E-60 | 0,38 | 0,702 | 0,6 | 4,23E-56 |
| Slamf1 | 2,80E-146 | 0,38 | 0,556 | 0,353 | 5,19E-142 |
| Plk3 | 4,14E-134 | 0,38 | 0,459 | 0,28 | 7,69E-130 |
| Pstpip1 | 5,27E-161 | 0,38 | 0,887 | 0,809 | 9,80E-157 |
| Zc3h12d | 1,18E-117 | 0,38 | 0,691 | 0,528 | 2,18E-113 |
| Sla | 9,63E-100 | 0,38 | 0,877 | 0,816 | 1,79E-95 |
| Crispld2 | 4,84E-104 | 0,38 | 0,169 | 0,066 | 8,99E-100 |
| Pgk1 | 6,82E-119 | 0,37 | 0,537 | 0,381 | 1,27E-114 |
| Clic1 | 5,70E-193 | 0,37 | 0,883 | 0,774 | 1,06E-188 |
| Slc2a1 | 4,00E-119 | 0,37 | 0,511 | 0,337 | 7,44E-115 |
| Rora | 9,50E-151 | 0,37 | 0,76 | 0,556 | 1,77E-146 |
| Gbp5 | 4,06E-87 | 0,37 | 0,702 | 0,563 | 7,54E-83 |
| Sh2d2a | 5,47E-162 | 0,37 | 0,961 | 0,93 | 1,02E-157 |

| | | | | | |
|---|---|---|---|---|---|
| Mvb12a | 2,70E-195 | 0,37 | 0,587 | 0,368 | 5,03E-191 |
| Cdkl3 | 2,18E-238 | 0,37 | 0,331 | 0,122 | 4,05E-234 |
| Fam20a | 9,75E-272 | 0,37 | 0,15 | 0,005 | 1,81E-267 |
| H1f0 | 2,30E-131 | 0,37 | 0,299 | 0,148 | 4,27E-127 |
| Mpdu1 | 2,08E-188 | 0,37 | 0,634 | 0,421 | 3,86E-184 |
| Twsg1 | 4,31E-152 | 0,37 | 0,382 | 0,204 | 8,00E-148 |
| Il1rl1 | 5,55E-187 | 0,36 | 0,151 | 0,026 | 1,03E-182 |
| Keap1 | 1,43E-171 | 0,36 | 0,658 | 0,448 | 2,66E-167 |
| Rbm47 | 1,14E-273 | 0,36 | 0,301 | 0,087 | 2,11E-269 |
| Tspan3 | 9,08E-111 | 0,36 | 0,713 | 0,543 | 1,69E-106 |
| Padi2 | 2,06E-142 | 0,36 | 0,703 | 0,536 | 3,83E-138 |
| Cnp | 4,50E-103 | 0,36 | 0,763 | 0,644 | 8,36E-99 |
| Znfx1 | 1,72E-104 | 0,36 | 0,585 | 0,422 | 3,20E-100 |
| Itm2c | 7,07E-122 | 0,36 | 0,767 | 0,64 | 1,31E-117 |
| Trim21 | 1,23E-93 | 0,36 | 0,607 | 0,454 | 2,29E-89 |
| Ppp1r3b | 5,56E-153 | 0,35 | 0,253 | 0,103 | 1,03E-148 |
| Nfil3 | 2,48E-132 | 0,35 | 0,375 | 0,207 | 4,61E-128 |
| S100a13 | 1,47E-133 | 0,35 | 0,767 | 0,61 | 2,73E-129 |
| Ndufa11 | 1,07E-164 | 0,35 | 0,662 | 0,471 | 1,99E-160 |
| Trim34a | 3,69E-119 | 0,35 | 0,643 | 0,469 | 6,86E-115 |
| Sys1 | 1,39E-153 | 0,35 | 0,76 | 0,61 | 2,58E-149 |
| Slc25a20 | 2,67E-180 | 0,35 | 0,521 | 0,306 | 4,97E-176 |
| Ufc1 | 2,62E-148 | 0,35 | 0,698 | 0,537 | 4,87E-144 |
| Pomp | 7,65E-149 | 0,35 | 0,754 | 0,606 | 1,42E-144 |
| Sirt2 | 1,79E-156 | 0,35 | 0,753 | 0,591 | 3,32E-152 |
| Xdh | 9,97E-108 | 0,35 | 0,373 | 0,221 | 1,85E-103 |
| Ndufa8 | 1,62E-150 | 0,34 | 0,75 | 0,584 | 3,01E-146 |
| Zfp52 | 8,95E-145 | 0,34 | 0,468 | 0,281 | 1,66E-140 |
| Adprm | 2,82E-137 | 0,34 | 0,513 | 0,331 | 5,24E-133 |
| Chsy1 | 3,99E-104 | 0,34 | 0,842 | 0,728 | 7,42E-100 |
| Ehd4 | 1,04E-176 | 0,34 | 0,33 | 0,149 | 1,93E-172 |
| Tmem160 | 7,98E-155 | 0,34 | 0,692 | 0,503 | 1,48E-150 |
| Cblb | 4,80E-77 | 0,34 | 0,829 | 0,763 | 8,92E-73 |
| Fam110a | 1,90E-145 | 0,34 | 0,389 | 0,213 | 3,54E-141 |
| Cd53 | 5,46E-127 | 0,34 | 0,972 | 0,961 | 1,02E-122 |
| Ehd1 | 6,49E-109 | 0,34 | 0,592 | 0,441 | 1,21E-104 |
| Ndufb7 | 1,13E-155 | 0,34 | 0,816 | 0,664 | 2,10E-151 |
| Npc2 | 2,86E-108 | 0,33 | 0,935 | 0,888 | 5,32E-104 |
| Ssr2 | 1,05E-142 | 0,33 | 0,681 | 0,517 | 1,96E-138 |
| Ap1s3 | 2,29E-193 | 0,33 | 0,4 | 0,195 | 4,25E-189 |
| Fxyd5 | 1,17E-120 | 0,33 | 0,955 | 0,934 | 2,18E-116 |
| Csrp1 | 6,10E-90 | 0,33 | 0,522 | 0,373 | 1,13E-85 |
| Ostm1 | 1,72E-147 | 0,33 | 0,651 | 0,465 | 3,19E-143 |
| Mndal | 8,03E-53 | 0,33 | 0,897 | 0,859 | 1,49E-48 |
| Cdkn2d | 6,98E-94 | 0,33 | 0,674 | 0,536 | 1,30E-89 |
| Prdx5 | 8,55E-100 | 0,33 | 0,729 | 0,596 | 1,59E-95 |

| | | | | | |
|---|---|---|---|---|---|
| Pkp3 | 1,46E-106 | 0,33 | 0,603 | 0,441 | 2,72E-102 |
| Etnk1 | 1,03E-86 | 0,33 | 0,891 | 0,831 | 1,92E-82 |
| Arap2 | 5,73E-91 | 0,33 | 0,746 | 0,617 | 1,06E-86 |
| Cisd3 | 2,64E-131 | 0,33 | 0,444 | 0,27 | 4,91E-127 |
| Ppp1r16b | 8,57E-106 | 0,33 | 0,882 | 0,801 | 1,59E-101 |
| Pld3 | 1,60E-106 | 0,32 | 0,899 | 0,842 | 2,98E-102 |
| Slc25a22 | 2,23E-124 | 0,32 | 0,502 | 0,323 | 4,14E-120 |
| Stk39 | 9,63E-157 | 0,32 | 0,6 | 0,392 | 1,79E-152 |
| Cox17 | 2,86E-98 | 0,32 | 0,789 | 0,692 | 5,32E-94 |
| Psma4 | 1,35E-133 | 0,32 | 0,825 | 0,721 | 2,51E-129 |
| Snrpd1 | 9,69E-112 | 0,32 | 0,629 | 0,474 | 1,80E-107 |
| Tap1 | 9,82E-181 | 0,32 | 0,961 | 0,94 | 1,82E-176 |
| Psmb2 | 2,09E-129 | 0,32 | 0,77 | 0,641 | 3,89E-125 |
| Mapk6 | 6,20E-128 | 0,32 | 0,563 | 0,381 | 1,15E-123 |
| Hist1h1b | 9,07E-34 | 0,32 | 0,174 | 0,11 | 1,69E-29 |
| Cytip | 2,92E-115 | 0,32 | 0,965 | 0,961 | 5,43E-111 |
| Tgfb1 | 1,88E-64 | 0,32 | 0,729 | 0,638 | 3,49E-60 |
| Arf4 | 1,97E-104 | 0,32 | 0,772 | 0,654 | 3,66E-100 |
| Car5b | 7,99E-203 | 0,32 | 0,292 | 0,107 | 1,48E-198 |
| Camk4 | 1,28E-112 | 0,32 | 0,8 | 0,684 | 2,38E-108 |
| Aars | 9,92E-86 | 0,31 | 0,604 | 0,462 | 1,84E-81 |
| Resf1 | 8,94E-80 | 0,31 | 0,706 | 0,562 | 1,66E-75 |
| Uba7 | 3,55E-61 | 0,31 | 0,662 | 0,544 | 6,59E-57 |
| Fam49b | 5,58E-157 | 0,31 | 0,945 | 0,918 | 1,04E-152 |
| Trp53i11 | 2,74E-102 | 0,31 | 0,704 | 0,553 | 5,10E-98 |
| Ndfip1 | 1,05E-91 | 0,31 | 0,942 | 0,928 | 1,96E-87 |
| Ssr4 | 9,19E-93 | 0,31 | 0,752 | 0,644 | 1,71E-88 |
| Ube2v1 | 5,00E-143 | 0,31 | 0,829 | 0,721 | 9,30E-139 |
| Psmb10 | 3,47E-84 | 0,31 | 0,852 | 0,764 | 6,45E-80 |
| Gypc | 9,94E-128 | 0,31 | 0,314 | 0,159 | 1,85E-123 |
| Rnf19b | 3,02E-80 | 0,31 | 0,825 | 0,722 | 5,62E-76 |
| Slc39a4 | 7,59E-164 | 0,31 | 0,391 | 0,2 | 1,41E-159 |
| Ndufs5 | 2,16E-290 | 0,31 | 0,383 | 0,14 | 4,01E-286 |
| Cables1 | 4,58E-114 | 0,31 | 0,27 | 0,132 | 8,51E-110 |
| Mki67 | 1,36E-32 | 0,31 | 0,319 | 0,237 | 2,52E-28 |
| Ermn | 1,43E-179 | 0,31 | 0,12 | 0,012 | 2,65E-175 |
| Slfn3 | 4,53E-209 | 0,31 | 0,229 | 0,064 | 8,41E-205 |
| Nucb1 | 8,35E-103 | 0,31 | 0,848 | 0,76 | 1,55E-98 |
| Apobec3 | 1,53E-124 | 0,31 | 0,943 | 0,903 | 2,84E-120 |
| Cfl1 | 2,98E-169 | 0,31 | 0,967 | 0,941 | 5,53E-165 |
| Cdk6 | 7,57E-74 | 0,30 | 0,57 | 0,447 | 1,41E-69 |
| Cacnb3 | 2,75E-177 | 0,30 | 0,226 | 0,073 | 5,10E-173 |
| Asns | 1,00E-107 | 0,30 | 0,184 | 0,074 | 1,86E-103 |
| Elob | 2,87E-131 | 0,30 | 0,871 | 0,775 | 5,33E-127 |
| Cox5b | 1,63E-128 | 0,30 | 0,807 | 0,69 | 3,02E-124 |
| Ecm1 | 8,94E-62 | 0,30 | 0,139 | 0,065 | 1,66E-57 |

| | | | | | |
|---|---|---|---|---|---|
| Nfkbib | 3,94E-100 | 0,30 | 0,651 | 0,492 | 7,33E-96 |
| Frmd4a | 2,11E-147 | 0,30 | 0,262 | 0,111 | 3,93E-143 |
| Pttg1 | 4,30E-67 | 0,30 | 0,739 | 0,623 | 7,99E-63 |
| Atxn1 | 2,58E-118 | 0,30 | 0,654 | 0,473 | 4,79E-114 |
| Psmb8 | 2,36E-137 | 0,30 | 0,96 | 0,94 | 4,39E-133 |
| BC147527 | 8,71E-142 | 0,30 | 0,326 | 0,162 | 1,62E-137 |
| Coro1a | 2,68E-199 | 0,30 | 0,985 | 0,985 | 4,98E-195 |
| Snx10 | 3,07E-122 | 0,30 | 0,52 | 0,339 | 5,71E-118 |
| Sdhb | 8,23E-115 | 0,30 | 0,785 | 0,662 | 1,53E-110 |
| F2r | 4,06E-96 | 0,30 | 0,837 | 0,721 | 7,54E-92 |
| Stmn1 | 1,56E-13 | 0,29 | 0,188 | 0,151 | 2,90E-09 |
| Adamts14 | 5,52E-172 | 0,29 | 0,25 | 0,09 | 1,03E-167 |
| Erp44 | 1,34E-87 | 0,29 | 0,712 | 0,597 | 2,48E-83 |
| Gsr | 1,73E-114 | 0,29 | 0,587 | 0,413 | 3,22E-110 |
| Il2rb | 4,64E-131 | 0,29 | 0,964 | 0,946 | 8,63E-127 |
| Rab1b | 1,37E-123 | 0,29 | 0,805 | 0,675 | 2,55E-119 |
| Adam19 | 6,05E-103 | 0,29 | 0,804 | 0,685 | 1,12E-98 |
| Vasp | 2,49E-145 | 0,29 | 0,966 | 0,945 | 4,62E-141 |
| Plaat3 | 2,82E-67 | 0,29 | 0,873 | 0,8 | 5,24E-63 |
| Pcgf5 | 4,47E-92 | 0,29 | 0,549 | 0,387 | 8,31E-88 |
| Anapc16 | 1,53E-131 | 0,29 | 0,576 | 0,387 | 2,85E-127 |
| Ctsb | 1,97E-106 | 0,29 | 0,829 | 0,725 | 3,66E-102 |
| Parp10 | 8,68E-67 | 0,29 | 0,66 | 0,534 | 1,61E-62 |
| Irf9 | 5,97E-113 | 0,29 | 0,765 | 0,604 | 1,11E-108 |
| Tmem128 | 1,40E-103 | 0,29 | 0,743 | 0,618 | 2,61E-99 |
| Ddit3 | 6,78E-72 | 0,29 | 0,483 | 0,346 | 1,26E-67 |
| Emp1 | 3,49E-72 | 0,28 | 0,289 | 0,174 | 6,49E-68 |
| Ube2c | 4,09E-65 | 0,28 | 0,13 | 0,056 | 7,61E-61 |
| Trim56 | 1,55E-64 | 0,28 | 0,839 | 0,746 | 2,89E-60 |
| Ccl5 | 1,59E-68 | 0,28 | 0,971 | 0,901 | 2,96E-64 |
| Slc15a3 | 3,06E-172 | 0,28 | 0,166 | 0,038 | 5,69E-168 |
| Pnpla2 | 7,06E-119 | 0,28 | 0,695 | 0,517 | 1,31E-114 |
| Ttc39b | 4,33E-62 | 0,28 | 0,886 | 0,812 | 8,05E-58 |
| Wipf1 | 2,36E-143 | 0,28 | 0,96 | 0,94 | 4,39E-139 |
| Mov10 | 3,56E-63 | 0,28 | 0,526 | 0,399 | 6,61E-59 |
| Csnk2b | 5,78E-116 | 0,28 | 0,769 | 0,643 | 1,07E-111 |
| Abracl | 1,31E-94 | 0,28 | 0,85 | 0,743 | 2,43E-90 |
| Hk1 | 2,13E-99 | 0,28 | 0,806 | 0,7 | 3,95E-95 |
| Unc119b | 5,34E-92 | 0,28 | 0,689 | 0,544 | 9,92E-88 |
| Cst7 | 2,99E-78 | 0,28 | 0,806 | 0,701 | 5,55E-74 |
| Psme1 | 7,73E-122 | 0,28 | 0,954 | 0,91 | 1,44E-117 |
| Snw1 | 1,22E-88 | 0,28 | 0,754 | 0,626 | 2,26E-84 |
| Ctla2b | 2,29E-79 | 0,28 | 0,542 | 0,395 | 4,26E-75 |
| Tbkbp1 | 2,50E-117 | 0,28 | 0,345 | 0,19 | 4,64E-113 |
| Crlf2 | 3,69E-87 | 0,28 | 0,697 | 0,558 | 6,86E-83 |
| Fen1 | 1,26E-112 | 0,28 | 0,379 | 0,218 | 2,35E-108 |

| | | | | | |
|---|---|---|---|---|---|
| Aebp2 | 3,69E-79 | 0,27 | 0,884 | 0,839 | 6,86E-75 |
| Adap1 | 5,37E-102 | 0,27 | 0,508 | 0,345 | 9,98E-98 |
| Ctss | 4,50E-115 | 0,27 | 0,851 | 0,737 | 8,36E-111 |
| Tmbim6 | 4,30E-128 | 0,27 | 0,912 | 0,866 | 8,00E-124 |
| Stk32c | 2,28E-124 | 0,27 | 0,45 | 0,271 | 4,23E-120 |
| Epas1 | 6,73E-207 | 0,27 | 0,161 | 0,026 | 1,25E-202 |
| Sdf4 | 4,05E-89 | 0,27 | 0,922 | 0,895 | 7,53E-85 |
| Ier3 | 5,35E-123 | 0,27 | 0,17 | 0,056 | 9,95E-119 |
| Tmed9 | 1,64E-109 | 0,27 | 0,862 | 0,78 | 3,05E-105 |
| Rab5c | 7,55E-123 | 0,27 | 0,872 | 0,775 | 1,40E-118 |
| Agpat4 | 1,83E-112 | 0,27 | 0,452 | 0,284 | 3,41E-108 |
| Sipa1l1 | 2,99E-87 | 0,27 | 0,687 | 0,539 | 5,55E-83 |
| Bag1 | 2,00E-103 | 0,27 | 0,805 | 0,678 | 3,71E-99 |
| Mctp2 | 1,75E-76 | 0,27 | 0,786 | 0,682 | 3,26E-72 |
| Ostf1 | 4,73E-63 | 0,27 | 0,884 | 0,826 | 8,79E-59 |
| Mitd1 | 3,42E-111 | 0,27 | 0,53 | 0,35 | 6,36E-107 |
| Litaf | 1,16E-109 | 0,27 | 0,677 | 0,492 | 2,15E-105 |
| Srp9 | 1,28E-103 | 0,27 | 0,764 | 0,627 | 2,39E-99 |
| Snrpe | 5,26E-100 | 0,27 | 0,71 | 0,569 | 9,78E-96 |
| Fermt3 | 6,81E-100 | 0,27 | 0,853 | 0,779 | 1,26E-95 |
| Sp100 | 2,22E-107 | 0,27 | 0,962 | 0,956 | 4,12E-103 |
| Rabepk | 8,88E-182 | 0,27 | 0,383 | 0,182 | 1,65E-177 |
| Tnfrsf8 | 1,47E-88 | 0,27 | 0,142 | 0,053 | 2,73E-84 |
| Slc2a3 | 1,87E-31 | 0,27 | 0,7 | 0,624 | 3,47E-27 |
| Adss | 2,85E-91 | 0,27 | 0,696 | 0,561 | 5,30E-87 |
| Rtca | 4,57E-135 | 0,27 | 0,451 | 0,265 | 8,50E-131 |
| Themis2 | 1,29E-144 | 0,27 | 0,268 | 0,113 | 2,40E-140 |
| Rnf19a | 2,52E-81 | 0,26 | 0,627 | 0,481 | 4,69E-77 |
| Arhgef10 | 1,05E-182 | 0,26 | 0,171 | 0,037 | 1,95E-178 |
| Adam8 | 1,71E-67 | 0,26 | 0,433 | 0,301 | 3,18E-63 |
| Slc25a5 | 1,26E-74 | 0,26 | 0,777 | 0,685 | 2,34E-70 |
| Samd9l | 1,78E-39 | 0,26 | 0,725 | 0,644 | 3,32E-35 |
| Pmepa1 | 1,25E-38 | 0,26 | 0,57 | 0,48 | 2,32E-34 |
| Ccnb2 | 7,72E-45 | 0,26 | 0,109 | 0,052 | 1,43E-40 |
| Arpc2 | 5,32E-114 | 0,26 | 0,862 | 0,776 | 9,89E-110 |
| Atp5d | 4,69E-106 | 0,26 | 0,878 | 0,807 | 8,72E-102 |
| Spsb3 | 2,84E-100 | 0,26 | 0,691 | 0,528 | 5,27E-96 |
| Rnh1 | 1,89E-119 | 0,26 | 0,639 | 0,463 | 3,51E-115 |
| Tpgs1 | 2,99E-131 | 0,26 | 0,54 | 0,347 | 5,56E-127 |
| Lrrk1 | 7,91E-105 | 0,26 | 0,592 | 0,42 | 1,47E-100 |
| Actg1 | 5,12E-80 | 0,26 | 0,961 | 0,957 | 9,52E-76 |
| Uqcrfs1 | 1,28E-90 | 0,26 | 0,806 | 0,696 | 2,37E-86 |
| F2rl2 | 5,91E-148 | 0,26 | 0,241 | 0,094 | 1,10E-143 |
| Mif | 6,55E-72 | 0,26 | 0,502 | 0,369 | 1,22E-67 |
| Tuba1c | 1,57E-40 | 0,26 | 0,524 | 0,423 | 2,92E-36 |
| Mlkl | 6,30E-223 | 0,26 | 0,197 | 0,041 | 1,17E-218 |

| | | | | | |
|---|---|---|---|---|---|
| Vim | 2,04E-42 | 0,26 | 0,937 | 0,919 | 3,79E-38 |
| Txndc17 | 1,87E-86 | 0,26 | 0,743 | 0,605 | 3,48E-82 |
| Cmc1 | 5,53E-81 | 0,26 | 0,641 | 0,485 | 1,03E-76 |
| Gas2 | 1,13E-172 | 0,26 | 0,21 | 0,064 | 2,10E-168 |
| Tor1aip2 | 3,00E-48 | 0,25 | 0,766 | 0,67 | 5,57E-44 |
| Calm3 | 1,19E-82 | 0,25 | 0,783 | 0,668 | 2,21E-78 |
| Neb | 3,11E-190 | 0,25 | 0,152 | 0,025 | 5,79E-186 |
| Dcp2 | 5,32E-95 | 0,25 | 0,577 | 0,412 | 9,88E-91 |
| Pdlim2 | 1,79E-111 | 0,25 | 0,548 | 0,37 | 3,32E-107 |
| Gpi1 | 1,77E-74 | 0,25 | 0,822 | 0,738 | 3,29E-70 |
| Prelid3b | 1,45E-100 | 0,25 | 0,558 | 0,394 | 2,69E-96 |
| Morc3 | 9,32E-90 | 0,25 | 0,581 | 0,417 | 1,73E-85 |
| Psma6 | 1,44E-82 | 0,25 | 0,738 | 0,62 | 2,68E-78 |
| Otulin | 2,27E-65 | 0,25 | 0,793 | 0,704 | 4,21E-61 |
| Smox | 2,52E-17 | 0,25 | 0,211 | 0,164 | 4,68E-13 |
| Gstt2 | 1,47E-106 | 0,25 | 0,355 | 0,203 | 2,73E-102 |
| Tapbp | 3,83E-59 | 0,25 | 0,96 | 0,958 | 7,13E-55 |
| Myo1f | 1,16E-29 | -0,25 | 0,714 | 0,699 | 2,16E-25 |
| Tnks2 | 6,13E-65 | -0,25 | 0,726 | 0,746 | 1,14E-60 |
| Ganab | 8,55E-58 | -0,25 | 0,689 | 0,71 | 1,59E-53 |
| Ubqln2 | 2,41E-43 | -0,25 | 0,435 | 0,486 | 4,49E-39 |
| Mgst2 | 7,72E-44 | -0,25 | 0,312 | 0,379 | 1,43E-39 |
| mt-Nd4l | 2,41E-60 | -0,25 | 0,712 | 0,768 | 4,47E-56 |
| Idnk | 1,67E-51 | -0,25 | 0,669 | 0,684 | 3,10E-47 |
| Sh2b1 | 4,89E-60 | -0,25 | 0,605 | 0,647 | 9,08E-56 |
| Ddx3x | 1,97E-65 | -0,25 | 0,767 | 0,794 | 3,66E-61 |
| Pnn | 1,88E-61 | -0,25 | 0,567 | 0,611 | 3,49E-57 |
| Tle3 | 3,03E-75 | -0,25 | 0,777 | 0,819 | 5,63E-71 |
| Trim39 | 1,01E-42 | -0,25 | 0,334 | 0,392 | 1,88E-38 |
| Glg1 | 2,59E-63 | -0,25 | 0,72 | 0,741 | 4,81E-59 |
| Gabbr1 | 1,13E-30 | -0,25 | 0,606 | 0,617 | 2,10E-26 |
| Xcl1 | 1,39E-30 | -0,25 | 0,125 | 0,183 | 2,59E-26 |
| Dhx15 | 1,38E-70 | -0,25 | 0,765 | 0,796 | 2,56E-66 |
| Adgre5 | 6,37E-42 | -0,25 | 0,823 | 0,831 | 1,18E-37 |
| Phf21a | 2,90E-40 | -0,25 | 0,433 | 0,481 | 5,39E-36 |
| Yme1l1 | 4,74E-72 | -0,25 | 0,729 | 0,75 | 8,81E-68 |
| Dtnb | 4,66E-48 | -0,25 | 0,439 | 0,491 | 8,66E-44 |
| Ptpn6 | 4,02E-31 | -0,25 | 0,779 | 0,808 | 7,47E-27 |
| Map4k2 | 8,22E-64 | -0,25 | 0,713 | 0,74 | 1,53E-59 |
| Gprasp1 | 2,93E-95 | -0,25 | 0,07 | 0,162 | 5,45E-91 |
| Foxk1 | 8,20E-47 | -0,25 | 0,462 | 0,509 | 1,52E-42 |
| Abcg3 | 3,76E-117 | -0,25 | 0,089 | 0,202 | 6,99E-113 |
| Atp8a1 | 1,36E-46 | -0,25 | 0,258 | 0,331 | 2,52E-42 |
| Scai | 1,08E-59 | -0,25 | 0,265 | 0,351 | 2,00E-55 |
| Bcl9l | 3,14E-31 | -0,25 | 0,5 | 0,525 | 5,84E-27 |
| Sap25 | 6,79E-41 | -0,26 | 0,343 | 0,4 | 1,26E-36 |

| | | | | | |
|---|---|---|---|---|---|
| Zrsr1 | 4,27E-62 | -0,26 | 0,148 | 0,232 | 7,94E-58 |
| Usp48 | 1,73E-69 | -0,26 | 0,717 | 0,741 | 3,21E-65 |
| Naa30 | 5,72E-50 | -0,26 | 0,565 | 0,596 | 1,06E-45 |
| Myb | 1,73E-35 | -0,26 | 0,206 | 0,129 | 3,22E-31 |
| Dennd1c | 2,27E-64 | -0,26 | 0,752 | 0,777 | 4,21E-60 |
| Znrf2 | 1,50E-91 | -0,26 | 0,868 | 0,882 | 2,79E-87 |
| B4galt1 | 7,79E-69 | -0,26 | 0,73 | 0,763 | 1,45E-64 |
| Eif4b | 3,13E-82 | -0,26 | 0,739 | 0,772 | 5,81E-78 |
| Cdc25b | 3,92E-56 | -0,26 | 0,26 | 0,349 | 7,28E-52 |
| Ddb2 | 1,17E-55 | -0,26 | 0,244 | 0,324 | 2,18E-51 |
| Sec11a | 1,55E-56 | -0,26 | 0,788 | 0,797 | 2,88E-52 |
| March7 | 2,11E-64 | -0,26 | 0,786 | 0,8 | 3,92E-60 |
| Pan3 | 2,46E-44 | -0,26 | 0,511 | 0,543 | 4,57E-40 |
| Hsp90aa1 | 2,05E-58 | -0,26 | 0,612 | 0,666 | 3,81E-54 |
| Paxbp1 | 6,07E-51 | -0,26 | 0,44 | 0,493 | 1,13E-46 |
| Ss18 | 1,70E-62 | -0,26 | 0,757 | 0,776 | 3,16E-58 |
| Brd8 | 1,88E-51 | -0,26 | 0,466 | 0,519 | 3,50E-47 |
| Rbm4 | 3,19E-47 | -0,26 | 0,588 | 0,612 | 5,93E-43 |
| Xpc | 7,81E-63 | -0,26 | 0,238 | 0,326 | 1,45E-58 |
| Trim33 | 5,40E-45 | -0,26 | 0,55 | 0,579 | 1,00E-40 |
| Wls | 5,35E-105 | -0,26 | 0,07 | 0,17 | 9,93E-101 |
| Atn1 | 4,12E-36 | -0,26 | 0,413 | 0,457 | 7,65E-32 |
| Usp12 | 1,84E-30 | -0,26 | 0,319 | 0,36 | 3,41E-26 |
| Prkcz | 1,19E-125 | -0,26 | 0,095 | 0,214 | 2,21E-121 |
| Acss1 | 1,06E-63 | -0,26 | 0,243 | 0,338 | 1,97E-59 |
| Dyrk2 | 1,04E-34 | -0,26 | 0,471 | 0,506 | 1,93E-30 |
| Trim44 | 2,16E-50 | -0,26 | 0,551 | 0,593 | 4,01E-46 |
| Hnrnpa2b1 | 1,24E-125 | -0,26 | 0,901 | 0,923 | 2,31E-121 |
| Hexim1 | 4,83E-31 | -0,26 | 0,633 | 0,637 | 8,98E-27 |
| Phc3 | 1,18E-63 | -0,26 | 0,603 | 0,645 | 2,19E-59 |
| Exoc6b | 8,76E-85 | -0,26 | 0,158 | 0,262 | 1,63E-80 |
| Itsn2 | 6,71E-81 | -0,26 | 0,739 | 0,761 | 1,25E-76 |
| Notch2 | 8,03E-47 | -0,26 | 0,451 | 0,501 | 1,49E-42 |
| Sdha | 4,84E-86 | -0,26 | 0,788 | 0,817 | 9,00E-82 |
| Lysmd2 | 6,33E-46 | -0,26 | 0,319 | 0,396 | 1,18E-41 |
| Prpf4b | 7,10E-68 | -0,26 | 0,708 | 0,741 | 1,32E-63 |
| Abcb1b | 1,24E-22 | -0,26 | 0,296 | 0,34 | 2,31E-18 |
| Dennd4a | 1,88E-21 | -0,26 | 0,784 | 0,756 | 3,49E-17 |
| Kmt2a | 5,39E-82 | -0,26 | 0,791 | 0,816 | 1,00E-77 |
| St3gal1 | 1,95E-58 | -0,26 | 0,465 | 0,537 | 3,62E-54 |
| Atf7 | 4,16E-38 | -0,26 | 0,462 | 0,5 | 7,74E-34 |
| Carmil2 | 6,57E-46 | -0,26 | 0,596 | 0,624 | 1,22E-41 |
| Rmnd5a | 1,92E-58 | -0,26 | 0,557 | 0,597 | 3,57E-54 |
| Ice1 | 4,31E-55 | -0,26 | 0,388 | 0,454 | 8,01E-51 |
| Phf20l1 | 1,35E-48 | -0,26 | 0,612 | 0,636 | 2,51E-44 |
| Prpf38b | 5,60E-62 | -0,26 | 0,726 | 0,735 | 1,04E-57 |

| | | | | | |
|---|---|---|---|---|---|
| Cd37 | 4,65E-81 | -0,26 | 0,864 | 0,872 | 8,64E-77 |
| Hnrnpr | 2,49E-66 | -0,26 | 0,608 | 0,653 | 4,63E-62 |
| Ddhd1 | 1,87E-46 | -0,26 | 0,331 | 0,394 | 3,47E-42 |
| Gnl3 | 1,91E-39 | -0,26 | 0,305 | 0,373 | 3,54E-35 |
| Ncln | 3,16E-41 | -0,26 | 0,418 | 0,469 | 5,88E-37 |
| Als2cl | 3,43E-17 | -0,26 | 0,337 | 0,362 | 6,38E-13 |
| Sik2 | 8,57E-53 | -0,26 | 0,388 | 0,452 | 1,59E-48 |
| Pink1 | 2,43E-33 | -0,26 | 0,545 | 0,57 | 4,51E-29 |
| Herc1 | 7,43E-60 | -0,26 | 0,636 | 0,668 | 1,38E-55 |
| Cux1 | 3,67E-57 | -0,26 | 0,449 | 0,52 | 6,82E-53 |
| Slc37a1 | 6,28E-44 | -0,26 | 0,455 | 0,501 | 1,17E-39 |
| Clcf1 | 3,98E-29 | -0,26 | 0,484 | 0,512 | 7,40E-25 |
| Tspyl2 | 4,97E-58 | -0,26 | 0,179 | 0,261 | 9,23E-54 |
| Rabep1 | 6,88E-46 | -0,26 | 0,466 | 0,51 | 1,28E-41 |
| Stx16 | 1,33E-51 | -0,26 | 0,558 | 0,596 | 2,48E-47 |
| Trip4 | 1,70E-43 | -0,27 | 0,428 | 0,472 | 3,17E-39 |
| P2rx7 | 2,21E-18 | -0,27 | 0,384 | 0,42 | 4,10E-14 |
| Rbbp6 | 5,77E-45 | -0,27 | 0,617 | 0,633 | 1,07E-40 |
| Junb | 3,22E-24 | -0,27 | 0,919 | 0,908 | 5,98E-20 |
| Fam117a | 4,73E-70 | -0,27 | 0,377 | 0,478 | 8,78E-66 |
| Atp1b3 | 2,81E-50 | -0,27 | 0,839 | 0,827 | 5,23E-46 |
| Spg7 | 1,57E-50 | -0,27 | 0,488 | 0,539 | 2,91E-46 |
| Ndrg3 | 2,47E-42 | -0,27 | 0,443 | 0,488 | 4,59E-38 |
| Tespa1 | 1,05E-66 | -0,27 | 0,282 | 0,386 | 1,96E-62 |
| Tgfbr1 | 1,36E-82 | -0,27 | 0,162 | 0,265 | 2,54E-78 |
| Aff1 | 3,28E-46 | -0,27 | 0,47 | 0,516 | 6,10E-42 |
| Tbc1d5 | 2,40E-60 | -0,27 | 0,265 | 0,349 | 4,47E-56 |
| Ccnt2 | 3,34E-56 | -0,27 | 0,594 | 0,635 | 6,21E-52 |
| Mgat5 | 1,22E-57 | -0,27 | 0,346 | 0,427 | 2,26E-53 |
| Nebl | 1,15E-73 | -0,27 | 0,109 | 0,199 | 2,14E-69 |
| Mapk1ip1 | 2,03E-54 | -0,27 | 0,222 | 0,302 | 3,77E-50 |
| Elf2 | 4,05E-54 | -0,27 | 0,564 | 0,597 | 7,53E-50 |
| Vps13a | 4,48E-55 | -0,27 | 0,527 | 0,574 | 8,32E-51 |
| Swap70 | 6,43E-47 | -0,27 | 0,138 | 0,208 | 1,19E-42 |
| Fubp1 | 6,74E-66 | -0,27 | 0,662 | 0,713 | 1,25E-61 |
| Rbm5 | 6,01E-79 | -0,27 | 0,756 | 0,778 | 1,12E-74 |
| Impact | 4,25E-38 | -0,27 | 0,297 | 0,354 | 7,91E-34 |
| Golga4 | 8,46E-54 | -0,27 | 0,52 | 0,566 | 1,57E-49 |
| Atp11b | 2,00E-43 | -0,27 | 0,639 | 0,65 | 3,72E-39 |
| Ncf1 | 1,66E-22 | -0,27 | 0,359 | 0,403 | 3,08E-18 |
| Slc35g1 | 1,28E-48 | -0,27 | 0,249 | 0,321 | 2,38E-44 |
| Ezh1 | 3,18E-57 | -0,27 | 0,366 | 0,438 | 5,91E-53 |
| Man2a2 | 1,35E-54 | -0,27 | 0,467 | 0,532 | 2,51E-50 |
| Patz1 | 2,02E-40 | -0,27 | 0,417 | 0,468 | 3,76E-36 |
| Nufip2 | 4,15E-64 | -0,27 | 0,622 | 0,653 | 7,71E-60 |
| Dalrd3 | 7,08E-52 | -0,27 | 0,321 | 0,389 | 1,32E-47 |

| | | | | | |
|---|---|---|---|---|---|
| Ifrd1 | 3,40E-23 | -0,27 | 0,506 | 0,524 | 6,32E-19 |
| Lbh | 5,69E-74 | -0,27 | 0,787 | 0,813 | 1,06E-69 |
| Nup210 | 2,08E-112 | -0,27 | 0,913 | 0,942 | 3,87E-108 |
| Vipr1 | 9,69E-131 | -0,27 | 0,051 | 0,156 | 1,80E-126 |
| Cdk17 | 4,66E-78 | -0,27 | 0,802 | 0,822 | 8,67E-74 |
| Araf | 2,40E-53 | -0,27 | 0,467 | 0,513 | 4,46E-49 |
| Ddx42 | 7,15E-60 | -0,27 | 0,452 | 0,502 | 1,33E-55 |
| Psip1 | 5,92E-73 | -0,27 | 0,671 | 0,724 | 1,10E-68 |
| Tcf12 | 7,49E-66 | -0,27 | 0,535 | 0,589 | 1,39E-61 |
| Nipbl | 5,18E-85 | -0,27 | 0,745 | 0,771 | 9,62E-81 |
| Fam214a | 1,20E-73 | -0,27 | 0,205 | 0,303 | 2,23E-69 |
| Elovl6 | 1,59E-44 | -0,27 | 0,178 | 0,247 | 2,95E-40 |
| Lbr | 4,05E-119 | -0,27 | 0,869 | 0,903 | 7,52E-115 |
| Tmx4 | 1,79E-49 | -0,27 | 0,301 | 0,373 | 3,33E-45 |
| Gnpat | 3,96E-54 | -0,27 | 0,45 | 0,506 | 7,36E-50 |
| Skp1a | 1,06E-67 | -0,27 | 0,541 | 0,586 | 1,98E-63 |
| Ip6k1 | 1,80E-69 | -0,27 | 0,745 | 0,772 | 3,35E-65 |
| Otulinl | 3,04E-71 | -0,27 | 0,757 | 0,782 | 5,65E-67 |
| Bcl7a | 1,33E-110 | -0,28 | 0,101 | 0,213 | 2,47E-106 |
| Rgs3 | 3,91E-59 | -0,28 | 0,872 | 0,867 | 7,26E-55 |
| Setd7 | 4,58E-44 | -0,28 | 0,471 | 0,51 | 8,51E-40 |
| Uvrag | 1,34E-59 | -0,28 | 0,427 | 0,494 | 2,50E-55 |
| Dnajc9 | 3,19E-47 | -0,28 | 0,692 | 0,714 | 5,92E-43 |
| Pds5a | 2,14E-77 | -0,28 | 0,705 | 0,734 | 3,97E-73 |
| Luc7l3 | 1,81E-58 | -0,28 | 0,53 | 0,578 | 3,36E-54 |
| Slc4a7 | 4,03E-33 | -0,28 | 0,398 | 0,436 | 7,48E-29 |
| Tmem63a | 5,17E-53 | -0,28 | 0,316 | 0,394 | 9,60E-49 |
| Cd200r1 | 1,15E-118 | -0,28 | 0,038 | 0,129 | 2,15E-114 |
| Plekha5 | 2,23E-64 | -0,28 | 0,336 | 0,418 | 4,14E-60 |
| Ube3a | 1,51E-71 | -0,28 | 0,642 | 0,681 | 2,81E-67 |
| Hsd17b11 | 7,59E-81 | -0,28 | 0,166 | 0,269 | 1,41E-76 |
| Zfand6 | 8,69E-71 | -0,28 | 0,573 | 0,615 | 1,62E-66 |
| Sp140 | 5,26E-106 | -0,28 | 0,366 | 0,481 | 9,78E-102 |
| Tmem245 | 1,33E-61 | -0,28 | 0,291 | 0,374 | 2,48E-57 |
| Adrb2 | 2,41E-42 | -0,28 | 0,171 | 0,243 | 4,48E-38 |
| Srpk1 | 2,68E-91 | -0,28 | 0,836 | 0,864 | 4,99E-87 |
| Uggt1 | 1,41E-67 | -0,28 | 0,644 | 0,678 | 2,62E-63 |
| Tprgl | 3,98E-76 | -0,28 | 0,639 | 0,673 | 7,39E-72 |
| Slc9a1 | 1,64E-52 | -0,28 | 0,51 | 0,549 | 3,05E-48 |
| Zbtb44 | 5,74E-49 | -0,28 | 0,442 | 0,497 | 1,07E-44 |
| Ppp1r13b | 3,39E-64 | -0,28 | 0,213 | 0,305 | 6,30E-60 |
| Kansl1 | 4,84E-104 | -0,28 | 0,8 | 0,836 | 8,99E-100 |
| Cnot2 | 1,19E-83 | -0,28 | 0,726 | 0,754 | 2,21E-79 |
| Dnaja4 | 1,77E-49 | -0,28 | 0,295 | 0,364 | 3,28E-45 |
| Abcg1 | 4,26E-79 | -0,28 | 0,26 | 0,371 | 7,92E-75 |
| Camk1d | 8,82E-64 | -0,28 | 0,396 | 0,48 | 1,64E-59 |

| | | | | | |
|---|---|---|---|---|---|
| Strn3 | 8,71E-70 | -0,28 | 0,606 | 0,646 | 1,62E-65 |
| Kmt2e | 3,06E-91 | -0,28 | 0,773 | 0,801 | 5,68E-87 |
| Elk4 | 1,89E-84 | -0,28 | 0,765 | 0,786 | 3,51E-80 |
| Smad3 | 6,83E-76 | -0,28 | 0,441 | 0,549 | 1,27E-71 |
| Ppp2r5a | 1,64E-84 | -0,28 | 0,811 | 0,827 | 3,04E-80 |
| Stk4 | 1,07E-96 | -0,28 | 0,784 | 0,812 | 1,99E-92 |
| Gas7 | 2,22E-47 | -0,28 | 0,14 | 0,213 | 4,13E-43 |
| Cdc42se2 | 1,38E-76 | -0,28 | 0,754 | 0,776 | 2,57E-72 |
| Anapc1 | 1,61E-67 | -0,28 | 0,573 | 0,619 | 2,99E-63 |
| Acin1 | 8,31E-99 | -0,28 | 0,796 | 0,821 | 1,54E-94 |
| Atad2 | 2,25E-68 | -0,28 | 0,389 | 0,481 | 4,19E-64 |
| Safb2 | 9,11E-62 | -0,28 | 0,614 | 0,651 | 1,69E-57 |
| Donson | 1,57E-40 | -0,28 | 0,503 | 0,538 | 2,91E-36 |
| Rnf130 | 6,55E-104 | -0,28 | 0,113 | 0,223 | 1,22E-99 |
| Mapk8ip3 | 2,91E-57 | -0,28 | 0,43 | 0,489 | 5,40E-53 |
| Son | 1,14E-157 | -0,28 | 0,944 | 0,957 | 2,11E-153 |
| Mef2d | 2,64E-99 | -0,28 | 0,866 | 0,893 | 4,90E-95 |
| Calm2 | 9,97E-84 | -0,28 | 0,548 | 0,615 | 1,85E-79 |
| Dhx40 | 1,72E-31 | -0,28 | 0,532 | 0,548 | 3,21E-27 |
| Msl2 | 1,33E-75 | -0,28 | 0,755 | 0,769 | 2,47E-71 |
| Sesn1 | 1,02E-99 | -0,28 | 0,123 | 0,231 | 1,89E-95 |
| Ephx1 | 2,68E-121 | -0,29 | 0,05 | 0,148 | 4,98E-117 |
| Stip1 | 3,68E-58 | -0,29 | 0,653 | 0,692 | 6,85E-54 |
| Mast3 | 9,51E-84 | -0,29 | 0,699 | 0,74 | 1,77E-79 |
| Pex6 | 4,48E-89 | -0,29 | 0,195 | 0,303 | 8,33E-85 |
| Prr12 | 1,41E-78 | -0,29 | 0,574 | 0,642 | 2,63E-74 |
| Tbc1d9b | 4,22E-61 | -0,29 | 0,56 | 0,608 | 7,83E-57 |
| Ppm1b | 1,59E-86 | -0,29 | 0,699 | 0,742 | 2,95E-82 |
| Cyth3 | 2,12E-43 | -0,29 | 0,22 | 0,291 | 3,94E-39 |
| N4bp2l2 | 4,33E-77 | -0,29 | 0,651 | 0,685 | 8,04E-73 |
| Taz | 3,72E-51 | -0,29 | 0,398 | 0,459 | 6,92E-47 |
| Rbm25 | 3,75E-86 | -0,29 | 0,777 | 0,806 | 6,96E-82 |
| Nfya | 1,93E-51 | -0,29 | 0,325 | 0,405 | 3,58E-47 |
| Slc25a36 | 1,25E-44 | -0,29 | 0,347 | 0,408 | 2,33E-40 |
| Slc49a4 | 3,53E-119 | -0,29 | 0,114 | 0,237 | 6,56E-115 |
| Sgsh | 3,27E-77 | -0,29 | 0,212 | 0,313 | 6,07E-73 |
| Zfp280d | 1,25E-50 | -0,29 | 0,415 | 0,472 | 2,32E-46 |
| Zbtb20 | 2,27E-172 | -0,29 | 0,042 | 0,162 | 4,22E-168 |
| Hspe1 | 1,36E-39 | -0,29 | 0,526 | 0,556 | 2,52E-35 |
| Pdk1 | 8,69E-25 | -0,29 | 0,239 | 0,283 | 1,61E-20 |
| Pik3c3 | 3,78E-66 | -0,29 | 0,527 | 0,576 | 7,03E-62 |
| Gimap8 | 2,93E-140 | -0,29 | 0,918 | 0,941 | 5,45E-136 |
| Stap1 | 4,31E-57 | -0,29 | 0,433 | 0,495 | 8,02E-53 |
| Zfp683 | 3,07E-25 | -0,29 | 0,33 | 0,383 | 5,70E-21 |
| Pik3c2a | 1,02E-51 | -0,29 | 0,352 | 0,418 | 1,89E-47 |
| Hsp90b1 | 1,10E-75 | -0,29 | 0,691 | 0,734 | 2,04E-71 |

| | | | | | |
|---|---|---|---|---|---|
| Trappc12 | 4,36E-59 | -0,29 | 0,503 | 0,552 | 8,10E-55 |
| mt-Co1 | 2,31E-95 | -0,29 | 0,95 | 0,969 | 4,30E-91 |
| Exoc4 | 6,79E-77 | -0,29 | 0,661 | 0,71 | 1,26E-72 |
| Tesc | 8,42E-47 | -0,29 | 0,259 | 0,335 | 1,56E-42 |
| Arhgap15 | 7,62E-86 | -0,29 | 0,784 | 0,805 | 1,42E-81 |
| Ctnnb1 | 2,25E-83 | -0,29 | 0,719 | 0,75 | 4,18E-79 |
| Rbm6 | 4,06E-77 | -0,29 | 0,586 | 0,635 | 7,54E-73 |
| Tgfbr3 | 1,63E-78 | -0,29 | 0,229 | 0,334 | 3,03E-74 |
| Klhl24 | 5,66E-60 | -0,29 | 0,431 | 0,496 | 1,05E-55 |
| Zfp445 | 2,05E-73 | -0,30 | 0,55 | 0,604 | 3,82E-69 |
| Rprd2 | 2,26E-82 | -0,30 | 0,653 | 0,694 | 4,19E-78 |
| Rasgrp1 | 2,86E-80 | -0,30 | 0,774 | 0,81 | 5,31E-76 |
| Fam120b | 2,33E-55 | -0,30 | 0,367 | 0,431 | 4,34E-51 |
| Tnrc6b | 5,85E-89 | -0,30 | 0,733 | 0,773 | 1,09E-84 |
| Bcl6 | 8,94E-55 | -0,30 | 0,256 | 0,343 | 1,66E-50 |
| Cxcr5 | 4,93E-107 | -0,30 | 0,03 | 0,11 | 9,16E-103 |
| Foxo3 | 4,50E-72 | -0,30 | 0,274 | 0,37 | 8,37E-68 |
| Cited2 | 2,94E-24 | -0,30 | 0,422 | 0,456 | 5,47E-20 |
| Ubr2 | 2,94E-89 | -0,30 | 0,719 | 0,755 | 5,47E-85 |
| Cdkal1 | 9,80E-70 | -0,30 | 0,406 | 0,482 | 1,82E-65 |
| Rassf2 | 2,39E-80 | -0,30 | 0,713 | 0,75 | 4,43E-76 |
| Nxf1 | 7,74E-87 | -0,30 | 0,79 | 0,818 | 1,44E-82 |
| Trappc8 | 9,15E-78 | -0,30 | 0,611 | 0,652 | 1,70E-73 |
| Dmtf1 | 7,72E-72 | -0,30 | 0,46 | 0,529 | 1,43E-67 |
| Hvcn1 | 5,03E-70 | -0,30 | 0,322 | 0,415 | 9,36E-66 |
| Abcc5 | 1,98E-68 | -0,30 | 0,216 | 0,309 | 3,68E-64 |
| Hdac7 | 9,92E-62 | -0,30 | 0,654 | 0,693 | 1,84E-57 |
| Klhdc2 | 2,06E-30 | -0,30 | 0,354 | 0,398 | 3,82E-26 |
| Zkscan3 | 5,45E-69 | -0,30 | 0,278 | 0,369 | 1,01E-64 |
| Oga | 2,97E-96 | -0,30 | 0,793 | 0,821 | 5,52E-92 |
| Ptprc | 5,60E-278 | -0,30 | 0,984 | 0,994 | 1,04E-273 |
| Mat2a | 1,15E-85 | -0,30 | 0,611 | 0,69 | 2,13E-81 |
| Bcor | 9,91E-64 | -0,30 | 0,397 | 0,472 | 1,84E-59 |
| Srsf11 | 1,25E-84 | -0,30 | 0,722 | 0,755 | 2,32E-80 |
| Mppe1 | 2,62E-68 | -0,30 | 0,416 | 0,49 | 4,86E-64 |
| Atp1b1 | 1,23E-145 | -0,30 | 0,034 | 0,136 | 2,28E-141 |
| Hdgfl3 | 1,92E-74 | -0,30 | 0,307 | 0,407 | 3,56E-70 |
| Plekhn1 | 1,52E-111 | -0,30 | 0,14 | 0,261 | 2,82E-107 |
| Epm2aip1 | 2,00E-44 | -0,30 | 0,293 | 0,356 | 3,71E-40 |
| Diaph2 | 9,04E-79 | -0,30 | 0,297 | 0,393 | 1,68E-74 |
| Ctdsp2 | 2,69E-70 | -0,30 | 0,481 | 0,56 | 4,99E-66 |
| Klrd1 | 2,41E-68 | -0,30 | 0,685 | 0,751 | 4,48E-64 |
| Rb1 | 2,81E-69 | -0,31 | 0,532 | 0,579 | 5,22E-65 |
| Polr2a | 3,50E-103 | -0,31 | 0,834 | 0,853 | 6,51E-99 |
| Atxn2l | 4,06E-144 | -0,31 | 0,879 | 0,909 | 7,54E-140 |
| Maz | 4,66E-119 | -0,31 | 0,782 | 0,835 | 8,65E-115 |

| | | | | | |
|---|---|---|---|---|---|
| Smyd3 | 3,72E-86 | -0,31 | 0,492 | 0,583 | 6,91E-82 |
| Ralgapa1 | 1,28E-69 | -0,31 | 0,531 | 0,581 | 2,38E-65 |
| Piezo1 | 1,33E-70 | -0,31 | 0,397 | 0,482 | 2,47E-66 |
| Slamf6 | 3,38E-38 | -0,31 | 0,196 | 0,264 | 6,29E-34 |
| Wbp1 | 4,44E-50 | -0,31 | 0,387 | 0,447 | 8,26E-46 |
| Pcmtd1 | 3,56E-89 | -0,31 | 0,327 | 0,427 | 6,62E-85 |
| Ankrd12 | 2,82E-79 | -0,31 | 0,492 | 0,563 | 5,25E-75 |
| Mbtps1 | 6,60E-81 | -0,31 | 0,507 | 0,574 | 1,23E-76 |
| Cdc37l1 | 4,56E-74 | -0,31 | 0,537 | 0,592 | 8,47E-70 |
| Ppm1h | 1,02E-54 | -0,31 | 0,626 | 0,653 | 1,90E-50 |
| Nlk | 8,54E-67 | -0,31 | 0,56 | 0,612 | 1,59E-62 |
| Mbnl1 | 5,71E-169 | -0,31 | 0,896 | 0,924 | 1,06E-164 |
| Zfp827 | 3,00E-76 | -0,31 | 0,222 | 0,323 | 5,57E-72 |
| Ifngr1 | 2,07E-89 | -0,31 | 0,958 | 0,965 | 3,85E-85 |
| Stim1 | 4,41E-84 | -0,31 | 0,578 | 0,636 | 8,19E-80 |
| Il17ra | 6,19E-89 | -0,31 | 0,774 | 0,801 | 1,15E-84 |
| Calr | 7,61E-77 | -0,31 | 0,665 | 0,709 | 1,41E-72 |
| Lrp6 | 1,02E-91 | -0,31 | 0,301 | 0,414 | 1,90E-87 |
| Arid4a | 7,20E-75 | -0,31 | 0,417 | 0,487 | 1,34E-70 |
| Atp2b1 | 8,24E-48 | -0,31 | 0,594 | 0,615 | 1,53E-43 |
| Gpr34 | 1,10E-55 | -0,31 | 0,155 | 0,235 | 2,05E-51 |
| Hexa | 1,55E-106 | -0,32 | 0,165 | 0,284 | 2,88E-102 |
| Trap1 | 1,50E-78 | -0,32 | 0,389 | 0,476 | 2,78E-74 |
| Gimap3 | 1,56E-142 | -0,32 | 0,936 | 0,954 | 2,91E-138 |
| Smarca2 | 9,51E-73 | -0,32 | 0,412 | 0,488 | 1,77E-68 |
| Snrnp48 | 4,67E-77 | -0,32 | 0,465 | 0,535 | 8,69E-73 |
| Leng8 | 1,35E-115 | -0,32 | 0,846 | 0,886 | 2,51E-111 |
| Adamts10 | 1,20E-69 | -0,32 | 0,651 | 0,684 | 2,23E-65 |
| Jade2 | 3,46E-82 | -0,32 | 0,432 | 0,526 | 6,42E-78 |
| Tcrg-C1 | 5,75E-66 | -0,32 | 0,044 | 0,108 | 1,07E-61 |
| Kdm6a | 7,11E-101 | -0,32 | 0,636 | 0,692 | 1,32E-96 |
| Zmynd11 | 1,59E-93 | -0,32 | 0,758 | 0,789 | 2,96E-89 |
| Polg2 | 1,96E-33 | -0,32 | 0,254 | 0,309 | 3,64E-29 |
| Tomm34 | 1,28E-80 | -0,32 | 0,631 | 0,674 | 2,38E-76 |
| Tmem131l | 6,88E-84 | -0,32 | 0,406 | 0,503 | 1,28E-79 |
| Ccpg1 | 2,00E-81 | -0,32 | 0,432 | 0,515 | 3,72E-77 |
| Kmt2d | 2,15E-151 | -0,32 | 0,896 | 0,923 | 3,99E-147 |
| Unc5cl | 7,80E-38 | -0,32 | 0,454 | 0,503 | 1,45E-33 |
| H2-Ke6 | 2,79E-92 | -0,32 | 0,389 | 0,492 | 5,18E-88 |
| Stat6 | 2,58E-117 | -0,32 | 0,765 | 0,801 | 4,79E-113 |
| Dync1h1 | 4,32E-137 | -0,32 | 0,844 | 0,869 | 8,03E-133 |
| Clk4 | 3,97E-91 | -0,32 | 0,631 | 0,682 | 7,38E-87 |
| Nr1d2 | 9,56E-84 | -0,32 | 0,276 | 0,389 | 1,78E-79 |
| Ccdc117 | 1,31E-56 | -0,32 | 0,335 | 0,408 | 2,43E-52 |
| Nfatc3 | 1,44E-123 | -0,32 | 0,806 | 0,838 | 2,67E-119 |
| Cxcr3 | 8,96E-66 | -0,32 | 0,583 | 0,662 | 1,67E-61 |

| | | | | | |
|---|---|---|---|---|---|
| Pnrc1 | 1,41E-147 | -0,32 | 0,928 | 0,944 | 2,62E-143 |
| Rbm39 | 2,67E-131 | -0,32 | 0,775 | 0,824 | 4,95E-127 |
| Itpr3 | 6,62E-87 | -0,32 | 0,321 | 0,42 | 1,23E-82 |
| Creb1 | 7,72E-66 | -0,32 | 0,617 | 0,656 | 1,43E-61 |
| Bin2 | 1,09E-140 | -0,33 | 0,917 | 0,935 | 2,03E-136 |
| Eef1g | 1,65E-60 | -0,33 | 0,771 | 0,777 | 3,07E-56 |
| Psap | 1,38E-114 | -0,33 | 0,771 | 0,806 | 2,56E-110 |
| Pag1 | 5,09E-68 | -0,33 | 0,355 | 0,446 | 9,47E-64 |
| Hmg20a | 7,02E-63 | -0,33 | 0,323 | 0,406 | 1,30E-58 |
| Ank | 4,62E-100 | -0,33 | 0,293 | 0,415 | 8,59E-96 |
| Akna | 8,63E-150 | -0,33 | 0,892 | 0,923 | 1,60E-145 |
| Pnpla7 | 4,92E-68 | -0,33 | 0,324 | 0,411 | 9,14E-64 |
| Cxcr4 | 1,44E-39 | -0,33 | 0,757 | 0,787 | 2,68E-35 |
| Tmem59 | 8,79E-111 | -0,33 | 0,743 | 0,772 | 1,63E-106 |
| Rere | 3,66E-120 | -0,33 | 0,216 | 0,348 | 6,80E-116 |
| Tardbp | 6,58E-132 | -0,33 | 0,733 | 0,79 | 1,22E-127 |
| Plcg2 | 1,20E-70 | -0,33 | 0,196 | 0,288 | 2,23E-66 |
| Pdlim1 | 1,04E-82 | -0,33 | 0,329 | 0,44 | 1,93E-78 |
| Ccdc88c | 6,03E-106 | -0,33 | 0,795 | 0,825 | 1,12E-101 |
| Txnip | 2,48E-72 | -0,33 | 0,805 | 0,832 | 4,61E-68 |
| Usp28 | 1,46E-111 | -0,33 | 0,106 | 0,218 | 2,72E-107 |
| Acap1 | 6,14E-124 | -0,33 | 0,858 | 0,901 | 1,14E-119 |
| Lats2 | 3,42E-74 | -0,33 | 0,325 | 0,422 | 6,35E-70 |
| Tnfrsf26 | 6,54E-67 | -0,33 | 0,227 | 0,331 | 1,22E-62 |
| Myh9 | 2,17E-150 | -0,33 | 0,855 | 0,89 | 4,02E-146 |
| Dzip1 | 4,05E-111 | -0,33 | 0,122 | 0,239 | 7,53E-107 |
| Ugcg | 6,78E-40 | -0,33 | 0,788 | 0,789 | 1,26E-35 |
| N4bp2l1 | 9,37E-75 | -0,34 | 0,441 | 0,53 | 1,74E-70 |
| Kmt2c | 1,76E-95 | -0,34 | 0,53 | 0,603 | 3,28E-91 |
| Pcmtd2 | 1,10E-77 | -0,34 | 0,46 | 0,534 | 2,04E-73 |
| Rsrc2 | 1,76E-119 | -0,34 | 0,715 | 0,769 | 3,27E-115 |
| Cd96 | 1,17E-91 | -0,34 | 0,686 | 0,743 | 2,17E-87 |
| Srsf2 | 2,44E-92 | -0,34 | 0,588 | 0,659 | 4,53E-88 |
| Sema6d | 5,01E-112 | -0,34 | 0,034 | 0,117 | 9,30E-108 |
| Tlr1 | 1,46E-123 | -0,34 | 0,082 | 0,195 | 2,71E-119 |
| Zfp638 | 4,15E-107 | -0,34 | 0,41 | 0,512 | 7,70E-103 |
| Fbxl12 | 3,02E-81 | -0,34 | 0,405 | 0,494 | 5,62E-77 |
| Arglu1 | 3,00E-115 | -0,34 | 0,747 | 0,797 | 5,57E-111 |
| Dapk2 | 2,24E-59 | -0,34 | 0,232 | 0,317 | 4,15E-55 |
| Crlf3 | 3,84E-77 | -0,34 | 0,799 | 0,809 | 7,14E-73 |
| Kat6b | 2,05E-96 | -0,34 | 0,229 | 0,342 | 3,80E-92 |
| Gramd1a | 2,76E-138 | -0,34 | 0,895 | 0,924 | 5,13E-134 |
| Dnajb9 | 1,62E-70 | -0,34 | 0,36 | 0,441 | 3,00E-66 |
| Sept6 | 9,18E-143 | -0,34 | 0,819 | 0,866 | 1,71E-138 |
| Prpf39 | 5,99E-91 | -0,34 | 0,617 | 0,671 | 1,11E-86 |
| Gdi1 | 1,05E-113 | -0,34 | 0,728 | 0,778 | 1,95E-109 |

| | | | | | |
|---|---|---|---|---|---|
| Atf7ip | 1,51E-100 | -0,34 | 0,709 | 0,765 | 2,81E-96 |
| Ankrd10 | 3,14E-100 | -0,34 | 0,519 | 0,599 | 5,84E-96 |
| Huwe1 | 2,71E-132 | -0,34 | 0,803 | 0,844 | 5,03E-128 |
| Jmjd1c | 9,30E-89 | -0,34 | 0,678 | 0,717 | 1,73E-84 |
| Fgf13 | 2,36E-119 | -0,34 | 0,118 | 0,24 | 4,38E-115 |
| Chordc1 | 7,51E-98 | -0,34 | 0,505 | 0,581 | 1,40E-93 |
| Nop56 | 4,96E-101 | -0,34 | 0,51 | 0,606 | 9,21E-97 |
| Sfpq | 1,36E-122 | -0,34 | 0,763 | 0,803 | 2,53E-118 |
| Ski | 3,09E-129 | -0,34 | 0,83 | 0,877 | 5,74E-125 |
| Mrtfa | 5,38E-114 | -0,34 | 0,671 | 0,727 | 9,99E-110 |
| mt-Nd2 | 1,40E-96 | -0,34 | 0,729 | 0,798 | 2,60E-92 |
| Retreg1 | 6,25E-86 | -0,34 | 0,382 | 0,476 | 1,16E-81 |
| Plcb2 | 3,41E-101 | -0,34 | 0,294 | 0,408 | 6,33E-97 |
| Akt3 | 1,42E-96 | -0,35 | 0,301 | 0,411 | 2,64E-92 |
| Esyt2 | 1,72E-152 | -0,35 | 0,835 | 0,877 | 3,19E-148 |
| Rapgef4 | 8,40E-168 | -0,35 | 0,055 | 0,18 | 1,56E-163 |
| Cbl | 3,80E-95 | -0,35 | 0,671 | 0,714 | 7,06E-91 |
| Fto | 7,09E-116 | -0,35 | 0,316 | 0,437 | 1,32E-111 |
| Aplp2 | 1,31E-65 | -0,35 | 0,463 | 0,52 | 2,43E-61 |
| Kcnn4 | 2,54E-75 | -0,35 | 0,495 | 0,575 | 4,72E-71 |
| Prdx6 | 2,46E-95 | -0,35 | 0,518 | 0,584 | 4,57E-91 |
| Per1 | 6,72E-50 | -0,35 | 0,71 | 0,729 | 1,25E-45 |
| Usp24 | 6,55E-95 | -0,35 | 0,53 | 0,597 | 1,22E-90 |
| Senp7 | 7,61E-107 | -0,35 | 0,542 | 0,621 | 1,41E-102 |
| Btg2 | 4,81E-63 | -0,35 | 0,867 | 0,875 | 8,93E-59 |
| Srsf3 | 1,06E-80 | -0,35 | 0,556 | 0,61 | 1,97E-76 |
| Cdc42ep3 | 1,24E-90 | -0,35 | 0,334 | 0,44 | 2,31E-86 |
| Rasa1 | 4,18E-96 | -0,35 | 0,583 | 0,647 | 7,77E-92 |
| Hmgb1 | 1,15E-54 | -0,35 | 0,46 | 0,513 | 2,15E-50 |
| Axin2 | 7,61E-147 | -0,35 | 0,108 | 0,245 | 1,41E-142 |
| Slc28a2 | 2,63E-109 | -0,36 | 0,672 | 0,736 | 4,89E-105 |
| Srsf7 | 3,11E-94 | -0,36 | 0,731 | 0,782 | 5,79E-90 |
| Gramd4 | 4,75E-16 | -0,36 | 0,398 | 0,401 | 8,82E-12 |
| Tcf20 | 1,14E-104 | -0,36 | 0,571 | 0,636 | 2,12E-100 |
| Bclaf1 | 5,63E-106 | -0,36 | 0,719 | 0,757 | 1,05E-101 |
| Rnf167 | 1,21E-105 | -0,36 | 0,641 | 0,706 | 2,25E-101 |
| Zgpat | 3,71E-128 | -0,36 | 0,694 | 0,759 | 6,90E-124 |
| Patj | 1,06E-169 | -0,36 | 0,058 | 0,186 | 1,97E-165 |
| Cxcr6 | 1,94E-17 | -0,36 | 0,796 | 0,767 | 3,60E-13 |
| Ddx17 | 2,03E-211 | -0,36 | 0,892 | 0,933 | 3,77E-207 |
| Stk10 | 9,14E-174 | -0,36 | 0,836 | 0,879 | 1,70E-169 |
| Sidt1 | 4,02E-28 | -0,36 | 0,319 | 0,353 | 7,47E-24 |
| Map3k3 | 2,31E-123 | -0,36 | 0,453 | 0,557 | 4,30E-119 |
| Stk17b | 1,00E-232 | -0,36 | 0,971 | 0,985 | 1,86E-228 |
| Pcf11 | 1,59E-102 | -0,36 | 0,724 | 0,762 | 2,96E-98 |
| Treml2 | 4,35E-43 | -0,36 | 0,212 | 0,283 | 8,08E-39 |

| | | | | | |
|---|---|---|---|---|---|
| Zhx2 | 5,09E-118 | -0,36 | 0,193 | 0,325 | 9,46E-114 |
| Camk2d | 1,70E-102 | -0,36 | 0,328 | 0,446 | 3,15E-98 |
| Bend4 | 3,99E-63 | -0,36 | 0,255 | 0,343 | 7,42E-59 |
| Rptor | 2,41E-127 | -0,36 | 0,33 | 0,454 | 4,48E-123 |
| Trat1 | 4,83E-133 | -0,36 | 0,134 | 0,27 | 8,97E-129 |
| Phf1 | 6,61E-107 | -0,37 | 0,488 | 0,579 | 1,23E-102 |
| Sgk1 | 7,88E-14 | -0,37 | 0,225 | 0,256 | 1,46E-09 |
| AW549877 | 1,63E-98 | -0,37 | 0,426 | 0,518 | 3,02E-94 |
| Anp32a | 2,08E-157 | -0,37 | 0,79 | 0,835 | 3,87E-153 |
| Tnrc6c | 1,84E-118 | -0,37 | 0,651 | 0,705 | 3,42E-114 |
| Nop10 | 3,33E-81 | -0,37 | 0,536 | 0,617 | 6,18E-77 |
| Nisch | 1,67E-136 | -0,37 | 0,772 | 0,818 | 3,10E-132 |
| Akap8 | 1,11E-128 | -0,37 | 0,684 | 0,749 | 2,06E-124 |
| Ltb | 8,60E-133 | -0,37 | 0,889 | 0,922 | 1,60E-128 |
| Ncor1 | 2,54E-181 | -0,37 | 0,871 | 0,901 | 4,72E-177 |
| Ipcef1 | 1,58E-96 | -0,37 | 0,727 | 0,781 | 2,95E-92 |
| Atp2a3 | 1,38E-134 | -0,37 | 0,763 | 0,821 | 2,57E-130 |
| Rabac1 | 1,01E-109 | -0,37 | 0,763 | 0,799 | 1,88E-105 |
| mt-Cytb | 2,11E-124 | -0,37 | 0,821 | 0,881 | 3,92E-120 |
| Ssbp2 | 1,94E-140 | -0,37 | 0,182 | 0,328 | 3,61E-136 |
| Cyld | 5,62E-159 | -0,37 | 0,782 | 0,831 | 1,04E-154 |
| Rarg | 4,49E-104 | -0,37 | 0,186 | 0,306 | 8,34E-100 |
| Ly9 | 1,97E-103 | -0,37 | 0,579 | 0,658 | 3,66E-99 |
| Hsf1 | 8,27E-133 | -0,37 | 0,328 | 0,46 | 1,54E-128 |
| Emb | 2,78E-89 | -0,37 | 0,865 | 0,883 | 5,16E-85 |
| Gbp8 | 6,43E-99 | -0,37 | 0,669 | 0,731 | 1,19E-94 |
| Lta4h | 5,29E-118 | -0,37 | 0,583 | 0,664 | 9,83E-114 |
| Epc1 | 1,81E-123 | -0,37 | 0,755 | 0,791 | 3,37E-119 |
| Hbp1 | 1,20E-110 | -0,37 | 0,537 | 0,624 | 2,23E-106 |
| Arid1a | 5,48E-177 | -0,37 | 0,8 | 0,846 | 1,02E-172 |
| Fam193b | 2,10E-114 | -0,38 | 0,527 | 0,611 | 3,91E-110 |
| Kif13b | 8,84E-79 | -0,38 | 0,442 | 0,515 | 1,64E-74 |
| Mdm4 | 2,10E-125 | -0,38 | 0,626 | 0,7 | 3,90E-121 |
| Fam53b | 6,19E-133 | -0,38 | 0,703 | 0,771 | 1,15E-128 |
| Lmna | 2,17E-104 | -0,38 | 0,082 | 0,185 | 4,03E-100 |
| Zmym2 | 1,40E-121 | -0,38 | 0,406 | 0,521 | 2,61E-117 |
| Ttc7 | 3,82E-142 | -0,38 | 0,651 | 0,743 | 7,09E-138 |
| Nnt | 3,13E-204 | -0,38 | 0,136 | 0,303 | 5,83E-200 |
| Arhgef3 | 1,58E-154 | -0,38 | 0,733 | 0,823 | 2,93E-150 |
| Ivns1abp | 3,65E-137 | -0,38 | 0,642 | 0,729 | 6,78E-133 |
| Fam169b | 3,26E-37 | -0,38 | 0,528 | 0,545 | 6,05E-33 |
| Pik3ip1 | 2,68E-76 | -0,38 | 0,211 | 0,316 | 4,97E-72 |
| Trp53inp1 | 5,06E-121 | -0,38 | 0,674 | 0,756 | 9,41E-117 |
| Tnfsf8 | 9,34E-184 | -0,38 | 0,082 | 0,233 | 1,74E-179 |
| Vgll4 | 2,08E-126 | -0,38 | 0,696 | 0,759 | 3,86E-122 |
| Hs3st3b1 | 4,88E-132 | -0,39 | 0,127 | 0,26 | 9,07E-128 |

| | | | | | |
|---|---|---|---|---|---|
| St3gal6 | 2,07E-132 | -0,39 | 0,379 | 0,518 | 3,85E-128 |
| Rfx1 | 1,21E-90 | -0,39 | 0,426 | 0,516 | 2,26E-86 |
| Scml4 | 4,36E-90 | -0,39 | 0,336 | 0,448 | 8,10E-86 |
| Il16 | 4,80E-146 | -0,39 | 0,756 | 0,818 | 8,92E-142 |
| Kdm7a | 1,52E-121 | -0,39 | 0,63 | 0,707 | 2,83E-117 |
| Pim2 | 4,73E-84 | -0,39 | 0,328 | 0,435 | 8,78E-80 |
| Tnfaip3 | 2,00E-108 | -0,39 | 0,952 | 0,96 | 3,71E-104 |
| Dnah8 | 1,29E-130 | -0,39 | 0,154 | 0,287 | 2,40E-126 |
| Tatdn2 | 2,47E-89 | -0,39 | 0,67 | 0,71 | 4,59E-85 |
| Kcnc1 | 1,19E-189 | -0,39 | 0,093 | 0,246 | 2,21E-185 |
| Crebzf | 1,21E-99 | -0,39 | 0,537 | 0,614 | 2,24E-95 |
| Elmsan1 | 1,06E-117 | -0,39 | 0,377 | 0,491 | 1,98E-113 |
| Ttc3 | 6,08E-134 | -0,39 | 0,269 | 0,406 | 1,13E-129 |
| Sptbn1 | 8,68E-71 | -0,39 | 0,682 | 0,693 | 1,61E-66 |
| Antxr2 | 1,30E-126 | -0,39 | 0,369 | 0,495 | 2,42E-122 |
| Arnt2 | 1,50E-109 | -0,39 | 0,165 | 0,286 | 2,78E-105 |
| Man1a | 7,44E-122 | -0,39 | 0,707 | 0,762 | 1,38E-117 |
| Prkacb | 1,29E-177 | -0,40 | 0,774 | 0,835 | 2,40E-173 |
| Rcsd1 | 3,73E-120 | -0,40 | 0,425 | 0,552 | 6,92E-116 |
| Rnf138 | 1,98E-98 | -0,40 | 0,802 | 0,806 | 3,68E-94 |
| Smc4 | 3,63E-59 | -0,40 | 0,74 | 0,756 | 6,74E-55 |
| Apbb1ip | 6,66E-222 | -0,40 | 0,818 | 0,89 | 1,24E-217 |
| Prkd3 | 8,60E-153 | -0,40 | 0,307 | 0,456 | 1,60E-148 |
| Zfp36 | 4,71E-22 | -0,40 | 0,697 | 0,69 | 8,76E-18 |
| Ifnar1 | 9,83E-136 | -0,40 | 0,735 | 0,782 | 1,83E-131 |
| Appl2 | 2,82E-133 | -0,40 | 0,18 | 0,317 | 5,24E-129 |
| Nek7 | 2,02E-149 | -0,40 | 0,528 | 0,647 | 3,76E-145 |
| Socs3 | 2,21E-19 | -0,40 | 0,465 | 0,474 | 4,10E-15 |
| Lta | 1,16E-46 | -0,40 | 0,17 | 0,244 | 2,16E-42 |
| Arhgap4 | 1,88E-124 | -0,40 | 0,584 | 0,662 | 3,49E-120 |
| Ube2h | 8,21E-176 | -0,40 | 0,775 | 0,827 | 1,52E-171 |
| Cdkn1b | 2,36E-180 | -0,40 | 0,816 | 0,859 | 4,38E-176 |
| Myadm | 1,73E-41 | -0,41 | 0,283 | 0,345 | 3,22E-37 |
| Znrf3 | 2,34E-191 | -0,41 | 0,193 | 0,369 | 4,34E-187 |
| Arl5c | 2,63E-99 | -0,41 | 0,197 | 0,316 | 4,89E-95 |
| Arl4c | 6,58E-120 | -0,41 | 0,599 | 0,687 | 1,22E-115 |
| Rflnb | 1,51E-217 | -0,41 | 0,03 | 0,161 | 2,81E-213 |
| Top2b | 3,08E-188 | -0,41 | 0,722 | 0,785 | 5,73E-184 |
| Skap1 | 1,96E-241 | -0,41 | 0,878 | 0,919 | 3,65E-237 |
| Susd6 | 7,52E-212 | -0,42 | 0,788 | 0,846 | 1,40E-207 |
| Galnt10 | 2,31E-134 | -0,42 | 0,248 | 0,39 | 4,30E-130 |
| Ppp1r15a | 2,54E-102 | -0,42 | 0,604 | 0,678 | 4,71E-98 |
| Runx3 | 9,99E-168 | -0,42 | 0,837 | 0,88 | 1,86E-163 |
| Irs2 | 4,54E-166 | -0,42 | 0,094 | 0,24 | 8,45E-162 |
| Elovl5 | 8,58E-146 | -0,42 | 0,551 | 0,65 | 1,59E-141 |
| Ctsc | 6,13E-63 | -0,42 | 0,631 | 0,67 | 1,14E-58 |

| | | | | | |
|---|---|---|---|---|---|
| St8sia1 | 7,13E-164 | -0,42 | 0,133 | 0,283 | 1,32E-159 |
| Ctps2 | 5,23E-129 | -0,42 | 0,585 | 0,67 | 9,72E-125 |
| Rbm14 | 9,44E-153 | -0,42 | 0,654 | 0,731 | 1,75E-148 |
| Cd101 | 8,99E-98 | -0,42 | 0,304 | 0,416 | 1,67E-93 |
| Rftn1 | 1,95E-134 | -0,42 | 0,546 | 0,65 | 3,63E-130 |
| Adcy7 | 1,04E-163 | -0,43 | 0,757 | 0,83 | 1,93E-159 |
| Zfp592 | 1,98E-149 | -0,43 | 0,631 | 0,705 | 3,69E-145 |
| Sorl1 | 9,18E-208 | -0,43 | 0,893 | 0,924 | 1,71E-203 |
| Ldlrap1 | 8,96E-133 | -0,43 | 0,208 | 0,347 | 1,66E-128 |
| Cd244a | 4,05E-57 | -0,43 | 0,106 | 0,179 | 7,53E-53 |
| Ikbke | 5,99E-78 | -0,43 | 0,448 | 0,52 | 1,11E-73 |
| Ifnar2 | 2,57E-171 | -0,43 | 0,506 | 0,645 | 4,78E-167 |
| Snx29 | 2,68E-127 | -0,43 | 0,266 | 0,403 | 4,99E-123 |
| Itpr2 | 6,77E-148 | -0,43 | 0,493 | 0,608 | 1,26E-143 |
| Ppp2r2c | 1,49E-76 | -0,43 | 0,254 | 0,355 | 2,76E-72 |
| Parp8 | 7,08E-205 | -0,43 | 0,161 | 0,337 | 1,32E-200 |
| Akap8l | 4,05E-171 | -0,43 | 0,398 | 0,541 | 7,53E-167 |
| Elmo1 | 8,59E-238 | -0,43 | 0,866 | 0,923 | 1,60E-233 |
| Tox | 6,50E-166 | -0,44 | 0,121 | 0,275 | 1,21E-161 |
| Tle4 | 6,26E-162 | -0,44 | 0,437 | 0,571 | 1,16E-157 |
| Znrf1 | 2,61E-175 | -0,44 | 0,752 | 0,823 | 4,85E-171 |
| Zfp652 | 1,88E-156 | -0,44 | 0,498 | 0,623 | 3,50E-152 |
| Slc20a1 | 2,52E-109 | -0,44 | 0,696 | 0,743 | 4,69E-105 |
| Itgax | 3,94E-15 | -0,44 | 0,423 | 0,426 | 7,32E-11 |
| Slc9a9 | 2,44E-178 | -0,44 | 0,407 | 0,561 | 4,53E-174 |
| Ccm2 | 6,54E-135 | -0,44 | 0,585 | 0,663 | 1,22E-130 |
| Spsb1 | 3,97E-125 | -0,44 | 0,194 | 0,329 | 7,37E-121 |
| Itk | 3,49E-268 | -0,44 | 0,976 | 0,985 | 6,49E-264 |
| Dock8 | 1,56E-244 | -0,44 | 0,83 | 0,89 | 2,90E-240 |
| Map4k4 | 1,69E-166 | -0,45 | 0,649 | 0,731 | 3,13E-162 |
| F2rl1 | 9,95E-143 | -0,45 | 0,15 | 0,292 | 1,85E-138 |
| Tob1 | 4,50E-117 | -0,45 | 0,321 | 0,443 | 8,37E-113 |
| Tra2a | 6,19E-158 | -0,45 | 0,571 | 0,675 | 1,15E-153 |
| Srsf6 | 1,02E-167 | -0,45 | 0,792 | 0,854 | 1,90E-163 |
| Zfp831 | 3,39E-170 | -0,45 | 0,427 | 0,571 | 6,29E-166 |
| Srrm2 | 0 | -0,45 | 0,884 | 0,933 | 0 |
| Arrdc3 | 1,75E-107 | -0,45 | 0,243 | 0,365 | 3,26E-103 |
| Rasal3 | 0 | -0,45 | 0,923 | 0,961 | 0 |
| Lpar6 | 7,00E-141 | -0,45 | 0,308 | 0,45 | 1,30E-136 |
| Kif21b | 4,80E-248 | -0,46 | 0,842 | 0,911 | 8,91E-244 |
| Ahnak | 1,39E-109 | -0,46 | 0,883 | 0,842 | 2,58E-105 |
| Zscan26 | 2,46E-158 | -0,46 | 0,269 | 0,429 | 4,58E-154 |
| Ccnl1 | 8,49E-218 | -0,46 | 0,767 | 0,836 | 1,58E-213 |
| Rasa3 | 1,10E-139 | -0,46 | 0,619 | 0,709 | 2,05E-135 |
| Itpkb | 1,65E-181 | -0,46 | 0,633 | 0,729 | 3,07E-177 |
| Nr3c1 | 9,02E-167 | -0,46 | 0,469 | 0,599 | 1,68E-162 |

| | | | | | |
|---|---|---|---|---|---|
| Qpct | 9,27E-248 | -0,46 | 0,07 | 0,241 | 1,72E-243 |
| Arhgap45 | 0 | -0,46 | 0,959 | 0,982 | 0 |
| Add3 | 3,05E-166 | -0,47 | 0,672 | 0,756 | 5,66E-162 |
| Txk | 2,80E-30 | -0,47 | 0,612 | 0,599 | 5,20E-26 |
| Smpdl3a | 1,49E-151 | -0,47 | 0,486 | 0,605 | 2,77E-147 |
| Pde7a | 4,71E-188 | -0,47 | 0,673 | 0,758 | 8,75E-184 |
| Dgkd | 6,15E-196 | -0,47 | 0,658 | 0,744 | 1,14E-191 |
| Ablim1 | 2,67E-270 | -0,47 | 0,909 | 0,951 | 4,97E-266 |
| Igflr1 | 1,06E-121 | -0,47 | 0,46 | 0,571 | 1,97E-117 |
| Pnisr | 9,29E-209 | -0,48 | 0,639 | 0,735 | 1,73E-204 |
| Cbx7 | 1,12E-178 | -0,48 | 0,375 | 0,54 | 2,09E-174 |
| Ikzf2 | 6,44E-46 | -0,48 | 0,216 | 0,288 | 1,20E-41 |
| H2-Ab1 | 9,90E-25 | -0,49 | 0,061 | 0,1 | 1,84E-20 |
| Selenop | 6,67E-157 | -0,49 | 0,152 | 0,306 | 1,24E-152 |
| St6gal1 | 2,26E-306 | -0,49 | 0,032 | 0,201 | 4,19E-302 |
| Pik3cd | 0 | -0,49 | 0,92 | 0,957 | 0 |
| Gtf2i | 3,67E-235 | -0,50 | 0,732 | 0,819 | 6,81E-231 |
| Lrig1 | 3,27E-290 | -0,50 | 0,121 | 0,329 | 6,07E-286 |
| Ypel3 | 3,85E-252 | -0,50 | 0,771 | 0,854 | 7,16E-248 |
| Rsrp1 | 9,60E-236 | -0,50 | 0,856 | 0,911 | 1,78E-231 |
| Dnajb1 | 5,70E-62 | -0,50 | 0,566 | 0,598 | 1,06E-57 |
| Il4ra | 1,99E-59 | -0,50 | 0,487 | 0,533 | 3,71E-55 |
| Zfp36l2 | 5,84E-159 | -0,50 | 0,92 | 0,952 | 1,09E-154 |
| Fam78a | 5,43E-232 | -0,51 | 0,793 | 0,863 | 1,01E-227 |
| Saraf | 0 | -0,51 | 0,932 | 0,958 | 0 |
| Mxd4 | 1,20E-144 | -0,51 | 0,468 | 0,589 | 2,24E-140 |
| Srsf5 | 5,93E-297 | -0,51 | 0,805 | 0,869 | 1,10E-292 |
| Tgfbr2 | 0 | -0,51 | 0,907 | 0,953 | 0 |
| Tmem181a | 6,34E-304 | -0,52 | 0,182 | 0,402 | 1,18E-299 |
| Rgs10 | 5,45E-183 | -0,52 | 0,408 | 0,582 | 1,01E-178 |
| Prkd2 | 7,33E-244 | -0,52 | 0,68 | 0,788 | 1,36E-239 |
| Sgms1 | 3,14E-198 | -0,52 | 0,241 | 0,418 | 5,83E-194 |
| Clk1 | 1,02E-281 | -0,52 | 0,904 | 0,939 | 1,90E-277 |
| Tcp11l2 | 1,29E-136 | -0,52 | 0,546 | 0,65 | 2,39E-132 |
| Fam241a | 4,76E-53 | -0,52 | 0,547 | 0,58 | 8,85E-49 |
| Mbp | 1,42E-267 | -0,52 | 0,593 | 0,736 | 2,64E-263 |
| Rab3ip | 2,15E-72 | -0,53 | 0,356 | 0,428 | 3,99E-68 |
| Rapgef6 | 1,22E-237 | -0,53 | 0,856 | 0,898 | 2,27E-233 |
| Tdrp | 5,10E-253 | -0,53 | 0,047 | 0,205 | 9,48E-249 |
| Inpp4b | 9,17E-177 | -0,54 | 0,685 | 0,778 | 1,70E-172 |
| Fosb | 1,48E-18 | -0,54 | 0,363 | 0,396 | 2,76E-14 |
| Dph5 | 2,20E-127 | -0,54 | 0,242 | 0,371 | 4,09E-123 |
| Thada | 9,07E-178 | -0,54 | 0,582 | 0,692 | 1,68E-173 |
| Ptger4 | 5,18E-88 | -0,54 | 0,52 | 0,589 | 9,63E-84 |
| Faah | 1,45E-221 | -0,55 | 0,302 | 0,482 | 2,69E-217 |
| Rasgrp2 | 6,28E-119 | -0,55 | 0,372 | 0,496 | 1,17E-114 |

| | | | | | |
|---|---|---|---|---|---|
| Itgad | 1,85E-132 | -0,56 | 0,038 | 0,135 | 3,44E-128 |
| Ogt | 0 | -0,56 | 0,838 | 0,906 | 0 |
| Gpr183 | 3,78E-138 | -0,56 | 0,521 | 0,64 | 7,02E-134 |
| Kif1b | 3,86E-222 | -0,56 | 0,294 | 0,479 | 7,17E-218 |
| Itgb7 | 0 | -0,57 | 0,895 | 0,949 | 0 |
| Sik1 | 8,33E-92 | -0,57 | 0,515 | 0,586 | 1,55E-87 |
| Ripor2 | 1,92E-222 | -0,57 | 0,66 | 0,773 | 3,56E-218 |
| St8sia4 | 2,82E-203 | -0,57 | 0,369 | 0,54 | 5,24E-199 |
| Numa1 | 0 | -0,57 | 0,785 | 0,877 | 0 |
| Tspan13 | 5,58E-271 | -0,57 | 0,392 | 0,593 | 1,04E-266 |
| Samd3 | 2,34E-192 | -0,57 | 0,225 | 0,406 | 4,35E-188 |
| Sh3bp5 | 6,00E-232 | -0,57 | 0,106 | 0,284 | 1,12E-227 |
| Bcl11b | 7,47E-297 | -0,58 | 0,818 | 0,88 | 1,39E-292 |
| Macf1 | 0 | -0,58 | 0,899 | 0,944 | 0 |
| Smad7 | 1,77E-223 | -0,58 | 0,665 | 0,767 | 3,29E-219 |
| Dusp1 | 6,12E-113 | -0,59 | 0,641 | 0,728 | 1,14E-108 |
| Ifngr2 | 8,19E-217 | -0,59 | 0,028 | 0,156 | 1,52E-212 |
| Tcrg-C4 | 4,03E-181 | -0,59 | 0,05 | 0,179 | 7,50E-177 |
| Il6ra | 1,47E-285 | -0,60 | 0,044 | 0,218 | 2,73E-281 |
| Stk38 | 2,58E-264 | -0,60 | 0,658 | 0,764 | 4,80E-260 |
| Tcrg-C2 | 9,84E-87 | -0,61 | 0,177 | 0,284 | 1,83E-82 |
| Bach2 | 1,37E-265 | -0,61 | 0,147 | 0,354 | 2,54E-261 |
| Pde2a | 8,69E-91 | -0,62 | 0,225 | 0,338 | 1,62E-86 |
| Pde3b | 1,39E-286 | -0,62 | 0,683 | 0,787 | 2,59E-282 |
| Gpr132 | 2,25E-213 | -0,62 | 0,665 | 0,764 | 4,19E-209 |
| Dnaja1 | 0 | -0,62 | 0,699 | 0,791 | 0 |
| Kctd12 | 2,76E-252 | -0,63 | 0,182 | 0,389 | 5,13E-248 |
| Klf3 | 1,05E-142 | -0,63 | 0,257 | 0,411 | 1,95E-138 |
| Eomes | 2,46E-103 | -0,63 | 0,242 | 0,362 | 4,57E-99 |
| Il27ra | 0 | -0,63 | 0,657 | 0,796 | 0 |
| Spry2 | 2,79E-60 | -0,63 | 0,157 | 0,243 | 5,18E-56 |
| Gramd3 | 2,20E-119 | -0,64 | 0,866 | 0,847 | 4,09E-115 |
| Sesn3 | 0 | -0,64 | 0,568 | 0,747 | 0 |
| Sun2 | 0 | -0,65 | 0,756 | 0,872 | 0 |
| Arhgef18 | 6,15E-280 | -0,65 | 0,704 | 0,839 | 1,14E-275 |
| Nsg2 | 6,64E-308 | -0,66 | 0,073 | 0,275 | 1,23E-303 |
| Foxp1 | 5,76E-265 | -0,66 | 0,596 | 0,724 | 1,07E-260 |
| Tagap | 1,71E-253 | -0,66 | 0,68 | 0,819 | 3,18E-249 |
| Rgcc | 4,34E-128 | -0,66 | 0,172 | 0,305 | 8,06E-124 |
| Klre1 | 7,40E-26 | -0,67 | 0,22 | 0,266 | 1,38E-21 |
| Rgs2 | 3,46E-119 | -0,67 | 0,318 | 0,452 | 6,43E-115 |
| Ddx5 | 0 | -0,68 | 0,902 | 0,951 | 0 |
| Dapl1 | 6,59E-192 | -0,68 | 0,022 | 0,134 | 1,23E-187 |
| Sntb1 | 0 | -0,69 | 0,295 | 0,559 | 0 |
| Dusp10 | 5,00E-67 | -0,70 | 0,461 | 0,514 | 9,29E-63 |
| Itgb1 | 2,24E-127 | -0,71 | 0,649 | 0,707 | 4,17E-123 |

| | | | | | |
|---|---|---|---|---|---|
| Jmy | 0 | -0,72 | 0,294 | 0,516 | 0 |
| Cd7 | 1,67E-153 | -0,72 | 0,352 | 0,518 | 3,11E-149 |
| Ifi27 | 0 | -0,72 | 0,585 | 0,806 | 0 |
| Ldlrad4 | 1,11E-307 | -0,72 | 0,327 | 0,549 | 2,06E-303 |
| Itga1 | 1,54E-112 | -0,74 | 0,387 | 0,483 | 2,87E-108 |
| P2ry10 | 0 | -0,75 | 0,775 | 0,872 | 0 |
| Cdh1 | 4,24E-268 | -0,75 | 0,155 | 0,363 | 7,88E-264 |
| Evl | 0 | -0,75 | 0,636 | 0,844 | 0 |
| Hsph1 | 0 | -0,77 | 0,391 | 0,593 | 0 |
| Ier5l | 4,72E-82 | -0,77 | 0,203 | 0,304 | 8,77E-78 |
| Chd3 | 0 | -0,78 | 0,585 | 0,766 | 0 |
| Rhob | 4,84E-198 | -0,83 | 0,218 | 0,39 | 9,00E-194 |
| Dgka | 0 | -0,84 | 0,829 | 0,885 | 0 |
| Slc12a7 | 0 | -0,85 | 0,469 | 0,693 | 0 |
| Klf6 | 0 | -0,86 | 0,89 | 0,946 | 0 |
| Cd74 | 7,30E-90 | -0,86 | 0,132 | 0,241 | 1,36E-85 |
| Fos | 1,14E-132 | -0,87 | 0,457 | 0,577 | 2,12E-128 |
| Igfbp4 | 2,75E-228 | -0,88 | 0,053 | 0,209 | 5,10E-224 |
| Eif4a2 | 0 | -0,90 | 0,878 | 0,958 | 0 |
| Cmah | 0 | -0,91 | 0,13 | 0,383 | 0 |
| Neurl3 | 7,20E-277 | -0,92 | 0,478 | 0,657 | 1,34E-272 |
| Lef1 | 8,54E-94 | -0,92 | 0,49 | 0,531 | 1,59E-89 |
| Il6st | 1,40E-201 | -0,96 | 0,453 | 0,571 | 2,59E-197 |
| Hspa1a | 7,40E-219 | -1,01 | 0,012 | 0,123 | 1,38E-214 |
| Vps37b | 7,45E-253 | -1,01 | 0,83 | 0,862 | 1,38E-248 |
| Actn1 | 0 | -1,06 | 0,256 | 0,52 | 0 |
| Jun | 0 | -1,19 | 0,584 | 0,741 | 0 |
| Tcf7 | 0 | -1,23 | 0,287 | 0,633 | 0 |
| S1pr1 | 0 | -1,25 | 0,283 | 0,502 | 0 |
| Ighm | 0 | -1,32 | 0,095 | 0,353 | 0 |
| Itgae | 0 | -1,32 | 0,185 | 0,43 | 0 |
| Sell | 3,19E-274 | -1,34 | 0,211 | 0,427 | 5,92E-270 |
| Trdc | 1,41E-128 | -1,38 | 0,036 | 0,13 | 2,62E-124 |
| Hspa1b | 0 | -1,45 | 0,021 | 0,2 | 0 |
| Klf2 | 0 | -1,54 | 0,441 | 0,631 | 0 |
| Ccr7 | 0 | -1,82 | 0,095 | 0,374 | 0 |

### Table S5

Cluster 0

| | p_val | avg_log2FC | pct.1 | pct.2 | p_val_adj |
|---|---|---|---|---|---|
| Ifit3 | 8.359210797 | 2,30196596 | 0,731 | 0,15 | 1,55E-302 |
| Gzmb | 6.379583909 | 2,12348227 | 0,911 | 0,389 | 1,19E-241 |
| Ifi27l2a | 3.295351082 | 1,62620285 | 0,825 | 0,572 | 6,12E-142 |
| Isg15 | 8.294509466 | 1,55095138 | 0,622 | 0,217 | 1,54E-162 |
| Rtp4 | 3.090855912 | 1,42142982 | 0,758 | 0,299 | 5,74E-216 |
| Ifit1 | 1.060410259 | 1,38934589 | 0,499 | 0,084 | 1,97E-170 |
| Plac8 | 6.621921374 | 1,35124621 | 0,905 | 0,688 | 1,23E-141 |
| Ifi208 | 7.691011369 | 1,24978255 | 0,655 | 0,172 | 1,43E-217 |
| Zbp1 | 3.964665414 | 1,176173 | 0,933 | 0,776 | 7,37E-179 |
| Rnf213 | 1.505260547 | 1,17466313 | 0,88 | 0,753 | 2,80E-112 |
| Usp18 | 1.534197769 | 1,14701584 | 0,515 | 0,142 | 2,85E-133 |
| Bst2 | 2.427093552 | 1,14437318 | 0,847 | 0,677 | 4,51E-113 |
| AA467197 | 2.656649885 | 1,07897853 | 0,53 | 0,231 | 4,94E-84 |
| Ifit1bl1 | 3.081544393 | 1,0203866 | 0,832 | 0,61 | 5,73E-110 |
| Rbm3 | 2.406372940 | 0,98539571 | 0,909 | 0,7 | 4,47E-196 |
| Xaf1 | 2.619680303 | 0,91436698 | 0,753 | 0,464 | 4,87E-119 |
| Irf7 | 5.199219931 | 0,91357209 | 0,772 | 0,483 | 9,66E-108 |
| Ly6a | 4.587701271 | 0,91237456 | 0,986 | 0,903 | 8,53E-148 |
| Oas3 | 1.898227552 | 0,85366742 | 0,595 | 0,327 | 3,53E-77 |
| Slfn1 | 3.425571861 | 0,84860245 | 0,825 | 0,616 | 6,37E-97 |
| Bcl2l11 | 3.670639231 | 0,7758117 | 0,642 | 0,457 | 6,82E-40 |
| Dhx58 | 1.080837550 | 0,77006702 | 0,478 | 0,201 | 2,01E-77 |
| Ccl3 | 8.771047954 | 0,74851971 | 0,387 | 0,166 | 1,63E-44 |
| Ms4a4c | 2.022222118 | 0,7342783 | 0,353 | 0,149 | 3,76E-48 |
| Parp9 | 1.217235370 | 0,73132817 | 0,737 | 0,52 | 2,26E-77 |
| Phf11c | 8.267303310 | 0,71957883 | 0,605 | 0,355 | 1,54E-69 |
| Lgals3bp | 6.611540389 | 0,70820242 | 0,917 | 0,848 | 1,23E-87 |
| Serpinb9 | 6.942522628 | 0,70711451 | 0,61 | 0,428 | 1,29E-38 |
| Isg20 | 3.580253054 | 0,69554677 | 0,68 | 0,537 | 6,65E-36 |
| Phf11b | 4.448330196 | 0,69166711 | 0,807 | 0,633 | 8,27E-74 |
| Id2 | 3.592436526 | 0,68271481 | 0,971 | 0,929 | 6,68E-72 |
| Pim1 | 1.025574395 | 0,68067491 | 0,794 | 0,63 | 1,91E-43 |
| Ifi209 | 1.234815208 | 0,66888656 | 0,891 | 0,781 | 2,29E-64 |
| Pik3ap1 | 2.510546179 | 0,66707281 | 0,716 | 0,537 | 4,67E-48 |
| Oas1a | 8.923360268 | 0,66635482 | 0,397 | 0,148 | 1,66E-65 |
| Gbp9 | 2.094750354 | 0,6622305 | 0,74 | 0,55 | 3,89E-59 |
| Il18rap | 2.408169070 | 0,66062255 | 0,893 | 0,764 | 4,48E-60 |
| Serpina3g | 4.186589858 | 0,66046858 | 0,765 | 0,67 | 7,78E-26 |
| Bcl2 | 3.288513791 | 0,65582491 | 0,567 | 0,384 | 6,11E-32 |
| Daxx | 4.461449508 | 0,63972294 | 0,635 | 0,462 | 8,29E-39 |
| Ifih1 | 4.438261775 | 0,63118446 | 0,418 | 0,167 | 8,25E-65 |
| Gbp2 | 3.025048420 | 0,62520426 | 0,82 | 0,708 | 5,62E-40 |
| Ifi206 | 2.756644918 | 0,61579065 | 0,808 | 0,664 | 5,12E-60 |
| Dtx3l | 1.637662316 | 0,5848048 | 0,855 | 0,77 | 3,04E-59 |
| Rab27a | 1.417615398 | 0,5756761 | 0,822 | 0,677 | 2,63E-58 |

Up in Sp140-/-

Up in Sp140+/+

| | | | | | |
|---|---|---|---|---|---|
| Il2ra | 1.115510929 | 0,57358777 | 0,345 | 0,193 | 2,07E-23 |
| Arsb | 3.203447859 | 0,57120043 | 0,748 | 0,579 | 5,95E-49 |
| Fkbp5 | 2.561845498 | 0,5669972 | 0,49 | 0,259 | 4,76E-48 |
| Cish | 2.160911615 | 0,55865508 | 0,632 | 0,453 | 4,02E-33 |
| Cd86 | 5.830627576 | 0,55250284 | 0,465 | 0,28 | 1,08E-33 |
| Ifit2 | 2.233221836 | 0,55246411 | 0,449 | 0,228 | 4,15E-47 |
| H2-T24 | 7.306644145 | 0,54791612 | 0,498 | 0,283 | 1,36E-40 |
| Irgm1 | 2.271677049 | 0,54765977 | 0,781 | 0,637 | 4,22E-43 |
| Ifit3b | 1.856528399 | 0,54695232 | 0,257 | 0,018 | 3,45E-92 |
| Slfn8 | 1.974340339 | 0,54358532 | 0,651 | 0,453 | 3,67E-45 |
| Gbp4 | 1.379127805 | 0,53508518 | 0,845 | 0,748 | 2,56E-35 |
| Gbp6 | 1.145494004 | 0,52933571 | 0,681 | 0,513 | 2,13E-37 |
| Nfkbia | 3.146299011 | 0,52624799 | 0,808 | 0,696 | 5,85E-28 |
| Ide | 2.984842149 | 0,52586157 | 0,757 | 0,6 | 5,55E-54 |
| Ifi47 | 1.327915669 | 0,5213982 | 0,929 | 0,853 | 2,47E-49 |
| Gzma | 1.226488276 | 0,51623106 | 0,665 | 0,274 | 2,28E-89 |
| Trafd1 | 2.897096325 | 0,51262704 | 0,652 | 0,525 | 5,38E-29 |
| Herc6 | 1.534472531 | 0,51192405 | 0,609 | 0,462 | 2,85E-27 |
| Gadd45b | 2.093506436 | 0,50244491 | 0,453 | 0,36 | 3,89E-09 |
| Ddx5 | 5.656024631 | -0,5039387 | 0,897 | 0,95 | 1,05E-77 |
| Sntb1 | 2.377393812 | -0,515582 | 0,346 | 0,564 | 4,42E-49 |
| Ptger4 | 7.494295205 | -0,5205411 | 0,566 | 0,688 | 1,39E-23 |
| Dnaja1 | 5.414409358 | -0,5331034 | 0,646 | 0,784 | 1,01E-44 |
| Hist1h1e | 4.652598328 | -0,534646 | 0,634 | 0,687 | 8,65E-12 |
| Vps37b | 9.703424742 | -0,5524949 | 0,735 | 0,794 | 1,80E-17 |
| Fosb | 1.638402339 | -0,557475 | 0,298 | 0,411 | 3,04E-11 |
| Gpr183 | 1.576155447 | -0,5587314 | 0,482 | 0,62 | 2,93E-24 |
| Actn1 | 1.636842206 | -0,5733772 | 0,216 | 0,391 | 3,04E-34 |
| Ldlrad4 | 5.278990501 | -0,5847816 | 0,256 | 0,472 | 9,81E-46 |
| Rgs2 | 1.679368704 | -0,5849529 | 0,255 | 0,381 | 3,12E-15 |
| Chd3 | 2.554879015 | -0,5892751 | 0,55 | 0,741 | 4,75E-57 |
| Rbpj | 7.778148216 | -0,597163 | 0,514 | 0,652 | 1,45E-28 |
| Fos | 2.872836462 | -0,5990056 | 0,347 | 0,448 | 5,34E-10 |
| Sgms1 | 7.826864592 | -0,6006299 | 0,251 | 0,492 | 1,45E-58 |
| Itgb1 | 7.317695479 | -0,6049563 | 0,79 | 0,87 | 1,36E-28 |
| Cxcr6 | 8.801200608 | -0,605254 | 0,794 | 0,846 | 1,64E-27 |
| Ppp2r2c | 7.347050898 | -0,6084719 | 0,184 | 0,405 | 1,37E-48 |
| Itga1 | 1.207596144 | -0,6277222 | 0,36 | 0,505 | 2,24E-23 |
| Eif4a2 | 3.362194930 | -0,628116 | 0,857 | 0,943 | 6,25E-93 |
| Cdh1 | 3.572518292 | -0,6286916 | 0,152 | 0,349 | 6,64E-41 |
| Dusp1 | 5.198757563 | -0,6298237 | 0,57 | 0,682 | 9,66E-16 |
| Myadm | 6.625106006 | -0,6388281 | 0,247 | 0,424 | 1,23E-32 |
| Eomes | 7.385716900 | -0,6738132 | 0,249 | 0,394 | 1,37E-22 |
| Ifi27 | 1.896931550 | -0,7015123 | 0,585 | 0,83 | 3,53E-91 |
| Hsph1 | 2.789472271 | -0,7104807 | 0,339 | 0,561 | 5,18E-59 |
| Tcf7 | 2.609020256 | -0,7445652 | 0,402 | 0,64 | 4,85E-53 |
| Rhob | 1.909399389 | -0,8230993 | 0,149 | 0,337 | 3,55E-42 |
| Ier5l | 6.874062502 | -0,8303033 | 0,155 | 0,289 | 1,28E-21 |

| | | | | | |
|---|---|---|---|---|---|
| Klf6 | 1.551305110 | -0,8874783 | 0,882 | 0,953 | 2,88E-106 |
| Itgae | 1.759391136 | -1,0726746 | 0,146 | 0,34 | 3,27E-41 |
| Jun | 3.288865001 | -1,286883 | 0,438 | 0,661 | 6,11E-62 |

Cluster 1

| | p_val | avg_log2FC | pct.1 | pct.2 | p_val_adj |
|---|---|---|---|---|---|
| Gzmb | 2,25E-282 | 2,91584746 | 0,99 | 0,632 | 4,17E-278 |
| Ccl3 | 7,58E-126 | 2,68539027 | 0,755 | 0,322 | 1,41E-121 |
| Ifi27l2a | 4,18E-226 | 2,58405822 | 0,91 | 0,568 | 7,77E-222 |
| Ifit3 | 3,10E-257 | 2,5464408 | 0,81 | 0,166 | 5,76E-253 |
| Isg15 | 3,02E-267 | 2,51024267 | 0,917 | 0,395 | 5,62E-263 |
| Gzma | 1,30E-141 | 2,36043015 | 0,778 | 0,263 | 2,41E-137 |
| Usp18 | 6,30E-304 | 2,31308996 | 0,896 | 0,235 | 1,17E-299 |
| Gzmc | 7,24E-59 | 2,23229891 | 0,284 | 0,038 | 1,35E-54 |
| Tnfrsf9 | 6,60E-161 | 2,18405726 | 0,692 | 0,183 | 1,23E-156 |
| Ifitm3 | 5,62E-142 | 2,16799521 | 0,543 | 0,075 | 1,05E-137 |
| Ifitm2 | 1,81E-79 | 2,15869042 | 0,398 | 0,077 | 3,37E-75 |
| Ifitm1 | 2,76E-54 | 1,89319943 | 0,315 | 0,064 | 5,14E-50 |
| Prf1 | 1,80E-163 | 1,8678773 | 0,986 | 0,909 | 3,34E-159 |
| AA467197 | 1,57E-169 | 1,7717627 | 0,949 | 0,752 | 2,91E-165 |
| Ms4a4c | 9,11E-189 | 1,74427621 | 0,712 | 0,161 | 1,69E-184 |
| Ifit1bl1 | 7,37E-161 | 1,68328628 | 0,905 | 0,664 | 1,37E-156 |
| Bst2 | 5,34E-198 | 1,62597822 | 0,964 | 0,819 | 9,91E-194 |
| Rnf213 | 4,84E-168 | 1,56308539 | 0,954 | 0,831 | 9,00E-164 |
| Tnfrsf18 | 1,34E-182 | 1,55574668 | 0,919 | 0,596 | 2,50E-178 |
| Ifit1 | 4,60E-172 | 1,51751244 | 0,716 | 0,179 | 8,54E-168 |
| Rtp4 | 4,74E-208 | 1,47017257 | 0,878 | 0,389 | 8,81E-204 |
| Irf8 | 9,66E-159 | 1,43877398 | 0,779 | 0,273 | 1,80E-154 |
| Irf7 | 3,67E-178 | 1,4295951 | 0,906 | 0,55 | 6,81E-174 |
| Oas3 | 1,66E-165 | 1,42234323 | 0,823 | 0,415 | 3,08E-161 |
| Mxd1 | 2,65E-147 | 1,38742182 | 0,956 | 0,729 | 4,92E-143 |
| Ifi211 | 9,86E-178 | 1,35956828 | 0,66 | 0,117 | 1,83E-173 |
| Tnfrsf4 | 4,06E-124 | 1,34252888 | 0,489 | 0,06 | 7,54E-120 |
| Pik3ap1 | 8,44E-187 | 1,33856375 | 0,944 | 0,653 | 1,57E-182 |
| Acadl | 3,84E-145 | 1,31757941 | 0,951 | 0,816 | 7,13E-141 |
| Lag3 | 1,16E-123 | 1,29532606 | 0,97 | 0,793 | 2,16E-119 |
| Dhx58 | 2,06E-185 | 1,28179827 | 0,81 | 0,298 | 3,83E-181 |
| Oas1a | 5,09E-166 | 1,24200719 | 0,713 | 0,205 | 9,45E-162 |
| Rsad2 | 1,50E-153 | 1,22434693 | 0,525 | 0,034 | 2,78E-149 |
| Isg20 | 2,57E-152 | 1,18954338 | 0,969 | 0,825 | 4,78E-148 |
| Ifit2 | 4,59E-110 | 1,17870004 | 0,694 | 0,304 | 8,53E-106 |
| Xaf1 | 4,95E-176 | 1,17836216 | 0,92 | 0,61 | 9,19E-172 |
| Slfn1 | 1,13E-164 | 1,17460463 | 0,938 | 0,699 | 2,10E-160 |
| Sema7a | 1,04E-149 | 1,1711565 | 0,657 | 0,146 | 1,93E-145 |
| Icos | 1,47E-122 | 1,14530992 | 0,989 | 0,889 | 2,73E-118 |
| Havcr2 | 5,61E-109 | 1,13094911 | 0,84 | 0,535 | 1,04E-104 |
| Il2ra | 2,52E-51 | 1,12651496 | 0,719 | 0,491 | 4,67E-47 |
| Daxx | 1,77E-112 | 1,10310428 | 0,835 | 0,55 | 3,29E-108 |
| Pkm | 6,53E-203 | 1,09329396 | 0,993 | 0,911 | 1,21E-198 |
| Phf11b | 2,80E-153 | 1,0854493 | 0,954 | 0,764 | 5,20E-149 |
| Trafd1 | 7,98E-156 | 1,08461783 | 0,924 | 0,647 | 1,48E-151 |
| Casp3 | 6,21E-152 | 1,08337478 | 0,913 | 0,603 | 1,15E-147 |
| Plac8 | 9,88E-99 | 1,08021081 | 0,979 | 0,898 | 1,84E-94 |
| Ifi208 | 7,28E-159 | 1,07892245 | 0,677 | 0,181 | 1,35E-154 |
| Zbp1 | 1,86E-163 | 1,05098925 | 0,991 | 0,92 | 3,46E-159 |
| Entpd1 | 1,63E-138 | 1,04577843 | 0,898 | 0,621 | 3,02E-134 |

Up in Sp140-/-

Up in Sp140+/+

| | | | | | |
|---|---|---|---|---|---|
| Gstt1 | 2,97E-39 | 1,01619592 | 0,453 | 0,224 | 5,51E-35 |
| Ifih1 | 1,59E-134 | 1,00285275 | 0,642 | 0,171 | 2,96E-130 |
| Ifi209 | 6,27E-133 | 1,00246963 | 0,968 | 0,859 | 1,17E-128 |
| Hif1a | 4,83E-119 | 0,99660233 | 0,966 | 0,809 | 8,97E-115 |
| Herc6 | 3,13E-106 | 0,99026314 | 0,839 | 0,571 | 5,82E-102 |
| Nt5c3 | 6,09E-83 | 0,98935742 | 0,756 | 0,465 | 1,13E-78 |
| Sv2c | 1,47E-88 | 0,9842195 | 0,447 | 0,09 | 2,73E-84 |
| Tpi1 | 7,07E-88 | 0,98240382 | 0,667 | 0,305 | 1,31E-83 |
| Cdkn1a | 1,26E-69 | 0,9812027 | 0,454 | 0,134 | 2,34E-65 |
| Phf11c | 2,06E-122 | 0,9698867 | 0,796 | 0,417 | 3,84E-118 |
| Hlx | 7,55E-82 | 0,96023271 | 0,649 | 0,292 | 1,40E-77 |
| Slc7a5 | 2,33E-80 | 0,94097809 | 0,567 | 0,202 | 4,33E-76 |
| Chchd10 | 2,02E-66 | 0,93561809 | 0,526 | 0,217 | 3,75E-62 |
| Helz2 | 2,27E-123 | 0,93182868 | 0,952 | 0,818 | 4,22E-119 |
| Sdc3 | 1,43E-130 | 0,92818366 | 0,451 | 0,02 | 2,65E-126 |
| Ccl4 | 1,97E-23 | 0,8961773 | 0,945 | 0,894 | 3,66E-19 |
| Pml | 1,83E-117 | 0,89296567 | 0,844 | 0,514 | 3,40E-113 |
| Zbtb32 | 8,07E-123 | 0,88702761 | 0,457 | 0,038 | 1,50E-118 |
| Mapkapk2 | 1,05E-138 | 0,87858651 | 0,981 | 0,876 | 1,95E-134 |
| Rilpl2 | 2,50E-86 | 0,87186587 | 0,787 | 0,471 | 4,64E-82 |
| Hk2 | 3,03E-85 | 0,85491984 | 0,528 | 0,16 | 5,64E-81 |
| Sytl3 | 2,82E-116 | 0,8447899 | 0,872 | 0,531 | 5,24E-112 |
| Pdcd1 | 6,73E-72 | 0,84218916 | 0,755 | 0,42 | 1,25E-67 |
| Trim30d | 5,71E-124 | 0,839722 | 0,719 | 0,268 | 1,06E-119 |
| Trim30a | 4,61E-112 | 0,83567371 | 0,86 | 0,579 | 8,56E-108 |
| Gnptab | 1,01E-108 | 0,82894959 | 0,851 | 0,537 | 1,88E-104 |
| Il10ra | 1,97E-88 | 0,82582419 | 0,817 | 0,506 | 3,67E-84 |
| Ifng | 1,63E-22 | 0,82359475 | 0,966 | 0,939 | 3,03E-18 |
| Lgals3bp | 8,90E-126 | 0,8082187 | 0,991 | 0,979 | 1,65E-121 |
| Cblb | 4,50E-75 | 0,80746574 | 0,899 | 0,745 | 8,35E-71 |
| Serpina3f | 1,77E-62 | 0,79796954 | 0,47 | 0,165 | 3,29E-58 |
| Chd7 | 1,87E-95 | 0,79758091 | 0,912 | 0,696 | 3,48E-91 |
| Ddt | 3,79E-39 | 0,79495957 | 0,753 | 0,556 | 7,05E-35 |
| Aldoa | 8,41E-98 | 0,79444538 | 0,978 | 0,906 | 1,56E-93 |
| Id2 | 1,79E-123 | 0,78845274 | 0,999 | 0,994 | 3,32E-119 |
| Cmpk2 | 8,44E-103 | 0,78483504 | 0,443 | 0,062 | 1,57E-98 |
| Slc16a3 | 3,03E-70 | 0,7809307 | 0,483 | 0,162 | 5,63E-66 |
| Mif4gd | 1,41E-107 | 0,78048798 | 0,958 | 0,783 | 2,62E-103 |
| Tg | 9,78E-75 | 0,78048074 | 0,332 | 0,037 | 1,82E-70 |
| Rbm3 | 2,59E-134 | 0,77525711 | 0,955 | 0,81 | 4,82E-130 |
| Parp9 | 4,31E-99 | 0,77309944 | 0,901 | 0,67 | 8,01E-95 |
| Slfn8 | 1,04E-102 | 0,77222743 | 0,881 | 0,61 | 1,94E-98 |
| Satb1 | 9,36E-73 | 0,76525464 | 0,959 | 0,778 | 1,74E-68 |
| Slc25a19 | 4,63E-70 | 0,76190989 | 0,804 | 0,548 | 8,60E-66 |
| Cd27 | 2,34E-70 | 0,76142871 | 0,801 | 0,528 | 4,35E-66 |
| Pim1 | 9,48E-84 | 0,75982887 | 0,964 | 0,788 | 1,76E-79 |
| Phf11a | 6,26E-108 | 0,74616761 | 0,64 | 0,216 | 1,16E-103 |
| Naa20 | 2,13E-88 | 0,74206799 | 0,856 | 0,587 | 3,95E-84 |
| Ccr2 | 4,05E-36 | 0,74095815 | 0,684 | 0,439 | 7,53E-32 |
| Smpdl3b | 1,81E-87 | 0,73637462 | 0,614 | 0,221 | 3,36E-83 |
| Il18rap | 2,16E-55 | 0,72412603 | 0,947 | 0,827 | 4,01E-51 |
| Serpinb9 | 4,68E-45 | 0,72364135 | 0,714 | 0,475 | 8,69E-41 |
| Glrx | 1,39E-64 | 0,719111 | 0,963 | 0,941 | 2,59E-60 |

| | | | | | |
|---|---|---|---|---|---|
| Slc2a1 | 1,17E-63 | 0,71507531 | 0,716 | 0,412 | 2,18E-59 |
| Setbp1 | 7,47E-79 | 0,71039133 | 0,763 | 0,427 | 1,39E-74 |
| Wdfy1 | 3,92E-111 | 0,70721649 | 0,769 | 0,356 | 7,29E-107 |
| Sub1 | 4,87E-88 | 0,7039865 | 0,993 | 0,961 | 9,06E-84 |
| Ddx60 | 2,40E-88 | 0,70063204 | 0,558 | 0,184 | 4,46E-84 |
| Crispld2 | 3,37E-23 | 0,69617361 | 0,392 | 0,227 | 6,25E-19 |
| Sco1 | 4,25E-93 | 0,68414895 | 0,53 | 0,144 | 7,90E-89 |
| Il12rb1 | 2,88E-66 | 0,67355498 | 0,92 | 0,757 | 5,36E-62 |
| Cd53 | 1,74E-111 | 0,66612253 | 0,999 | 0,994 | 3,24E-107 |
| Eif2ak2 | 8,56E-81 | 0,6640232 | 0,683 | 0,337 | 1,59E-76 |
| Ifi214 | 1,85E-80 | 0,66380336 | 0,763 | 0,43 | 3,43E-76 |
| Sdcbp2 | 2,43E-68 | 0,65765852 | 0,618 | 0,289 | 4,51E-64 |
| Ifit3b | 1,75E-102 | 0,65663576 | 0,391 | 0,026 | 3,25E-98 |
| Oas2 | 1,17E-89 | 0,65659032 | 0,328 | 0,011 | 2,17E-85 |
| Chmp4b | 9,63E-110 | 0,65433068 | 0,951 | 0,781 | 1,79E-105 |
| Plek | 1,13E-58 | 0,65169124 | 0,863 | 0,634 | 2,10E-54 |
| H2-T24 | 3,12E-73 | 0,64571311 | 0,637 | 0,283 | 5,80E-69 |
| Trim25 | 1,24E-71 | 0,63919107 | 0,788 | 0,486 | 2,30E-67 |
| Glipr2 | 3,10E-77 | 0,6390248 | 0,952 | 0,828 | 5,76E-73 |
| Ccrl2 | 2,51E-89 | 0,63575955 | 0,379 | 0,041 | 4,66E-85 |
| Rhbdf2 | 8,04E-72 | 0,6345085 | 0,77 | 0,482 | 1,49E-67 |
| Prelid1 | 8,89E-91 | 0,63197839 | 0,969 | 0,902 | 1,65E-86 |
| Gem | 8,33E-32 | 0,62934304 | 0,81 | 0,648 | 1,55E-27 |
| Frmd4a | 1,81E-63 | 0,62546315 | 0,466 | 0,163 | 3,37E-59 |
| Peli1 | 4,03E-67 | 0,6224511 | 0,961 | 0,857 | 7,49E-63 |
| Anxa2 | 5,81E-49 | 0,6198678 | 0,964 | 0,936 | 1,08E-44 |
| Dgat1 | 1,60E-60 | 0,61500497 | 0,896 | 0,679 | 2,98E-56 |
| Prdm1 | 7,34E-46 | 0,61458392 | 0,777 | 0,537 | 1,36E-41 |
| Hilpda | 6,90E-51 | 0,61011317 | 0,432 | 0,157 | 1,28E-46 |
| Bcl2l1 | 5,13E-59 | 0,60947089 | 0,692 | 0,376 | 9,53E-55 |
| Slfn5 | 3,24E-53 | 0,60877059 | 0,283 | 0,049 | 6,03E-49 |
| Ddx58 | 5,91E-59 | 0,60115768 | 0,817 | 0,596 | 1,10E-54 |
| Ctss | 1,20E-59 | 0,5988058 | 0,913 | 0,741 | 2,23E-55 |
| Fam20a | 4,43E-78 | 0,59691001 | 0,302 | 0,014 | 8,23E-74 |
| Camk4 | 7,24E-64 | 0,59375262 | 0,897 | 0,729 | 1,35E-59 |
| Ppp1r3b | 2,61E-56 | 0,58883857 | 0,533 | 0,226 | 4,85E-52 |
| Fkbp5 | 6,65E-69 | 0,5865776 | 0,626 | 0,284 | 1,24E-64 |
| Sh3bp2 | 8,91E-69 | 0,58494573 | 0,68 | 0,336 | 1,66E-64 |
| Stat2 | 3,68E-53 | 0,58253414 | 0,714 | 0,455 | 6,83E-49 |
| Dtx3l | 7,47E-69 | 0,58232012 | 0,953 | 0,899 | 1,39E-64 |
| Epas1 | 7,15E-65 | 0,58169955 | 0,292 | 0,032 | 1,33E-60 |
| Samsn1 | 1,19E-43 | 0,5808629 | 0,843 | 0,644 | 2,21E-39 |
| Adprm | 2,17E-55 | 0,57634005 | 0,676 | 0,409 | 4,04E-51 |
| Ly6a | 1,01E-83 | 0,573338 | 1 | 0,993 | 1,87E-79 |
| Cytip | 2,49E-94 | 0,57237377 | 0,999 | 0,989 | 4,63E-90 |
| Arsb | 1,25E-51 | 0,56633383 | 0,895 | 0,737 | 2,32E-47 |
| Spats2 | 4,31E-60 | 0,55607964 | 0,404 | 0,112 | 8,02E-56 |
| Alcam | 3,33E-72 | 0,55581077 | 0,367 | 0,064 | 6,19E-68 |
| Ube2l6 | 1,30E-68 | 0,55522045 | 0,429 | 0,111 | 2,41E-64 |
| Rgs16 | 8,91E-41 | 0,5498847 | 0,692 | 0,443 | 1,66E-36 |
| Slc39a10 | 1,58E-50 | 0,54582891 | 0,681 | 0,407 | 2,94E-46 |
| Clic4 | 8,84E-60 | 0,54434071 | 0,733 | 0,428 | 1,64E-55 |
| Ifi35 | 1,61E-53 | 0,53913189 | 0,869 | 0,688 | 3,00E-49 |

| | | | | | |
|---|---|---|---|---|---|
| Oasl2 | 9,77E-67 | 0,53810462 | 0,257 | 0,009 | 1,81E-62 |
| Ldha | 6,12E-57 | 0,53404785 | 0,99 | 0,976 | 1,14E-52 |
| Ccnyl1 | 3,42E-61 | 0,52688674 | 0,562 | 0,236 | 6,35E-57 |
| Capza2 | 2,08E-64 | 0,5266815 | 0,853 | 0,644 | 3,86E-60 |
| Parp14 | 1,69E-48 | 0,52624029 | 0,936 | 0,806 | 3,14E-44 |
| Flot1 | 1,67E-53 | 0,52198392 | 0,794 | 0,535 | 3,10E-49 |
| Klhl6 | 6,99E-33 | 0,51926596 | 0,762 | 0,585 | 1,30E-28 |
| Ugcg | 1,20E-40 | 0,51537309 | 0,935 | 0,828 | 2,23E-36 |
| Chsy1 | 1,69E-48 | 0,51413786 | 0,976 | 0,917 | 3,15E-44 |
| Zc3h12d | 1,11E-40 | 0,51324019 | 0,812 | 0,594 | 2,07E-36 |
| Gpr171 | 1,08E-50 | 0,51118031 | 0,977 | 0,919 | 2,01E-46 |
| Frmd4b | 5,09E-52 | 0,51054847 | 0,679 | 0,382 | 9,45E-48 |
| Ier3 | 4,96E-53 | 0,50743273 | 0,309 | 0,062 | 9,22E-49 |
| Srgn | 1,12E-76 | 0,50648278 | 0,999 | 1 | 2,08E-72 |
| Gbp7 | 2,72E-58 | 0,50644774 | 0,943 | 0,834 | 5,06E-54 |
| Usp25 | 2,23E-64 | 0,50594618 | 0,973 | 0,921 | 4,15E-60 |
| Kbtbd11 | 1,15E-44 | 0,50433044 | 0,75 | 0,471 | 2,14E-40 |
| Ms4a6d | 6,00E-55 | 0,50321013 | 0,509 | 0,211 | 1,11E-50 |
| Parp12 | 1,96E-47 | 0,50188414 | 0,526 | 0,25 | 3,64E-43 |
| Serpinb6b | 3,50E-46 | 0,50122482 | 0,425 | 0,164 | 6,50E-42 |
| Cd86 | 1,31E-31 | 0,50057174 | 0,694 | 0,471 | 2,44E-27 |
| Trib2 | 2,61E-35 | 0,50008931 | 0,587 | 0,352 | 4,84E-31 |
| Lpar6 | 4,73E-38 | -0,5020899 | 0,323 | 0,51 | 8,79E-34 |
| Tnfsf10 | 3,84E-48 | -0,5025789 | 0,786 | 0,914 | 7,14E-44 |
| Jmy | 3,35E-48 | -0,5051589 | 0,225 | 0,447 | 6,23E-44 |
| mt-Cytb | 3,50E-42 | -0,5053769 | 0,758 | 0,884 | 6,51E-38 |
| Prkd2 | 1,86E-48 | -0,5085636 | 0,757 | 0,848 | 3,46E-44 |
| Wbp1 | 1,50E-35 | -0,5088912 | 0,512 | 0,658 | 2,78E-31 |
| Saraf | 1,71E-86 | -0,5108165 | 0,978 | 0,99 | 3,18E-82 |
| Cd82 | 1,32E-58 | -0,5113443 | 0,953 | 0,996 | 2,46E-54 |
| Ogt | 3,96E-56 | -0,5122527 | 0,892 | 0,939 | 7,35E-52 |
| Ltb | 4,20E-46 | -0,5159578 | 0,957 | 0,989 | 7,81E-42 |
| Dgkd | 1,90E-47 | -0,5159582 | 0,691 | 0,792 | 3,54E-43 |
| Nr1d2 | 3,31E-55 | -0,5173309 | 0,171 | 0,414 | 6,15E-51 |
| Slc9a9 | 1,97E-52 | -0,5174746 | 0,422 | 0,624 | 3,65E-48 |
| Susd6 | 8,38E-63 | -0,5180525 | 0,84 | 0,908 | 1,56E-58 |
| Antxr2 | 1,81E-47 | -0,5192772 | 0,413 | 0,616 | 3,37E-43 |
| Camk1d | 4,89E-52 | -0,5193715 | 0,338 | 0,565 | 9,08E-48 |
| Fam78a | 1,32E-58 | -0,5236425 | 0,877 | 0,938 | 2,45E-54 |
| Sh3bgrl3 | 8,09E-73 | -0,5309472 | 0,994 | 0,998 | 1,50E-68 |
| St8sia4 | 2,22E-37 | -0,5318241 | 0,346 | 0,524 | 4,12E-33 |
| Ski | 2,07E-61 | -0,532967 | 0,894 | 0,955 | 3,85E-57 |
| Il16 | 3,03E-59 | -0,5352437 | 0,781 | 0,885 | 5,63E-55 |
| Tgfbr2 | 9,74E-80 | -0,5368466 | 0,951 | 0,979 | 1,81E-75 |
| Fgl2 | 1,27E-47 | -0,5370374 | 0,943 | 0,97 | 2,35E-43 |
| Gdf11 | 1,55E-45 | -0,5379732 | 0,333 | 0,537 | 2,88E-41 |
| Emp3 | 2,33E-57 | -0,5395664 | 0,868 | 0,951 | 4,32E-53 |
| Maz | 6,97E-65 | -0,5421705 | 0,823 | 0,901 | 1,30E-60 |
| Tgfbr3 | 9,73E-67 | -0,5426314 | 0,252 | 0,537 | 1,81E-62 |
| Rb1 | 4,14E-65 | -0,5427464 | 0,608 | 0,775 | 7,69E-61 |
| Cd96 | 2,50E-59 | -0,5427485 | 0,769 | 0,904 | 4,64E-55 |
| Anp32a | 3,36E-74 | -0,5441539 | 0,813 | 0,917 | 6,25E-70 |
| Rarg | 3,90E-45 | -0,5441659 | 0,24 | 0,455 | 7,24E-41 |

| | | | | | |
|---|---|---|---|---|---|
| Fkbp3 | 6,61E-54 | -0,5459059 | 0,696 | 0,824 | 1,23E-49 |
| Gbp8 | 4,49E-49 | -0,5497601 | 0,739 | 0,839 | 8,34E-45 |
| Rgcc | 6,56E-33 | -0,5540386 | 0,123 | 0,29 | 1,22E-28 |
| Cd44 | 8,74E-57 | -0,5543601 | 0,911 | 0,968 | 1,62E-52 |
| Ap1s2 | 8,01E-54 | -0,5592339 | 0,512 | 0,695 | 1,49E-49 |
| Zfp652 | 2,82E-56 | -0,5611278 | 0,483 | 0,691 | 5,23E-52 |
| Nek7 | 2,76E-59 | -0,5645233 | 0,498 | 0,706 | 5,13E-55 |
| Arhgef3 | 1,76E-51 | -0,565791 | 0,731 | 0,87 | 3,26E-47 |
| Gabarapl2 | 2,47E-75 | -0,5712161 | 0,873 | 0,936 | 4,59E-71 |
| Adgre5 | 2,26E-47 | -0,5716111 | 0,858 | 0,934 | 4,20E-43 |
| Smad3 | 2,66E-61 | -0,572535 | 0,319 | 0,602 | 4,95E-57 |
| Ttc7 | 1,63E-67 | -0,5850899 | 0,647 | 0,817 | 3,02E-63 |
| Camk2n1 | 4,89E-44 | -0,5880483 | 0,497 | 0,681 | 9,10E-40 |
| Kcnc1 | 5,42E-84 | -0,5885954 | 0,095 | 0,385 | 1,01E-79 |
| Prkd3 | 3,81E-79 | -0,5898701 | 0,286 | 0,575 | 7,07E-75 |
| Gpr132 | 7,41E-31 | -0,5957108 | 0,703 | 0,786 | 1,38E-26 |
| Cdc42ep3 | 7,24E-62 | -0,5970238 | 0,376 | 0,621 | 1,34E-57 |
| Dennd4a | 1,38E-49 | -0,5971609 | 0,865 | 0,918 | 2,57E-45 |
| Tmem59 | 3,92E-81 | -0,599339 | 0,833 | 0,925 | 7,28E-77 |
| Tnfaip3 | 1,09E-44 | -0,6045402 | 0,983 | 0,987 | 2,03E-40 |
| Spsb1 | 2,80E-35 | -0,6102465 | 0,296 | 0,483 | 5,21E-31 |
| Atp2a3 | 9,54E-86 | -0,6108069 | 0,853 | 0,94 | 1,77E-81 |
| Numa1 | 2,27E-80 | -0,6109085 | 0,813 | 0,91 | 4,22E-76 |
| Mbp | 4,54E-63 | -0,6127864 | 0,519 | 0,705 | 8,44E-59 |
| Ly6g5b | 8,47E-46 | -0,6151359 | 0,275 | 0,511 | 1,57E-41 |
| St3gal6 | 7,49E-65 | -0,6203567 | 0,373 | 0,636 | 1,39E-60 |
| Top2b | 4,48E-78 | -0,6220596 | 0,747 | 0,858 | 8,32E-74 |
| Tcf7 | 2,06E-37 | -0,6278392 | 0,085 | 0,254 | 3,83E-33 |
| Sntb1 | 8,89E-80 | -0,6301146 | 0,288 | 0,581 | 1,65E-75 |
| Tspan13 | 3,65E-69 | -0,6339242 | 0,452 | 0,702 | 6,78E-65 |
| Il27ra | 9,61E-79 | -0,6344798 | 0,784 | 0,896 | 1,79E-74 |
| Sgms1 | 7,60E-71 | -0,6378754 | 0,22 | 0,509 | 1,41E-66 |
| Bcl11b | 1,12E-82 | -0,6382675 | 0,901 | 0,955 | 2,08E-78 |
| Serpinb1a | 1,82E-19 | -0,640203 | 0,161 | 0,291 | 3,38E-15 |
| Pim2 | 2,89E-34 | -0,6402106 | 0,37 | 0,544 | 5,38E-30 |
| Dnajc9 | 6,30E-57 | -0,6403564 | 0,768 | 0,865 | 1,17E-52 |
| Rftn1 | 2,00E-77 | -0,6425056 | 0,643 | 0,86 | 3,72E-73 |
| Ank | 1,88E-85 | -0,6447518 | 0,236 | 0,551 | 3,49E-81 |
| Ypel3 | 1,26E-71 | -0,649215 | 0,833 | 0,896 | 2,35E-67 |
| Lmna | 1,10E-61 | -0,6497964 | 0,086 | 0,324 | 2,05E-57 |
| Cmtm7 | 2,04E-82 | -0,6503774 | 0,729 | 0,897 | 3,78E-78 |
| Zfp683 | 6,71E-54 | -0,6591131 | 0,373 | 0,646 | 1,25E-49 |
| Aplp2 | 6,10E-77 | -0,6619389 | 0,482 | 0,727 | 1,13E-72 |
| Sesn3 | 1,72E-70 | -0,6634603 | 0,597 | 0,796 | 3,20E-66 |
| Rbpj | 4,03E-69 | -0,664287 | 0,847 | 0,948 | 7,49E-65 |
| Inpp4b | 5,32E-68 | -0,668404 | 0,793 | 0,919 | 9,89E-64 |
| Lrig1 | 2,97E-114 | -0,6694969 | 0,085 | 0,432 | 5,51E-110 |
| Itgax | 7,74E-36 | -0,6720727 | 0,568 | 0,715 | 1,44E-31 |
| Cd101 | 4,11E-60 | -0,6905188 | 0,289 | 0,536 | 7,63E-56 |
| Ppp2r2c | 2,38E-71 | -0,6945684 | 0,431 | 0,714 | 4,42E-67 |
| Ifnar2 | 1,08E-80 | -0,6997069 | 0,458 | 0,722 | 2,01E-76 |
| Spry2 | 4,76E-17 | -0,7002749 | 0,16 | 0,274 | 8,84E-13 |
| Emb | 1,94E-56 | -0,70768 | 0,862 | 0,928 | 3,60E-52 |

| | | | | | |
|---|---|---|---|---|---|
| Sorl1 | 6,33E-133 | -0,7076844 | 0,945 | 0,986 | 1,18E-128 |
| Rgs10 | 3,04E-64 | -0,7102623 | 0,472 | 0,715 | 5,64E-60 |
| Cd226 | 1,27E-97 | -0,7155711 | 0,832 | 0,96 | 2,37E-93 |
| Gpr34 | 4,03E-65 | -0,7165271 | 0,266 | 0,551 | 7,49E-61 |
| Igflr1 | 9,42E-64 | -0,7192516 | 0,515 | 0,74 | 1,75E-59 |
| Ddx5 | 1,71E-137 | -0,721292 | 0,953 | 0,985 | 3,18E-133 |
| Prkacb | 2,16E-101 | -0,7243929 | 0,807 | 0,929 | 4,02E-97 |
| Slc20a1 | 1,84E-71 | -0,7316789 | 0,733 | 0,854 | 3,42E-67 |
| Tesc | 6,96E-68 | -0,7345829 | 0,281 | 0,555 | 1,29E-63 |
| Btg2 | 4,66E-48 | -0,7356358 | 0,919 | 0,938 | 8,66E-44 |
| Crip1 | 2,45E-55 | -0,7363125 | 0,937 | 0,961 | 4,54E-51 |
| Neurl3 | 9,43E-23 | -0,7450259 | 0,526 | 0,62 | 1,75E-18 |
| Apbb1ip | 4,12E-107 | -0,7485312 | 0,808 | 0,942 | 7,66E-103 |
| Itm2b | 1,97E-195 | -0,7496354 | 0,995 | 0,998 | 3,67E-191 |
| Itgb1 | 4,03E-46 | -0,7498788 | 0,556 | 0,707 | 7,48E-42 |
| Znrf1 | 4,99E-101 | -0,7563514 | 0,782 | 0,915 | 9,28E-97 |
| Kctd12 | 4,26E-69 | -0,7627318 | 0,19 | 0,478 | 7,91E-65 |
| Myadm | 4,87E-53 | -0,7702783 | 0,424 | 0,646 | 9,05E-49 |
| Pdcd4 | 1,18E-96 | -0,7703435 | 0,73 | 0,908 | 2,19E-92 |
| Smpdl3a | 8,30E-92 | -0,7844685 | 0,518 | 0,773 | 1,54E-87 |
| Dnaja1 | 7,59E-82 | -0,7977096 | 0,747 | 0,854 | 1,41E-77 |
| Ifi27 | 6,13E-115 | -0,7980024 | 0,695 | 0,92 | 1,14E-110 |
| Itgb7 | 1,67E-140 | -0,800627 | 0,966 | 0,992 | 3,11E-136 |
| Evl | 3,40E-66 | -0,8010495 | 0,511 | 0,735 | 6,32E-62 |
| Cxcr3 | 1,31E-75 | -0,8105576 | 0,556 | 0,794 | 2,44E-71 |
| Fosb | 2,11E-19 | -0,8150831 | 0,348 | 0,477 | 3,92E-15 |
| Rgs1 | 3,20E-52 | -0,8174821 | 0,952 | 0,973 | 5,94E-48 |
| Eif4a2 | 2,25E-145 | -0,833129 | 0,917 | 0,979 | 4,18E-141 |
| Tcrg-C2 | 6,16E-28 | -0,83468 | 0,228 | 0,391 | 1,14E-23 |
| Rgs2 | 1,56E-29 | -0,8351846 | 0,34 | 0,509 | 2,90E-25 |
| Arnt2 | 4,87E-120 | -0,8380389 | 0,233 | 0,633 | 9,05E-116 |
| Sun2 | 1,67E-106 | -0,8469081 | 0,709 | 0,891 | 3,10E-102 |
| Hsph1 | 1,28E-82 | -0,8542942 | 0,399 | 0,663 | 2,38E-78 |
| P2ry10 | 3,71E-82 | -0,8672533 | 0,8 | 0,916 | 6,89E-78 |
| Lta | 1,95E-35 | -0,8693794 | 0,177 | 0,364 | 3,63E-31 |
| Mxd4 | 8,11E-97 | -0,8853705 | 0,49 | 0,759 | 1,51E-92 |
| Fos | 1,43E-17 | -0,8920459 | 0,463 | 0,57 | 2,67E-13 |
| Tagap | 2,88E-68 | -0,8955688 | 0,597 | 0,803 | 5,35E-64 |
| Gpr183 | 8,58E-68 | -0,9012182 | 0,496 | 0,737 | 1,59E-63 |
| Cxcr6 | 1,09E-124 | -0,9040392 | 0,91 | 0,988 | 2,02E-120 |
| Ldlrad4 | 1,37E-105 | -0,9112533 | 0,323 | 0,67 | 2,55E-101 |
| Ahnak | 3,54E-162 | -0,9384062 | 0,958 | 0,996 | 6,58E-158 |
| Dusp1 | 1,05E-33 | -0,9457855 | 0,676 | 0,779 | 1,95E-29 |
| Zfp36l2 | 1,88E-55 | -0,9610564 | 0,843 | 0,926 | 3,50E-51 |
| Ier5l | 1,36E-23 | -0,9851227 | 0,268 | 0,407 | 2,52E-19 |
| P2rx7 | 2,26E-76 | -0,9895181 | 0,287 | 0,581 | 4,20E-72 |
| Rhob | 3,88E-49 | -1,0036116 | 0,256 | 0,48 | 7,21E-45 |
| Qpct | 1,16E-166 | -1,0257083 | 0,065 | 0,506 | 2,15E-162 |
| Klf6 | 3,29E-119 | -1,0463278 | 0,952 | 0,987 | 6,12E-115 |
| Ptger4 | 1,55E-89 | -1,0535654 | 0,46 | 0,717 | 2,88E-85 |
| S100a4 | 3,47E-89 | -1,0537085 | 0,732 | 0,92 | 6,44E-85 |
| Klrg1 | 8,59E-71 | -1,0789679 | 0,34 | 0,613 | 1,60E-66 |
| Itga1 | 9,12E-142 | -1,1298396 | 0,569 | 0,884 | 1,69E-137 |

| | | | | | |
|---|---|---|---|---|---|
| Cdh1 | 1,57E-138 | -1,2432345 | 0,224 | 0,651 | 2,91E-134 |
| Jun | 8,05E-74 | -1,3619539 | 0,651 | 0,83 | 1,50E-69 |
| Itgae | 2,49E-127 | -1,9960653 | 0,193 | 0,587 | 4,63E-123 |

Cluster 2

| | p_val | avg_log2FC | pct.1 | pct.2 | p_val_adj |
|---|---|---|---|---|---|
| Gzmb | 5,71E-128 | 2,56291115 | 0,978 | 0,51 | 1,06E-123 |
| Isg15 | 1,54E-92 | 2,04578404 | 0,796 | 0,276 | 2,86E-88 |
| Ifit3 | 1,40E-77 | 2,04470437 | 0,631 | 0,116 | 2,59E-73 |
| Usp18 | 2,19E-118 | 1,84600579 | 0,837 | 0,223 | 4,08E-114 |
| AA467197 | 2,20E-81 | 1,84596578 | 0,834 | 0,404 | 4,09E-77 |
| Ccl3 | 3,12E-52 | 1,80939929 | 0,885 | 0,55 | 5,80E-48 |
| Ifi27l2a | 6,35E-63 | 1,6560355 | 0,897 | 0,668 | 1,18E-58 |
| Ifitm3 | 3,86E-49 | 1,64067764 | 0,403 | 0,028 | 7,18E-45 |
| Ifit1bl1 | 2,26E-55 | 1,6117399 | 0,708 | 0,313 | 4,21E-51 |
| Ifng | 1,58E-34 | 1,58653538 | 0,954 | 0,828 | 2,93E-30 |
| Icos | 3,16E-84 | 1,54018188 | 0,972 | 0,798 | 5,86E-80 |
| Tnfrsf4 | 2,53E-72 | 1,53930099 | 0,695 | 0,211 | 4,70E-68 |
| Cdkn1a | 1,13E-63 | 1,41996909 | 0,719 | 0,269 | 2,11E-59 |
| Rnf213 | 2,29E-46 | 1,40811372 | 0,885 | 0,694 | 4,25E-42 |
| Gzma | 1,09E-57 | 1,40073108 | 0,714 | 0,237 | 2,03E-53 |
| Bst2 | 1,18E-67 | 1,39876744 | 0,943 | 0,756 | 2,20E-63 |
| Sv2c | 9,75E-68 | 1,37133395 | 0,555 | 0,086 | 1,81E-63 |
| Sytl3 | 6,00E-103 | 1,3682149 | 0,928 | 0,578 | 1,12E-98 |
| Plac8 | 6,53E-45 | 1,29906121 | 0,918 | 0,735 | 1,21E-40 |
| Tnfrsf18 | 7,77E-79 | 1,29337726 | 0,929 | 0,705 | 1,44E-74 |
| Prf1 | 7,65E-43 | 1,28596391 | 0,949 | 0,824 | 1,42E-38 |
| Zbtb32 | 1,38E-90 | 1,27687337 | 0,689 | 0,128 | 2,57E-86 |
| Ifit1 | 1,00E-50 | 1,26315539 | 0,48 | 0,081 | 1,86E-46 |
| Ifi211 | 7,30E-73 | 1,26072481 | 0,7 | 0,223 | 1,36E-68 |
| Isg20 | 9,07E-51 | 1,24640863 | 0,829 | 0,555 | 1,68E-46 |
| Rtp4 | 3,65E-56 | 1,23155676 | 0,639 | 0,227 | 6,78E-52 |
| Pik3ap1 | 5,78E-68 | 1,21488909 | 0,912 | 0,613 | 1,07E-63 |
| Zbp1 | 1,09E-55 | 1,18879912 | 0,902 | 0,647 | 2,03E-51 |
| Irf7 | 4,66E-58 | 1,18757494 | 0,749 | 0,357 | 8,66E-54 |
| Ifih1 | 1,22E-39 | 1,17381407 | 0,572 | 0,216 | 2,26E-35 |
| Ms4a4c | 1,47E-44 | 1,16138241 | 0,608 | 0,251 | 2,74E-40 |
| Entpd1 | 1,92E-69 | 1,14685934 | 0,752 | 0,297 | 3,56E-65 |
| Havcr2 | 2,96E-63 | 1,14538782 | 0,865 | 0,487 | 5,50E-59 |
| Id2 | 7,74E-67 | 1,10017276 | 0,989 | 0,935 | 1,44E-62 |
| Il1r2 | 1,31E-25 | 1,09889691 | 0,299 | 0,053 | 2,43E-21 |
| Rsad2 | 1,32E-41 | 1,086509 | 0,368 | 0,03 | 2,45E-37 |
| Slc7a5 | 4,24E-43 | 1,07477615 | 0,812 | 0,503 | 7,88E-39 |
| Xaf1 | 2,10E-60 | 1,06017891 | 0,77 | 0,385 | 3,90E-56 |
| Oas3 | 1,82E-42 | 1,04527084 | 0,594 | 0,244 | 3,38E-38 |
| Mxd1 | 2,85E-36 | 1,02749753 | 0,796 | 0,536 | 5,29E-32 |
| Pkm | 6,94E-81 | 1,02523176 | 0,975 | 0,875 | 1,29E-76 |
| Chchd10 | 4,60E-29 | 1,02050746 | 0,625 | 0,36 | 8,55E-25 |
| Cytip | 2,80E-73 | 1,00621959 | 0,984 | 0,937 | 5,20E-69 |
| Daxx | 3,55E-41 | 1,00326533 | 0,783 | 0,483 | 6,60E-37 |
| Tpi1 | 1,04E-45 | 1,00128431 | 0,753 | 0,439 | 1,93E-41 |
| Sdcbp2 | 2,87E-49 | 1,00091803 | 0,619 | 0,23 | 5,33E-45 |
| Arl14ep | 1,03E-40 | 0,97054848 | 0,583 | 0,255 | 1,92E-36 |
| Oas1a | 1,13E-56 | 0,96244505 | 0,548 | 0,116 | 2,10E-52 |
| Trafd1 | 3,29E-51 | 0,94634965 | 0,794 | 0,492 | 6,12E-47 |
| Hif1a | 8,29E-67 | 0,94115257 | 0,965 | 0,749 | 1,54E-62 |

Up in Sp140-/-
Up in Sp140+/+

| | | | | | |
|---|---|---|---|---|---|
| Serpinb9 | 5,55E-34 | 0,92400826 | 0,733 | 0,457 | 1,03E-29 |
| Hk2 | 1,71E-46 | 0,91632706 | 0,72 | 0,341 | 3,17E-42 |
| Ifi209 | 1,31E-35 | 0,91103083 | 0,847 | 0,582 | 2,43E-31 |
| Ifit2 | 3,40E-30 | 0,90558487 | 0,507 | 0,223 | 6,33E-26 |
| Slfn8 | 6,66E-60 | 0,90347057 | 0,838 | 0,492 | 1,24E-55 |
| Dhx58 | 2,23E-50 | 0,90066802 | 0,656 | 0,262 | 4,15E-46 |
| Slfn1 | 1,92E-41 | 0,89902 | 0,82 | 0,515 | 3,57E-37 |
| Il2ra | 7,87E-31 | 0,89145981 | 0,862 | 0,668 | 1,46E-26 |
| Il21 | 1,24E-22 | 0,88219019 | 0,288 | 0,06 | 2,30E-18 |
| Pstpip1 | 1,32E-59 | 0,87636437 | 0,942 | 0,768 | 2,46E-55 |
| Slc39a10 | 5,86E-47 | 0,86949219 | 0,808 | 0,52 | 1,09E-42 |
| Phf11b | 1,43E-35 | 0,86853126 | 0,829 | 0,587 | 2,65E-31 |
| Rbm3 | 2,87E-70 | 0,86537362 | 0,948 | 0,791 | 5,34E-66 |
| Batf | 1,92E-51 | 0,86292105 | 0,924 | 0,673 | 3,57E-47 |
| Chd7 | 1,04E-52 | 0,86278957 | 0,931 | 0,759 | 1,93E-48 |
| Slc16a3 | 1,32E-30 | 0,85536744 | 0,482 | 0,197 | 2,46E-26 |
| Alcam | 8,87E-42 | 0,84030532 | 0,656 | 0,288 | 1,65E-37 |
| Gnptab | 2,21E-56 | 0,83024113 | 0,889 | 0,661 | 4,11E-52 |
| Lgals3bp | 3,37E-42 | 0,82233669 | 0,955 | 0,858 | 6,26E-38 |
| Fam20a | 7,42E-40 | 0,82013987 | 0,327 | 0,012 | 1,38E-35 |
| Dgat1 | 3,32E-45 | 0,8201138 | 0,83 | 0,543 | 6,16E-41 |
| Ccl4 | 1,96E-20 | 0,81713629 | 0,955 | 0,803 | 3,64E-16 |
| Emp1 | 1,02E-24 | 0,79855727 | 0,432 | 0,174 | 1,90E-20 |
| Hilpda | 5,20E-34 | 0,79750449 | 0,547 | 0,23 | 9,66E-30 |
| Ctla2a | 1,97E-20 | 0,79282971 | 0,91 | 0,789 | 3,66E-16 |
| Tg | 9,66E-42 | 0,78511756 | 0,52 | 0,144 | 1,79E-37 |
| Slamf1 | 8,12E-37 | 0,78194762 | 0,779 | 0,434 | 1,51E-32 |
| Sema7a | 5,13E-38 | 0,7697302 | 0,834 | 0,534 | 9,54E-34 |
| Lag3 | 1,98E-27 | 0,76254992 | 0,967 | 0,893 | 3,69E-23 |
| Ermn | 2,28E-28 | 0,75608488 | 0,275 | 0,028 | 4,23E-24 |
| Ly6a | 4,02E-46 | 0,73863348 | 0,995 | 0,886 | 7,48E-42 |
| Herc6 | 1,09E-27 | 0,73499556 | 0,697 | 0,439 | 2,02E-23 |
| Helz2 | 1,09E-30 | 0,7333061 | 0,91 | 0,794 | 2,03E-26 |
| Clic4 | 2,27E-36 | 0,72688279 | 0,739 | 0,459 | 4,22E-32 |
| Frmd4b | 1,61E-42 | 0,72329164 | 0,628 | 0,267 | 2,99E-38 |
| Phf11c | 2,19E-33 | 0,72028526 | 0,606 | 0,297 | 4,07E-29 |
| Gem | 4,46E-21 | 0,71855525 | 0,798 | 0,608 | 8,29E-17 |
| Lamc1 | 1,86E-48 | 0,71612151 | 0,833 | 0,506 | 3,45E-44 |
| Slc25a19 | 5,95E-39 | 0,71329172 | 0,812 | 0,552 | 1,11E-34 |
| Twsg1 | 8,08E-36 | 0,70653633 | 0,675 | 0,341 | 1,50E-31 |
| Aldoa | 1,56E-38 | 0,70643034 | 0,964 | 0,87 | 2,90E-34 |
| Setdb1 | 3,04E-18 | 0,70463444 | 0,609 | 0,418 | 5,65E-14 |
| Bcl2l1 | 3,87E-45 | 0,7044585 | 0,814 | 0,476 | 7,18E-41 |
| Slfn5 | 6,05E-25 | 0,70356397 | 0,279 | 0,046 | 1,12E-20 |
| Rilpl2 | 1,36E-41 | 0,70269994 | 0,922 | 0,701 | 2,53E-37 |
| Slc2a1 | 1,02E-31 | 0,70021724 | 0,706 | 0,422 | 1,90E-27 |
| Mapkapk2 | 2,54E-55 | 0,69784565 | 0,975 | 0,891 | 4,72E-51 |
| Ifi208 | 9,30E-32 | 0,69693979 | 0,408 | 0,118 | 1,73E-27 |
| Irf8 | 2,47E-30 | 0,6968705 | 0,92 | 0,703 | 4,59E-26 |
| Serpinb6b | 4,03E-35 | 0,68713251 | 0,506 | 0,179 | 7,49E-31 |
| Smap2 | 2,78E-37 | 0,68265257 | 0,945 | 0,858 | 5,18E-33 |
| Satb1 | 1,04E-40 | 0,67066833 | 0,976 | 0,789 | 1,93E-36 |
| Asns | 8,18E-21 | 0,66888056 | 0,475 | 0,237 | 1,52E-16 |

| | | | | | |
|---|---|---|---|---|---|
| Il18rap | 6,85E-30 | 0,66507875 | 0,958 | 0,814 | 1,27E-25 |
| Parp9 | 2,19E-36 | 0,6620497 | 0,782 | 0,48 | 4,08E-32 |
| Ctla4 | 2,70E-22 | 0,66140909 | 0,952 | 0,861 | 5,02E-18 |
| Ldha | 7,09E-43 | 0,66049089 | 0,989 | 0,93 | 1,32E-38 |
| Smim3 | 3,47E-41 | 0,65230625 | 0,661 | 0,329 | 6,44E-37 |
| Mif4gd | 3,21E-38 | 0,650309 | 0,913 | 0,747 | 5,97E-34 |
| Parp14 | 3,23E-20 | 0,64196001 | 0,833 | 0,661 | 6,00E-16 |
| Glrx | 2,24E-12 | 0,63601944 | 0,74 | 0,599 | 4,17E-08 |
| Glipr2 | 8,32E-34 | 0,63546566 | 0,874 | 0,65 | 1,55E-29 |
| Pml | 1,14E-26 | 0,63382396 | 0,777 | 0,575 | 2,12E-22 |
| Dtx3l | 4,79E-28 | 0,63190869 | 0,872 | 0,733 | 8,90E-24 |
| Hlx | 5,93E-29 | 0,63190373 | 0,391 | 0,104 | 1,10E-24 |
| Marcksl1 | 2,75E-11 | 0,62912449 | 0,542 | 0,374 | 5,11E-07 |
| Farp1 | 7,87E-33 | 0,62833889 | 0,703 | 0,387 | 1,46E-28 |
| Nt5c3 | 4,61E-18 | 0,62806087 | 0,695 | 0,499 | 8,57E-14 |
| Ddit4 | 1,35E-23 | 0,62646883 | 0,704 | 0,455 | 2,52E-19 |
| Klrc1 | 2,56E-27 | 0,62549832 | 0,927 | 0,696 | 4,77E-23 |
| Tmem163 | 4,90E-42 | 0,62375514 | 0,778 | 0,413 | 9,10E-38 |
| Sco1 | 1,42E-39 | 0,61821469 | 0,579 | 0,227 | 2,64E-35 |
| Il1rl1 | 7,49E-20 | 0,61780505 | 0,341 | 0,118 | 1,39E-15 |
| Ddt | 4,30E-23 | 0,61572597 | 0,756 | 0,513 | 7,99E-19 |
| Gstt1 | 1,82E-13 | 0,61042608 | 0,434 | 0,248 | 3,38E-09 |
| Gbp7 | 1,26E-31 | 0,60866688 | 0,842 | 0,578 | 2,35E-27 |
| Zc3h12d | 1,16E-27 | 0,60853422 | 0,868 | 0,654 | 2,15E-23 |
| Cd247 | 3,46E-35 | 0,60510117 | 0,953 | 0,845 | 6,44E-31 |
| Capza2 | 1,20E-46 | 0,60229828 | 0,875 | 0,659 | 2,23E-42 |
| Camk4 | 3,08E-37 | 0,60203158 | 0,91 | 0,708 | 5,73E-33 |
| Epas1 | 1,67E-29 | 0,60126483 | 0,294 | 0,032 | 3,11E-25 |
| Smpdl3b | 1,22E-37 | 0,60104248 | 0,534 | 0,188 | 2,26E-33 |
| Ccr2 | 1,36E-19 | 0,59952273 | 0,484 | 0,23 | 2,52E-15 |
| Aars | 5,52E-22 | 0,59488945 | 0,764 | 0,578 | 1,03E-17 |
| Cblb | 3,34E-27 | 0,59421521 | 0,943 | 0,865 | 6,21E-23 |
| Eif2ak2 | 2,45E-32 | 0,59278411 | 0,596 | 0,276 | 4,55E-28 |
| Acadl | 1,04E-14 | 0,58894437 | 0,797 | 0,682 | 1,93E-10 |
| Cd2 | 1,29E-33 | 0,58877701 | 0,983 | 0,926 | 2,39E-29 |
| Sdc3 | 1,75E-27 | 0,58861047 | 0,3 | 0,049 | 3,25E-23 |
| Rhbdf2 | 2,93E-31 | 0,5877585 | 0,761 | 0,497 | 5,44E-27 |
| Trim30a | 8,54E-31 | 0,58176923 | 0,763 | 0,508 | 1,59E-26 |
| Ier3 | 1,76E-22 | 0,58013051 | 0,331 | 0,093 | 3,27E-18 |
| Sys1 | 1,74E-38 | 0,5787466 | 0,881 | 0,68 | 3,23E-34 |
| Trim25 | 2,72E-31 | 0,57798837 | 0,79 | 0,543 | 5,06E-27 |
| Wdfy1 | 1,83E-39 | 0,57304646 | 0,776 | 0,457 | 3,40E-35 |
| Themis | 7,37E-27 | 0,57160832 | 0,953 | 0,803 | 1,37E-22 |
| Arhgef10 | 5,91E-35 | 0,56855985 | 0,388 | 0,074 | 1,10E-30 |
| Itm2c | 5,07E-27 | 0,56652987 | 0,896 | 0,738 | 9,42E-23 |
| Ostf1 | 2,85E-25 | 0,56628879 | 0,947 | 0,87 | 5,30E-21 |
| Map2k3 | 2,24E-24 | 0,56224731 | 0,92 | 0,763 | 4,17E-20 |
| Spats2 | 4,03E-38 | 0,56133712 | 0,374 | 0,051 | 7,49E-34 |
| Ifi206 | 5,06E-16 | 0,55192854 | 0,613 | 0,425 | 9,41E-12 |
| Smox | 2,10E-10 | 0,55023921 | 0,428 | 0,276 | 3,89E-06 |
| Mdfic | 3,18E-31 | 0,55020909 | 0,897 | 0,735 | 5,91E-27 |
| Cd53 | 2,09E-33 | 0,54770323 | 0,995 | 0,981 | 3,89E-29 |
| Npc2 | 2,80E-27 | 0,54667599 | 0,959 | 0,868 | 5,20E-23 |

| | | | | | |
|---|---|---|---|---|---|
| Srgn | 7,63E-31 | 0,54544206 | 0,999 | 0,974 | 1,42E-26 |
| Cd47 | 4,39E-35 | 0,54481217 | 0,99 | 0,914 | 8,17E-31 |
| Pim1 | 6,80E-28 | 0,54419868 | 0,964 | 0,777 | 1,26E-23 |
| Il7r | 1,36E-23 | 0,54297782 | 0,653 | 0,376 | 2,54E-19 |
| Phf11a | 5,99E-37 | 0,54157147 | 0,473 | 0,132 | 1,11E-32 |
| Trim30d | 7,70E-34 | 0,54048087 | 0,584 | 0,253 | 1,43E-29 |
| Prdm1 | 6,41E-28 | 0,53967669 | 0,771 | 0,485 | 1,19E-23 |
| Prelid1 | 1,27E-27 | 0,53727775 | 0,935 | 0,858 | 2,36E-23 |
| Adamts14 | 1,23E-30 | 0,5372438 | 0,484 | 0,169 | 2,29E-26 |
| Arsb | 9,40E-24 | 0,53133791 | 0,789 | 0,592 | 1,75E-19 |
| Setbp1 | 1,81E-30 | 0,5265227 | 0,868 | 0,624 | 3,36E-26 |
| Ccrl2 | 2,55E-21 | 0,52586938 | 0,337 | 0,104 | 4,74E-17 |
| Tubb6 | 8,79E-30 | 0,51938927 | 0,304 | 0,039 | 1,63E-25 |
| Cdkn2d | 4,91E-24 | 0,51827694 | 0,673 | 0,413 | 9,12E-20 |
| Ifi35 | 7,31E-23 | 0,51506838 | 0,82 | 0,638 | 1,36E-18 |
| Myd88 | 7,18E-31 | 0,51348147 | 0,799 | 0,545 | 1,33E-26 |
| Lmnb1 | 3,12E-23 | 0,51304771 | 0,898 | 0,747 | 5,80E-19 |
| Trib2 | 2,62E-13 | 0,51241936 | 0,549 | 0,378 | 4,88E-09 |
| Rbm47 | 1,30E-37 | 0,51076702 | 0,528 | 0,16 | 2,42E-33 |
| Tnfrsf9 | 1,17E-22 | 0,50778281 | 0,917 | 0,756 | 2,18E-18 |
| Nfil3 | 4,42E-23 | 0,50598518 | 0,688 | 0,406 | 8,21E-19 |
| Adprm | 5,33E-30 | 0,50521507 | 0,717 | 0,425 | 9,91E-26 |
| Chmp4b | 3,81E-37 | 0,50328516 | 0,945 | 0,803 | 7,08E-33 |
| Camk2n1 | 2,61E-11 | -0,5000559 | 0,416 | 0,529 | 4,84E-07 |
| Susd6 | 3,85E-28 | -0,5027743 | 0,787 | 0,868 | 7,16E-24 |
| Prkd2 | 4,34E-15 | -0,506764 | 0,592 | 0,689 | 8,07E-11 |
| Ifrd1 | 5,53E-10 | -0,5067802 | 0,651 | 0,712 | 1,03E-05 |
| Rasgrp1 | 4,34E-21 | -0,5089954 | 0,792 | 0,868 | 8,07E-17 |
| Inpp4a | 8,39E-20 | -0,5090191 | 0,429 | 0,606 | 1,56E-15 |
| Cd52 | 2,07E-16 | -0,510526 | 0,984 | 0,984 | 3,84E-12 |
| Klf6 | 1,13E-13 | -0,5148278 | 0,923 | 0,937 | 2,10E-09 |
| Stat6 | 2,52E-32 | -0,5171093 | 0,852 | 0,907 | 4,68E-28 |
| Dgkd | 1,13E-22 | -0,5252393 | 0,711 | 0,784 | 2,10E-18 |
| Bcor | 1,23E-15 | -0,5257823 | 0,48 | 0,61 | 2,28E-11 |
| Csk | 1,33E-24 | -0,5261516 | 0,846 | 0,884 | 2,48E-20 |
| Il2rg | 6,94E-36 | -0,5277568 | 0,99 | 0,988 | 1,29E-31 |
| Fam53b | 9,72E-18 | -0,5290631 | 0,649 | 0,726 | 1,81E-13 |
| Stim2 | 7,14E-32 | -0,5291284 | 0,357 | 0,585 | 1,33E-27 |
| Ogt | 1,57E-23 | -0,5317222 | 0,923 | 0,942 | 2,92E-19 |
| Tob1 | 7,02E-16 | -0,5356525 | 0,37 | 0,515 | 1,30E-11 |
| Foxp1 | 1,48E-17 | -0,5393597 | 0,646 | 0,745 | 2,75E-13 |
| Rasal3 | 2,19E-24 | -0,5412982 | 0,893 | 0,923 | 4,07E-20 |
| Ppp2r2c | 5,03E-16 | -0,5419635 | 0,309 | 0,485 | 9,35E-12 |
| Ifnar1 | 5,76E-21 | -0,5428544 | 0,73 | 0,789 | 1,07E-16 |
| Emb | 7,24E-12 | -0,5433807 | 0,841 | 0,852 | 1,35E-07 |
| Numa1 | 1,94E-31 | -0,5435402 | 0,788 | 0,849 | 3,61E-27 |
| Igflr1 | 4,39E-20 | -0,5437799 | 0,483 | 0,643 | 8,16E-16 |
| Rftn1 | 7,59E-22 | -0,5443089 | 0,602 | 0,749 | 1,41E-17 |
| Akna | 8,87E-26 | -0,5449778 | 0,898 | 0,93 | 1,65E-21 |
| Tox | 1,02E-42 | -0,5471051 | 0,068 | 0,292 | 1,89E-38 |
| Chd3 | 2,46E-17 | -0,5473979 | 0,565 | 0,664 | 4,57E-13 |
| Lasp1 | 1,00E-22 | -0,5474444 | 0,721 | 0,794 | 1,86E-18 |
| Ctps2 | 1,27E-14 | -0,549041 | 0,567 | 0,661 | 2,35E-10 |

| | | | | | |
|---|---|---|---|---|---|
| Ncf1 | 1,48E-09 | -0,5496272 | 0,407 | 0,51 | 2,75E-05 |
| Actn1 | 2,50E-19 | -0,5514811 | 0,184 | 0,353 | 4,64E-15 |
| Cdh1 | 2,84E-35 | -0,5573183 | 0,076 | 0,278 | 5,28E-31 |
| Hsp90b1 | 2,13E-23 | -0,5595475 | 0,829 | 0,87 | 3,97E-19 |
| Ahnak | 2,42E-19 | -0,5602461 | 0,913 | 0,926 | 4,50E-15 |
| Cd44 | 1,84E-17 | -0,5626407 | 0,864 | 0,898 | 3,42E-13 |
| Plcg2 | 2,12E-26 | -0,5636923 | 0,21 | 0,415 | 3,94E-22 |
| Smpdl3a | 1,05E-21 | -0,5658439 | 0,374 | 0,552 | 1,94E-17 |
| Itm2b | 2,19E-29 | -0,5659038 | 0,986 | 0,972 | 4,07E-25 |
| Mgat5 | 8,26E-21 | -0,5662309 | 0,347 | 0,531 | 1,53E-16 |
| Gtf2i | 4,22E-28 | -0,5663914 | 0,766 | 0,842 | 7,84E-24 |
| Saraf | 1,34E-29 | -0,5677186 | 0,958 | 0,947 | 2,49E-25 |
| Tle4 | 1,91E-24 | -0,5706539 | 0,385 | 0,571 | 3,56E-20 |
| Myo1f | 1,54E-18 | -0,5717193 | 0,659 | 0,733 | 2,87E-14 |
| Sun2 | 9,53E-21 | -0,5729181 | 0,763 | 0,817 | 1,77E-16 |
| mt-Cytb | 3,58E-17 | -0,5739252 | 0,821 | 0,884 | 6,65E-13 |
| Il27ra | 7,35E-27 | -0,5739487 | 0,669 | 0,773 | 1,37E-22 |
| Cd8a | 2,87E-41 | -0,5743967 | 0,998 | 0,993 | 5,32E-37 |
| Pdia4 | 6,01E-22 | -0,5749114 | 0,713 | 0,796 | 1,12E-17 |
| Ifnar2 | 6,92E-30 | -0,5752353 | 0,4 | 0,626 | 1,29E-25 |
| Tspan13 | 1,19E-33 | -0,5775626 | 0,351 | 0,608 | 2,22E-29 |
| Junb | 5,73E-10 | -0,5812665 | 0,985 | 0,958 | 1,07E-05 |
| Dnaja1 | 2,32E-32 | -0,5850817 | 0,788 | 0,856 | 4,32E-28 |
| Tnfsf10 | 1,52E-15 | -0,5865802 | 0,689 | 0,759 | 2,83E-11 |
| Itgb2 | 1,88E-43 | -0,5901305 | 0,985 | 0,991 | 3,49E-39 |
| Rgs10 | 6,16E-22 | -0,5917587 | 0,23 | 0,427 | 1,14E-17 |
| Cdk17 | 1,93E-38 | -0,5922874 | 0,908 | 0,958 | 3,60E-34 |
| Matk | 2,36E-35 | -0,5971105 | 0,106 | 0,325 | 4,39E-31 |
| Sntb1 | 8,18E-44 | -0,6097442 | 0,148 | 0,418 | 1,52E-39 |
| Adgrl1 | 7,29E-39 | -0,6163725 | 0,253 | 0,529 | 1,36E-34 |
| Lta4h | 1,74E-31 | -0,6225163 | 0,606 | 0,747 | 3,23E-27 |
| Tesc | 2,67E-38 | -0,6245525 | 0,196 | 0,469 | 4,97E-34 |
| Nfatc1 | 3,95E-12 | -0,6246982 | 0,8 | 0,803 | 7,34E-08 |
| Mcm6 | 3,45E-23 | -0,6263087 | 0,795 | 0,872 | 6,41E-19 |
| Lat2 | 1,28E-17 | -0,6316634 | 0,189 | 0,362 | 2,37E-13 |
| Calr | 6,64E-32 | -0,6325823 | 0,799 | 0,87 | 1,23E-27 |
| Mmd | 9,62E-16 | -0,6330532 | 0,478 | 0,613 | 1,79E-11 |
| Cxcr3 | 7,77E-23 | -0,6346966 | 0,351 | 0,557 | 1,44E-18 |
| Ddx5 | 5,13E-52 | -0,6435772 | 0,955 | 0,972 | 9,52E-48 |
| Eif4a2 | 4,79E-46 | -0,6473134 | 0,949 | 0,968 | 8,91E-42 |
| Pdia6 | 2,63E-22 | -0,6504842 | 0,806 | 0,875 | 4,89E-18 |
| Lrig1 | 1,00E-63 | -0,6512506 | 0,115 | 0,439 | 1,86E-59 |
| Per1 | 5,47E-16 | -0,6517508 | 0,855 | 0,858 | 1,02E-11 |
| Ctsw | 2,21E-46 | -0,6524903 | 0,958 | 0,986 | 4,11E-42 |
| Sh3bgrl3 | 1,67E-30 | -0,6526516 | 0,948 | 0,935 | 3,09E-26 |
| Atp2a3 | 3,92E-32 | -0,6540959 | 0,648 | 0,784 | 7,28E-28 |
| St8sia4 | 1,09E-30 | -0,6546342 | 0,218 | 0,448 | 2,02E-26 |
| Trat1 | 2,68E-64 | -0,6662258 | 0,066 | 0,348 | 4,98E-60 |
| Ptger4 | 2,15E-19 | -0,6718975 | 0,616 | 0,742 | 4,00E-15 |
| Inpp4b | 1,38E-31 | -0,6728872 | 0,75 | 0,863 | 2,56E-27 |
| Stap1 | 1,13E-35 | -0,6729874 | 0,473 | 0,696 | 2,10E-31 |
| Tle3 | 7,46E-30 | -0,6760517 | 0,781 | 0,893 | 1,39E-25 |
| Irf2bp2 | 2,52E-31 | -0,6766231 | 0,771 | 0,877 | 4,69E-27 |

| | | | | | |
|---|---|---|---|---|---|
| Bach2 | 1,04E-37 | -0,6789273 | 0,139 | 0,387 | 1,93E-33 |
| Kcnn4 | 1,84E-25 | -0,6870663 | 0,582 | 0,735 | 3,42E-21 |
| Arhgef3 | 1,26E-26 | -0,6871432 | 0,677 | 0,814 | 2,34E-22 |
| Nek7 | 5,63E-35 | -0,6876725 | 0,436 | 0,668 | 1,05E-30 |
| Jun | 6,23E-12 | -0,6911549 | 0,685 | 0,738 | 1,16E-07 |
| Tgfbr2 | 6,64E-43 | -0,6925042 | 0,899 | 0,94 | 1,23E-38 |
| Irs2 | 1,02E-29 | -0,7005992 | 0,075 | 0,262 | 1,90E-25 |
| Dusp1 | 3,51E-14 | -0,7053522 | 0,727 | 0,777 | 6,52E-10 |
| Ldlrad4 | 1,38E-34 | -0,7135789 | 0,269 | 0,524 | 2,56E-30 |
| Kif13b | 1,91E-17 | -0,7162247 | 0,476 | 0,585 | 3,56E-13 |
| Cd5 | 1,09E-08 | -0,7195776 | 0,805 | 0,752 | 0,0002028 |
| Itgb7 | 9,87E-19 | -0,7223065 | 0,809 | 0,842 | 1,83E-14 |
| Sorl1 | 3,02E-38 | -0,7289095 | 0,809 | 0,884 | 5,62E-34 |
| Itpkb | 5,34E-35 | -0,7295643 | 0,637 | 0,787 | 9,91E-31 |
| Chn2 | 7,35E-24 | -0,732889 | 0,178 | 0,371 | 1,37E-19 |
| Hsph1 | 5,98E-33 | -0,7512758 | 0,583 | 0,756 | 1,11E-28 |
| P2rx7 | 6,75E-32 | -0,7529319 | 0,226 | 0,471 | 1,25E-27 |
| Eomes | 2,72E-08 | -0,7548462 | 0,255 | 0,357 | 0,00050575 |
| Ptpn6 | 2,78E-28 | -0,7575649 | 0,863 | 0,923 | 5,16E-24 |
| Pdcd4 | 4,18E-38 | -0,7623017 | 0,697 | 0,838 | 7,76E-34 |
| Dgkz | 1,41E-22 | -0,7753472 | 0,925 | 0,912 | 2,61E-18 |
| Egr2 | 2,00E-17 | -0,7761776 | 0,409 | 0,589 | 3,71E-13 |
| Rgs3 | 1,32E-40 | -0,7820553 | 0,918 | 0,958 | 2,45E-36 |
| Cyth3 | 8,57E-31 | -0,7883743 | 0,336 | 0,566 | 1,59E-26 |
| Itga1 | 1,10E-26 | -0,7945984 | 0,389 | 0,592 | 2,05E-22 |
| Tagap | 6,45E-33 | -0,8084499 | 0,793 | 0,93 | 1,20E-28 |
| Fgl2 | 3,31E-14 | -0,8095559 | 0,694 | 0,752 | 6,16E-10 |
| Klf2 | 8,68E-08 | -0,8101507 | 0,307 | 0,408 | 0,00161279 |
| Rasa1 | 3,84E-41 | -0,8179093 | 0,659 | 0,826 | 7,14E-37 |
| P2ry10 | 1,01E-35 | -0,8197055 | 0,728 | 0,845 | 1,88E-31 |
| Tcrg-C4 | 3,69E-69 | -0,8198225 | 0,029 | 0,274 | 6,87E-65 |
| Btg2 | 5,04E-30 | -0,8259805 | 0,933 | 0,94 | 9,36E-26 |
| Rhob | 9,91E-20 | -0,8271146 | 0,303 | 0,487 | 1,84E-15 |
| Cd101 | 6,84E-43 | -0,8283652 | 0,296 | 0,568 | 1,27E-38 |
| Sesn3 | 8,39E-50 | -0,84225 | 0,477 | 0,742 | 1,56E-45 |
| Zfp683 | 4,85E-33 | -0,843605 | 0,199 | 0,441 | 9,02E-29 |
| Ephx1 | 4,22E-67 | -0,858532 | 0,045 | 0,306 | 7,84E-63 |
| Hivep3 | 8,75E-16 | -0,8726055 | 0,35 | 0,524 | 1,63E-11 |
| Mxd4 | 6,79E-26 | -0,8875079 | 0,39 | 0,571 | 1,26E-21 |
| Samd3 | 6,42E-61 | -0,8886666 | 0,123 | 0,441 | 1,19E-56 |
| Bcl6 | 5,02E-37 | -0,9151364 | 0,285 | 0,545 | 9,33E-33 |
| Sema6d | 3,09E-69 | -0,9249505 | 0,034 | 0,285 | 5,75E-65 |
| Adgre5 | 2,15E-38 | -0,9454831 | 0,813 | 0,879 | 3,99E-34 |
| Itgb1 | 1,96E-33 | -0,9496391 | 0,607 | 0,754 | 3,64E-29 |
| Tcrg-C2 | 9,67E-44 | -0,9821379 | 0,124 | 0,381 | 1,80E-39 |
| S100a4 | 2,75E-18 | -0,9914776 | 0,547 | 0,691 | 5,11E-14 |
| Ipcef1 | 9,41E-28 | -1,0294363 | 0,663 | 0,775 | 1,75E-23 |
| Neurl3 | 2,22E-38 | -1,0350449 | 0,386 | 0,645 | 4,13E-34 |
| Tcf7 | 3,82E-62 | -1,0427347 | 0,107 | 0,418 | 7,10E-58 |
| Ltb | 1,03E-51 | -1,0458144 | 0,856 | 0,933 | 1,91E-47 |
| Itgax | 9,12E-19 | -1,0585869 | 0,428 | 0,559 | 1,70E-14 |
| Evl | 4,00E-69 | -1,1211848 | 0,345 | 0,717 | 7,43E-65 |
| Klre1 | 6,20E-17 | -1,1255153 | 0,23 | 0,381 | 1,15E-12 |

| | | | | | |
|---|---|---|---|---|---|
| Ikzf2 | 8,11E-46 | -1,1258047 | 0,195 | 0,483 | 1,51E-41 |
| Cd7 | 2,06E-31 | -1,1936042 | 0,237 | 0,492 | 3,82E-27 |
| Spry2 | 2,77E-33 | -1,1965944 | 0,183 | 0,425 | 5,16E-29 |
| Crtam | 1,17E-35 | -1,2692148 | 0,458 | 0,701 | 2,17E-31 |
| Lta | 8,54E-33 | -1,2722138 | 0,257 | 0,499 | 1,59E-28 |
| Ier5l | 2,14E-48 | -1,2724427 | 0,24 | 0,548 | 3,98E-44 |
| Cd74 | 2,78E-49 | -1,3176406 | 0,15 | 0,443 | 5,17E-45 |
| Rgs2 | 2,61E-39 | -1,3715761 | 0,35 | 0,615 | 4,85E-35 |
| Itgae | 9,38E-98 | -1,9673502 | 0,116 | 0,531 | 1,74E-93 |
| Xcl1 | 4,35E-60 | -1,981446 | 0,24 | 0,608 | 8,08E-56 |

Cluster 3

| | p_val | avg_log2FC | pct.1 | pct.2 | p_val_adj |
|---|---|---|---|---|---|
| Ifit3 | 1.009399226 | 2,68995538 | 0,735 | 0,164 | 1,88E-197 |
| Gzmb | 5.045524247 | 2,1428604 | 0,933 | 0,437 | 9,38E-162 |
| Isg15 | 6.262011667 | 2,03025924 | 0,622 | 0,164 | 1,16E-131 |
| Ifi27l2a | 1.410630773 | 1,92551332 | 0,756 | 0,334 | 2,62E-123 |
| Usp18 | 3.219141524 | 1,86320189 | 0,643 | 0,212 | 5,98E-121 |
| Rnf213 | 1.246610785 | 1,80422157 | 0,919 | 0,79 | 2,32E-132 |
| Ccl3 | 2.771806445 | 1,69460028 | 0,48 | 0,176 | 5,15E-57 |
| Ifit1 | 1.148649774 | 1,60216287 | 0,523 | 0,112 | 2,13E-107 |
| Bst2 | 4.880662918 | 1,51841796 | 0,716 | 0,402 | 9,07E-87 |
| Ifit1bl1 | 2.349840368 | 1,48679868 | 0,722 | 0,433 | 4,37E-83 |
| Rtp4 | 1.377542026 | 1,44566741 | 0,726 | 0,353 | 2,56E-110 |
| Ifi208 | 4.480535748 | 1,41350823 | 0,607 | 0,18 | 8,33E-121 |
| AA467197 | 1.779425002 | 1,364499 | 0,491 | 0,208 | 3,31E-52 |
| Gzma | 7.294222922 | 1,33292698 | 0,72 | 0,273 | 1,36E-96 |
| Oas3 | 1.355006239 | 1,33050989 | 0,61 | 0,295 | 2,52E-74 |
| Rsad2 | 1.948438188 | 1,32545594 | 0,395 | 0,046 | 3,62E-94 |
| Irf7 | 2.254604532 | 1,30067431 | 0,793 | 0,534 | 4,19E-86 |
| Ifih1 | 6.588124790 | 1,27577479 | 0,529 | 0,203 | 1,22E-71 |
| Daxx | 1.449199807 | 1,26583823 | 0,666 | 0,447 | 2,69E-55 |
| Isg20 | 5.569273045 | 1,2496122 | 0,698 | 0,416 | 1,03E-65 |
| Slfn1 | 3.820733610 | 1,24230856 | 0,749 | 0,501 | 7,10E-74 |
| Dhx58 | 8.677529675 | 1,20121719 | 0,557 | 0,237 | 1,61E-71 |
| Ifit2 | 1.239676268 | 1,19834998 | 0,51 | 0,228 | 2,30E-53 |
| Zbp1 | 2.794287849 | 1,18385072 | 0,87 | 0,696 | 5,19E-91 |
| Xaf1 | 1.238462347 | 1,16152018 | 0,761 | 0,491 | 2,30E-80 |
| Ifi209 | 4.180216765 | 1,10252101 | 0,827 | 0,669 | 7,77E-70 |
| Slfn5 | 9.878292338 | 1,09667067 | 0,302 | 0,059 | 1,84E-53 |
| Trafd1 | 8.658961340 | 1,07120832 | 0,748 | 0,588 | 1,61E-54 |
| Pik3ap1 | 1.862393871 | 1,05109019 | 0,757 | 0,533 | 3,46E-55 |
| Ms4a4c | 1.076218500 | 1,03274295 | 0,417 | 0,189 | 2,00E-37 |
| Ly6a | 3.452566619 | 1,03071976 | 0,953 | 0,842 | 6,42E-93 |
| Phf11b | 4.158146587 | 1,00217684 | 0,657 | 0,396 | 7,73E-58 |
| Herc6 | 2.892274691 | 0,99837317 | 0,738 | 0,568 | 5,37E-45 |
| Parp9 | 7.348329948 | 0,98190266 | 0,68 | 0,466 | 1,37E-48 |
| Pml | 1.530748911 | 0,9811553 | 0,679 | 0,47 | 2,84E-49 |
| Plac8 | 1.153416855 | 0,98007892 | 0,834 | 0,611 | 2,14E-49 |
| Oas1a | 4.249344322 | 0,9731416 | 0,458 | 0,191 | 7,90E-49 |
| Phf11c | 1.274877217 | 0,94781471 | 0,608 | 0,351 | 2,37E-50 |
| Lag3 | 1.108339438 | 0,94640676 | 0,712 | 0,554 | 2,06E-24 |
| Tnfrsf9 | 8.599455687 | 0,94408345 | 0,325 | 0,141 | 1,60E-24 |
| Sema7a | 5.199511283 | 0,93741947 | 0,304 | 0,067 | 9,66E-49 |
| Lgals3bp | 3.155043964 | 0,92779621 | 0,894 | 0,807 | 5,86E-58 |
| Il2ra | 2.336225320 | 0,92350335 | 0,391 | 0,252 | 4,34E-12 |
| Pim1 | 5.109580366 | 0,91922188 | 0,777 | 0,628 | 9,50E-35 |
| Wdfy1 | 1.128978193 | 0,90578112 | 0,553 | 0,282 | 2,10E-48 |
| Cmpk2 | 1.279552981 | 0,89510749 | 0,335 | 0,082 | 2,38E-52 |
| Slfn8 | 9.757074327 | 0,89314725 | 0,719 | 0,505 | 1,81E-50 |
| Oasl2 | 2.518883260 | 0,88417101 | 0,264 | 0,05 | 4,68E-46 |
| Trim30d | 6.254055626 | 0,87945238 | 0,505 | 0,258 | 1,16E-43 |
| Sdc3 | 8.547805919 | 0,87765766 | 0,265 | 0,03 | 1,59E-57 |

Up in Sp140-/-
Up in Sp140+/+

| | | | | | |
|---|---|---|---|---|---|
| Id2 | 3.972002413 | 0,86224856 | 0,934 | 0,879 | 7,38E-51 |
| Il18rap | 1.233843452 | 0,86064975 | 0,879 | 0,78 | 2,29E-40 |
| Mxd1 | 4.609881207 | 0,85020416 | 0,707 | 0,594 | 8,57E-20 |
| Helz2 | 1.723497933 | 0,8474221 | 0,83 | 0,764 | 3,20E-47 |
| Dtx3l | 1.393988291 | 0,83989679 | 0,8 | 0,665 | 2,59E-45 |
| Gbp9 | 5.344327355 | 0,82999148 | 0,778 | 0,641 | 9,93E-44 |
| Ifit3b | 4.586447377 | 0,82808753 | 0,304 | 0,027 | 8,52E-74 |
| Irf8 | 1.176554380 | 0,82674521 | 0,345 | 0,18 | 2,19E-19 |
| Il10ra | 1.793405294 | 0,81464304 | 0,648 | 0,464 | 3,33E-28 |
| Trim30a | 8.526046303 | 0,80628794 | 0,65 | 0,46 | 1,58E-38 |
| Eif2ak2 | 1.909512334 | 0,79702529 | 0,525 | 0,324 | 3,55E-28 |
| Prf1 | 8.355478882 | 0,77369285 | 0,739 | 0,656 | 1,55E-12 |
| Ccl4 | 9.879092519 | 0,76842862 | 0,833 | 0,647 | 1,84E-25 |
| Bcl2l11 | 1.760978169 | 0,76739564 | 0,715 | 0,604 | 3,27E-17 |
| Rilpl2 | 4.895283303 | 0,7572198 | 0,432 | 0,245 | 9,10E-23 |
| Ddx60 | 2.559149886 | 0,75272295 | 0,484 | 0,249 | 4,76E-35 |
| Havcr2 | 3.557727498 | 0,7409288 | 0,37 | 0,208 | 6,61E-17 |
| Ifi211 | 4.212589654 | 0,73615782 | 0,262 | 0,059 | 7,83E-40 |
| Parp14 | 1.161914944 | 0,73595847 | 0,805 | 0,745 | 2,16E-31 |
| Parp12 | 5.451855477 | 0,73236267 | 0,426 | 0,247 | 1,01E-22 |
| Pkm | 4.112894781 | 0,72844397 | 0,679 | 0,586 | 7,64E-17 |
| H2-T24 | 1.392971431 | 0,71790389 | 0,491 | 0,293 | 2,59E-24 |
| Fkbp5 | 3.871757474 | 0,71095838 | 0,451 | 0,27 | 7,19E-22 |
| Cd86 | 1.683911014 | 0,70549242 | 0,462 | 0,266 | 3,13E-24 |
| Serpinb9 | 2.865741245 | 0,70470693 | 0,558 | 0,43 | 5,33E-13 |
| Stat2 | 1.761037434 | 0,70263778 | 0,546 | 0,384 | 3,27E-21 |
| Slc7a5 | 8.101997740 | 0,69188707 | 0,328 | 0,21 | 1,51E-08 |
| Acadl | 2.215017367 | 0,68585902 | 0,567 | 0,48 | 4,12E-09 |
| Ddx58 | 1.247739231 | 0,68152316 | 0,635 | 0,49 | 2,32E-25 |
| Naa20 | 4.433251206 | 0,68091521 | 0,593 | 0,436 | 8,24E-21 |
| Tnfrsf18 | 7.600863469 | 0,675203 | 0,52 | 0,404 | 1,41E-11 |
| Trim34a | 1.397820081 | 0,66893258 | 0,629 | 0,471 | 2,60E-23 |
| Sytl3 | 1.433278931 | 0,66507723 | 0,609 | 0,429 | 2,66E-25 |
| Gbp7 | 1.082874556 | 0,66287321 | 0,744 | 0,59 | 2,01E-31 |
| Ccr2 | 3.364852472 | 0,66076818 | 0,476 | 0,298 | 6,25E-18 |
| Rab27a | 3.843374945 | 0,66058771 | 0,733 | 0,578 | 7,14E-32 |
| Setbp1 | 2.598615588 | 0,6454765 | 0,489 | 0,331 | 4,83E-16 |
| Ifi206 | 5.145113895 | 0,63902106 | 0,671 | 0,502 | 9,56E-25 |
| Ifi214 | 6.727016594 | 0,63543599 | 0,544 | 0,352 | 1,25E-25 |
| Hif1a | 1.509982520 | 0,63289885 | 0,643 | 0,567 | 2,81E-13 |
| Casp4 | 3.334202764 | 0,62729441 | 0,561 | 0,408 | 6,20E-18 |
| Sh3bp2 | 1.865524655 | 0,62630247 | 0,384 | 0,209 | 3,47E-19 |
| Ifi35 | 2.124596291 | 0,62124687 | 0,66 | 0,506 | 3,95E-24 |
| Trim25 | 7.016590926 | 0,62089223 | 0,618 | 0,451 | 1,30E-21 |
| Phf11a | 1.953696551 | 0,61988239 | 0,327 | 0,103 | 3,63E-38 |
| Stx11 | 1.386383511 | 0,61065602 | 0,574 | 0,446 | 2,58E-12 |
| Arsb | 1.120135068 | 0,61003201 | 0,642 | 0,472 | 2,08E-22 |
| Ccnyl1 | 5.294806395 | 0,60120899 | 0,459 | 0,283 | 9,84E-19 |
| Casp3 | 4.126833495 | 0,59998579 | 0,547 | 0,405 | 7,67E-15 |
| Ptms | 7.128263168 | 0,59911013 | 0,654 | 0,519 | 1,32E-13 |
| Hsh2d | 2.395086916 | 0,59459886 | 0,377 | 0,173 | 4,45E-28 |
| Mapkapk2 | 5.270283783 | 0,5916774 | 0,706 | 0,591 | 9,79E-20 |
| Pdcd1 | 2.677082909 | 0,58806832 | 0,467 | 0,312 | 4,97E-11 |

| | | | | | |
|---|---|---|---|---|---|
| N4bp1 | 7.858992124 | 0,58764097 | 0,68 | 0,559 | 1,46E-17 |
| Smchd1 | 2.248993951 | 0,58311829 | 0,781 | 0,719 | 4,18E-20 |
| Nfkbia | 3.516904196 | 0,58215857 | 0,772 | 0,702 | 6,54E-10 |
| Gbp2 | 2.563282615 | 0,57605343 | 0,726 | 0,624 | 4,76E-14 |
| Il12rb1 | 1.043700867 | 0,57453317 | 0,691 | 0,587 | 1,94E-14 |
| Nt5c3 | 7.681256169 | 0,57051084 | 0,433 | 0,32 | 1,43E-09 |
| Nop53 | 6.494071325 | 0,56880903 | 0,818 | 0,716 | 1,21E-25 |
| Clic1 | 5.171276091 | 0,56709313 | 0,654 | 0,514 | 9,61E-13 |
| Tor3a | 1.862454696 | 0,5592521 | 0,345 | 0,182 | 3,46E-18 |
| Chmp4b | 9.115846442 | 0,55626875 | 0,588 | 0,453 | 1,69E-18 |
| Ctss | 3.947602377 | 0,55226757 | 0,682 | 0,567 | 7,34E-15 |
| Chd7 | 6.735551201 | 0,53861805 | 0,762 | 0,687 | 1,25E-17 |
| Plek | 7.104777849 | 0,53718186 | 0,605 | 0,462 | 1,32E-15 |
| Mif4gd | 1.077973672 | 0,53138661 | 0,667 | 0,591 | 2,00E-10 |
| Gpsm3 | 3.790432837 | 0,53063981 | 0,808 | 0,709 | 7,04E-19 |
| Coro1a | 4.760959254 | 0,52019002 | 0,975 | 0,968 | 8,85E-55 |
| Tmem184b | 5.743034267 | 0,51908908 | 0,477 | 0,359 | 1,07E-09 |
| Uba7 | 3.893852347 | 0,51706946 | 0,701 | 0,618 | 7,24E-13 |
| Dgkh | 3.004508987 | 0,51423312 | 0,282 | 0,113 | 5,58E-22 |
| Ddit4 | 8.232829572 | 0,51228151 | 0,544 | 0,4 | 1,53E-14 |
| Serpina3g | 6.820368114 | 0,50766245 | 0,727 | 0,648 | 1,27E-10 |
| Srgn | 5.737958743 | 0,50584583 | 0,936 | 0,873 | 1,07E-14 |
| Glrx | 6.721358354 | 0,50460732 | 0,535 | 0,451 | 1,25E-05 |
| Rnf114 | 1.976008886 | 0,50409291 | 0,713 | 0,599 | 3,67E-18 |
| Smpdl3b | 9.562312356 | 0,50171517 | 0,371 | 0,198 | 1,78E-17 |
| Numa1 | 1.874913192 | -0,501698 | 0,703 | 0,852 | 3,48E-32 |
| Emb | 3.742509528 | -0,5018884 | 0,784 | 0,849 | 6,95E-09 |
| Pde3b | 3.467783212 | -0,5032289 | 0,671 | 0,806 | 6,44E-25 |
| Chd3 | 8.506641335 | -0,5051942 | 0,607 | 0,77 | 1,58E-24 |
| Rarg | 1.654571333 | -0,5062977 | 0,154 | 0,332 | 3,07E-19 |
| Appl2 | 5.093402423 | -0,512481 | 0,195 | 0,384 | 9,47E-20 |
| Cmah | 4.216892454 | -0,5166915 | 0,093 | 0,264 | 7,84E-22 |
| Spsb1 | 1.928509025 | -0,5201897 | 0,198 | 0,402 | 3,58E-21 |
| Trat1 | 2.095787247 | -0,5212585 | 0,15 | 0,359 | 3,89E-25 |
| Slc20a1 | 3.486353257 | -0,5225956 | 0,716 | 0,85 | 6,48E-20 |
| Kif1b | 6.151231826 | -0,523025 | 0,249 | 0,448 | 1,14E-22 |
| Abcb1b | 3.792614949 | -0,5244296 | 0,249 | 0,4 | 7,05E-12 |
| Actn1 | 2.228235355 | -0,5256848 | 0,209 | 0,398 | 4,14E-19 |
| Nebl | 1.138810582 | -0,5282628 | 0,125 | 0,326 | 2,12E-24 |
| Itgax | 5.333553915 | -0,5321354 | 0,369 | 0,505 | 9,91E-10 |
| Tspan13 | 5.953856415 | -0,5332689 | 0,242 | 0,452 | 1,11E-25 |
| Srsf7 | 3.296020295 | -0,5359686 | 0,683 | 0,836 | 6,12E-31 |
| Sesn3 | 5.410544437 | -0,5376394 | 0,522 | 0,72 | 1,01E-27 |
| Il27ra | 9.297341122 | -0,5414305 | 0,662 | 0,812 | 1,73E-27 |
| Ern1 | 1.198810640 | -0,5423472 | 0,403 | 0,502 | 0,00222775 |
| Snx29 | 3.912840364 | -0,5442746 | 0,331 | 0,524 | 7,27E-19 |
| Ikzf2 | 5.438240220 | -0,544486 | 0,205 | 0,336 | 1,01E-08 |
| Lpar6 | 2.632591339 | -0,5515227 | 0,292 | 0,505 | 4,89E-22 |
| Nr3c1 | 1.918492309 | -0,5517682 | 0,397 | 0,603 | 3,57E-24 |
| S1pr1 | 4.171786690 | -0,5525776 | 0,217 | 0,337 | 7,75E-08 |
| Rgcc | 2.715026252 | -0,5664535 | 0,106 | 0,275 | 5,05E-19 |
| Gpr183 | 6.581683977 | -0,5710598 | 0,532 | 0,691 | 1,22E-17 |
| Dnajb1 | 4.853464388 | -0,5811453 | 0,335 | 0,467 | 9,02E-09 |

| | | | | | |
|---|---|---|---|---|---|
| Arrdc3 | 2.898295267 | -0,5888682 | 0,249 | 0,423 | 5,39E-16 |
| Dnaja1 | 3.483559527 | -0,5900256 | 0,551 | 0,708 | 6,47E-25 |
| Itgb7 | 5.061285538 | -0,5917949 | 0,85 | 0,95 | 9,41E-43 |
| Itgb1 | 8.787693908 | -0,6024154 | 0,595 | 0,748 | 1,63E-17 |
| Sgms1 | 2.716822209 | -0,6065153 | 0,263 | 0,507 | 5,05E-30 |
| Ptger4 | 1.661433211 | -0,6167506 | 0,48 | 0,69 | 3,09E-22 |
| Gpr132 | 2.104155472 | -0,6185243 | 0,667 | 0,811 | 3,91E-25 |
| Faah | 1.323084682 | -0,6201541 | 0,312 | 0,567 | 2,46E-33 |
| Eif4a2 | 3.649785939 | -0,6238336 | 0,817 | 0,929 | 6,78E-58 |
| Rgs2 | 1.055355328 | -0,6251483 | 0,259 | 0,412 | 1,96E-11 |
| Kctd12 | 7.512696876 | -0,6275393 | 0,193 | 0,414 | 1,40E-26 |
| Lrig1 | 2.186680019 | -0,6307858 | 0,165 | 0,448 | 4,06E-41 |
| St8sia4 | 2.043380220 | -0,6310129 | 0,363 | 0,589 | 3,80E-26 |
| Sik1 | 7.448606343 | -0,6522349 | 0,485 | 0,672 | 1,38E-21 |
| Arnt2 | 2.093701996 | -0,6528491 | 0,174 | 0,434 | 3,89E-35 |
| Tcrg-C2 | 6.793555324 | -0,6543105 | 0,173 | 0,392 | 1,26E-24 |
| Sntb1 | 1.700221496 | -0,6553822 | 0,39 | 0,673 | 3,16E-43 |
| Kcnc1 | 5.146706917 | -0,6570638 | 0,122 | 0,349 | 9,56E-32 |
| Eomes | 3.932536845 | -0,660429 | 0,179 | 0,312 | 7,31E-10 |
| Jmy | 9.526990408 | -0,6740531 | 0,374 | 0,592 | 1,77E-28 |
| Slc12a7 | 2.421151035 | -0,6814976 | 0,46 | 0,709 | 4,50E-42 |
| Tmem181a | 3.831382852 | -0,6885304 | 0,205 | 0,509 | 7,12E-48 |
| Ppp2r2c | 1.120115550 | -0,6889346 | 0,236 | 0,48 | 2,08E-30 |
| Cd244a | 1.126398830 | -0,690378 | 0,114 | 0,271 | 2,09E-17 |
| Spry2 | 1.155739482 | -0,7041218 | 0,135 | 0,29 | 2,15E-14 |
| Ddx5 | 1.620280022 | -0,71527 | 0,783 | 0,911 | 3,01E-71 |
| P2ry10 | 3.277981108 | -0,7222934 | 0,672 | 0,822 | 6,09E-38 |
| Cxcr6 | 8.543947102 | -0,73273 | 0,719 | 0,87 | 1,59E-33 |
| Vps37b | 2.363383398 | -0,7596978 | 0,759 | 0,845 | 4,39E-24 |
| Klf2 | 4.683961442 | -0,7606208 | 0,385 | 0,513 | 8,70E-10 |
| Klre1 | 1.319684435 | -0,7806723 | 0,139 | 0,277 | 2,45E-12 |
| Tcf7 | 2.151739241 | -0,8206077 | 0,284 | 0,528 | 4,00E-32 |
| Klf6 | 1.076300593 | -0,8509512 | 0,837 | 0,938 | 2,00E-51 |
| Sell | 3.209070381 | -0,8711903 | 0,175 | 0,32 | 5,96E-12 |
| Neurl3 | 2.648158841 | -0,8737167 | 0,522 | 0,691 | 4,92E-24 |
| Ifi27 | 1.742906810 | -0,8962947 | 0,371 | 0,718 | 3,24E-69 |
| Ldlrad4 | 2.640240340 | -0,9081205 | 0,355 | 0,662 | 4,91E-53 |
| Hsph1 | 8.841228554 | -0,9573631 | 0,267 | 0,568 | 1,64E-51 |
| Itga1 | 3.354133296 | -0,9925829 | 0,419 | 0,693 | 6,23E-48 |
| Cdh1 | 1.663086508 | -1,0231351 | 0,215 | 0,538 | 3,09E-52 |
| Ccr7 | 1.046581159 | -1,0252519 | 0,073 | 0,271 | 1,94E-29 |
| Ier5l | 8.461835029 | -1,0773052 | 0,192 | 0,425 | 1,57E-31 |
| Rhob | 1.754139526 | -1,1260504 | 0,16 | 0,403 | 3,26E-36 |
| Jun | 2.504676436 | -1,1793912 | 0,498 | 0,707 | 4,65E-37 |
| Itgae | 1.868025706 | -1,4142687 | 0,275 | 0,591 | 3,47E-55 |

| Cluster 4 | | | | | |
|---|---|---|---|---|---|
| | p_val | avg_log2FC | pct.1 | pct.2 | p_val_adj |
| Gzmb | 2.03713316( | 2,41245706 | 0,77 | 0,157 | 3,79E-168 |
| Ifit3 | 5.49156713( | 1,8331169 | 0,693 | 0,294 | 1,02E-78 |
| Isg15 | 1.07828436( | 1,55900833 | 0,63 | 0,285 | 2,00E-59 |
| Ifi27l2a | 1.30130761: | 1,32693619 | 0,87 | 0,663 | 2,42E-50 |
| Isg20 | 3.735222181 | 1,2951772 | 0,563 | 0,322 | 6,94E-32 |
| Ccl5 | 1.580010994 | 1,19596765 | 0,866 | 0,724 | 2,94E-20 |
| Gzma | 1.19081450ξ | 1,18239558 | 0,577 | 0,184 | 2,21E-63 |
| Ifit1 | 4.92568255ξ | 1,08857794 | 0,417 | 0,122 | 9,15E-53 |
| Irf7 | 2.994895332 | 1,07194186 | 0,785 | 0,574 | 5,57E-41 |
| Rnf213 | 5.01532227( | 1,07027494 | 0,931 | 0,882 | 9,32E-26 |
| Oas3 | 5.33946049ξ | 1,02212156 | 0,685 | 0,464 | 9,92E-33 |
| Plac8 | 4.10436486( | 1,0138978 | 0,894 | 0,818 | 7,63E-29 |
| Slfn5 | 3.13951916( | 1,00515015 | 0,352 | 0,111 | 5,83E-37 |
| Lag3 | 2.85105708ξ | 0,96486818 | 0,394 | 0,13 | 5,30E-38 |
| Ly6a | 1.00725048( | 0,90604852 | 0,965 | 0,834 | 1,87E-42 |
| Rsad2 | 1.483043552 | 0,89142697 | 0,27 | 0,031 | 2,76E-61 |
| Oas2 | 4.63254684ξ | 0,88281257 | 0,358 | 0,142 | 8,61E-27 |
| Rbm3 | 1.758003057 | 0,88122994 | 0,907 | 0,727 | 3,27E-73 |
| Ifi208 | 3.42404749ξ | 0,86935894 | 0,715 | 0,399 | 6,36E-48 |
| Bhlhe40 | 7.231623761 | 0,85467645 | 0,496 | 0,269 | 1,34E-20 |
| Ccl4 | 1.531267328 | 0,81724681 | 0,571 | 0,237 | 2,85E-42 |
| Rtp4 | 4.257606922 | 0,81277525 | 0,776 | 0,534 | 7,91E-36 |
| Usp18 | 1.34451817: | 0,80702901 | 0,632 | 0,449 | 2,50E-16 |
| Dhx58 | 2.124515587 | 0,79643591 | 0,488 | 0,232 | 3,95E-31 |
| Zbp1 | 1.61256614ξ | 0,78678296 | 0,925 | 0,84 | 3,00E-28 |
| Oasl2 | 1.84720615: | 0,77003553 | 0,396 | 0,169 | 3,43E-26 |
| Lgals3bp | 4.83720091ξ | 0,76817826 | 0,909 | 0,83 | 8,99E-32 |
| Klrk1 | 6.92515326ξ | 0,76721872 | 0,307 | 0,156 | 1,29E-10 |
| Oas1a | 5.631828272 | 0,76036817 | 0,425 | 0,179 | 1,05E-30 |
| Phf11b | 2.64118202( | 0,75845786 | 0,713 | 0,471 | 4,91E-30 |
| Herc6 | 1.41576976( | 0,74519125 | 0,677 | 0,53 | 2,63E-14 |
| AA467197 | 1.02126501: | 0,74230193 | 0,256 | 0,05 | 1,90E-39 |
| Id2 | 1.355369451 | 0,73198263 | 0,781 | 0,608 | 2,52E-17 |
| Xaf1 | 1.28914901( | 0,73027238 | 0,699 | 0,501 | 2,40E-24 |
| Ifit1bl1 | 4.661691601 | 0,72199685 | 0,699 | 0,575 | 8,66E-12 |
| Serpina3g | 5.90365534: | 0,71995003 | 0,417 | 0,242 | 1,10E-12 |
| Fkbp5 | 8.01248468ξ | 0,7148374 | 0,518 | 0,276 | 1,49E-27 |
| Daxx | 3.95584663ξ | 0,71098366 | 0,764 | 0,631 | 7,35E-16 |
| S100a6 | 2.10232611( | 0,69039347 | 0,372 | 0,145 | 3,91E-27 |
| Fgl2 | 4.48311198: | 0,68500644 | 0,297 | 0,151 | 8,33E-11 |
| Jaml | 3.758973827 | 0,67506725 | 0,683 | 0,527 | 6,99E-16 |
| Bst2 | 4.128023667 | 0,64312984 | 0,917 | 0,829 | 7,67E-17 |
| Trafd1 | 6.395578182 | 0,63359902 | 0,614 | 0,501 | 1,19E-08 |
| Slfn1 | 2.876109924 | 0,63220367 | 0,854 | 0,753 | 5,34E-16 |
| Anxa2 | 1.23339748: | 0,6302433 | 0,51 | 0,28 | 2,29E-19 |
| Ifng | 3.87771356: | 0,61314796 | 0,455 | 0,142 | 7,21E-48 |
| Phf11c | 5.15364638: | 0,59230914 | 0,632 | 0,406 | 9,58E-22 |
| Ifih1 | 3.741196511 | 0,5922943 | 0,429 | 0,231 | 6,95E-19 |
| Ide | 4.19970317: | 0,58703113 | 0,724 | 0,499 | 7,80E-29 |
| Ifit2 | 1.56412110ξ | 0,58696385 | 0,378 | 0,219 | 2,91E-11 |

Up in Sp140-/-

Up in Sp140+/+

| | | | | | |
|---|---|---|---|---|---|
| Klrc1 | 2.048235938 | 0,58021208 | 0,437 | 0,266 | 3,81E-08 |
| Ccl3 | 9.457468860 | 0,5698363 | 0,333 | 0,054 | 1,76E-61 |
| Ddx60 | 1.061310043 | 0,56592963 | 0,437 | 0,257 | 1,97E-14 |
| Pml | 3.197949226 | 0,56024535 | 0,715 | 0,546 | 5,94E-14 |
| Coro2a | 1.945439806 | 0,55653419 | 0,427 | 0,249 | 3,62E-13 |
| Ptms | 3.928003315 | 0,54005749 | 0,38 | 0,22 | 7,30E-10 |
| Pik3ap1 | 7.336721749 | 0,5369625 | 0,331 | 0,181 | 1,36E-10 |
| mt-Nd5 | 4.354875312 | 0,52084312 | 0,949 | 0,879 | 8,09E-17 |
| Il18rap | 4.402353698 | 0,51884569 | 0,555 | 0,398 | 8,18E-11 |
| Helz2 | 1.076914213 | 0,51257041 | 0,829 | 0,763 | 2,00E-07 |
| mt-Atp8 | 2.088414193 | 0,50421512 | 0,998 | 0,994 | 3,88E-18 |
| Oasl1 | 8.961953943 | 0,50389875 | 0,268 | 0,072 | 1,67E-30 |
| Serpinb9 | 2.574215598 | 0,50360156 | 0,435 | 0,335 | 0,00047837 |
| mt-Co2 | 2.921039382 | 0,50115449 | 0,998 | 0,986 | 5,43E-18 |
| Cmpk2 | 2.542939443 | 0,50043974 | 0,295 | 0,094 | 4,73E-27 |
| Macf1 | 1.460619192 | -0,500649 | 0,933 | 0,972 | 2,71E-32 |
| Smad7 | 5.868761959 | -0,507296 | 0,748 | 0,843 | 1,09E-13 |
| Gbp8 | 1.278700734 | -0,5217985 | 0,657 | 0,776 | 2,38E-10 |
| Sik1 | 8.481890194 | -0,5455553 | 0,496 | 0,624 | 1,58E-08 |
| Hist1h1e | 2.242911756 | -0,5462946 | 0,665 | 0,729 | 4,17E-05 |
| Il6st | 1.764537510 | -0,5493065 | 0,807 | 0,917 | 3,28E-23 |
| Dapl1 | 6.602394629 | -0,577463 | 0,283 | 0,436 | 1,23E-06 |
| Jun | 5.289505748 | -0,5868737 | 0,754 | 0,836 | 9,83E-09 |
| Ccr7 | 6.101787262 | -0,5888072 | 0,854 | 0,923 | 1,13E-12 |
| Smc4 | 7.853362183 | -0,5910229 | 0,774 | 0,855 | 1,46E-16 |
| S1pr1 | 1.078445948 | -0,5912454 | 0,844 | 0,929 | 2,00E-24 |
| Ifi27 | 9.623257300 | -0,6038365 | 0,652 | 0,841 | 1,79E-27 |
| Klf2 | 9.875938843 | -0,6065083 | 0,945 | 0,984 | 1,84E-24 |
| Dnajb1 | 7.340864894 | -0,6139096 | 0,545 | 0,695 | 1,36E-12 |
| Chd3 | 5.068623263 | -0,6185529 | 0,685 | 0,862 | 9,42E-32 |
| Sp140 | 3.985116355 | -0,6266228 | 0,287 | 0,558 | 7,41E-30 |
| Tdrp | 6.953980151 | -0,6286818 | 0,374 | 0,576 | 1,29E-17 |
| Klf6 | 9.095310119 | -0,6496914 | 0,85 | 0,948 | 1,69E-26 |
| Ighm | 2.153673743 | -0,6634825 | 0,817 | 0,923 | 4,00E-29 |
| Cd28 | 2.269767815 | -0,7075624 | 0,604 | 0,763 | 4,22E-21 |
| Tagap | 2.759526370 | -0,7223133 | 0,732 | 0,88 | 5,13E-24 |
| Hsph1 | 8.156423186 | -0,7244056 | 0,352 | 0,609 | 1,52E-28 |
| Vps37b | 9.815123980 | -0,8084528 | 0,931 | 0,97 | 1,82E-35 |
| Fos | 7.630540718 | -0,8922604 | 0,547 | 0,712 | 1,42E-16 |
| Dusp10 | 2.715830617 | -0,8982911 | 0,555 | 0,694 | 5,05E-16 |
| Rgcc | 1.750522061 | -0,953254 | 0,209 | 0,397 | 3,25E-14 |

Cluster 5

| | p_val | avg_log2FC | pct.1 | pct.2 | p_val_adj |
|---|---|---|---|---|---|
| Ifit3 | 9.717589314 | 2,67262582 | 0,998 | 0,503 | 1,81E-203 |
| Slfn5 | 6.744625509 | 2,29662586 | 0,75 | 0,126 | 1,25E-115 |
| Isg15 | 2.935519401 | 2,23841078 | 0,987 | 0,491 | 5,46E-172 |
| Ifi208 | 3.858697118 | 2,08865631 | 0,942 | 0,27 | 7,17E-184 |
| Ifit1 | 3.691607915 | 2,06617342 | 0,976 | 0,371 | 6,86E-162 |
| Ifit2 | 1.197271888 | 2,06246937 | 0,875 | 0,377 | 2,22E-119 |
| Ms4a4c | 3.298751663 | 2,05568832 | 0,684 | 0,167 | 6,13E-89 |
| Usp18 | 2.401479391 | 2,03375376 | 0,969 | 0,367 | 4,46E-163 |
| Ifi211 | 1.003068675 | 1,96744079 | 0,658 | 0,107 | 1,86E-91 |
| Rsad2 | 9.570886455 | 1,95070553 | 0,89 | 0,21 | 1,78E-132 |
| Daxx | 1.526341618 | 1,92036148 | 0,98 | 0,592 | 2,84E-164 |
| Gzmb | 1.456788079 | 1,82532973 | 0,966 | 0,548 | 2,71E-87 |
| Ifit1bl1 | 2.144482085 | 1,79759788 | 0,991 | 0,786 | 3,99E-155 |
| Phf11b | 4.655665382 | 1,78196078 | 0,982 | 0,682 | 8,65E-157 |
| Bst2 | 1.209198653 | 1,75837883 | 0,995 | 0,769 | 2,25E-144 |
| Cxcl10 | 1.463282617 | 1,74433874 | 0,498 | 0,122 | 2,72E-43 |
| Ifi27l2a | 3.282014612 | 1,74002946 | 0,898 | 0,68 | 6,10E-76 |
| Isg20 | 1.940636064 | 1,7374069 | 0,982 | 0,693 | 3,61E-137 |
| Oasl2 | 8.402753597 | 1,71908917 | 0,658 | 0,043 | 1,56E-104 |
| Pik3ap1 | 1.101806224 | 1,68121303 | 0,945 | 0,542 | 2,05E-123 |
| Sdc3 | 5.550100670 | 1,6686175 | 0,712 | 0,107 | 1,03E-103 |
| Plac8 | 1.807806971 | 1,65186765 | 0,969 | 0,695 | 3,36E-101 |
| Oas1a | 1.188076920 | 1,65183999 | 0,746 | 0,249 | 2,21E-89 |
| Ifih1 | 5.264269019 | 1,64303625 | 0,914 | 0,359 | 9,78E-129 |
| Oas3 | 1.605343135 | 1,61834441 | 0,97 | 0,515 | 2,98E-139 |
| Dhx58 | 2.678406171 | 1,61183381 | 0,938 | 0,396 | 4,98E-137 |
| Ifit3b | 5.457252092 | 1,58266883 | 0,815 | 0,159 | 1,01E-124 |
| Rnf213 | 1.249635599 | 1,57595806 | 0,997 | 0,849 | 2,32E-147 |
| Cmpk2 | 2.624003028 | 1,51834514 | 0,838 | 0,252 | 4,88E-103 |
| Irf7 | 2.275500336 | 1,50693245 | 0,987 | 0,672 | 4,23E-140 |
| Nt5c3 | 9.900166590 | 1,50469718 | 0,816 | 0,452 | 1,84E-74 |
| Ifitm3 | 1.658095783 | 1,4926843 | 0,298 | 0,021 | 3,08E-32 |
| Ccrl2 | 8.830795899 | 1,46457726 | 0,606 | 0,085 | 1,64E-79 |
| Slfn1 | 4.492977593 | 1,45690928 | 0,987 | 0,718 | 8,35E-138 |
| Ube2l6 | 1.849079953 | 1,38495954 | 0,566 | 0,109 | 3,44E-63 |
| Phf11a | 2.464630237 | 1,37955488 | 0,799 | 0,249 | 4,58E-101 |
| Cd86 | 1.783797999 | 1,37515843 | 0,779 | 0,32 | 3,31E-76 |
| Trafd1 | 4.437839964 | 1,35972695 | 0,968 | 0,647 | 8,25E-120 |
| Ifi209 | 8.314476384 | 1,35802443 | 0,997 | 0,858 | 1,55E-141 |
| Phf11c | 3.897760831 | 1,34792366 | 0,908 | 0,511 | 7,24E-107 |
| AA467197 | 1.016564526 | 1,33766566 | 0,461 | 0,138 | 1,89E-35 |
| Ifi204 | 3.056766816 | 1,33502085 | 0,384 | 0,019 | 5,68E-48 |
| Oas2 | 7.603416801 | 1,30563908 | 0,497 | 0,033 | 1,41E-65 |
| Lgals9 | 1.543509634 | 1,28703535 | 0,935 | 0,635 | 2,87E-95 |
| Lgals3bp | 7.117845059 | 1,28636262 | 0,993 | 0,849 | 1,32E-131 |
| Slfn8 | 1.643148722 | 1,25894117 | 0,949 | 0,542 | 3,05E-105 |
| Herc6 | 2.803922373 | 1,24931384 | 0,949 | 0,567 | 5,21E-106 |
| Rtp4 | 2.722682817 | 1,2258379 | 0,98 | 0,625 | 5,06E-111 |
| Parp12 | 1.450920404 | 1,20376798 | 0,761 | 0,278 | 2,70E-80 |
| Xaf1 | 3.771830469 | 1,19135496 | 0,978 | 0,678 | 7,01E-118 |

Up in Sp140-/-
Up in Sp140+/+

| | | | | | |
|---|---|---|---|---|---|
| Helz2 | 1.211757592 | 1,17615085 | 0,982 | 0,765 | 2,25E-107 |
| Cd69 | 1.520198563 | 1,15508419 | 0,905 | 0,701 | 2,82E-37 |
| Pml | 3.598352039 | 1,11516491 | 0,951 | 0,598 | 6,69E-96 |
| Zbp1 | 1.519385158 | 1,10348101 | 0,998 | 0,913 | 2,82E-124 |
| Oasl1 | 1.233707807 | 1,09776278 | 0,442 | 0,033 | 2,29E-55 |
| Iigp1 | 3.988239667 | 1,09526948 | 0,519 | 0,221 | 7,41E-31 |
| Tor3a | 6.820456047 | 1,09487677 | 0,721 | 0,258 | 1,27E-68 |
| Ddx60 | 4.743330657 | 1,08849664 | 0,694 | 0,247 | 8,81E-60 |
| Smchd1 | 1.462119701 | 1,0742055 | 0,941 | 0,715 | 2,72E-75 |
| Parp9 | 5.268001782 | 1,06646466 | 0,96 | 0,658 | 9,79E-93 |
| Ifi214 | 5.592757791 | 1,05203781 | 0,891 | 0,447 | 1,04E-82 |
| Tmem184b | 1.827771355 | 1,03384591 | 0,808 | 0,408 | 3,40E-64 |
| Gbp2 | 2.190108723 | 1,01063365 | 0,984 | 0,893 | 4,07E-63 |
| Gbp9 | 1.572977928 | 1,00953691 | 0,965 | 0,753 | 2,92E-77 |
| Dtx3l | 2.176160357 | 1,00829405 | 0,989 | 0,858 | 4,04E-100 |
| Naa20 | 3.708015882 | 1,00730242 | 0,856 | 0,534 | 6,89E-62 |
| Samhd1 | 9.384928002 | 1,00224075 | 0,996 | 0,951 | 1,74E-70 |
| Eif2ak2 | 1.642049043 | 0,9918348 | 0,788 | 0,361 | 3,05E-64 |
| Mxd1 | 3.944774147 | 0,99180579 | 0,889 | 0,645 | 7,33E-40 |
| Slco3a1 | 3.562365438 | 0,98734713 | 0,906 | 0,596 | 6,62E-63 |
| Parp14 | 2.735642415 | 0,96470944 | 0,979 | 0,802 | 5,08E-76 |
| Ddx58 | 3.917894544 | 0,95786804 | 0,941 | 0,647 | 7,28E-75 |
| B4galt5 | 1.652716998 | 0,95001519 | 0,896 | 0,608 | 3,07E-59 |
| Dpp4 | 1.658320891 | 0,94386683 | 0,71 | 0,396 | 3,08E-40 |
| Bcl2l11 | 3.052640388 | 0,93586398 | 0,764 | 0,546 | 5,67E-25 |
| Il18rap | 1.744065659 | 0,93509723 | 0,903 | 0,647 | 3,24E-46 |
| Znfx1 | 1.640943893 | 0,91207905 | 0,77 | 0,419 | 3,05E-49 |
| H2-T24 | 1.507066238 | 0,90489637 | 0,751 | 0,41 | 2,80E-42 |
| Clic4 | 6.976659709 | 0,90266772 | 0,785 | 0,476 | 1,30E-45 |
| Jaml | 8.183919168 | 0,90096079 | 0,957 | 0,645 | 1,52E-57 |
| Mndal | 7.107721074 | 0,89305499 | 0,987 | 0,858 | 1,32E-82 |
| Ifi206 | 7.386919439 | 0,86675867 | 0,963 | 0,718 | 1,37E-66 |
| Ctss | 8.812691508 | 0,86613948 | 0,951 | 0,759 | 1,64E-63 |
| Pcgf5 | 7.287084903 | 0,86369073 | 0,713 | 0,346 | 1,35E-41 |
| Trim30a | 3.247810489 | 0,86167777 | 0,907 | 0,608 | 6,04E-66 |
| Stat2 | 3.005884712 | 0,86010751 | 0,822 | 0,449 | 5,59E-55 |
| Tor1aip2 | 2.599219639 | 0,85935172 | 0,869 | 0,606 | 4,83E-46 |
| Il12rb2 | 1.296663907 | 0,85776023 | 0,715 | 0,375 | 2,41E-31 |
| Evi2a | 1.807915518 | 0,85644855 | 0,619 | 0,299 | 3,36E-35 |
| Ifi35 | 2.292124151 | 0,85098593 | 0,907 | 0,606 | 4,26E-61 |
| Gypc | 2.516814561 | 0,84932927 | 0,466 | 0,134 | 4,68E-36 |
| Uba7 | 1.565109978 | 0,84272165 | 0,847 | 0,586 | 2,91E-47 |
| Usp25 | 8.083592710 | 0,84108071 | 0,982 | 0,862 | 1,50E-70 |
| Epsti1 | 6.671981750 | 0,83870987 | 0,995 | 0,934 | 1,24E-75 |
| Csrp1 | 1.081150576 | 0,83264731 | 0,621 | 0,324 | 2,01E-29 |
| Slc25a22 | 1.938897567 | 0,82749024 | 0,685 | 0,297 | 3,60E-45 |
| Zup1 | 3.203364506 | 0,82731832 | 0,807 | 0,499 | 5,95E-44 |
| Chmp4b | 6.595194794 | 0,82368011 | 0,906 | 0,658 | 1,23E-59 |
| Ifi203 | 4.387931692 | 0,81906613 | 0,978 | 0,874 | 8,15E-64 |
| Setdb2 | 4.139155972 | 0,81528065 | 0,594 | 0,287 | 7,69E-31 |
| Adgrg5 | 7.615004470 | 0,81123471 | 0,865 | 0,577 | 1,42E-39 |
| Lag3 | 5.319070446 | 0,80435384 | 0,744 | 0,54 | 9,88E-19 |
| Sh3bp2 | 1.022324511 | 0,7989509 | 0,6 | 0,198 | 1,90E-43 |

| | | | | | |
|---|---|---|---|---|---|
| Ccl3 | 3.977576411 | 0,79580368 | 0,404 | 0,159 | 7,39E-17 |
| Etnk1 | 1.17069132[illegible] | 0,79363808 | 0,959 | 0,804 | 2,18E-54 |
| Samd9l | 1.844571334 | 0,79115267 | 0,9 | 0,639 | 3,43E-45 |
| Ly6a | 4.158396087 | 0,78794203 | 0,999 | 0,942 | 7,73E-64 |
| Casp4 | 1.155255282 | 0,78166878 | 0,804 | 0,592 | 2,15E-33 |
| Slfn3 | 2.25590700[illegible] | 0,77637908 | 0,378 | 0,045 | 4,19E-39 |
| Trim25 | 5.412341077 | 0,7760407 | 0,868 | 0,538 | 1,01E-49 |
| Gbp7 | 3.04340230[illegible] | 0,77465347 | 0,974 | 0,858 | 5,66E-60 |
| Tor1aip1 | 7.67969380[illegible] | 0,77084624 | 0,96 | 0,852 | 1,43E-56 |
| Hif1a | 9.27693198[illegible] | 0,7699841 | 0,819 | 0,649 | 1,72E-24 |
| Trim30d | 2.268859537 | 0,76411558 | 0,702 | 0,33 | 4,22E-41 |
| Ifi47 | 1.87102440[illegible] | 0,76402622 | 0,99 | 0,899 | 3,48E-61 |
| Trim30c | 6.191546011 | 0,76160826 | 0,268 | 0,035 | 1,15E-23 |
| Mov10 | 1.10244760[illegible] | 0,75854977 | 0,772 | 0,466 | 2,05E-37 |
| N4bp1 | 3.441827731 | 0,75445919 | 0,807 | 0,563 | 6,40E-36 |
| Atp10a | 5.74803960[illegible] | 0,74430691 | 0,749 | 0,427 | 1,07E-36 |
| Nfkbid | 5.468314507 | 0,7395494 | 0,409 | 0,206 | 1,02E-12 |
| Cnp | 4.722608201 | 0,7380357 | 0,877 | 0,674 | 8,78E-40 |
| Sytl3 | 8.21069681[illegible] | 0,73551219 | 0,626 | 0,293 | 1,53E-33 |
| Wdfy1 | 2.28838700[illegible] | 0,73375285 | 0,595 | 0,268 | 4,25E-32 |
| Trim21 | 8.749631704 | 0,72759997 | 0,805 | 0,538 | 1,63E-33 |
| Irgm1 | 2.081084582 | 0,72144163 | 0,962 | 0,773 | 3,87E-46 |
| Sco1 | 1.02323551[illegible] | 0,71865366 | 0,38 | 0,124 | 1,90E-23 |
| Ms4a4b | 5.46422806[illegible] | 0,71566394 | 0,999 | 0,994 | 1,02E-84 |
| Rbm3 | 8.952028611 | 0,70852941 | 0,841 | 0,647 | 1,66E-36 |
| Lamc1 | 1.030869064 | 0,70775208 | 0,618 | 0,326 | 1,92E-27 |
| Satb1 | 3.39965561[illegible] | 0,70653735 | 0,866 | 0,588 | 6,32E-31 |
| Nampt | 3.56319896[illegible] | 0,70582307 | 0,689 | 0,369 | 6,62E-33 |
| Tgtp1 | 2.404258394 | 0,70550917 | 0,619 | 0,299 | 4,47E-32 |
| Icos | 4.37119525[illegible] | 0,70008395 | 0,885 | 0,678 | 8,12E-21 |
| Il12rb1 | 2.18557262[illegible] | 0,69846308 | 0,766 | 0,52 | 4,06E-24 |
| Ap1s3 | 3.07525844[illegible] | 0,69821225 | 0,503 | 0,181 | 5,71E-33 |
| Ccnyl1 | 7.134119927 | 0,69530636 | 0,55 | 0,268 | 1,33E-23 |
| Tapbp | 2.661384004 | 0,68804363 | 0,995 | 0,965 | 4,95E-63 |
| Grina | 8.857490307 | 0,68429761 | 0,923 | 0,771 | 1,65E-33 |
| Il2ra | 1.04797790[illegible] | 0,68316646 | 0,308 | 0,13 | 1,95E-10 |
| Npc2 | 4.76483827[illegible] | 0,68234099 | 0,977 | 0,953 | 8,85E-39 |
| Adar | 1.51841870[illegible] | 0,68097897 | 0,925 | 0,742 | 2,82E-46 |
| Xdh | 1.693938757 | 0,67941643 | 0,408 | 0,198 | 3,15E-14 |
| Nmi | 5.598985374 | 0,67704946 | 0,877 | 0,682 | 1,04E-33 |
| Ms4a6d | 1.358903814 | 0,66962765 | 0,516 | 0,219 | 2,53E-25 |
| Ly6e | 2.187051254 | 0,66515358 | 0,999 | 0,984 | 4,06E-61 |
| Il18bp | 3.18566254[illegible] | 0,66466008 | 0,376 | 0,126 | 5,92E-22 |
| Herc3 | 3.282450427 | 0,66047366 | 0,835 | 0,573 | 6,10E-36 |
| Resf1 | 1.240843037 | 0,65109527 | 0,804 | 0,563 | 2,31E-26 |
| Keap1 | 7.84959760[illegible] | 0,64576445 | 0,69 | 0,4 | 1,46E-29 |
| Pttg1 | 1.13223281[illegible] | 0,63897992 | 0,841 | 0,695 | 2,10E-23 |
| Ncoa7 | 1.306338117 | 0,63800015 | 0,497 | 0,241 | 2,43E-20 |
| Hsh2d | 8.973674954 | 0,63693086 | 0,619 | 0,295 | 1,67E-30 |
| Nfkbia | 9.342042984 | 0,63575803 | 0,836 | 0,697 | 1,74E-10 |
| Tspo | 3.825373977 | 0,63503929 | 0,933 | 0,837 | 7,11E-33 |
| BC147527 | 7.81880532[illegible] | 0,62891824 | 0,55 | 0,243 | 1,45E-28 |
| Gpr171 | 5.446328001 | 0,62878108 | 0,952 | 0,833 | 1,01E-28 |

| | | | | | |
|---|---|---|---|---|---|
| Cdkl3 | 2.690517164 | 0,62590741 | 0,48 | 0,202 | 5,00E-24 |
| Acadl | 1.191004205 | 0,62544793 | 0,71 | 0,619 | 2,21E-07 |
| Tnfrsf9 | 2.809497292 | 0,62510031 | 0,264 | 0,097 | 5,22E-10 |
| Casp3 | 2.666791433 | 0,6073513 | 0,692 | 0,48 | 4,96E-18 |
| Id2 | 1.631251670 | 0,60323256 | 0,965 | 0,895 | 3,03E-20 |
| Cd47 | 1.262750064 | 0,59938403 | 0,994 | 0,973 | 2,35E-56 |
| Arf4 | 1.244920188 | 0,5946926 | 0,823 | 0,637 | 2,31E-23 |
| Trim56 | 4.641991282 | 0,59346553 | 0,943 | 0,777 | 8,63E-29 |
| Dcp2 | 3.028215862 | 0,58569693 | 0,613 | 0,346 | 5,63E-22 |
| Xpo1 | 5.803332928 | 0,58312718 | 0,816 | 0,693 | 1,08E-12 |
| Dgkh | 9.594973627 | 0,58294698 | 0,35 | 0,109 | 1,78E-19 |
| Cish | 8.971750229 | 0,58039994 | 0,695 | 0,485 | 1,67E-15 |
| Txn1 | 2.167659122 | 0,57564077 | 0,854 | 0,755 | 4,03E-18 |
| Ogfr | 8.462340550 | 0,57225168 | 0,905 | 0,761 | 1,57E-28 |
| Gbp5 | 1.482850157 | 0,57098572 | 0,845 | 0,693 | 2,76E-16 |
| Arhgap26 | 2.734645768 | 0,56766933 | 0,791 | 0,559 | 5,08E-24 |
| Irgm2 | 3.014720985 | 0,56718199 | 0,828 | 0,548 | 5,60E-26 |
| Parp10 | 5.812879748 | 0,56354198 | 0,826 | 0,604 | 1,08E-24 |
| Sema7a | 1.894267085 | 0,56096219 | 0,262 | 0,054 | 3,52E-18 |
| Mgat1 | 4.578374180 | 0,55962882 | 0,653 | 0,435 | 8,51E-17 |
| Egr1 | 8.167550234 | 0,55693846 | 0,292 | 0,159 | 0,00015178 |
| Arsb | 3.192128171 | 0,55668944 | 0,725 | 0,53 | 5,93E-18 |
| Hk1 | 8.193737765 | 0,55229534 | 0,827 | 0,647 | 1,52E-23 |
| Tspan3 | 4.882719245 | 0,55219875 | 0,84 | 0,695 | 9,07E-17 |
| Slfn2 | 6.239913850 | 0,55191072 | 0,992 | 0,965 | 1,16E-37 |
| Gbp3 | 1.211853015 | 0,54840535 | 0,866 | 0,705 | 2,25E-20 |
| Grn | 4.563733297 | 0,54684743 | 0,414 | 0,196 | 8,48E-15 |
| Klhl6 | 2.299845072 | 0,54682426 | 0,743 | 0,546 | 4,27E-14 |
| Ehd4 | 5.035766365 | 0,54573647 | 0,353 | 0,146 | 9,36E-14 |
| Abcb1a | 7.755255389 | 0,54221256 | 0,78 | 0,594 | 1,44E-15 |
| Vps54 | 6.759871043 | 0,53777073 | 0,974 | 0,922 | 1,26E-25 |
| Atp8b4 | 6.514150948 | 0,5366018 | 0,819 | 0,621 | 1,21E-18 |
| Ascc3 | 1.656913951 | 0,53627063 | 0,859 | 0,654 | 3,08E-23 |
| Gbp6 | 2.011957643 | 0,53561619 | 0,941 | 0,728 | 3,74E-34 |
| Tdrd7 | 7.387461436 | 0,53379303 | 0,257 | 0,035 | 1,37E-21 |
| Capza2 | 3.273031920 | 0,53277251 | 0,711 | 0,489 | 6,08E-24 |
| Aida | 5.380550035 | 0,52524615 | 0,61 | 0,408 | 1,00E-12 |
| Psmb10 | 6.848363112 | 0,52268558 | 0,934 | 0,837 | 1,27E-24 |
| Rnf114 | 2.579337882 | 0,51765024 | 0,946 | 0,852 | 4,79E-27 |
| Trim14 | 2.289457121 | 0,516949 | 0,882 | 0,676 | 4,25E-23 |
| Ppa1 | 5.217286091 | 0,51598263 | 0,485 | 0,264 | 9,70E-14 |
| Psme2b | 6.607192419 | 0,51540788 | 0,924 | 0,81 | 1,23E-23 |
| Apobec1 | 6.343455868 | 0,51449283 | 0,378 | 0,134 | 1,18E-19 |
| Selenow | 4.128376287 | 0,5130689 | 0,932 | 0,829 | 7,67E-20 |
| Psmb8 | 6.316611374 | 0,51246391 | 0,989 | 0,969 | 1,17E-40 |
| Snw1 | 8.391792278 | 0,5117615 | 0,799 | 0,619 | 1,56E-18 |
| Pkm | 1.891023994 | 0,51106424 | 0,834 | 0,703 | 3,51E-15 |
| Casp8 | 1.839263705 | 0,50705421 | 0,79 | 0,594 | 3,42E-17 |
| Gbp4 | 1.480193102 | 0,50658917 | 0,976 | 0,866 | 2,75E-22 |
| Ogfrl1 | 1.232184183 | 0,50037967 | 0,298 | 0,072 | 2,29E-19 |
| Ctsc | 1.448603904 | -0,5025485 | 0,468 | 0,596 | 2,69E-07 |
| Myo1f | 1.281237446 | -0,5057976 | 0,624 | 0,736 | 2,38E-11 |
| Calm2 | 8.696303869 | -0,5080612 | 0,415 | 0,577 | 1,62E-15 |

| | | | | | |
|---|---|---|---|---|---|
| Kif13b | 4.900585226 | -0,5084853 | 0,356 | 0,546 | 9,11E-16 |
| Ptger4 | 1.704599246 | -0,5093371 | 0,441 | 0,532 | 0,00031677 |
| Aplp2 | 6.729040770 | -0,5138857 | 0,494 | 0,645 | 1,25E-13 |
| Pim2 | 8.822651632 | -0,5163539 | 0,26 | 0,408 | 1,64E-09 |
| Tspan13 | 6.014368465 | -0,5167001 | 0,246 | 0,468 | 1,12E-19 |
| St3gal6 | 1.289869516 | -0,5192607 | 0,396 | 0,569 | 2,40E-15 |
| Jmy | 1.114166428 | -0,5241748 | 0,239 | 0,412 | 2,07E-14 |
| Tmem123 | 1.541581093 | -0,5299771 | 0,72 | 0,814 | 2,86E-14 |
| Runx3 | 1.024419088 | -0,5308608 | 0,819 | 0,903 | 1,90E-22 |
| Cdc42ep3 | 1.480989721 | -0,5368212 | 0,286 | 0,485 | 2,75E-18 |
| Ddx17 | 6.406687209 | -0,5375499 | 0,816 | 0,928 | 1,19E-27 |
| Antxr2 | 5.253298112 | -0,546703 | 0,312 | 0,493 | 9,76E-16 |
| Gnas | 1.412281718 | -0,5568615 | 0,878 | 0,922 | 2,62E-17 |
| Acot7 | 2.963351906 | -0,5624544 | 0,458 | 0,584 | 5,51E-11 |
| Il27ra | 1.847990426 | -0,5703536 | 0,453 | 0,641 | 3,43E-20 |
| Thy1 | 3.942608940 | -0,5786256 | 0,963 | 0,984 | 7,33E-34 |
| Rftn1 | 3.053979564 | -0,5822826 | 0,328 | 0,54 | 5,68E-21 |
| Chd3 | 5.695385842 | -0,5931175 | 0,535 | 0,701 | 1,06E-20 |
| Ahnak | 6.914354291 | -0,5936747 | 0,847 | 0,936 | 1,28E-23 |
| Gpr132 | 5.033010331 | -0,5939764 | 0,601 | 0,73 | 9,35E-15 |
| Ddx5 | 9.416315224 | -0,5968982 | 0,882 | 0,936 | 1,75E-33 |
| Sik1 | 1.404653814 | -0,5987202 | 0,316 | 0,414 | 0,00026103 |
| Ube2h | 5.970540835 | -0,6050475 | 0,718 | 0,841 | 1,11E-20 |
| P2ry10 | 4.564221073 | -0,6052856 | 0,816 | 0,918 | 8,48E-23 |
| Lrrc8c | 1.648706089 | -0,6062193 | 0,714 | 0,843 | 3,06E-18 |
| Bcl11b | 9.447030027 | -0,6097341 | 0,756 | 0,878 | 1,76E-27 |
| Samd3 | 1.021429562 | -0,6149696 | 0,294 | 0,485 | 1,90E-16 |
| Dusp5 | 2.842655507 | -0,6176189 | 0,738 | 0,798 | 5,28E-07 |
| Smpdl3a | 1.238380195 | -0,6275098 | 0,495 | 0,666 | 2,30E-19 |
| S1pr1 | 6.845988001 | -0,6299989 | 0,297 | 0,408 | 1,27E-05 |
| Vps37b | 9.788637861 | -0,6430805 | 0,878 | 0,889 | 1,82E-10 |
| Emp3 | 4.206611899 | -0,6441156 | 0,578 | 0,734 | 7,82E-21 |
| Tagap | 4.974122729 | -0,6533541 | 0,57 | 0,718 | 9,24E-15 |
| Itgax | 5.638055842 | -0,6837765 | 0,316 | 0,476 | 1,05E-11 |
| St8sia4 | 2.190093164 | -0,6851742 | 0,375 | 0,594 | 4,07E-22 |
| Sntb1 | 4.233085520 | -0,6913267 | 0,27 | 0,536 | 7,87E-33 |
| Itga1 | 4.558086529 | -0,703211 | 0,246 | 0,377 | 8,47E-08 |
| Dnajc9 | 2.386650863 | -0,7041939 | 0,636 | 0,786 | 4,44E-25 |
| Zfp36l2 | 2.867735478 | -0,7138657 | 0,962 | 0,984 | 5,33E-29 |
| Hsph1 | 8.255129104 | -0,7654756 | 0,238 | 0,497 | 1,53E-32 |
| Dusp1 | 1.011685408 | -0,8172085 | 0,555 | 0,691 | 1,88E-07 |
| Jun | 8.032709871 | -0,8256264 | 0,553 | 0,619 | 0,00149272 |
| Klf6 | 6.529516615 | -0,8291161 | 0,868 | 0,93 | 1,21E-37 |
| Hist1h1e | 9.308318581 | -0,8488716 | 0,597 | 0,697 | 1,73E-11 |
| Ier5l | 5.987302255 | -0,8833292 | 0,105 | 0,27 | 1,11E-16 |
| Eif4a2 | 3.830492024 | -0,8905408 | 0,814 | 0,944 | 7,12E-63 |
| Tcf7 | 1.135203207 | -0,9571614 | 0,386 | 0,641 | 2,11E-31 |
| Tox | 1.249800238 | -0,9729288 | 0,112 | 0,351 | 2,32E-34 |
| Klre1 | 2.182183892 | -1,0441462 | 0,139 | 0,285 | 4,06E-12 |
| Ifi27 | 2.891351166 | -1,0889394 | 0,332 | 0,68 | 5,37E-63 |
| Cd7 | 5.543524241 | -1,1618968 | 0,303 | 0,474 | 1,03E-14 |
| Eomes | 9.243193927 | -1,2134017 | 0,261 | 0,542 | 1,72E-38 |
| Itgb1 | 9.486543875 | -1,3123409 | 0,649 | 0,885 | 1,76E-59 |

Cluster 6

| | p_val | avg_log2FC | pct.1 | pct.2 | p_val_adj |
|---|---|---|---|---|---|
| Ifit3 | 3.516244236 | 2,14689468 | 0,702 | 0,163 | 6,53E-78 |
| Isg15 | 5.252344616 | 2,05952233 | 0,808 | 0,368 | 9,76E-74 |
| Gzmb | 1.082285404 | 1,81211402 | 0,967 | 0,663 | 2,01E-54 |
| Usp18 | 2.359022419 | 1,6986503 | 0,747 | 0,255 | 4,38E-74 |
| Ifitm3 | 1.432704456 | 1,57558756 | 0,401 | 0,047 | 2,66E-37 |
| Ccl3 | 3.476309485 | 1,56409786 | 0,627 | 0,337 | 6,46E-20 |
| Ifi27l2a | 5.433736537 | 1,55339931 | 0,896 | 0,62 | 1,01E-51 |
| Ifit1bl1 | 4.509198215 | 1,4690295 | 0,78 | 0,439 | 8,38E-48 |
| Rtp4 | 2.240362251 | 1,40449391 | 0,814 | 0,333 | 4,16E-78 |
| Rnf213 | 2.957253992 | 1,38903459 | 0,954 | 0,861 | 5,50E-56 |
| Bst2 | 5.213556191 | 1,35747266 | 0,941 | 0,781 | 9,69E-62 |
| Irf7 | 2.904079971 | 1,28162483 | 0,851 | 0,55 | 5,40E-57 |
| AA467197 | 1.798626480 | 1,28106052 | 0,745 | 0,458 | 3,34E-30 |
| Isg20 | 4.651872147 | 1,24853172 | 0,854 | 0,665 | 8,64E-43 |
| Ifit1 | 6.248642567 | 1,24706317 | 0,555 | 0,085 | 1,16E-55 |
| Zbp1 | 5.279213029 | 1,20791888 | 0,964 | 0,868 | 9,81E-54 |
| Plac8 | 1.903767794 | 1,20520126 | 0,959 | 0,84 | 3,54E-41 |
| Ccl4 | 7.917527973 | 1,19710861 | 0,856 | 0,715 | 1,47E-09 |
| Oas3 | 6.521873784 | 1,17890348 | 0,798 | 0,37 | 1,21E-59 |
| Gzma | 1.154844061 | 1,17381886 | 0,793 | 0,37 | 2,15E-29 |
| Dhx58 | 8.659345554 | 1,14015354 | 0,749 | 0,325 | 1,61E-59 |
| Il2ra | 7.806021560 | 1,13005166 | 0,676 | 0,495 | 1,45E-16 |
| Slfn1 | 1.554237297 | 1,11697315 | 0,847 | 0,557 | 2,89E-48 |
| Ifitm2 | 1.476330586 | 1,09818144 | 0,298 | 0,068 | 2,74E-18 |
| Rsad2 | 1.075746942 | 1,08510977 | 0,44 | 0,047 | 2,00E-43 |
| Prf1 | 5.404385839 | 1,07913667 | 0,865 | 0,705 | 1,00E-20 |
| Xaf1 | 1.018533696 | 1,05540957 | 0,875 | 0,568 | 1,89E-58 |
| Lag3 | 3.546425231 | 1,05174804 | 0,836 | 0,601 | 6,59E-25 |
| Pik3ap1 | 3.695298982 | 1,03605335 | 0,856 | 0,63 | 6,87E-45 |
| Pdcd1 | 2.640061292 | 1,01725578 | 0,696 | 0,392 | 4,91E-26 |
| Cxcl10 | 4.919864838 | 1,00593034 | 0,354 | 0,219 | 0,00091426 |
| Ifi208 | 6.812861327 | 0,99529674 | 0,672 | 0,179 | 1,27E-65 |
| Daxx | 1.054799259 | 0,9916767 | 0,855 | 0,665 | 1,96E-36 |
| Ifih1 | 1.204299852 | 0,9714067 | 0,633 | 0,193 | 2,24E-52 |
| Lgals3bp | 1.170481687 | 0,93988458 | 0,99 | 0,92 | 2,18E-58 |
| Oas1a | 3.051072976 | 0,92753761 | 0,582 | 0,191 | 5,67E-43 |
| Ly6a | 1.164371350 | 0,92039963 | 0,992 | 0,92 | 2,16E-44 |
| Ifi209 | 5.178754784 | 0,91332508 | 0,916 | 0,811 | 9,62E-32 |
| Ifi211 | 1.039322616 | 0,90798692 | 0,553 | 0,205 | 1,93E-33 |
| Phf11b | 9.822244127 | 0,89722649 | 0,932 | 0,792 | 1,83E-39 |
| Cmpk2 | 3.171186874 | 0,88916548 | 0,523 | 0,118 | 5,89E-44 |
| Trafd1 | 3.081615744 | 0,8795537 | 0,9 | 0,759 | 5,73E-38 |
| Ifit2 | 4.345026624 | 0,85689256 | 0,704 | 0,458 | 8,07E-26 |
| Rbm3 | 6.642457199 | 0,85687912 | 0,975 | 0,906 | 1,23E-63 |
| Phf11c | 1.560275563 | 0,85151119 | 0,811 | 0,521 | 2,90E-43 |

Up in Sp140-/-

Up in Sp140+/+

| | | | | | |
|---|---|---|---|---|---|
| Samhd1 | 2.955076345 | 0,84538186 | 0,964 | 0,913 | 5,49E-26 |
| Tnfrsf18 | 1.189117280 | 0,83113768 | 0,815 | 0,59 | 2,21E-30 |
| Icos | 9.366037835 | 0,80267896 | 0,907 | 0,809 | 1,74E-17 |
| Irf8 | 4.864143376 | 0,79768934 | 0,7 | 0,446 | 9,04E-20 |
| Slfn8 | 3.116808946 | 0,78580402 | 0,822 | 0,521 | 5,79E-41 |
| Helz2 | 8.505637411 | 0,77961793 | 0,933 | 0,83 | 1,58E-31 |
| Parp9 | 1.148554518 | 0,77539377 | 0,878 | 0,667 | 2,13E-37 |
| Havcr2 | 5.292438009 | 0,7743641 | 0,632 | 0,41 | 9,83E-16 |
| Sema7a | 3.577252861 | 0,74905112 | 0,516 | 0,16 | 6,65E-32 |
| Mxd1 | 1.371773759 | 0,74318297 | 0,792 | 0,63 | 2,55E-14 |
| Tnfrsf9 | 2.524326002 | 0,74262974 | 0,621 | 0,42 | 4,69E-12 |
| Herc6 | 1.872021774 | 0,72099199 | 0,864 | 0,7 | 3,48E-24 |
| Ifi206 | 1.906290664 | 0,71888085 | 0,837 | 0,66 | 3,54E-26 |
| Nt5c3 | 1.721985883 | 0,7107802 | 0,785 | 0,616 | 3,20E-19 |
| Eif2ak2 | 1.140918125 | 0,70670427 | 0,756 | 0,436 | 2,12E-37 |
| Tnfrsf4 | 1.105449970 | 0,68335184 | 0,332 | 0,108 | 2,05E-16 |
| Myc | 7.731504690 | 0,66457559 | 0,548 | 0,366 | 1,44E-07 |
| Serpina3g | 5.627872773 | 0,66011521 | 0,883 | 0,738 | 1,05E-16 |
| Gbp9 | 2.107732895 | 0,6530358 | 0,844 | 0,62 | 3,92E-26 |
| Id2 | 2.936148796 | 0,65065423 | 0,99 | 0,955 | 5,46E-20 |
| Pml | 1.702732584 | 0,65011555 | 0,879 | 0,752 | 3,16E-27 |
| Sytl3 | 2.531895936 | 0,64835289 | 0,875 | 0,672 | 4,71E-29 |
| Mt1 | 1.693714528 | 0,64757069 | 0,506 | 0,245 | 3,15E-16 |
| Slc7a5 | 4.246558295 | 0,64557285 | 0,738 | 0,545 | 7,89E-14 |
| Phf11a | 1.336839619 | 0,63741225 | 0,621 | 0,274 | 2,48E-36 |
| Parp14 | 4.889943040 | 0,63209272 | 0,917 | 0,814 | 9,09E-19 |
| Ctss | 5.334221340 | 0,63171421 | 0,893 | 0,686 | 9,91E-28 |
| Gbp7 | 4.529294187 | 0,62701313 | 0,948 | 0,866 | 8,42E-26 |
| Ms4a4c | 1.172156014 | 0,62645337 | 0,511 | 0,255 | 2,18E-16 |
| Dtx3l | 3.374600058 | 0,62588052 | 0,941 | 0,849 | 6,27E-25 |
| Ifi47 | 1.454554712 | 0,62230917 | 0,978 | 0,92 | 2,70E-22 |
| Parp12 | 2.841324363 | 0,61980671 | 0,578 | 0,271 | 5,28E-25 |
| Ifi35 | 1.751975439 | 0,61665288 | 0,882 | 0,708 | 3,26E-29 |
| Acadl | 8.042425018 | 0,61057404 | 0,931 | 0,882 | 1,49E-18 |
| Sv2c | 1.057051404 | 0,60637331 | 0,383 | 0,099 | 1,96E-24 |
| Hif1a | 8.749129849 | 0,60432183 | 0,898 | 0,811 | 1,63E-18 |
| Irgm1 | 3.244612850 | 0,6025927 | 0,909 | 0,726 | 6,03E-26 |
| Sdc3 | 1.184062556 | 0,60150797 | 0,375 | 0,08 | 2,20E-25 |
| Trim30a | 7.437277963 | 0,59461022 | 0,79 | 0,62 | 1,38E-21 |
| Bcl2l11 | 7.128824241 | 0,59377943 | 0,717 | 0,575 | 1,32E-06 |
| Ddx58 | 3.848142450 | 0,58592402 | 0,809 | 0,634 | 7,15E-18 |
| Ifi214 | 2.083921170 | 0,58548767 | 0,647 | 0,33 | 3,87E-27 |
| Smpdl3b | 7.566621023 | 0,57252984 | 0,583 | 0,217 | 1,41E-33 |
| Trim25 | 3.515362471 | 0,56629719 | 0,836 | 0,59 | 6,53E-26 |
| Serpinb9 | 1.734581024 | 0,56233032 | 0,754 | 0,62 | 3,22E-07 |
| Naa20 | 6.281428076 | 0,56183773 | 0,865 | 0,7 | 1,17E-19 |
| Il10ra | 6.375299629 | 0,54980953 | 0,769 | 0,552 | 1,18E-18 |
| Rilpl2 | 3.809916022 | 0,54941053 | 0,781 | 0,618 | 7,08E-13 |

| | | | | | |
|---|---|---|---|---|---|
| Cd86 | 1.276517380 | 0,53646052 | 0,594 | 0,318 | 2,37E-20 |
| Hlx | 9.755969326 | 0,53582032 | 0,391 | 0,186 | 1,81E-11 |
| Lgals9 | 6.209369282 | 0,53406459 | 0,886 | 0,783 | 1,15E-14 |
| Gbp2 | 1.459695265 | 0,53188147 | 0,888 | 0,809 | 0,00027126 |
| Gstt1 | 2.415538956 | 0,53092764 | 0,415 | 0,182 | 4,49E-13 |
| H2-T24 | 2.195648928 | 0,52755817 | 0,617 | 0,333 | 4,08E-20 |
| Il18rap | 1.747830182 | 0,52705633 | 0,929 | 0,833 | 3,25E-13 |
| Trim30d | 4.864204308 | 0,52678653 | 0,546 | 0,281 | 9,04E-21 |
| Casp4 | 1.516426071 | 0,51906025 | 0,779 | 0,604 | 2,82E-15 |
| Vps54 | 2.383167891 | 0,51479227 | 0,958 | 0,915 | 4,43E-14 |
| Nfkbid | 9.690508580 | 0,51195371 | 0,629 | 0,458 | 1,80E-06 |
| Ctla4 | 2.307880994 | 0,51015532 | 0,902 | 0,828 | 4,29E-05 |
| Zc3h12d | 2.526405804 | 0,50750792 | 0,767 | 0,592 | 4,69E-13 |
| Entpd1 | 3.279396970 | 0,50742806 | 0,71 | 0,443 | 6,09E-21 |
| Mapkapk2 | 1.752417147 | 0,50396421 | 0,931 | 0,873 | 3,26E-21 |
| Tpi1 | 1.879590400 | 0,50324835 | 0,702 | 0,507 | 3,49E-15 |
| Chd3 | 7.875155357 | -0,5019183 | 0,665 | 0,792 | 1,46E-17 |
| Sesn3 | 4.594600036 | -0,5051315 | 0,717 | 0,825 | 8,54E-18 |
| Lta | 4.713266809 | -0,5279308 | 0,294 | 0,443 | 8,76E-06 |
| Samd3 | 1.204292634 | -0,546255 | 0,235 | 0,446 | 2,24E-16 |
| Dnaja1 | 4.032707067 | -0,5560006 | 0,83 | 0,903 | 7,49E-31 |
| Chst2 | 6.417611091 | -0,5581508 | 0,279 | 0,55 | 1,19E-25 |
| Eif4a2 | 2.105864450 | -0,558215 | 0,943 | 0,969 | 3,91E-29 |
| Itgax | 1.671614193 | -0,5769728 | 0,622 | 0,691 | 3,11E-07 |
| Ptger4 | 4.843481110 | -0,5814959 | 0,751 | 0,861 | 9,00E-18 |
| Ppp2r2c | 1.501150995 | -0,6135182 | 0,454 | 0,67 | 2,79E-19 |
| Hsph1 | 6.024481723 | -0,6164501 | 0,577 | 0,748 | 1,12E-24 |
| Ier5l | 1.931322268 | -0,6266477 | 0,311 | 0,436 | 0,0003589 |
| Klrg1 | 2.193284320 | -0,6482247 | 0,486 | 0,59 | 0,00040758 |
| Lmna | 7.891284628 | -0,6583538 | 0,232 | 0,507 | 1,47E-25 |
| Ldlrad4 | 6.294121628 | -0,663465 | 0,489 | 0,667 | 1,17E-15 |
| Itgb1 | 5.699226611 | -0,6679028 | 0,839 | 0,922 | 1,06E-21 |
| Tppp3 | 2.258796786 | -0,6682976 | 0,053 | 0,25 | 4,20E-25 |
| Cd74 | 8.049841883 | -0,6876986 | 0,236 | 0,436 | 1,50E-12 |
| Rhob | 2.647975826 | -0,6877958 | 0,266 | 0,429 | 4,92E-10 |
| Tcrg-C2 | 2.147211463 | -0,6890004 | 0,321 | 0,446 | 3,99E-05 |
| Cxcr6 | 3.136949830 | -0,7168766 | 0,842 | 0,884 | 5,83E-18 |
| Itga1 | 2.476149368 | -0,7206878 | 0,525 | 0,698 | 4,60E-18 |
| Tcf7 | 7.032296646 | -0,7392992 | 0,22 | 0,441 | 1,31E-16 |
| Cdh1 | 8.862768293 | -0,7406078 | 0,232 | 0,5 | 1,65E-25 |
| Fos | 1.161132239 | -0,7557578 | 0,568 | 0,642 | 0,00215773 |
| Tcrg-C4 | 3.304361835 | -0,7586953 | 0,106 | 0,325 | 6,14E-22 |
| Klf6 | 2.297833620 | -0,7808082 | 0,957 | 0,976 | 4,27E-22 |
| Myadm | 1.902754436 | -0,7921685 | 0,549 | 0,752 | 3,54E-25 |
| Jun | 1.050058996 | -1,1547655 | 0,647 | 0,797 | 1,95E-20 |
| Itgae | 1.670433802 | -1,1915891 | 0,297 | 0,568 | 3,10E-24 |
| Hspa1b | 5.142200810 | -1,3811693 | 0,036 | 0,262 | 9,56E-37 |

Cluster 7

| | p_val | avg_log2FC | pct.1 | pct.2 | p_val_adj |
|---|---|---|---|---|---|
| Gzmb | 2.258576541 | 2,16722301 | 0,736 | 0,12 | 4,20E-51 |
| Ifit3 | 5.858357944 | 1,44383438 | 0,512 | 0,169 | 1,09E-15 |
| Ccl5 | 2.867028387 | 1,33761053 | 0,791 | 0,624 | 0,0053278 |
| Ly6a | 4.50184153€ | 1,0534269 | 0,938 | 0,645 | 8,37E-16 |
| Isg15 | 1.00355761€ | 0,99360522 | 0,395 | 0,094 | 1,86E-14 |
| Gzma | 1.05528817C | 0,98197307 | 0,566 | 0,156 | 1,96E-20 |
| Ccl4 | 5.929695087 | 0,94523063 | 0,527 | 0,176 | 1,10E-12 |
| Xaf1 | 1.863325681 | 0,92144732 | 0,729 | 0,422 | 3,46E-10 |
| Ccl3 | 4.533057232 | 0,85102252 | 0,256 | 0,046 | 8,42E-11 |
| Ifi27l2a | 5.11867712€ | 0,77301459 | 0,628 | 0,268 | 9,51E-12 |
| Rtp4 | 3.538425367 | 0,77260791 | 0,62 | 0,32 | 6,58E-08 |
| Trim34a | 1.772650817 | 0,7657276 | 0,729 | 0,438 | 3,29E-07 |
| Ifi208 | 3.236762317 | 0,75248176 | 0,581 | 0,287 | 6,01E-08 |
| Slfn8 | 1.969381122 | 0,75080297 | 0,643 | 0,445 | 0,0036597 |
| Lgals3bp | 9.352926124 | 0,7361202 | 0,837 | 0,724 | 1,74E-06 |
| Rnf213 | 4.958419484 | 0,71414052 | 0,938 | 0,847 | 0,00921423 |
| Ccnd3 | 5.98042458€ | 0,71397258 | 0,876 | 0,626 | 1,11E-08 |
| Abca1 | 4.409708784 | 0,70720342 | 0,442 | 0,166 | 8,19E-08 |
| Ifit1 | 2.603428327 | 0,70432153 | 0,287 | 0,054 | 4,84E-12 |
| Fkbp5 | 1.006837012 | 0,69260077 | 0,457 | 0,184 | 1,87E-07 |
| Coro2a | 1.42766603C | 0,68621775 | 0,349 | 0,159 | 0,00265303 |
| Nkg7 | 7.69241907C | 0,68045492 | 0,891 | 0,755 | 0,00142948 |
| Ggt1 | 6.409962654 | 0,6759597 | 0,496 | 0,268 | 0,00119116 |
| Cysltr2 | 7.79946412€ | 0,65461671 | 0,403 | 0,192 | 0,00144937 |
| Cdkn2d | 4.14916487C | 0,64526175 | 0,527 | 0,325 | 0,00771039 |
| Oasl2 | 2.323033484 | 0,60678844 | 0,341 | 0,151 | 0,00431689 |
| Dusp7 | 3.08407651€ | 0,56450255 | 0,473 | 0,233 | 0,00057311 |
| Dhx58 | 2.94857937C | 0,53385644 | 0,349 | 0,158 | 0,00547935 |
| Sell | 1.633504304 | 0,52979097 | 0,977 | 0,961 | 3,04E-06 |
| Fermt3 | 3.324469331 | 0,52732072 | 0,736 | 0,599 | 0,00617786 |
| Apobec3 | 8.610146462 | 0,52703274 | 0,93 | 0,841 | 0,00016 |
| Gramd1a | 6.85607601§ | -0,5554522 | 0,791 | 0,903 | 0,00127406 |
| Il6st | 5.02743932£ | -0,5746205 | 0,822 | 0,895 | 0,00093425 |
| Actn1 | 2.20792942: | -0,5836724 | 0,62 | 0,839 | 0,0004103 |
| Chd3 | 2.27890206£ | -0,7015535 | 0,729 | 0,836 | 4,23E-05 |
| Vps37b | 1.85384594€ | -0,7416216 | 0,86 | 0,954 | 3,45E-05 |
| Ifi27 | 7.80222655: | -0,7742628 | 0,388 | 0,665 | 1,45E-05 |
| Sik1 | 2.92003766€ | -0,8357177 | 0,488 | 0,662 | 0,00542631 |
| Cd28 | 5.25260823: | -1,1467381 | 0,434 | 0,75 | 9,76E-10 |
| Hsph1 | 5.98523240£ | -1,1600504 | 0,194 | 0,506 | 1,11E-07 |

Up in Sp140-/-

Up in Sp140+/+

Cluster 8

| | p_val | avg_log2FC | pct.1 | pct.2 | p_val_adj |
|---|---|---|---|---|---|
| Gzmb | 5.321686267 | 2,3401997 | 0,978 | 0,62 | 9,89E-26 |
| Ifitm1 | 1.718111977 | 2,29083144 | 0,254 | 0,009 | 3,19E-19 |
| Ifit3 | 1.024301618 | 2,23533918 | 0,687 | 0,12 | 1,90E-39 |
| Ifi27l2a | 2.156361506 | 2,15180718 | 0,776 | 0,402 | 4,01E-18 |
| Gzma | 1.526996586 | 2,04345924 | 0,873 | 0,439 | 2,84E-17 |
| Isg15 | 1.060634013 | 1,84621266 | 0,694 | 0,239 | 1,97E-23 |
| Usp18 | 5.058205156 | 1,63078549 | 0,575 | 0,134 | 9,40E-26 |
| Rnf213 | 8.857304392 | 1,55085737 | 0,955 | 0,805 | 1,65E-19 |
| Plac8 | 9.292685406 | 1,44072025 | 0,97 | 0,786 | 1,73E-18 |
| AA467197 | 3.698794533 | 1,42693517 | 0,59 | 0,264 | 6,87E-11 |
| Rtp4 | 1.792437391 | 1,37654696 | 0,769 | 0,309 | 3,33E-24 |
| Bst2 | 1.171159524 | 1,36679401 | 0,851 | 0,67 | 2,18E-10 |
| Pdcd1 | 1.768699139 | 1,36362711 | 0,612 | 0,282 | 3,29E-10 |
| Slfn1 | 6.276200080 | 1,33350595 | 0,694 | 0,268 | 1,17E-20 |
| Ly6a | 4.983454167 | 1,32310254 | 0,948 | 0,791 | 9,26E-13 |
| Oas3 | 2.058353668 | 1,22024904 | 0,642 | 0,257 | 3,83E-18 |
| Il2ra | 2.359624261 | 1,21989514 | 0,44 | 0,166 | 4,38E-09 |
| Ifitm3 | 2.699348319 | 1,2094315 | 0,261 | 0,007 | 5,02E-21 |
| Fkbp5 | 2.820511131 | 1,20933654 | 0,672 | 0,309 | 5,24E-17 |
| Ifit1 | 1.382406411 | 1,2039796 | 0,545 | 0,077 | 2,57E-31 |
| Irf7 | 1.820632845 | 1,18571305 | 0,799 | 0,475 | 3,38E-15 |
| Mt1 | 2.706171126 | 1,16173623 | 0,321 | 0,057 | 5,03E-13 |
| Zbp1 | 1.008932815 | 1,15494673 | 0,888 | 0,755 | 1,87E-13 |
| Ifi208 | 3.776170001 | 1,14357662 | 0,59 | 0,141 | 7,02E-25 |
| Ifi209 | 1.092476349 | 1,13345787 | 0,866 | 0,691 | 2,03E-13 |
| Phf11b | 1.778651631 | 1,08872695 | 0,881 | 0,595 | 3,31E-16 |
| Ifit1bl1 | 1.603589318 | 1,07601361 | 0,672 | 0,457 | 2,98E-06 |
| Lgals3bp | 4.685442772 | 1,06274311 | 0,955 | 0,88 | 8,71E-16 |
| Rsad2 | 1.809700362 | 1,02453033 | 0,403 | 0,023 | 3,36E-30 |
| Oas1a | 1.170309976 | 1,01591725 | 0,567 | 0,227 | 2,17E-12 |
| Dhx58 | 2.454505335 | 0,99233475 | 0,582 | 0,23 | 4,56E-14 |
| Sytl3 | 3.738890330 | 0,98852855 | 0,813 | 0,518 | 6,95E-12 |
| Klrb1c | 4.055961997 | 0,97289782 | 0,291 | 0,03 | 7,54E-16 |
| Xaf1 | 7.358833466 | 0,96596629 | 0,776 | 0,461 | 1,37E-13 |
| Mxd1 | 3.390820814 | 0,92498931 | 0,776 | 0,625 | 0,00630116 |
| Parp9 | 1.956600737 | 0,92407716 | 0,821 | 0,493 | 3,64E-15 |
| Tnfrsf9 | 3.633493906 | 0,91425852 | 0,47 | 0,252 | 0,00675212 |
| Lag3 | 2.266414167 | 0,89084674 | 0,806 | 0,609 | 0,00042117 |
| Irf8 | 1.937326777 | 0,87925868 | 0,493 | 0,259 | 3,60E-05 |
| Klra7 | 2.771836389 | 0,87300424 | 0,403 | 0,168 | 0,00051509 |
| Ddit4 | 2.255723510 | 0,86756551 | 0,761 | 0,486 | 4,19E-07 |
| Cmpk2 | 9.179469649 | 0,85625622 | 0,388 | 0,086 | 1,71E-13 |
| Trafd1 | 1.914160717 | 0,84657372 | 0,836 | 0,589 | 3,56E-10 |
| Rbm3 | 7.239573453 | 0,83929999 | 0,888 | 0,757 | 1,35E-13 |
| Bcl2l11 | 1.944733305 | 0,83624007 | 0,679 | 0,466 | 0,00036139 |
| Entpd1 | 8.084443968 | 0,80201492 | 0,53 | 0,214 | 1,50E-10 |
| Nfkbia | 1.979345685 | 0,80187665 | 0,799 | 0,67 | 0,00367822 |
| Trim30a | 2.646675913 | 0,78384367 | 0,709 | 0,425 | 4,92E-10 |
| Serpinb9 | 7.153719914 | 0,77472229 | 0,769 | 0,543 | 0,00132938 |
| Samhd1 | 2.864874865 | 0,75602795 | 0,97 | 0,923 | 5,32E-08 |

Up in Sp140-/-
Up in Sp140+/+

| | | | | | |
|---|---|---|---|---|---|
| Slfn5 | 6.59359656 | 0,75433096 | 0,343 | 0,061 | 1,23E-13 |
| Eif2ak2 | 1.73452504 | 0,74057047 | 0,56 | 0,241 | 3,22E-10 |
| Ifi206 | 2.84368402 | 0,73961837 | 0,813 | 0,598 | 5,28E-07 |
| Pml | 5.12981737 | 0,72225935 | 0,694 | 0,477 | 9,53E-06 |
| Ifih1 | 4.44454511 | 0,71072596 | 0,381 | 0,1 | 8,26E-11 |
| Slfn8 | 1.988503701 | 0,71035162 | 0,769 | 0,493 | 3,70E-09 |
| Ifi35 | 8.20352786 | 0,70292666 | 0,784 | 0,555 | 1,52E-07 |
| Helz2 | 2.227348052 | 0,68275075 | 0,843 | 0,698 | 0,00413908 |
| Cd86 | 3.15430222 | 0,68238785 | 0,44 | 0,155 | 5,86E-09 |
| Rab27a | 1.06759042 | 0,68039713 | 0,851 | 0,732 | 1,98E-06 |
| Plek | 8.084412294 | 0,67894656 | 0,821 | 0,666 | 0,00150233 |
| Pim1 | 1.908568964 | 0,67579178 | 0,799 | 0,63 | 0,00035467 |
| Herc6 | 4.29558835 | 0,67494608 | 0,724 | 0,455 | 7,98E-06 |
| Acp5 | 2.456441844 | 0,6706429 | 0,851 | 0,655 | 4,56E-06 |
| Ifit2 | 1.11863304 | 0,65846349 | 0,418 | 0,211 | 0,00207876 |
| Ms4a4c | 1.87925788 | 0,64767153 | 0,269 | 0,057 | 3,49E-08 |
| Ifi211 | 2.750621371 | 0,63969398 | 0,276 | 0,03 | 5,11E-15 |
| Hif1a | 4.57507982 | 0,62987878 | 0,627 | 0,425 | 0,00085019 |
| Parp14 | 7.407244584 | 0,62838433 | 0,836 | 0,668 | 1,38E-05 |
| Dtx3l | 9.97964723 | 0,62139731 | 0,888 | 0,793 | 1,85E-05 |
| Phf11c | 9.23321287 | 0,62050411 | 0,664 | 0,35 | 1,72E-08 |
| Nt5c3 | 1.008337651 | 0,61768824 | 0,694 | 0,461 | 0,00187379 |
| Id2 | 1.60964246 | 0,6106804 | 0,985 | 0,9 | 0,0029912 |
| Ifit3b | 4.58067260 | 0,60602593 | 0,261 | 0,018 | 8,51E-17 |
| Casp3 | 5.066558682 | 0,60469507 | 0,799 | 0,57 | 9,42E-06 |
| Apobec3 | 1.44633907 | 0,60282977 | 0,985 | 0,907 | 2,69E-10 |
| Ctla4 | 4.859269302 | 0,59934193 | 0,828 | 0,677 | 0,00902998 |
| Pkm | 2.69480090 | 0,59817544 | 0,873 | 0,77 | 5,01E-05 |
| Phf11a | 1.10351342 | 0,59598795 | 0,396 | 0,143 | 2,05E-07 |
| B4galt5 | 2.417775784 | 0,58786458 | 0,858 | 0,732 | 0,00449295 |
| Ehd4 | 1.08077240 | 0,58381775 | 0,448 | 0,214 | 0,00020084 |
| Cd47 | 3.43461372 | 0,57903307 | 0,97 | 0,955 | 6,38E-10 |
| Xdh | 2.06280093 | 0,57808503 | 0,575 | 0,334 | 0,00038333 |
| Gbp9 | 1.48443555 | 0,57105143 | 0,813 | 0,582 | 0,00027585 |
| Arsb | 1.51497843 | 0,56968795 | 0,866 | 0,727 | 2,82E-05 |
| Trim30d | 4.76154436 | 0,56606298 | 0,425 | 0,2 | 8,85E-05 |
| Serpinb6b | 8.768821287 | 0,55578647 | 0,433 | 0,211 | 0,00162951 |
| Ifi47 | 2.145118621 | 0,54915806 | 0,933 | 0,836 | 0,00398627 |
| Ms4a6b | 6.19030368 | 0,53689921 | 0,978 | 0,914 | 0,00115034 |
| Mndal | 1.15988927 | 0,53546084 | 0,896 | 0,843 | 2,16E-05 |
| Gpr65 | 8.754964932 | 0,53364222 | 0,746 | 0,514 | 0,00162694 |
| Epsti1 | 2.68832071 | 0,52762808 | 0,985 | 0,914 | 5,00E-05 |
| Gsto1 | 1.421616531 | 0,52648497 | 0,313 | 0,084 | 2,64E-07 |
| Ccrl2 | 6.40217504 | 0,525529 | 0,276 | 0,064 | 1,19E-07 |
| Tmem181a | 1.25199284 | -0,5068461 | 0,194 | 0,43 | 0,00232658 |
| Jmy | 5.57897380 | -0,5388306 | 0,269 | 0,505 | 0,00103674 |
| Saraf | 3.63338275 | -0,5615597 | 0,948 | 0,993 | 6,75E-12 |
| Sntb1 | 1.243559162 | -0,5704763 | 0,336 | 0,591 | 2,31E-05 |
| Ypel3 | 6.975391471 | -0,5748298 | 0,761 | 0,925 | 1,30E-06 |
| Tgfbr2 | 2.809940227 | -0,5850412 | 0,903 | 0,977 | 5,22E-09 |
| Cd101 | 2.62054840 | -0,5885661 | 0,313 | 0,566 | 0,00048698 |
| Rabac1 | 3.464867411 | -0,5917445 | 0,784 | 0,898 | 6,44E-05 |
| Camk2n1 | 2.96638487 | -0,6214552 | 0,209 | 0,434 | 0,00551243 |

| | | | | | |
|---|---|---|---|---|---|
| Tcrg-C2 | 8.912204518 | -0,6256314 | 0,813 | 0,868 | 0,00016562 |
| Ptger4 | 4.480550179 | -0,6423715 | 0,634 | 0,798 | 0,00832621 |
| St3gal6 | 3.977607639 | -0,644547 | 0,403 | 0,657 | 7,39E-06 |
| Bcl11b | 1.513976762 | -0,6496715 | 0,731 | 0,866 | 2,81E-06 |
| St8sia4 | 6.466190241 | -0,656377 | 0,552 | 0,773 | 1,20E-06 |
| Slc20a1 | 2.373830031 | -0,6613626 | 0,552 | 0,743 | 0,00044113 |
| Hsph1 | 1.217241757 | -0,6695608 | 0,261 | 0,545 | 2,26E-05 |
| Ppp2r2c | 1.754089130 | -0,6911946 | 0,254 | 0,475 | 0,00325962 |
| Ifi27 | 2.250372068 | -0,7191237 | 0,545 | 0,814 | 4,18E-08 |
| Tcf7 | 4.646280040 | -0,7237576 | 0,343 | 0,561 | 0,00863418 |
| Trat1 | 1.278329763 | -0,7328476 | 0,269 | 0,584 | 2,38E-06 |
| Cdh1 | 2.996779818 | -0,7511304 | 0,246 | 0,511 | 0,00055689 |
| Samd3 | 2.795377095 | -0,7607931 | 0,299 | 0,559 | 0,00051946 |
| Klf6 | 6.422632212 | -0,7873645 | 0,918 | 0,964 | 1,19E-08 |
| Chd3 | 9.012370360 | -0,8352169 | 0,463 | 0,707 | 1,67E-08 |
| Eif4a2 | 1.822245838 | -0,8470298 | 0,888 | 0,961 | 3,39E-19 |
| Trdc | 2.566262268 | -0,8965893 | 0,448 | 0,634 | 0,00476889 |
| Itgb1 | 1.346017115 | -0,9689126 | 0,784 | 0,882 | 2,50E-05 |
| Itgax | 1.201398919 | -1,0007113 | 0,582 | 0,775 | 2,23E-06 |
| Eomes | 1.295210981 | -1,2419911 | 0,201 | 0,505 | 2,41E-07 |

| Cluster 9 | | | | | | | |
|---|---|---|---|---|---|---|---|
| | p_val | avg_log2FC | pct.1 | pct.2 | p_val_adj | | |
| Gzmb | 1.694883602 | 3,36730217 | 0,663 | 0,167 | 3,15E-13 | | Up in Sp140-/- |
| Gzma | 7.773510476 | 2,77109711 | 0,462 | 0,111 | 1,44E-05 | | Up in Sp140+/+ |
| B4galt7 | 2.537710425 | -0,502734 | 0,055 | 0,278 | 0,00471583 | | |
| Diaph2 | 1.841550929 | -0,5029843 | 0,075 | 0,322 | 0,00342215 | | |
| Sept11 | 2.269941451 | -0,5060844 | 0,131 | 0,422 | 0,00421823 | | |
| Rptor | 1.454937854 | -0,507736 | 0,035 | 0,256 | 0,00027037 | | |
| Wdr77 | 3.684595063 | -0,5082866 | 0,045 | 0,278 | 0,00068471 | | |
| Naa50 | 3.620428294 | -0,509295 | 0,131 | 0,422 | 0,00672784 | | |
| Atxn7l3b | 1.035582428 | -0,5104536 | 0,141 | 0,444 | 0,00192442 | | |
| Aplp2 | 1.848563941 | -0,5111812 | 0,07 | 0,333 | 0,00034352 | | |
| Ubtf | 1.930748494 | -0,5114075 | 0,141 | 0,444 | 0,00358791 | | |
| Atp6v1h | 4.201180449 | -0,5117186 | 0,126 | 0,444 | 0,00078071 | | |
| Idnk | 3.374669395 | -0,5125987 | 0,136 | 0,433 | 0,00627115 | | |
| Lrrc59 | 2.794117059 | -0,5131479 | 0,106 | 0,4 | 0,00051923 | | |
| Tmem173 | 1.118110999 | -0,5139662 | 0,101 | 0,367 | 0,00207779 | | |
| Drg1 | 6.536966842 | -0,5181293 | 0,106 | 0,389 | 0,00121476 | | |
| Rab3gap1 | 6.335269528 | -0,5199182 | 0,075 | 0,333 | 0,00117728 | | |
| Cyth1 | 1.922539111 | -0,5208407 | 0,161 | 0,478 | 0,00357265 | | |
| Slc25a28 | 1.235965854 | -0,5210583 | 0,06 | 0,322 | 0,00022968 | | |
| Bms1 | 1.868698379 | -0,5222329 | 0,04 | 0,267 | 0,00034726 | | |
| Susd6 | 1.671206836 | -0,5225014 | 0,166 | 0,5 | 0,00031056 | | |
| Hnrnpl | 4.226145328 | -0,5230467 | 0,126 | 0,433 | 0,00078534 | | |
| Uqcrc2 | 3.158414617 | -0,5231618 | 0,151 | 0,444 | 0,00586928 | | |
| Numa1 | 6.006674297 | -0,523801 | 0,141 | 0,456 | 0,00111622 | | |
| Ccnl1 | 4.069849330 | -0,5250698 | 0,166 | 0,467 | 0,007563 | | |
| Myo1e | 2.432239310 | -0,5265493 | 0,055 | 0,278 | 0,00451983 | | |
| Chordc1 | 4.915538109 | -0,5267367 | 0,06 | 0,3 | 0,00091345 | | |
| Map3k2 | 1.486031186 | -0,5269472 | 0,116 | 0,456 | 2,76E-05 | | |
| Trmt112 | 3.025208276 | -0,5271624 | 0,161 | 0,478 | 0,00562174 | | |
| Tomm34 | 3.880911270 | -0,5271641 | 0,106 | 0,422 | 7,21E-05 | | |
| Naip2 | 9.909209031 | -0,5271838 | 0,065 | 0,3 | 0,00184143 | | |
| Gpr107 | 2.236660076 | -0,5278342 | 0,09 | 0,344 | 0,00415639 | | |
| Psmc3 | 1.436544801 | -0,5294591 | 0,166 | 0,489 | 0,00266953 | | |
| Gusb | 1.448571149 | -0,5309716 | 0,085 | 0,344 | 0,00269188 | | |
| Glud1 | 4.333456050 | -0,5310779 | 0,216 | 0,533 | 0,00805286 | | |
| Anapc1 | 4.283027702 | -0,5312711 | 0,07 | 0,356 | 7,96E-05 | | |
| Galnt1 | 2.372432684 | -0,5321435 | 0,156 | 0,489 | 0,00044087 | | |
| Clk4 | 2.156291498 | -0,5326954 | 0,111 | 0,378 | 0,00400704 | | |
| Hnrnpk | 3.859100124 | -0,5344437 | 0,221 | 0,556 | 0,00717137 | | |
| Hprt | 2.194925717 | -0,5348388 | 0,166 | 0,467 | 0,00407883 | | |
| Sra1 | 3.260062714 | -0,5356969 | 0,141 | 0,444 | 0,00605817 | | |
| Ogt | 2.957183473 | -0,5365276 | 0,146 | 0,433 | 0,00549533 | | |
| Gmps | 4.789032630 | -0,5406716 | 0,075 | 0,311 | 0,00889946 | | |
| Pdia6 | 3.342840821 | -0,5410639 | 0,121 | 0,411 | 0,0006212 | | |
| Ndufs8 | 1.942199834 | -0,5411421 | 0,151 | 0,456 | 0,00360919 | | |
| Gbp6 | 4.474008927 | -0,5415203 | 0,136 | 0,422 | 0,00831405 | | |
| Siah2 | 7.301788122 | -0,5422585 | 0,05 | 0,278 | 0,00135689 | | |
| Tspan14 | 2.253556036 | -0,5435611 | 0,095 | 0,378 | 0,00041878 | | |
| Dgkd | 1.348969185 | -0,5455708 | 0,095 | 0,422 | 2,51E-05 | | |
| Ero1lb | 1.795934020 | -0,5456217 | 0,06 | 0,289 | 0,00333738 | | |

| | | | | | |
|---|---|---|---|---|---|
| Ap1ar | 4.713107577 | -0,5458474 | 0,06 | 0,278 | 0,00875837 |
| Dusp7 | 4.552840377 | -0,5464186 | 0,055 | 0,267 | 0,00846054 |
| Napsa | 1.349647998 | -0,5474737 | 0,06 | 0,322 | 0,00025081 |
| Purb | 1.876115423 | -0,5490107 | 0,151 | 0,467 | 0,00348639 |
| Prpf40a | 1.270090222 | -0,5495972 | 0,136 | 0,467 | 0,00023602 |
| Dnajc8 | 4.611266756 | -0,5504254 | 0,131 | 0,411 | 0,00856912 |
| Dnajc13 | 1.305002441 | -0,550572 | 0,095 | 0,367 | 0,00242509 |
| Hnrnpa2b1 | 1.609843690 | -0,5518193 | 0,236 | 0,6 | 0,00299157 |
| Stx16 | 6.455194473 | -0,5530153 | 0,06 | 0,367 | 1,20E-06 |
| Crbn | 2.927606484 | -0,5537949 | 0,111 | 0,378 | 0,00544037 |
| Lta4h | 3.835961185 | -0,5539135 | 0,085 | 0,356 | 0,00071284 |
| Trim26 | 1.524307998 | -0,5540236 | 0,085 | 0,344 | 0,00283262 |
| Scly | 5.105600825 | -0,5551559 | 0,075 | 0,311 | 0,00948774 |
| Set | 3.301759132 | -0,5555094 | 0,101 | 0,4 | 0,00061357 |
| Supt16 | 4.537457851 | -0,5557211 | 0,106 | 0,4 | 0,0008432 |
| Rnf13 | 2.038314842 | -0,5562406 | 0,101 | 0,411 | 3,79E-05 |
| Yrdc | 2.808403009 | -0,5567052 | 0,05 | 0,267 | 0,00521886 |
| Ddx21 | 9.723602017 | -0,5581113 | 0,166 | 0,5 | 0,00180694 |
| Nub1 | 2.570202365 | -0,5583942 | 0,131 | 0,444 | 0,00047762 |
| Cd2bp2 | 1.209724990 | -0,5595989 | 0,121 | 0,433 | 0,0002248 |
| Usp12 | 4.091727446 | -0,561834 | 0,075 | 0,367 | 7,60E-05 |
| Abcc5 | 1.157589881 | -0,562025 | 0,045 | 0,256 | 0,00215115 |
| Plekhj1 | 4.044103024 | -0,5627176 | 0,141 | 0,444 | 0,00751516 |
| Ywhaz | 1.138190097 | -0,5631607 | 0,07 | 0,311 | 0,0021151 |
| Rgs2 | 3.458669710 | -0,5631726 | 0,101 | 0,356 | 0,00642725 |
| Slc38a2 | 1.762489359 | -0,5642117 | 0,111 | 0,378 | 0,00327523 |
| Prkar1a | 2.152897138 | -0,5643307 | 0,196 | 0,533 | 0,00400073 |
| Mtdh | 2.216981818 | -0,5666374 | 0,126 | 0,467 | 4,12E-05 |
| Smarca2 | 1.875799703 | -0,5689839 | 0,055 | 0,3 | 0,00034858 |
| Romo1 | 2.516681306 | -0,5699971 | 0,07 | 0,333 | 0,00046767 |
| Pmpca | 1.662832755 | -0,5704274 | 0,04 | 0,311 | 3,09E-06 |
| Macf1 | 1.597288606 | -0,5708743 | 0,161 | 0,511 | 0,00029682 |
| Ncf4 | 3.434976724 | -0,5735378 | 0,176 | 0,489 | 0,00638322 |
| Yif1a | 1.475878254 | -0,5744247 | 0,06 | 0,289 | 0,00274262 |
| Esrra | 2.052077696 | -0,5755256 | 0,055 | 0,3 | 0,00038134 |
| Lrrc8c | 1.144740991 | -0,575732 | 0,116 | 0,4 | 0,00212727 |
| Trappc3 | 4.062246871 | -0,5774417 | 0,085 | 0,333 | 0,00754887 |
| Tpp2 | 6.933578864 | -0,5783027 | 0,146 | 0,489 | 0,00012885 |
| Bccip | 2.044262873 | -0,5797786 | 0,116 | 0,4 | 0,00379885 |
| Sipa1l3 | 9.384880802 | -0,5801233 | 0,025 | 0,278 | 1,74E-06 |
| Cept1 | 1.190321820 | -0,5835889 | 0,131 | 0,422 | 0,00221198 |
| Dazap2 | 3.410949319 | -0,5839111 | 0,241 | 0,6 | 0,00633857 |
| Nek7 | 3.490842852 | -0,5843006 | 0,116 | 0,444 | 6,49E-05 |
| Dnajc9 | 5.978290858 | -0,5855058 | 0,095 | 0,4 | 0,00011109 |
| Cbfb | 1.024286971 | -0,5857025 | 0,196 | 0,533 | 0,00190343 |
| Prkcb | 5.808978557 | -0,5860608 | 0,106 | 0,378 | 0,00107948 |
| Fli1 | 1.435978356 | -0,5866986 | 0,146 | 0,489 | 0,00026685 |
| Cog5 | 2.556068398 | -0,5870955 | 0,075 | 0,367 | 4,75E-05 |
| Maz | 1.651119764 | -0,5875108 | 0,171 | 0,511 | 0,00030683 |
| Abhd17a | 1.848563941 | -0,5879332 | 0,07 | 0,333 | 0,00034352 |
| Slc12a7 | 4.769158952 | -0,5884787 | 0,06 | 0,322 | 8,86E-05 |
| Bcor | 1.941169308 | -0,5888451 | 0,045 | 0,322 | 3,61E-06 |
| Rbbp4 | 2.306091369 | -0,5889066 | 0,176 | 0,5 | 0,00428541 |

| | | | | | |
|---|---|---|---|---|---|
| Ino80b | 2.307130018 | -0,5906451 | 0,06 | 0,289 | 0,00428734 |
| Bin2 | 4.282295863 | -0,5913629 | 0,216 | 0,544 | 0,00795779 |
| Trpc4ap | 1.534937169 | -0,591937 | 0,156 | 0,533 | 2,85E-06 |
| Cbl | 1.294792077 | -0,5922142 | 0,146 | 0,456 | 0,00240611 |
| Tmem222 | 3.279866699 | -0,5961706 | 0,045 | 0,267 | 0,0006095 |
| Prrc2a | 2.239603657 | -0,5963557 | 0,106 | 0,378 | 0,00416186 |
| Trap1 | 4.113640005 | -0,5972953 | 0,055 | 0,3 | 0,00076444 |
| Scyl1 | 2.693262795 | -0,5978621 | 0,095 | 0,356 | 0,00500489 |
| Rock1 | 1.061152012 | -0,5984864 | 0,136 | 0,5 | 1,97E-05 |
| Grb2 | 8.497142015 | -0,5987292 | 0,196 | 0,522 | 0,00157902 |
| Sh3kbp1 | 2.023606751 | -0,5989577 | 0,121 | 0,433 | 0,00037605 |
| Csnk1a1 | 6.450806808 | -0,5992689 | 0,126 | 0,433 | 0,00119875 |
| Mpv17l2 | 2.500226370 | -0,5994456 | 0,045 | 0,278 | 0,00046462 |
| Psen2 | 1.482041831 | -0,602113 | 0,05 | 0,289 | 0,00027541 |
| Dock8 | 1.376510556 | -0,6021693 | 0,111 | 0,389 | 0,00255797 |
| Dpysl2 | 4.744876741 | -0,6047783 | 0,151 | 0,433 | 0,0088174 |
| Tram1 | 2.128016947 | -0,6048959 | 0,206 | 0,544 | 0,00395449 |
| Rap1a | 2.788370086 | -0,6062876 | 0,211 | 0,578 | 0,00051816 |
| Tasor | 1.310892758 | -0,6084197 | 0,045 | 0,278 | 0,0002436 |
| Zmat5 | 1.655341567 | -0,6086459 | 0,055 | 0,278 | 0,00307612 |
| Snrpb | 9.042444107 | -0,6106541 | 0,111 | 0,422 | 0,00016804 |
| Csnk1g2 | 4.681066837 | -0,610828 | 0,136 | 0,444 | 0,00086988 |
| Pdpk1 | 7.759596683 | -0,611035 | 0,101 | 0,4 | 0,0001442 |
| Nme1 | 1.269855679 | -0,6150849 | 0,045 | 0,278 | 0,00023598 |
| Tmem234 | 5.040809868 | -0,6151205 | 0,151 | 0,444 | 0,00936734 |
| Tmed9 | 5.022903838 | -0,6155797 | 0,186 | 0,5 | 0,00933406 |
| Dgkz | 5.813720247 | -0,6163947 | 0,161 | 0,489 | 0,00108036 |
| Arid1b | 1.735856576 | -0,6171999 | 0,116 | 0,456 | 3,23E-05 |
| Ep400 | 1.049570974 | -0,6172848 | 0,121 | 0,411 | 0,00195042 |
| Tomm7 | 2.191371565 | -0,617721 | 0,126 | 0,433 | 0,00040722 |
| Ap3d1 | 2.030569452 | -0,6179427 | 0,111 | 0,4 | 0,00037734 |
| Ddx39b | 1.529011151 | -0,6181504 | 0,09 | 0,4 | 2,84E-05 |
| Eif4g3 | 1.117828207 | -0,6181656 | 0,131 | 0,478 | 2,08E-05 |
| B4galt1 | 1.991869945 | -0,6187816 | 0,136 | 0,456 | 0,00037015 |
| Cwc15 | 4.924389391 | -0,620211 | 0,146 | 0,444 | 0,00915099 |
| Nek9 | 1.641291813 | -0,6213675 | 0,106 | 0,378 | 0,00305001 |
| Arglu1 | 1.918315495 | -0,6219907 | 0,141 | 0,433 | 0,00356481 |
| Srsf5 | 1.668599621 | -0,6231491 | 0,161 | 0,467 | 0,00310076 |
| Acox3 | 7.516121002 | -0,6258909 | 0,075 | 0,367 | 1,40E-05 |
| Scpep1 | 1.925174321 | -0,6259431 | 0,09 | 0,378 | 0,00035776 |
| Map2k2 | 4.293129311 | -0,6282969 | 0,156 | 0,522 | 7,98E-05 |
| Mtss1 | 8.094047202 | -0,6308074 | 0,03 | 0,256 | 0,00015041 |
| Rasa1 | 1.582069533 | -0,6335099 | 0,08 | 0,333 | 0,00293996 |
| Prkx | 4.817098997 | -0,6369803 | 0,065 | 0,311 | 0,00089516 |
| Ctsh | 5.678664639 | -0,6372321 | 0,201 | 0,622 | 1,06E-06 |
| Agpat4 | 1.745573158 | -0,6374552 | 0,065 | 0,3 | 0,0032438 |
| Papola | 6.196637542 | -0,6378004 | 0,211 | 0,567 | 0,00115152 |
| Atp2c1 | 3.300580303 | -0,6383578 | 0,075 | 0,333 | 0,00061335 |
| Tnrc18 | 2.202948412 | -0,6385712 | 0,09 | 0,389 | 4,09E-05 |
| St8sia4 | 3.746081188 | -0,6387515 | 0,095 | 0,4 | 6,96E-05 |
| Stag2 | 9.874993925 | -0,6395773 | 0,111 | 0,444 | 1,84E-05 |
| Itsn2 | 1.597299291 | -0,6400879 | 0,166 | 0,5 | 0,00029683 |
| Irs2 | 1.027875392 | -0,6410025 | 0,055 | 0,289 | 0,0019101 |

| | | | | | |
|---|---|---|---|---|---|
| Rpia | 4.223835073 | -0,6418104 | 0,08 | 0,322 | 0,00784915 |
| Lpcat1 | 2.864280185 | -0,6426727 | 0,045 | 0,3 | 5,32E-05 |
| Rsrp1 | 1.639164512 | -0,6439277 | 0,196 | 0,533 | 0,00304606 |
| Lypla1 | 4.037691277 | -0,6441303 | 0,085 | 0,378 | 7,50E-05 |
| Tomm70a | 1.869863625 | -0,6455942 | 0,08 | 0,378 | 3,47E-05 |
| Elk3 | 5.608750027 | -0,6469365 | 0,121 | 0,411 | 0,00104227 |
| Myo1c | 5.448456686 | -0,6484743 | 0,121 | 0,422 | 0,00101249 |
| Mfsd1 | 1.687798285 | -0,6491807 | 0,095 | 0,367 | 0,00031364 |
| Cic | 2.714785883 | -0,6513787 | 0,131 | 0,467 | 5,04E-05 |
| Stk10 | 1.730420016 | -0,6513956 | 0,166 | 0,478 | 0,00321564 |
| Pxk | 4.870063586 | -0,6524007 | 0,126 | 0,4 | 0,00905004 |
| Bola2 | 5.411289602 | -0,6525015 | 0,121 | 0,411 | 0,00100558 |
| Grcc10 | 1.544259805 | -0,6528279 | 0,211 | 0,556 | 0,0028697 |
| Suds3 | 1.781470914 | -0,6533327 | 0,116 | 0,411 | 0,00033105 |
| Sgpp1 | 5.318812018 | -0,6534804 | 0,05 | 0,322 | 9,88E-06 |
| Mbp | 9.455651543 | -0,6541736 | 0,095 | 0,422 | 1,76E-05 |
| Calm2 | 1.482768599 | -0,6549155 | 0,131 | 0,444 | 0,00027554 |
| Mapk14 | 1.707605216 | -0,6551774 | 0,136 | 0,456 | 0,00031732 |
| Arpc1b | 1.652582746 | -0,655695 | 0,291 | 0,667 | 0,00307099 |
| Ncln | 2.280739712 | -0,6571061 | 0,085 | 0,378 | 4,24E-05 |
| Gdi2 | 1.058005605 | -0,6571926 | 0,266 | 0,633 | 0,00196609 |
| Cln3 | 3.756357913 | -0,6578724 | 0,045 | 0,289 | 6,98E-05 |
| Arid1a | 1.175058302 | -0,6588628 | 0,151 | 0,456 | 0,00218361 |
| Commd4 | 3.123726921 | -0,6617002 | 0,07 | 0,4 | 5,80E-07 |
| Ecpas | 1.394445866 | -0,6646191 | 0,08 | 0,356 | 0,00025913 |
| Mgrn1 | 1.658525486 | -0,6656762 | 0,065 | 0,356 | 3,08E-05 |
| Eef1d | 6.817044886 | -0,6660158 | 0,085 | 0,356 | 0,00126681 |
| Nxf1 | 5.038931909 | -0,6661945 | 0,161 | 0,467 | 0,00936385 |
| Smg7 | 1.783588551 | -0,6663298 | 0,111 | 0,378 | 0,00331444 |
| Dnajb11 | 7.917563890 | -0,6676758 | 0,111 | 0,389 | 0,00147132 |
| Vrk1 | 8.797877401 | -0,6676951 | 0,045 | 0,289 | 0,00016349 |
| Dnajc3 | 7.583205482 | -0,6678979 | 0,136 | 0,467 | 0,00014092 |
| Necap2 | 6.810011340 | -0,6682561 | 0,116 | 0,411 | 0,0012655 |
| Tspan13 | 1.590833794 | -0,668336 | 0,121 | 0,4 | 0,00295625 |
| Ech1 | 1.847308759 | -0,6687407 | 0,131 | 0,489 | 3,43E-05 |
| Ucp2 | 1.082938309 | -0,6688126 | 0,492 | 0,878 | 2,01E-05 |
| Micos13 | 5.665050784 | -0,6688555 | 0,08 | 0,389 | 1,05E-05 |
| Naca | 1.078665767 | -0,669568 | 0,231 | 0,578 | 0,00200448 |
| Matr3 | 2.786198230 | -0,6702311 | 0,09 | 0,367 | 0,00051776 |
| Ndufv2 | 2.683353062 | -0,6730377 | 0,151 | 0,456 | 0,00498647 |
| Abi1 | 4.166367991 | -0,6780292 | 0,186 | 0,556 | 7,74E-06 |
| Ccdc86 | 1.136344124 | -0,68074 | 0,055 | 0,311 | 0,00021117 |
| Pdlim1 | 2.284860820 | -0,6812202 | 0,075 | 0,311 | 0,00424596 |
| Atg7 | 1.337908408 | -0,6812359 | 0,035 | 0,256 | 0,00024862 |
| Camk1d | 1.332309044 | -0,6823901 | 0,085 | 0,389 | 2,48E-05 |
| Tmem50b | 4.733084238 | -0,6840161 | 0,035 | 0,289 | 8,80E-06 |
| Ndufb4 | 6.545572544 | -0,6841812 | 0,05 | 0,3 | 0,00012164 |
| Pten | 7.827440078 | -0,6849648 | 0,196 | 0,533 | 0,00145457 |
| Dym | 2.643611498 | -0,685509 | 0,05 | 0,333 | 4,91E-06 |
| Rab1b | 3.055038265 | -0,6870589 | 0,146 | 0,522 | 5,68E-06 |
| H2afz | 3.763451521 | -0,6892475 | 0,171 | 0,544 | 6,99E-05 |
| Sec24b | 1.624948123 | -0,6892694 | 0,116 | 0,389 | 0,00301964 |
| Gpx1 | 1.157851527 | -0,6910576 | 0,291 | 0,622 | 0,00215164 |

| | | | | | |
|---|---|---|---|---|---|
| Dus3l | 9.395999584 | -0,6938488 | 0,05 | 0,344 | 1,75E-06 |
| Pnpla8 | 4.985411714 | -0,6949957 | 0,055 | 0,333 | 9,26E-06 |
| Svil | 1.924310530 | -0,6965015 | 0,05 | 0,267 | 0,00357595 |
| Chmp5 | 1.366374880 | -0,6985529 | 0,146 | 0,489 | 2,54E-05 |
| Scarb2 | 2.487942332 | -0,6992432 | 0,101 | 0,344 | 0,00462334 |
| Acsl5 | 1.396615937 | -0,69959 | 0,131 | 0,422 | 0,00259533 |
| Ola1 | 8.445974410 | -0,6996327 | 0,106 | 0,411 | 0,00015695 |
| Saraf | 7.030628515 | -0,7029326 | 0,176 | 0,511 | 0,0013065 |
| Slc25a11 | 9.121140171 | -0,7034367 | 0,07 | 0,344 | 0,0001695 |
| Syngr2 | 6.323518116 | -0,7039966 | 0,136 | 0,467 | 0,00011751 |
| Nploc4 | 1.628052679 | -0,7048533 | 0,07 | 0,333 | 0,00030254 |
| Hmgcr | 2.030037453 | -0,7071559 | 0,06 | 0,311 | 0,00037724 |
| Ccng1 | 3.752431788 | -0,7077269 | 0,156 | 0,456 | 0,00697314 |
| Dync1h1 | 2.584028620 | -0,7093055 | 0,156 | 0,511 | 4,80E-05 |
| Stx8 | 8.949537730 | -0,7093948 | 0,09 | 0,356 | 0,00166309 |
| Sept6 | 3.148384531 | -0,7142965 | 0,136 | 0,456 | 0,00058506 |
| Tm6sf1 | 3.387403621 | -0,717096 | 0,126 | 0,456 | 6,29E-05 |
| Ing1 | 1.701995818 | -0,7186547 | 0,075 | 0,322 | 0,00316282 |
| Nagpa | 8.086222223 | -0,7194856 | 0,04 | 0,278 | 0,00015027 |
| Pdia4 | 2.875248005 | -0,7210085 | 0,106 | 0,389 | 0,00053431 |
| Ahsa1 | 9.923509890 | -0,7220038 | 0,146 | 0,456 | 0,00184409 |
| Hsp90ab1 | 1.350657060 | -0,72242 | 0,201 | 0,556 | 0,00025099 |
| Cat | 1.207252938 | -0,7227476 | 0,09 | 0,4 | 2,24E-05 |
| Tap2 | 6.916779102 | -0,7270799 | 0,251 | 0,656 | 0,00128535 |
| Arhgap17 | 8.725720624 | -0,7305632 | 0,08 | 0,356 | 0,00016215 |
| Itgb7 | 2.194125260 | -0,7306349 | 0,176 | 0,489 | 0,00407734 |
| Uqcr11 | 9.745985716 | -0,7306404 | 0,085 | 0,378 | 0,00018111 |
| Thada | 2.495665292 | -0,7349151 | 0,04 | 0,289 | 4,64E-05 |
| Atp5e | 6.539974946 | -0,7369875 | 0,176 | 0,522 | 0,00012153 |
| Safb2 | 1.705602657 | -0,7385452 | 0,121 | 0,4 | 0,00316952 |
| Slc8a1 | 4.231951701 | -0,7410379 | 0,045 | 0,278 | 0,00078642 |
| Ehbp1l1 | 5.736153079 | -0,7411863 | 0,101 | 0,4 | 0,00010659 |
| Rpe | 1.309979530 | -0,7417289 | 0,121 | 0,411 | 0,00243433 |
| Lcp1 | 5.227104197 | -0,7427235 | 0,467 | 0,767 | 0,00971353 |
| Klf4 | 1.245343827 | -0,7428167 | 0,045 | 0,289 | 0,00023142 |
| Tagln2 | 3.695052467 | -0,7430132 | 0,307 | 0,678 | 0,00068665 |
| Chd3 | 5.621759091 | -0,7437255 | 0,065 | 0,333 | 0,00010447 |
| S100a13 | 1.399265959 | -0,7469967 | 0,126 | 0,444 | 0,00026003 |
| Zdhhc5 | 2.774143480 | -0,7481401 | 0,116 | 0,378 | 0,00515519 |
| Arhgdia | 2.793146564 | -0,7489188 | 0,271 | 0,633 | 0,0051905 |
| Cdkn1b | 1.399213433 | -0,7489611 | 0,171 | 0,522 | 0,00026002 |
| Lamtor4 | 1.021598898 | -0,7492327 | 0,06 | 0,289 | 0,00189844 |
| Tns3 | 1.066135849 | -0,7504132 | 0,04 | 0,278 | 0,00019812 |
| Ppp1r10 | 1.887608053 | -0,7509087 | 0,095 | 0,356 | 0,00350774 |
| Vdac2 | 8.825354536 | -0,7511123 | 0,161 | 0,5 | 0,000164 |
| Cyb561a3 | 3.631813547 | -0,7532033 | 0,085 | 0,333 | 0,006749 |
| Cdk19 | 5.328959764 | -0,7542186 | 0,06 | 0,278 | 0,00990281 |
| Chd4 | 5.389822458 | -0,7562278 | 0,116 | 0,478 | 1,00E-06 |
| Gsk3b | 3.383686357 | -0,7563745 | 0,08 | 0,467 | 6,29E-09 |
| Capn2 | 7.251835251 | -0,7572165 | 0,095 | 0,367 | 0,00134761 |
| Ndufa4 | 1.698161140 | -0,7598285 | 0,09 | 0,344 | 0,00315569 |
| Plekhb2 | 3.002351861 | -0,7610832 | 0,151 | 0,5 | 5,58E-05 |
| Supt20 | 1.313627855 | -0,7611026 | 0,085 | 0,389 | 2,44E-05 |

| | | | | | |
|---|---|---|---|---|---|
| Dock2 | 1.100746174 | -0,7650433 | 0,236 | 0,611 | 0,00204552 |
| Anapc5 | 1.310892758 | -0,7667121 | 0,045 | 0,278 | 0,0002436 |
| Ccdc50 | 2.111881534 | -0,767855 | 0,095 | 0,378 | 0,00039245 |
| Agfg1 | 3.538135915 | -0,769169 | 0,191 | 0,544 | 0,00065749 |
| Scarb1 | 2.080563623 | -0,7692285 | 0,055 | 0,278 | 0,00386631 |
| Aars | 1.164300899 | -0,7699832 | 0,065 | 0,333 | 0,00021636 |
| Ctso | 6.479341983 | -0,7717948 | 0,05 | 0,278 | 0,00120406 |
| Preb | 9.792461682 | -0,7725618 | 0,131 | 0,422 | 0,00181973 |
| Pcf11 | 1.426159411 | -0,7727108 | 0,111 | 0,4 | 0,00026502 |
| Ikbkb | 2.165101727 | -0,7733299 | 0,126 | 0,433 | 0,00040234 |
| Canx | 7.325724046 | -0,7739381 | 0,106 | 0,478 | 1,36E-07 |
| Ybx1 | 2.193663051 | -0,7755805 | 0,191 | 0,533 | 0,00040765 |
| Dad1 | 6.217952885 | -0,7782033 | 0,186 | 0,522 | 0,00115548 |
| Ubfd1 | 1.677867610 | -0,7787879 | 0,035 | 0,3 | 3,12E-06 |
| Tgfbr2 | 4.680028885 | -0,7792883 | 0,206 | 0,522 | 0,0086969 |
| Cops4 | 1.692460272 | -0,7793591 | 0,08 | 0,378 | 3,15E-05 |
| Atp2a3 | 1.820385738 | -0,7795007 | 0,106 | 0,4 | 0,00033828 |
| Adam17 | 1.480447792 | -0,7803583 | 0,136 | 0,456 | 0,00027511 |
| Klhl9 | 3.598037984 | -0,7826186 | 0,08 | 0,367 | 6,69E-05 |
| Dynlrb1 | 1.019990570 | -0,7826482 | 0,206 | 0,556 | 0,00189545 |
| Arl6ip5 | 2.054990871 | -0,7838563 | 0,161 | 0,5 | 0,00038188 |
| Camk2d | 1.126902417 | -0,7850985 | 0,095 | 0,4 | 2,09E-05 |
| Mknk2 | 1.630004008 | -0,7864064 | 0,191 | 0,556 | 0,0003029 |
| Mtch1 | 4.700149931 | -0,787016 | 0,101 | 0,378 | 0,00087343 |
| Ifnar2 | 1.299792459 | -0,7898755 | 0,141 | 0,467 | 0,00024154 |
| Khsrp | 1.042319516 | -0,7914528 | 0,131 | 0,478 | 1,94E-05 |
| Foxp1 | 1.415798581 | -0,793103 | 0,106 | 0,456 | 2,63E-06 |
| Gosr2 | 3.824694721 | -0,7946016 | 0,151 | 0,478 | 0,00071074 |
| Vsir | 5.674286539 | -0,7947096 | 0,176 | 0,511 | 0,00105445 |
| Mink1 | 5.805886178 | -0,7952398 | 0,111 | 0,4 | 0,00107891 |
| Sh3bgrl | 3.199898647 | -0,7962741 | 0,166 | 0,489 | 0,00059464 |
| Skp1a | 2.566931481 | -0,7966519 | 0,111 | 0,411 | 0,00047701 |
| Dcaf8 | 9.273901252 | -0,7979976 | 0,075 | 0,378 | 1,72E-05 |
| Rnf167 | 1.705602657 | -0,798067 | 0,121 | 0,4 | 0,00316952 |
| Cdc37 | 2.334130504 | -0,7993286 | 0,136 | 0,467 | 4,34E-05 |
| Son | 8.353554670 | -0,8010212 | 0,251 | 0,611 | 0,00155234 |
| Sdha | 7.922602167 | -0,8011387 | 0,141 | 0,533 | 1,47E-06 |
| Psmb7 | 1.180703006 | -0,8014783 | 0,111 | 0,411 | 0,00021941 |
| Srf | 3.046480240 | -0,801691 | 0,126 | 0,411 | 0,00566127 |
| Ivns1abp | 2.270353019 | -0,8028438 | 0,156 | 0,533 | 4,22E-06 |
| Ak2 | 3.545319342 | -0,8036742 | 0,126 | 0,422 | 0,00065883 |
| Fgfr1op2 | 4.429890222 | -0,8049753 | 0,191 | 0,522 | 0,00082321 |
| Eif3f | 9.158952031 | -0,8060896 | 0,256 | 0,622 | 0,00170201 |
| Smap2 | 2.846655834 | -0,8066912 | 0,206 | 0,567 | 0,00052899 |
| Cers2 | 5.807731432 | -0,8067454 | 0,095 | 0,389 | 0,00010793 |
| Lamtor2 | 7.174390317 | -0,8079922 | 0,131 | 0,456 | 0,00013332 |
| Rexo2 | 2.778382978 | -0,8092073 | 0,126 | 0,4 | 0,00516307 |
| Sgpl1 | 1.181729231 | -0,8093308 | 0,101 | 0,356 | 0,00219601 |
| Lims1 | 2.562636938 | -0,818951 | 0,116 | 0,389 | 0,00476215 |
| BC005537 | 8.706331466 | -0,8198468 | 0,251 | 0,633 | 0,00016179 |
| Sbno2 | 6.072646128 | -0,8222606 | 0,176 | 0,5 | 0,00112848 |
| Sypl | 1.202802347 | -0,8252615 | 0,09 | 0,422 | 2,24E-06 |
| Tbl1xr1 | 4.447721497 | -0,8256413 | 0,101 | 0,444 | 8,27E-07 |

| | | | | | |
|---|---|---|---|---|---|
| Nfkb1 | 8.994705404 | -0,8268986 | 0,176 | 0,533 | 0,00016715 |
| Hvcn1 | 1.389619940 | -0,8283967 | 0,095 | 0,389 | 0,00025823 |
| Manf | 9.351059841 | -0,8330061 | 0,126 | 0,444 | 0,00017377 |
| Emsy | 6.020133926 | -0,8332935 | 0,04 | 0,278 | 0,00011187 |
| Prex1 | 2.797814607 | -0,8334411 | 0,241 | 0,6 | 0,00519918 |
| Rock2 | 4.578876942 | -0,8342235 | 0,065 | 0,356 | 8,51E-06 |
| Eif3l | 4.319938675 | -0,8350099 | 0,156 | 0,522 | 8,03E-06 |
| Golga4 | 9.121690358 | -0,8367301 | 0,085 | 0,344 | 0,00169508 |
| Evl | 4.137908132 | -0,8367375 | 0,196 | 0,556 | 0,00076895 |
| Psmb1 | 2.935689789 | -0,838368 | 0,146 | 0,556 | 5,46E-08 |
| Pik3ca | 7.643550501 | -0,8396196 | 0,075 | 0,333 | 0,0014204 |
| Ddx17 | 2.370879548 | -0,8410257 | 0,161 | 0,533 | 4,41E-06 |
| Snrnp200 | 4.554046605 | -0,8410598 | 0,126 | 0,444 | 8,46E-05 |
| Ilk | 5.408467465 | -0,8415727 | 0,146 | 0,456 | 0,00100506 |
| Tle5 | 3.647520523 | -0,843937 | 0,161 | 0,522 | 6,78E-05 |
| Fam102b | 5.250992439 | -0,8445849 | 0,05 | 0,278 | 0,00097579 |
| Cd300a | 1.434003799 | -0,8446871 | 0,05 | 0,267 | 0,00266481 |
| Rack1 | 2.110800447 | -0,8447373 | 0,317 | 0,678 | 0,0039225 |
| Clta | 2.207186011 | -0,8448467 | 0,382 | 0,822 | 4,10E-05 |
| Npepl1 | 4.713788209 | -0,8453363 | 0,131 | 0,411 | 0,00875963 |
| Eif4a2 | 4.756682105 | -0,8458532 | 0,246 | 0,633 | 8,84E-05 |
| Rp9 | 5.268285551 | -0,8468705 | 0,111 | 0,411 | 9,79E-05 |
| Tpr | 2.868047891 | -0,8508983 | 0,171 | 0,533 | 5,33E-05 |
| Rasgef1b | 4.885949820 | -0,8518238 | 0,065 | 0,289 | 0,00907956 |
| Slc25a5 | 5.535512487 | -0,8525312 | 0,216 | 0,556 | 0,00102866 |
| Atf6b | 3.723074554 | -0,8535125 | 0,06 | 0,367 | 6,92E-07 |
| Wdr82 | 1.419697884 | -0,8586101 | 0,121 | 0,433 | 0,00026382 |
| Skil | 8.446931485 | -0,8602116 | 0,171 | 0,5 | 0,00156969 |
| Ids | 3.473000276 | -0,8615455 | 0,07 | 0,367 | 6,45E-06 |
| Ndufa13 | 1.612421435 | -0,8618372 | 0,196 | 0,544 | 0,00029964 |
| Aagab | 1.253137699 | -0,8626186 | 0,095 | 0,356 | 0,00232871 |
| Tmem59 | 8.561409284 | -0,8639745 | 0,131 | 0,5 | 1,59E-06 |
| Itm2b | 9.318256283 | -0,8669695 | 0,452 | 0,8 | 0,00173161 |
| Rabac1 | 4.457479289 | -0,871045 | 0,181 | 0,511 | 0,00082833 |
| Rassf4 | 5.458732889 | -0,8724226 | 0,065 | 0,333 | 0,00010144 |
| Hspa5 | 1.609104288 | -0,874556 | 0,136 | 0,456 | 0,00029902 |
| Tfeb | 2.032688853 | -0,8787961 | 0,07 | 0,378 | 3,78E-06 |
| Plcg2 | 6.625958192 | -0,8796754 | 0,08 | 0,378 | 1,23E-05 |
| Cops9 | 5.200653050 | -0,880039 | 0,206 | 0,544 | 0,00096644 |
| Rab14 | 6.949420949 | -0,8814855 | 0,231 | 0,611 | 0,00012914 |
| Nipbl | 4.212148419 | -0,8821297 | 0,101 | 0,489 | 7,83E-09 |
| Ube2g2 | 5.195501645 | -0,8824135 | 0,121 | 0,422 | 0,00096548 |
| Lmbrd1 | 2.779448402 | -0,8849569 | 0,116 | 0,411 | 0,0005165 |
| Gabarap | 5.842689548 | -0,8852445 | 0,221 | 0,567 | 0,00108575 |
| Anxa5 | 1.557575737 | -0,8867458 | 0,191 | 0,511 | 0,00289444 |
| Prpsap2 | 7.699477089 | -0,8869403 | 0,04 | 0,278 | 0,00014308 |
| Hnrnpu | 8.190615116 | -0,8873403 | 0,166 | 0,622 | 1,52E-08 |
| Psmb9 | 1.277413723 | -0,8901654 | 0,332 | 0,7 | 0,00237382 |
| Usf1 | 1.526948985 | -0,8909671 | 0,095 | 0,4 | 2,84E-05 |
| Man2b2 | 3.127829914 | -0,8932677 | 0,075 | 0,356 | 5,81E-05 |
| Laptm5 | 7.457978858 | -0,8939916 | 0,467 | 0,778 | 0,00138592 |
| Akap13 | 1.100559484 | -0,8953518 | 0,211 | 0,544 | 0,00204517 |
| Srsf6 | 5.972003529 | -0,8960868 | 0,151 | 0,489 | 0,00011098 |

| | | | | | |
|---|---|---|---|---|---|
| Prpf8 | 2.563629412 | -0,9001054 | 0,166 | 0,522 | 4,76E-05 |
| Myo1f | 4.257094521 | -0,9034429 | 0,161 | 0,444 | 0,00791096 |
| Ptprc | 8.652501994 | -0,9040777 | 0,412 | 0,756 | 0,00160789 |
| Nabp1 | 7.920197771 | -0,9042402 | 0,176 | 0,578 | 1,47E-05 |
| Naxd | 7.601532405 | -0,9046661 | 0,035 | 0,267 | 0,00014126 |
| Stt3b | 2.492764557 | -0,9098425 | 0,121 | 0,456 | 4,63E-06 |
| Elmo1 | 2.042996608 | -0,9132604 | 0,186 | 0,567 | 3,80E-05 |
| Eloa | 4.857041909 | -0,9134401 | 0,116 | 0,478 | 9,03E-07 |
| Naaa | 1.913982470 | -0,9140557 | 0,065 | 0,344 | 3,56E-05 |
| Pdia3 | 4.861979377 | -0,9143244 | 0,171 | 0,489 | 0,0009035 |
| Adamts10 | 2.305935233 | -0,9202318 | 0,075 | 0,333 | 0,00042851 |
| Ncoa3 | 1.211790826 | -0,9205054 | 0,121 | 0,522 | 2,25E-08 |
| Tapbp | 1.165447307 | -0,9226248 | 0,317 | 0,689 | 0,00216575 |
| Ssb | 1.039803657 | -0,9245832 | 0,171 | 0,511 | 0,00019323 |
| Mat2a | 1.837862578 | -0,9266826 | 0,111 | 0,422 | 3,42E-05 |
| Man2b1 | 6.998229946 | -0,9277498 | 0,161 | 0,456 | 0,00130048 |
| Prrc2c | 1.362499250 | -0,928211 | 0,171 | 0,544 | 2,53E-05 |
| Rab43 | 1.418105020 | -0,9296603 | 0,095 | 0,411 | 2,64E-05 |
| Rnpepl1 | 4.848206440 | -0,9310416 | 0,08 | 0,356 | 9,01E-05 |
| Akr1b10 | 2.348836853 | -0,9317598 | 0,045 | 0,278 | 0,00043648 |
| H2-DMb1 | 2.916531646 | -0,9367916 | 0,075 | 0,333 | 0,00054198 |
| Swap70 | 3.458002944 | -0,9374819 | 0,055 | 0,278 | 0,00642601 |
| Psmc2 | 1.934198252 | -0,9383194 | 0,08 | 0,356 | 0,00035943 |
| Ppp3r1 | 1.368417216 | -0,9418717 | 0,106 | 0,478 | 2,54E-07 |
| Pdlim5 | 7.333863881 | -0,9452363 | 0,08 | 0,367 | 0,00013629 |
| Nop10 | 1.716738380 | -0,946194 | 0,095 | 0,411 | 3,19E-05 |
| Ip6k1 | 2.493124970 | -0,9467853 | 0,101 | 0,411 | 4,63E-05 |
| H2-D1 | 2.953215945 | -0,9475764 | 0,216 | 0,567 | 0,0005488 |
| Lamp1 | 2.409659614 | -0,9498256 | 0,181 | 0,511 | 0,00044779 |
| Ddx5 | 8.171643328 | -0,9498473 | 0,226 | 0,578 | 0,00015185 |
| Ldb1 | 7.042580098 | -0,9500104 | 0,176 | 0,522 | 0,00013087 |
| Uvrag | 4.569470971 | -0,9502133 | 0,085 | 0,4 | 8,49E-06 |
| Srrm2 | 2.685752766 | -0,9502466 | 0,156 | 0,522 | 4,99E-06 |
| Rtn3 | 7.146438641 | -0,955085 | 0,136 | 0,489 | 1,33E-06 |
| Elovl5 | 8.759897437 | -0,9552944 | 0,065 | 0,389 | 1,63E-07 |
| H2-K1 | 1.063750871 | -0,9576305 | 0,472 | 0,878 | 1,98E-08 |
| Serp1 | 8.562430426 | -0,9602255 | 0,065 | 0,356 | 1,59E-05 |
| Zmiz2 | 2.851996761 | -0,9632924 | 0,09 | 0,444 | 5,30E-08 |
| Hsp90b1 | 3.656869380 | -0,9635583 | 0,101 | 0,467 | 6,80E-08 |
| Lamp2 | 3.194883352 | -0,9669233 | 0,211 | 0,522 | 0,00059371 |
| Serf2 | 3.584192379 | -0,967307 | 0,266 | 0,644 | 0,00066605 |
| Iigp1 | 2.431664855 | -0,9690861 | 0,121 | 0,389 | 0,00451876 |
| Rdx | 2.791256188 | -0,9722253 | 0,085 | 0,356 | 0,0005187 |
| Smarcc2 | 5.625144323 | -0,973642 | 0,121 | 0,456 | 1,05E-05 |
| Atp5j | 8.929240592 | -0,9737966 | 0,131 | 0,422 | 0,00165932 |
| Rhoa | 7.433752453 | -0,9779859 | 0,246 | 0,611 | 0,00013814 |
| Colgalt1 | 6.575780662 | -0,9787296 | 0,106 | 0,378 | 0,00122198 |
| Serbp1 | 2.649865398 | -0,9852898 | 0,176 | 0,511 | 0,00049242 |
| Nfya | 1.420697518 | -0,986466 | 0,065 | 0,3 | 0,00264008 |
| Jmjd1c | 9.869536110 | -0,9878477 | 0,106 | 0,422 | 1,83E-05 |
| Lnpep | 7.612934778 | -0,9886111 | 0,166 | 0,556 | 1,41E-06 |
| Atp2b1 | 5.825060664 | -0,9895219 | 0,085 | 0,389 | 1,08E-05 |
| Arhgef6 | 5.895929173 | -0,9904691 | 0,131 | 0,456 | 0,00010956 |

| | | | | | |
|---|---|---|---|---|---|
| Ppp2r1a | 1.617447705 | -0,9906119 | 0,166 | 0,556 | 3,01E-06 |
| Rftn1 | 1.231670409 | -0,9907968 | 0,116 | 0,444 | 2,29E-05 |
| Snx2 | 1.750065506 | -0,9910807 | 0,131 | 0,489 | 3,25E-06 |
| Far1 | 1.125368501 | -0,9920469 | 0,116 | 0,467 | 2,09E-06 |
| Iqgap1 | 2.104185153 | -0,9925618 | 0,266 | 0,611 | 0,00391021 |
| Hdac7 | 2.064007932 | -0,9943558 | 0,08 | 0,322 | 0,00383555 |
| Ndufa3 | 3.384485989 | -0,9953597 | 0,186 | 0,511 | 0,00062894 |
| Bag1 | 3.382996815 | -0,996389 | 0,166 | 0,489 | 0,00062866 |
| Hmgcs1 | 1.628922975 | -0,9978172 | 0,06 | 0,311 | 0,0003027 |
| Osbpl8 | 1.745205257 | -1,0000624 | 0,111 | 0,456 | 3,24E-06 |
| Ddx3x | 9.316062984 | -1,005171 | 0,171 | 0,556 | 1,73E-06 |
| Plaat3 | 1.320528324 | -1,0058217 | 0,276 | 0,644 | 0,00245394 |
| Sh3bp5 | 7.353796383 | -1,0104462 | 0,04 | 0,256 | 0,00136656 |
| Eef1a1 | 2.746714570 | -1,0117211 | 0,749 | 0,978 | 5,10E-07 |
| Cotl1 | 4.199116021 | -1,0125692 | 0,412 | 0,8 | 0,00078032 |
| Nedd8 | 4.718814973 | -1,0136806 | 0,211 | 0,567 | 0,0008769 |
| Parp1 | 4.926854320 | -1,0140652 | 0,06 | 0,389 | 9,16E-08 |
| Ncf1 | 2.783377077 | -1,0179549 | 0,186 | 0,511 | 0,00051723 |
| Akr1a1 | 7.366281618 | -1,0191597 | 0,302 | 0,722 | 1,37E-05 |
| Ciita | 1.619089537 | -1,0209077 | 0,05 | 0,311 | 3,01E-05 |
| Dhx15 | 1.864384146 | -1,0214959 | 0,111 | 0,478 | 3,46E-07 |
| Csde1 | 6.459796148 | -1,0259154 | 0,136 | 0,544 | 1,20E-07 |
| Rnf130 | 6.580173224 | -1,0260602 | 0,121 | 0,478 | 1,22E-06 |
| Was | 5.134649691 | -1,026575 | 0,136 | 0,533 | 9,54E-08 |
| Nckap1l | 1.395207650 | -1,0295688 | 0,171 | 0,567 | 2,59E-07 |
| Lmo4 | 1.007444554 | -1,0370779 | 0,111 | 0,411 | 0,00018721 |
| Ifi27 | 2.038625600 | -1,047259 | 0,09 | 0,478 | 3,79E-09 |
| Gnas | 1.808742516 | -1,0612723 | 0,201 | 0,644 | 3,36E-08 |
| Gm2a | 2.512085101 | -1,0678888 | 0,221 | 0,578 | 0,00046682 |
| Tomm6 | 6.372338965 | -1,0731701 | 0,101 | 0,444 | 1,18E-06 |
| Lat2 | 3.925321216 | -1,0789199 | 0,035 | 0,333 | 7,29E-08 |
| Emg1 | 3.183867150 | -1,0828559 | 0,111 | 0,467 | 5,92E-07 |
| Ptges3 | 3.580718072 | -1,0877286 | 0,121 | 0,456 | 6,65E-06 |
| Stx7 | 2.676639754 | -1,0905822 | 0,171 | 0,533 | 4,97E-06 |
| Atp6ap2 | 1.728610343 | -1,0942829 | 0,146 | 0,444 | 0,00032123 |
| Ly86 | 3.747106664 | -1,0968691 | 0,07 | 0,367 | 6,96E-06 |
| Tcirg1 | 3.668028049 | -1,1072398 | 0,116 | 0,422 | 6,82E-05 |
| Cst3 | 1.118841895 | -1,1099119 | 0,281 | 0,7 | 2,08E-06 |
| Puf60 | 2.694832934 | -1,1104039 | 0,136 | 0,544 | 5,01E-08 |
| Ctsb | 2.182198372 | -1,1158947 | 0,302 | 0,667 | 0,00040552 |
| Tmsb4x | 2.133123913 | -1,116032 | 0,874 | 1 | 3,96E-09 |
| Mbd3 | 2.144040276 | -1,1172766 | 0,116 | 0,433 | 3,98E-05 |
| Ptpn6 | 1.050257871 | -1,1356915 | 0,261 | 0,644 | 1,95E-05 |
| Aldh2 | 2.866145432 | -1,1401121 | 0,07 | 0,411 | 5,33E-08 |
| Cd83 | 6.492163848 | -1,1465606 | 0,045 | 0,267 | 0,00120644 |
| Tgfbi | 3.195470225 | -1,1479827 | 0,196 | 0,511 | 0,00059381 |
| Lgmn | 6.091172506 | -1,1698418 | 0,085 | 0,344 | 0,00113192 |
| H2-DMa | 5.982741824 | -1,1770525 | 0,08 | 0,5 | 1,11E-10 |
| Eif3c | 2.342283141 | -1,1804432 | 0,095 | 0,5 | 4,35E-10 |
| Hexa | 2.143945959 | -1,1811773 | 0,116 | 0,4 | 0,00039841 |
| Irf8 | 8.328013605 | -1,1856065 | 0,166 | 0,522 | 1,55E-05 |
| Cd44 | 2.370180569 | -1,2101735 | 0,231 | 0,6 | 4,40E-05 |
| Tmsb10 | 2.580080888 | -1,2160119 | 0,432 | 0,833 | 4,79E-06 |

| | | | | | |
|---|---|---|---|---|---|
| Tppp3 | 2.389814272 | -1,2163571 | 0,045 | 0,256 | 0,00444099 |
| Prdx1 | 4.809686032 | -1,2164348 | 0,221 | 0,644 | 8,94E-07 |
| Crip1 | 1.191531658 | -1,2328965 | 0,236 | 0,722 | 2,21E-09 |
| Ndufs7 | 4.128465675 | -1,2350573 | 0,106 | 0,5 | 7,67E-09 |
| Gns | 1.194627941 | -1,275538 | 0,156 | 0,444 | 0,00221998 |
| Vps37b | 9.581774799 | -1,2783929 | 0,131 | 0,444 | 0,00017806 |
| Erp29 | 3.515559146 | -1,2856978 | 0,166 | 0,544 | 6,53E-07 |
| Ctsz | 1.196505743 | -1,3175806 | 0,251 | 0,622 | 2,22E-05 |
| Pecam1 | 5.111007668 | -1,3348435 | 0,055 | 0,289 | 0,00094978 |
| Hspa8 | 6.996436001 | -1,3545337 | 0,352 | 0,789 | 1,30E-06 |
| Pltp | 8.276236989 | -1,3749873 | 0,055 | 0,289 | 0,00153797 |
| Atp5b | 3.569804996 | -1,3807513 | 0,141 | 0,589 | 6,63E-11 |
| Itgb2 | 1.173406730 | -1,3940872 | 0,286 | 0,622 | 0,00021805 |
| Trp53 | 2.747147139 | -1,4128298 | 0,075 | 0,356 | 5,11E-05 |
| Ptms | 2.972301284 | -1,4138878 | 0,236 | 0,689 | 5,52E-08 |
| Aif1 | 6.288451336 | -1,4187858 | 0,101 | 0,422 | 1,17E-05 |
| Vim | 5.662670464 | -1,4980831 | 0,508 | 0,822 | 1,05E-05 |
| Ifi30 | 2.147479846 | -1,5664721 | 0,171 | 0,556 | 3,99E-07 |
| Ctsc | 6.427402910 | -1,648144 | 0,231 | 0,667 | 1,19E-08 |
| Fos | 1.028063915 | -1,6762401 | 0,181 | 0,489 | 0,00019105 |
| Sgk1 | 4.429250696 | -1,7005611 | 0,085 | 0,411 | 8,23E-07 |
| Cybb | 5.318379826 | -1,7019321 | 0,181 | 0,511 | 9,88E-06 |
| Unc93b1 | 5.376245728 | -1,771589 | 0,176 | 0,556 | 9,99E-08 |
| Selenop | 2.028931194 | -1,7796173 | 0,111 | 0,444 | 3,77E-06 |
| Cd9 | 4.314193017 | -1,8170676 | 0,136 | 0,433 | 0,00080171 |
| Ctss | 9.770784429 | -1,8564161 | 0,312 | 0,733 | 1,82E-07 |
| Mpeg1 | 1.009533888 | -1,9045984 | 0,216 | 0,533 | 0,0001876 |
| Psap | 9.374924000 | -2,0866997 | 0,261 | 0,811 | 1,74E-14 |
| H2-Eb1 | 1.963993863 | -2,1398063 | 0,216 | 0,9 | 3,65E-22 |
| Cd74 | 2.891301319 | -2,3157042 | 0,261 | 0,944 | 5,37E-24 |
| Ccl6 | 2.566269456 | -2,3329343 | 0,111 | 0,411 | 4,77E-05 |
| H2-Aa | 4.633566781 | -2,4984176 | 0,241 | 0,889 | 8,61E-22 |
| Ighm | 3.061009709 | -2,4998378 | 0,06 | 0,367 | 5,69E-07 |
| H2-Ab1 | 1.541717081 | -2,5315595 | 0,236 | 0,867 | 2,86E-20 |
| Lyz2 | 3.874250917 | -2,9425872 | 0,171 | 0,667 | 7,20E-14 |

Cluster 10

| | p_val | avg_log2FC | pct.1 | pct.2 | p_val_adj |
|---|---|---|---|---|---|
| Gzmb | 2.25730611З | 2,99907045 | 0,793 | 0,167 | 4,19E-08 |
| Ifi27l2a | 3.594792704 | 2,72044722 | 0,803 | 0,208 | 6,68E-09 |
| Isg15 | 7.403397734 | 1,93464709 | 0,697 | 0,146 | 1,38E-06 |
| Plac8 | 3.41883822 | 1,91409784 | 0,755 | 0,375 | 0,00063532 |
| Ifit3 | 3.16648906 | 1,74190625 | 0,585 | 0,125 | 0,00058843 |
| Usp18 | 1.23269950 | 1,59009954 | 0,574 | 0,042 | 2,29E-06 |
| Ifit1 | 1.76752972 | 1,36397572 | 0,537 | 0,104 | 0,0032846 |
| Irf7 | 2.09752352 | 1,30913093 | 0,803 | 0,396 | 3,90E-05 |
| Tox2 | 1.76390419 | 1,15940187 | 0,638 | 0,229 | 0,00327786 |
| Ifih1 | 7.89956863 | 1,13886926 | 0,622 | 0,062 | 1,47E-06 |
| Sytl3 | 2.91700078 | 1,12146807 | 0,771 | 0,396 | 0,00054207 |
| Xaf1 | 2.67628049 | 1,1030891 | 0,771 | 0,438 | 0,00049733 |
| Oas1a | 4.07933007 | 1,05674925 | 0,569 | 0,104 | 0,00075806 |
| Cd8b1 | 3.41672196 | 1,04363046 | 0,894 | 0,542 | 0,00634929 |
| Slfn1 | 3.656815261 | 1,040489 | 0,793 | 0,375 | 0,00067955 |
| Slfn8 | 6.75353215 | 0,90059955 | 0,846 | 0,479 | 0,00125501 |
| Mapkapk2 | 4.657570804 | 0,83667455 | 0,957 | 0,896 | 0,00865516 |
| Rbm3 | 3.686498714 | 0,8142944 | 0,91 | 0,75 | 0,00685062 |
| Kcnk1 | 2.93162677 | -0,5306597 | 0,069 | 0,333 | 0,00544784 |
| St3gal6 | 4.26813921 | -0,8117806 | 0,537 | 0,833 | 0,00079315 |
| Axin2 | 1.49374607 | -0,8191719 | 0,085 | 0,458 | 2,78E-06 |
| Qpct | 5.133510567 | -0,8199451 | 0,213 | 0,542 | 0,0095396 |
| Ikzf2 | 7.888455262 | -0,8941623 | 0,484 | 0,833 | 0,00014659 |
| Sdc1 | 2.12797572 | -1,1492712 | 0,021 | 0,25 | 0,00039544 |
| Tcf7 | 2.67897083 | -1,2178471 | 0,191 | 0,521 | 0,00497833 |
| Tmem64 | 1.603499657 | -1,3954972 | 0,479 | 0,854 | 2,98E-07 |

Up in Sp140-/-
Up in Sp140+/+
